# kASA: Taxonomic Analysis of Metagenomic Data on a Notebook

**DOI:** 10.1101/713966

**Authors:** Silvio Weging, Andreas Gogol-Döring, Ivo Grosse

## Abstract

The taxonomic analysis of sequencing data has become important in many areas of life sciences. However, currently available software tools for that purpose either consume large amounts of RAM or yield an insufficient quality of the results.

Here we present kASA, a *k*-mer based software capable of identifying and profiling metagenomic DNA sequences with high computational efficiency and a user-definable memory footprint. We ensure both high sensitivity and precision by using an amino acid-like encoding of *k*-mers with a dynamic length of multiple *k*’s. Custom algorithms and data structures optimized for external memory storage enable for the first time a full-scale metagenomic analysis without compromise on a standard notebook.

## Introduction

Decoding the complex composition of microbial communities is important, for example, to determine Earth’s biodiversity or its role in human diseases. However, this poses a major challenge. One of the biggest problems is the sheer number of different organisms living together in a biotope. Most organisms thrive only in the community of other organisms so specific cultivation of single species is usually impossible. Microbial communities therefore must be analyzed as a whole which is done by examining the totality of the genetic material in the community: the metagenome [1]. The technology of Next-Generation Sequencing (NGS) offers the possibility of creating vast amounts of DNA sequences (so called ‘reads’) from the metagenome, usually several million per dataset, each with a length between tens and hundreds of nucleotides [2]. The processing and taxonomic identification of these reads requires special bioinformatics methods.

In recent years, several software tools that are capable of comparing reads to a database containing genomic sequences of known organisms have been published [3]. Apart from the investigation of microbial communities, most of these tools can also be used to detect and quantify contaminations in any sequencing data, so they can be extremely valuable for data quality assurance in most application areas of NGS.

One of the best known programs for comparing sequences to large databases is MegaBLAST [4], which uses a seed-and-extend heuristic for finding local alignments between a read and the database. While this tool has a high usability and accuracy, it is not very well suited for quickly processing large numbers of sequences, since the analysis of complete NGS datasets using MegaBLAST would require high computational effort. Other tools like Kraken [5], KrakenUniq [6], and Clark [7] use a more time-efficient approach called *k-mer sampling* that compares short sub-sequences of a fixed length *k* (*k*-mers) taken from the reads with a pre-computed index derived from the database. Kraken uses a lowest-common-ancestor (LCA) approach to infer taxonomic memberships, whereas Clark needs a fixed taxonomic rank beforehand. These methods allow very fast identification but require huge amounts of random-access-memory (RAM), sometimes more than 100 GB, especially for comprehensive databases. Other solutions try to balance between time, accuracy and memory consumption like Centrifuge [8], Kraken2 [9], mash [10], sourmash [11], MetaCache [12], Ganon [13], Kaiju [14] and many more. However, due to the still growing number of reference genomes in, e.g., the NCBIs nucleotide sequence database [15], the amount of primary memory required by almost all of these tools finally scales beyond the scope of a conventional notebook. This means that the user is forced to rely on special and expensive hardware to perform metagenomic analysis.

In this paper we introduce kASA (<u>*k*</u>-mer <u>A</u>nalysis of <u>S</u>equences based on <u>A</u>mino acid-like encoding), a robust, accurate and deterministic *k*-mer based tool for the analysis of metagenomic sequencing data with a customizable memory footprint. The low memory requirement is achieved by an index that does not need to reside completely in the primary memory (RAM) but can remain largely on secondary memory like hard disk or solid state disk (using data structures from [16]). With this feature, kASA is the first software tool that is able to analyze large metagenomic datasets with superior accuracy on a standard commercial notebook within a reasonable period of time, even with large, comprehensive databases. kASA is written in `C++`, free, open source and available for all mainstream operating systems (Linux, Mac, Windows) and platforms (Desktop, HPCC, Notebook).

Unlike other *k*-mer based tools, kASA constructs *k*-mers using an amino acid-like encoding by converting triplets of nucleotides – regardless of coding or non-coding – via a translation table that corresponds to the genetic code. This way, storing *k*-mers requires less space (five bits per nucleotide triplet instead of six), and in addition the sensitivity is improved as synonymous DNA mutations (i.e. those that do not affect the encoded amino acid) no longer affect the matching process. The precision is maintained since differences at nucleotide level typically lead to differences in the corresponding amino acid sequence in at least one of the three possible reading frames. As long as there are differences at nucleotide level, kASA can therefore very well distinguish between, for example, homologous protein sequences in different species.

Another feature of kASA is the use of an entire range of word lengths *k*. Most other *k*-mer based methods use only a single *k* and thus have to make a singular decision how to the balance between sensitivity (small *k*) and precision (large *k*). With kASA the user can specify a small and a large *k* (12 is the current maximum) and the tool will try to match all word lengths of this closed interval. This interval can be changed each time the tool is run.

By dynamically adapting the word length *k*, kASA is able to optimize both sensitivity and precision. If a *k*-mer is too short for being specific for a particular organism it may become specific with longer word length. On the other hand, if a longer *k*-mer cannot be found in the index due to a mutation, it could possibly still be found when using a shorter word length. The index is designed in a way that its size does not increase when using multiple word lengths.

These capabilities allow kASA to compete with or even outperform other tools.

## Results and discussion

### Pre-Processing

The comparison of NGS data with a database can be significantly accelerated by pre-processing the database into an index data structure. This works as follows:

The DNA sequences from the database are scanned in three or six reading frames (depending on user choice) and converted and encoded via a given table to amino acid-like sequences, including coding and non-coding regions. This translation is lossless except for the first and last base (see supplementary file A). From the translated sequences all overlapping 12-mers (i.e. *k* = 12 amino acids, the maximum *k* supported by kASA) are extracted and saved into a file stored in secondary memory, together with the taxonomic ID of their source. After sorting and merging redundant entries (those, who share the same 12-mers and taxonomic ID), the index is ready to be used for identifying or profiling NGS data. Note, that this implies having multiple taxa for the same *k*-mer is allowed. In order to avoid having to rebuild the index every time the database is changed, kASA supports the addition of new reference sequences to the existing index without rebuilding.

In short the processing of sequencing reads works as follows: First the reads and their reverse complements are converted in the same manner as the database sequences. Then a set-intersection-like algorithm is used to find matches between *k*-mers from reads and the index, starting with the smallest *k*. If a hit is found, *k* is increased until the *k*-mers stop matching or the maximum *k* is reached. Any match is marked and scored. A more detailed description of these algorithms can be found in the Methods section.

### Taxonomic identification of simulated reads

To evaluate the accuracy and performance of kASA in relation to current state of the art tools, we designed a synthetic benchmark with snakemake [17] pipeline (see https://github.com/SilvioWeging/kASA_snakemake). This benchmark uses the largest chromosomes (or whole genomes in case of the prokaryotes) of multiple model organisms as a reference. From this reference, reads of length 100 are randomly sampled and randomly mutated via a mutation percentage (up to 20%). This simulates the problem of identifying new species which are related to known ones and tests the robustness of every tool. Results are shown in Figure 1. At the same time, the memory consumption during index building and identification as well as the run times are logged and shown in Figures 2 and 3.

**Figure 1:**
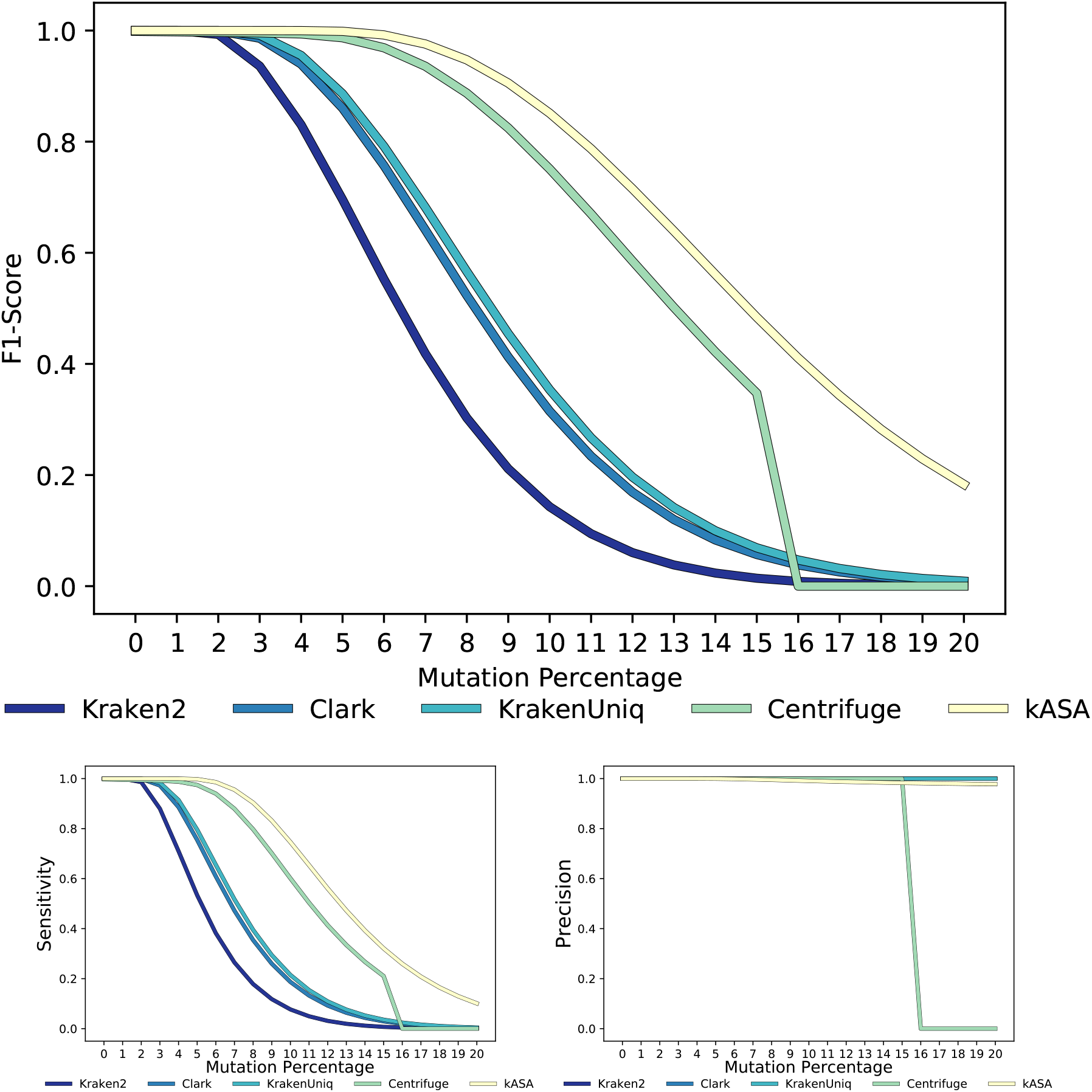
F1-Score (top), sensitivity (bottom left) and precision (bottom right) of tested tools for simulated data. kASA is shown with a lower *k* of 7 (default).

**Figure 2:**
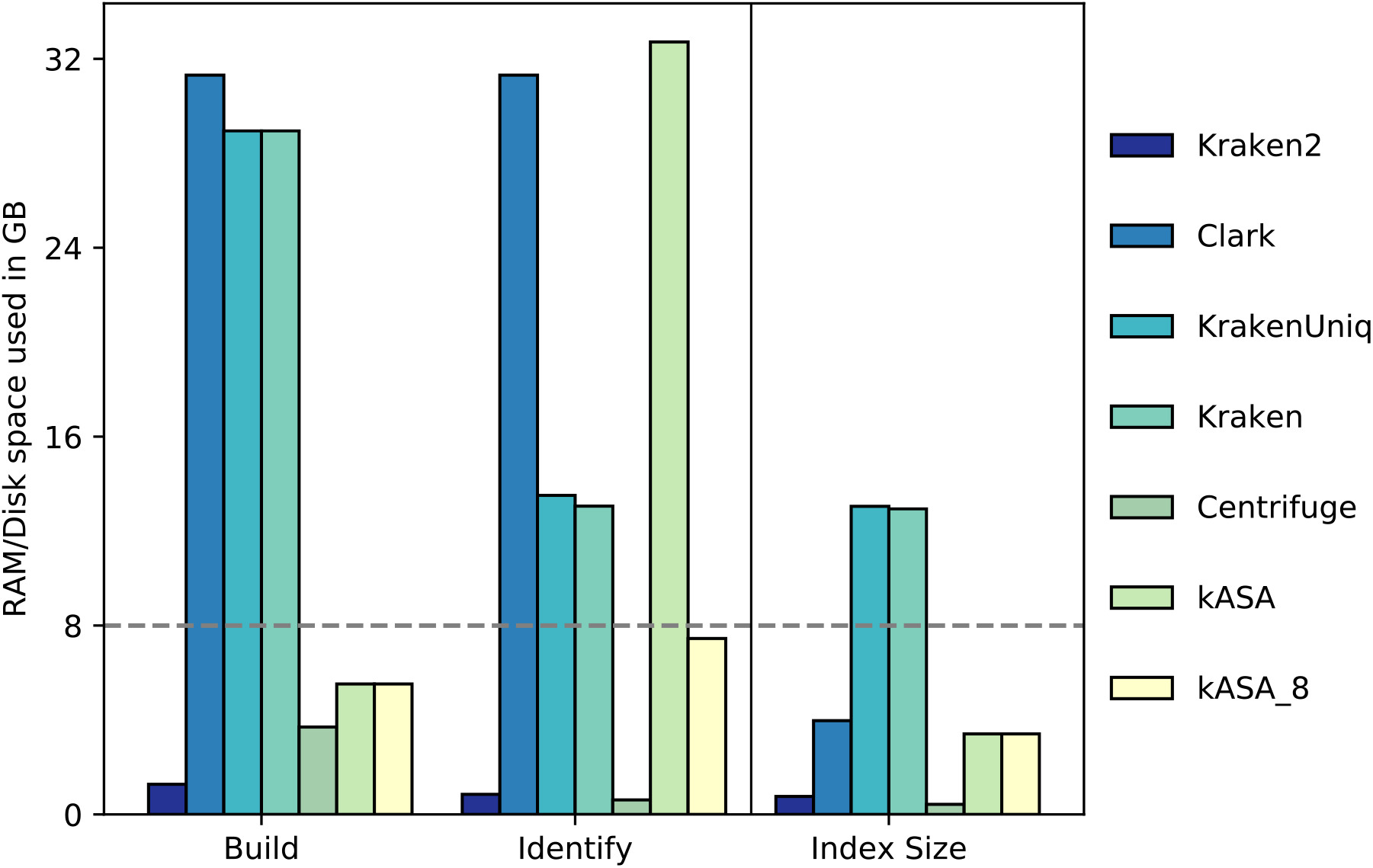
Memory and space, which is used to build the index or identify the test data, and sizes of indices. Colors and order are equivalent to the ones in Figure 1. kASA is shown in two ways: without RAM restrictions for index building and identification and with 8GB as limit with a parameter. The dashed gray line sets a boundary which tool would be able to complete this test on a system with 8GB RAM.

**Figure 3:**
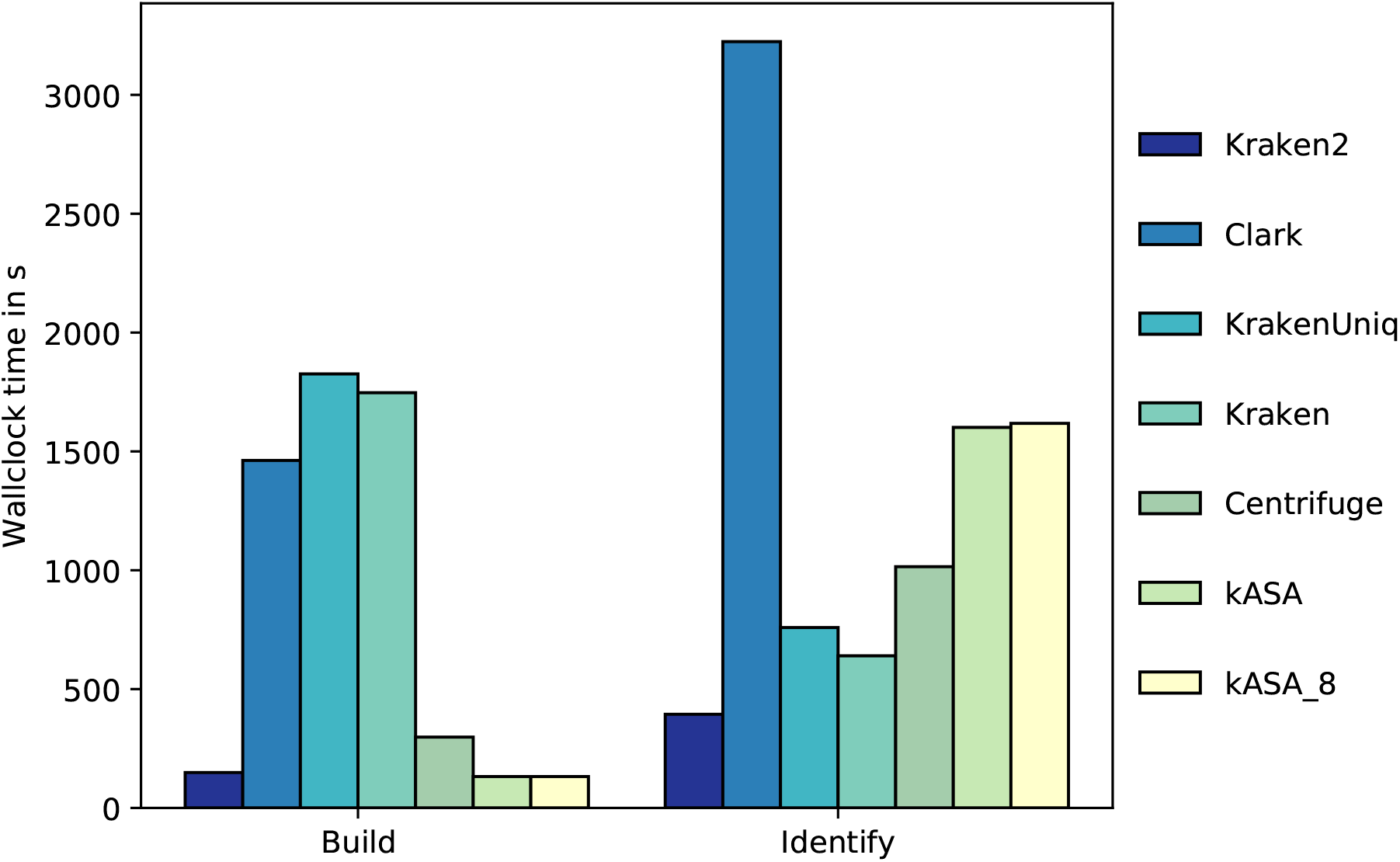
Wall clock time for the accuracy tests of kASA and other tools. Times were gathered with the benchmark option in snakemake and sorted in the same manner as in Figure 1. The plots y-Axis is marking seconds (s). Omitting read identification and creating only a profile speeds kASA up by a factor of ~1.3. Detailed measurements can be found in additional file C.

While the pipeline can be used with other organisms and tools, we would like to refer the reader to a more comprehensive benchmark given in: [18].

Detailed descriptions and tables of our evaluation methods, the tools used and their versions as well as results for different *k*’s and specificity can be found in the supplementary files A and B.

The plots in Figure 1 show the F1-Score, sensitivity and precision of selected tools for different mutation percentages. As expected, the values decrease non-linearly and tools allowing gaps perform better than exact matching algorithms with the exception of kASA. Because Kraken and KrakenUniq have exactly the same values only one of them is included in the plots. Centrifuge drops inexplicably to zero at 16% mutations per read. Our tool achieves superior accuracy even without explicitly modeling gaps since it is a *k*-mer based approach with dynamically adjusted word lengths. This ensures, that for a small enough *k* the non-mutated part matches. On top of that, we abstract to the amino acid-like level which makes at least one frame not likely to be affected by a single mutation due to robustness against synonymous mutations.

Because kASA was designed with minimal memory requirements in mind, we examined the memory consumption of each tool during index building and identification, as well as its index size. Results show (see Figure 2), that with standard settings only Kraken2, Centrifuge and kASA with a lower memory setting would be able to build an index and identify data on a standard notebook with 8GB RAM. However, this statement turns false when using more genomes as reference. Kraken2 can compensate this with a parameter limiting the index size but sacrificing sensitivity while Centrifuge exits with an error and the suggestion to run it on a system with larger primary memory.

For the sake of completeness, we would like to mention that other mentioned tools not included in the benchmark have strategies to cope with memory restrictions as well. Unfortunately, this often either influences run times by a large degree or reduces accuracy.

In terms of speed, Figure 3 shows that kASA is the fastest tool for index creation and Kraken2 for identification. KrakenUniq sometimes takes a very long time building the index (we suspect a bug) and kASA as well as Clark take their time identifying. Reducing the memory limit to 8GB increases the run time of kASA only by a small margin (writing and reading to disk is the limiting part, the rest is parallelized). This speed is mainly due to the index fitting into primary memory. Using an HDD increases run times by a larger degree. We therefore recommend using an SSD if not enough primary memory is available. We are confident that further optimizations in kASA can reduce the run time using user feedback and time. Further run times e.g. with different *k*’s can also be found in additional file C.

### Taxonomic profiling of simulated data

A taxonomic profile is a list of all organisms that have been identified in a dataset together with their respective abundance. This abundance is estimated by normalizing the relative frequency against genome length and ploidy.

Consistent with other studies (e.g. [19]), we use the following method to calculate the relative frequency of each taxon: Given an identification of the reads, we determine the most significant taxon for each read and count the number of reads per taxon. We then divide these read counts by the total number of reads in the sample. This relative frequency is sufficient to compare the accuracy between tools, so we do not normalize against the genome lengths and ploidy.

Apart from this method, kASA also offers a direct procedure for creating profiles by counting matched *k*-mers per taxon. See here for more details.

If taxonomic profiling is the main goal of a study, it is possible with kASA to completely dispense of the identification of the reads since this speeds up processing by a factor of ∼1.3. Furthermore, we advise to shrink the index by discarding a fixed proportion of *k*-mers from all taxa in equal proportions. This further speeds up the data processing without making significant sacrifices in terms of accuracy. For example, a reduction of the index by 50% decreases the F1-Score in our experiments only slightly but increases performance by a factor of ∼1.4. A detailed investigation of the effect of different index sizes on the profiles can be found in supplementary file F.

Regarding the quality of taxonomic profiles, we evaluated kASA with the same data, methods and quality measures as used in a metastudy by Lindgreen et. al. [19]. Although we did not run any of the tools studied in the paper except Clark again, we recalculated all measurements to ensure consistency (for more details, see supplementary file D).

Figure 4 and 5 show that kASA is the best tool regarding log-odd scores and only slightly differs in value for the Pearson correlation coefficient. Relative frequencies for this comparison were gathered per read for consistency, as mentioned above, but profiles generated by a direct use of *k*-mer frequencies deviate only slightly (see supplementary file D sheet 2).

**Figure 4:**
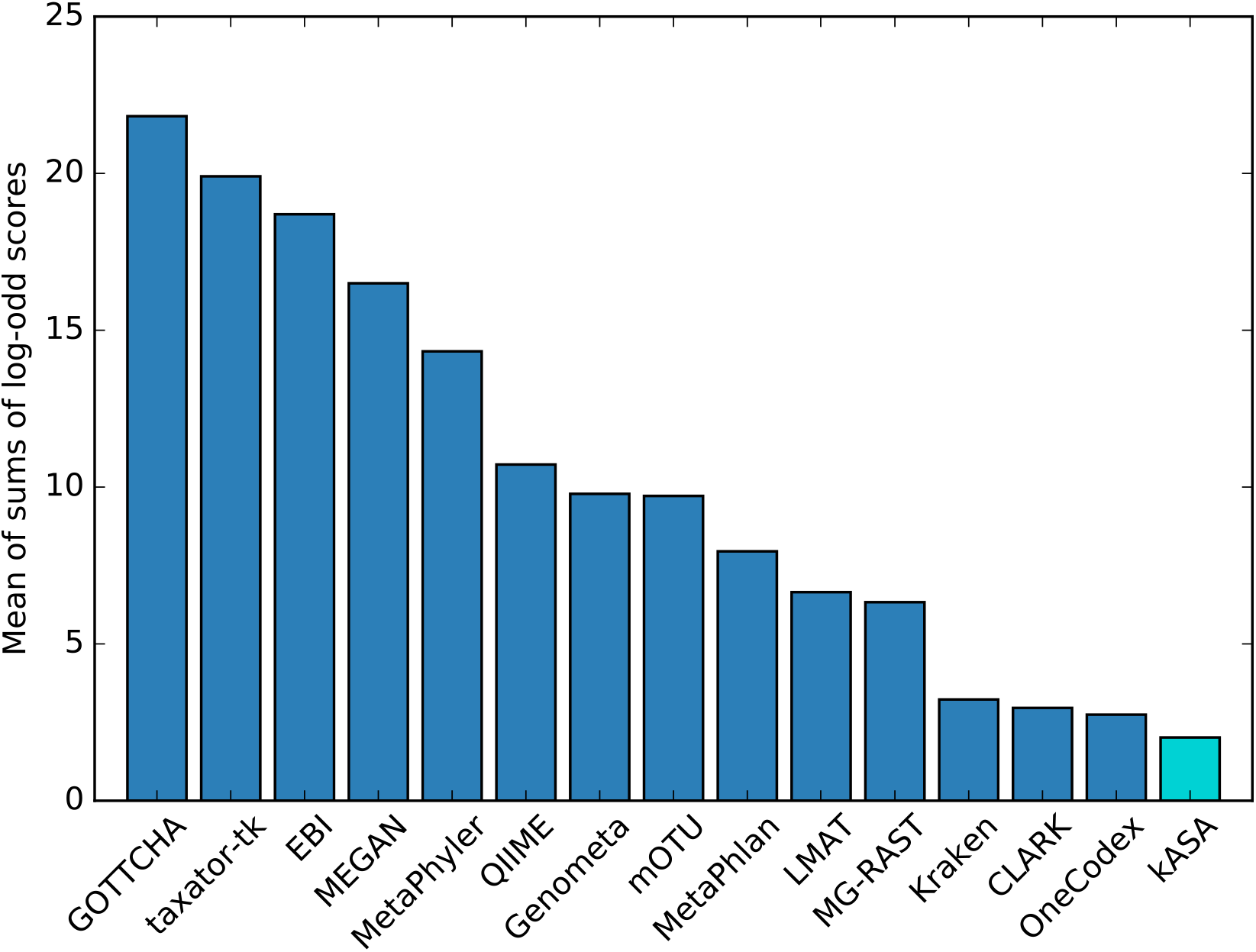
Profile quality measured by the mean of log-odd scores. The log-odds of absolute differences between relative frequency and gold standard were summed per dataset and averaged regarding all six datasets. Tools are sorted by this score meaning lower is better. Data from the corresponding publication [19] was also used in the publication of Centrifuge [8] where the authors wrote, that their accuracy was similar to that of Kraken. The newest version of Clark was used to verify its own results as can be seen in supplementary file D.

**Figure 5:**
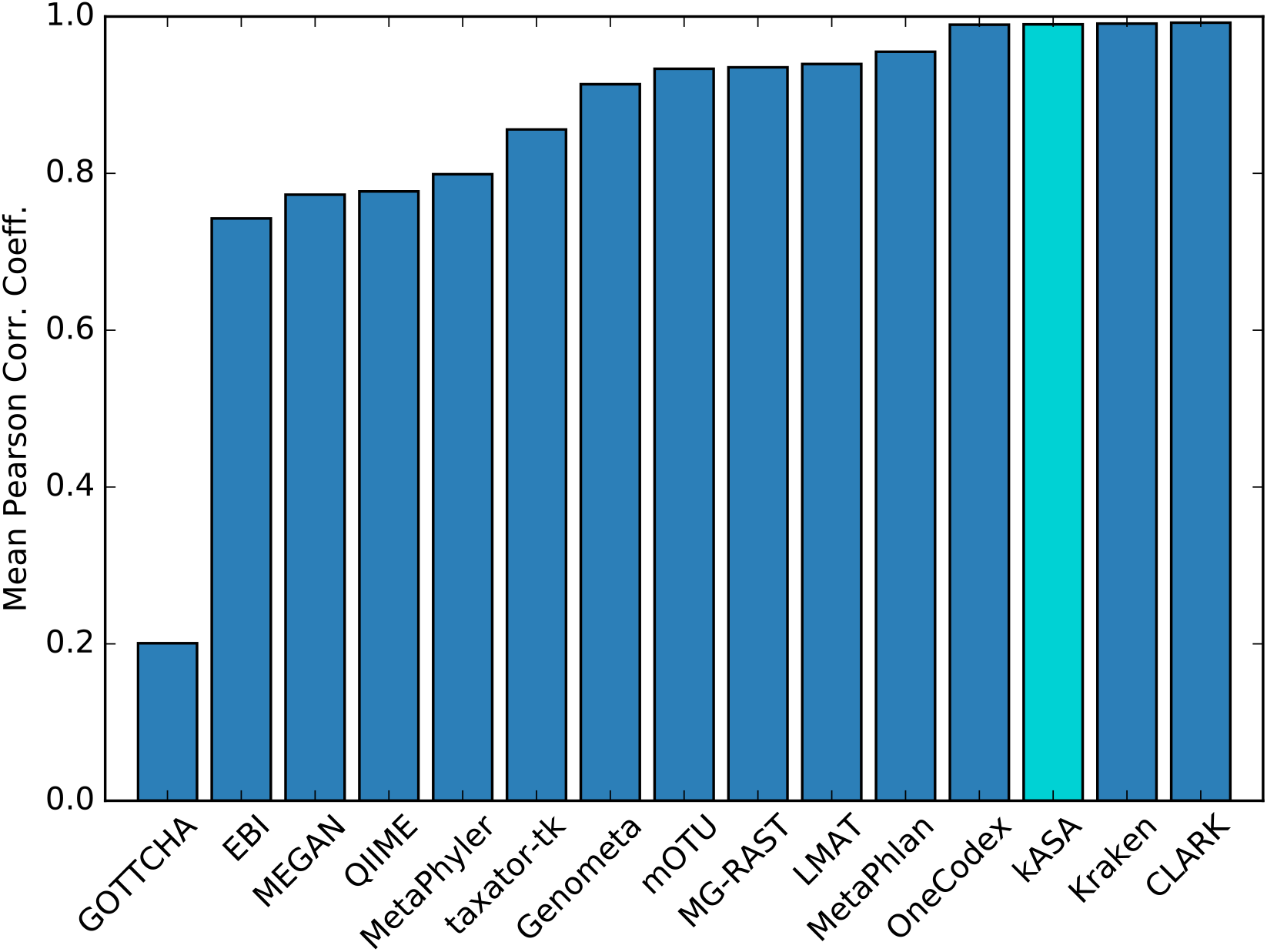
Profile quality measured by the mean of Pearson Correlation Coefficients. The Pearson Correlation Coefficients of vectors of relative frequencies in relation to the respective truth for every dataset were averaged and shown here. Tools are sorted by that value. Results for the best four tools differ only slightly.

The CAMI-Challenge [20] proposes another comparison principle. It measures accuracy not only by difference from a gold standard but also by correctly identified taxa at every level. Combined with OPAL [21] it creates a framework for profile testing which offers the opportunity to compare kASA with even more tools. OPAL treats additional entries in the taxonomic profile as false positives even if their percentages are very low. To address this, we set different thresholds in the profile on the species level. A threshold of 0.01 seemed to perform best while setting none at all performed worst regarding precision. We therefore recommend setting a cutoff value since any threshold performed better than none (see supplemental file E). Regarding the relative frequency used for the profile, the unique frequency, meaning the one relative to just the uniquely matched ones, is closest to the true frequency in the gold standard.

As shown in Table 1 and supplemental file E, kASA ranks first according to OPAL when using default settings. This indicates, that kASA could perform quite good in a CAMI-Challenge given the opportunity to discard entries with low frequency.

**Table 1:**
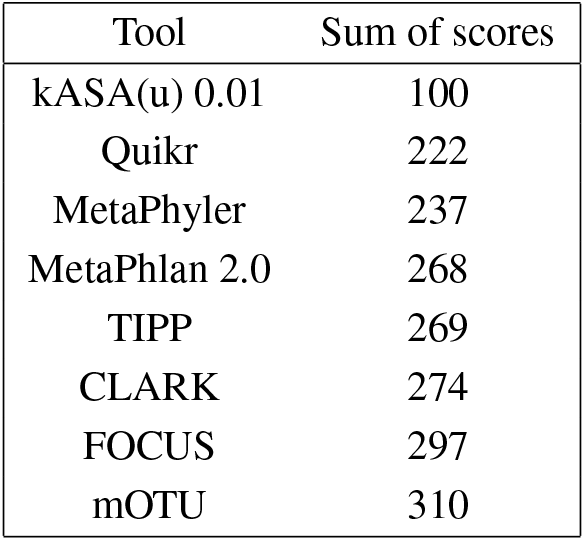
Sum of scores given by OPAL for the “CAMI low” dataset.

### Analysis of real data

To test kASA’s ability to identify known organisms in real data, we used data from the Human Microbiome Project [22] to create taxonomic profiles and compare the results with those created by Kraken2 [9] equipped with the same database. We chose Kraken2 because it was one of the few tools from our selection which was able to build an index that large (105 GB) on our system and finish in a reasonable time.

Table 2 shows that both tools agree in the most abundant genera although in different proportions depending on unique or non-unique counting. All detected genera are known to occur in the human oral flora which verifies our results. On phylum level both tools predict very similar proportions as shown in supplemental material G.

**Table 2:** Selection of genera with the highest relative frequency (in %) found in data sampled from human saliva (SRS147126). Results for kASA are shown with unique and non-unique values for *k*=12.

| Scientific Name | kASA u. | kASA n.-u. | Kraken2 |
| --- | --- | --- | --- |
| Prevotella | 43.5 | 27.9 | 30.7 |
| Streptococcus | 9.9 | 16.5 | 13.5 |
| Neisseria | 4.8 | 15.2 | 11.5 |
| Veillonella | 7.8 | 7.7 | 7.7 |
| Porphyromonas | 9.2 | 6.6 | 6.8 |
| Haemophilus | 3.9 | 5.1 | 4.5 |
| Campylobacter | 2.8 | 1.6 | 1.4 |

kASA can process reads of arbitrary length, so we were able to also analyze data from third generation sequencing technologies, namely from the Nanopore-Whole-Genome-Sequencing-Consortium [23] who used the Oxford Nanopore MinION technology [24] to sequence data from the GM12878 human cell line. Three samples were downloaded and analyzed. kASA detected, as expected, the human genome as well as phylogenetic relatives (but with much lower counts) and the Epstein-Barr virus. This leads to the conclusion that no contaminants were sampled which could have influenced an assembly. The taxonomic profiles are given in the supplemental material I. Note, that most other taxa appearing in the profiles only have relatively low unique *k*-mer counts and are therefore likely artifacts due to sequencing errors or short similarities.

## Conclusions

Our tests show that kASA performs taxonomic analyses with excellent sensitivity and precision. The use of a dynamic *k* instead of a fixed word length proves to be a valid method for improving sensitivity without sacrificing precision. The abstraction to an amino acid-like encoding further increases the robustness of kASA to mutations without the need for sequence alignment with gaps. This shows that our approach can compete with or even exceed commonly used approaches.

In addition, kASA is, to our knowledge, the only available tool providing a RAM consumption so highly adjustable that it also works on cheaper computers at reasonable speed and without loss of accuracy. It therefore allows scientists who do not have access to expensive hardware to run their analysis without much effort or help from other sources.

## Methods

### Index creation

As mentioned before, pre-processing a database to create an index ensures that only necessary calculations are performed while identifying or profiling NGS data. This step (named ‘build’ in kASA) converts a nucleotide database in fasta-format to a binary file containing *k*-mers and saves it to secondary memory, e.g. the hard drive. The index building step in kASA is depicted in Figure 6.

**Figure 6:**
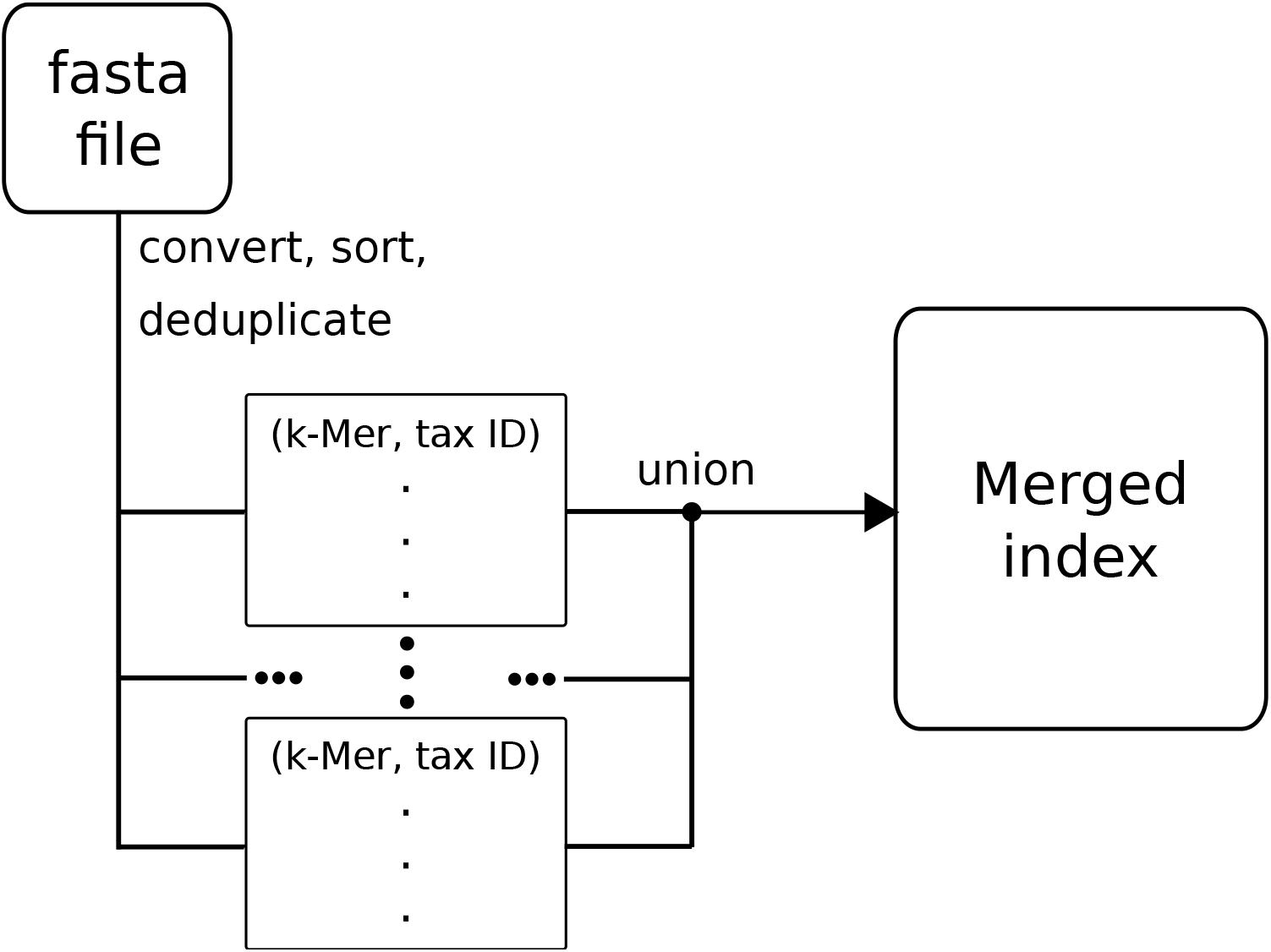
Flowchart for creating the index. In the first step, as much DNA from a fasta file as possible (determined by the memory restricting parameter) is transformed and stored into a container of *k*-mers which is stored onto the hard drive. If there is any DNA left in the file this step is repeated and at the end, the resulting containers are merged into the final index.

Accessing this file during a comparison would create a lot of slow I/O operations, even when using an SSD. To keep the number of these operations minimal and at the same time enable free choice of the values for *k*, we build a prefix trie [25] from the index containing the first six coded letters of each *k*-mer. Each leaf of this trie contains two integers representing the upper and lower boundaries of the range containing *k*-mers that share the same prefix. If this prefix is matched, only a fraction of the index must be searched for the remaining suffix. This significantly reduces the search space and since the trie is small enough to fit into primary memory, prefix matches are performed very quickly. Should the user provide more than 10GB of RAM and a lower *k* of at least 7, a much faster lookup table will be used instead.

Another important step in the construction of the index is the linking of DNA and its taxonomic ID. In order to recognize the origin of the DNA forming the nucleotide database, we use accession numbers from the fasta files and the NCBI taxonomy database. If there are no accession numbers or taxonomic IDs available, a dummy ID is given and the user receives a notification. The resulting so called “content file” is human readable and can be manually modified if necessary.

### Identification and Profiling

After building the index, the identification of NGS data can be performed. The identification algorithm is as follows: First, the DNA and its reverse complement, or already converted amino acid sequences, from a fastq or fasta file are (translated and) converted into *k*-mers in a similar way as in ‘build‘. While this is mainly an algorithmic necessity for the following algorithm, this method resembles “double indexing” [26, 27]. The difference to the building step is that each *k*-mer receives a read ID instead of a taxonomic ID and that duplicates are kept because they might hint at important motifs. For this reason, we advise the user and reader to de-duplicate reads beforehand to not distort abundances. The pairs are then sorted and passed to the identification algorithm to see which taxa match to which read. An overview of this process can be seen in Figures 7 and 8. A pseudocode of the algorithm can be found in additional file A.

Note, that if kASA is given a RAM limit then some input files may be too large to be processed at once so the files are read and processed in chunks.

**Figure 7:**
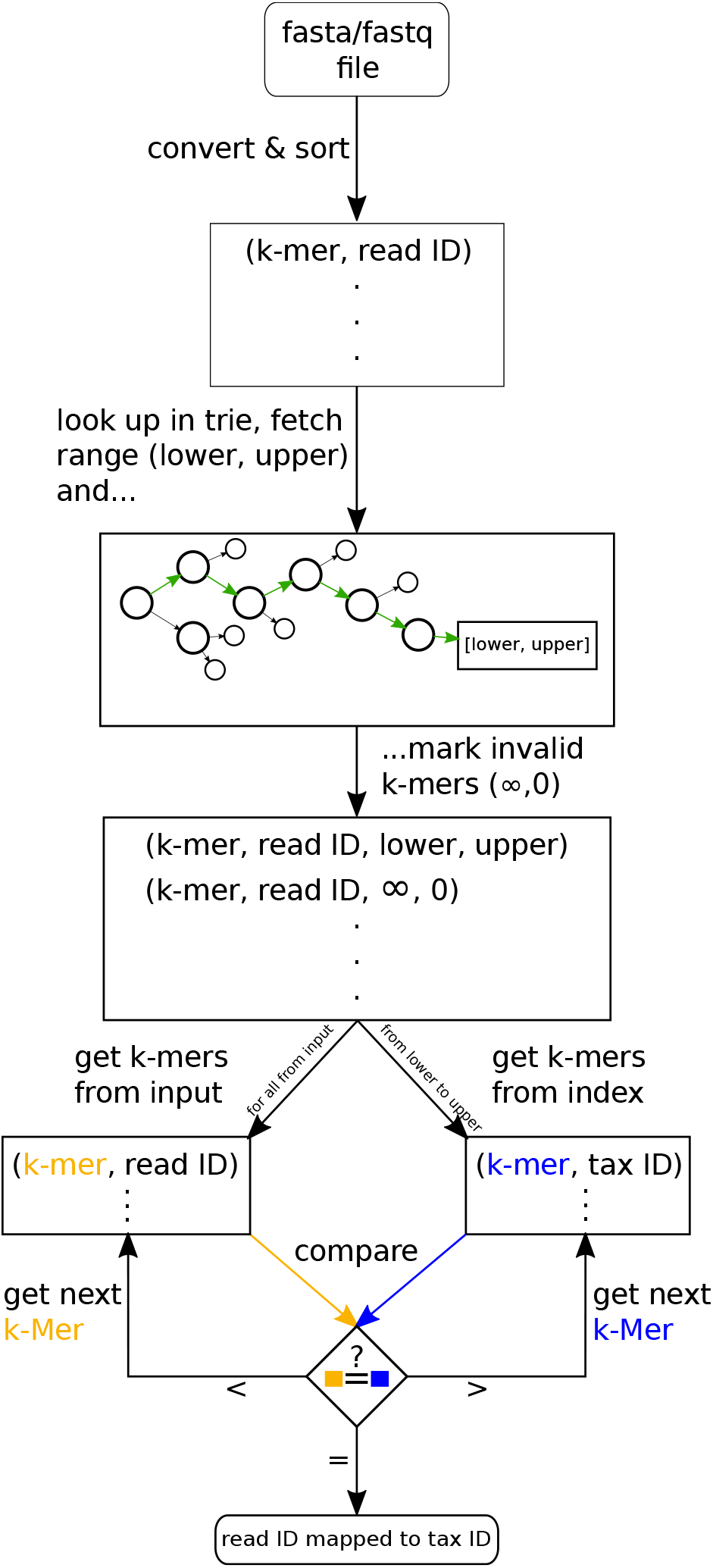
Flowchart of the identification algorithm. The DNA and its reverse complement are converted into pairs of *k*-mer and read ID. After sorting, the prefix of the first *k*-mer determines the suffix range over the precalculated index. From this range, the first *k*-mer is compared with the one from the input. If they match, the read ID and the taxonomic ID are scored accordingly. If not, either the next *k*-mer from the input or the next index’ *k*-mer is used. This loop continues until all *k*-mers from the input have been processed.

**Figure 8:**
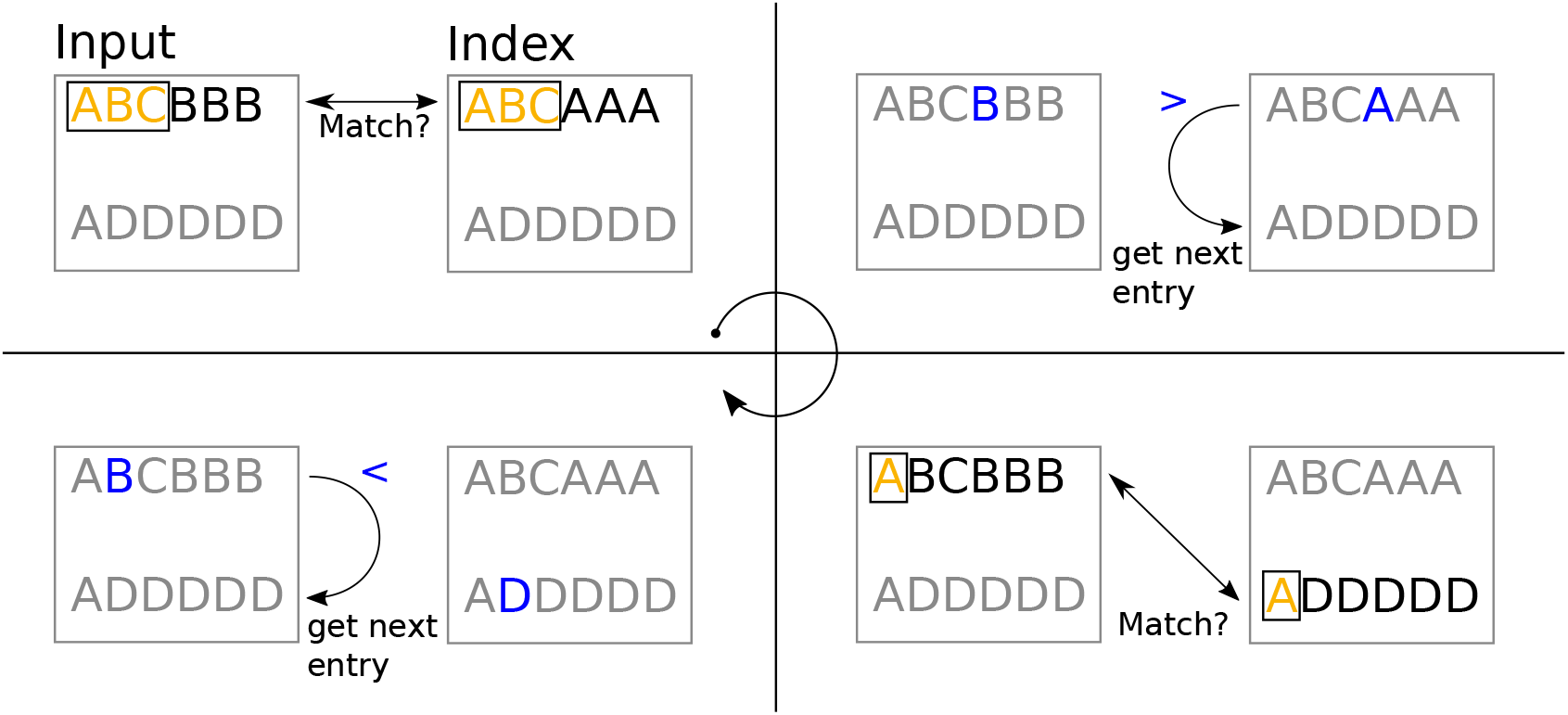
Matching algorithm. First, the *k*-mers are matched for *k* from lowest to highest. In case of a mismatch, either the pointer at the index or the input is iterated further. *k* is set to lowest again and the cycle repeats. A figure with more cases is shown in additional file A.

The scores in the identification algorithm are calculated as follows: Every read which shares at least one *k*-mer with the index will be noted to create a relation from read ID to taxonomic ID. Since matches of smaller lengths are less significant than longer ones, this relation is represented with a weighted sum. Weights *w_k_* are gained by a normalized quadratic function to stress a non-linear increase of significance and to strike a balance between rewarding long matches but not devaluing short matches too much. This means, that the values of *k*^2^ are normalized to (0, 1] for *k* = 1, …, 12. The sum using those *k*-dependent weights is called: *k*-mer Score of the taxon *t* ∈ *T* and read *r* ∈ *R*

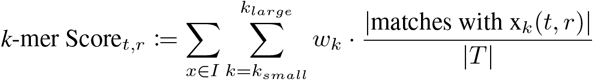

where *I* denotes the *k*-mers from the Input, |matches with *x_k_*(*t*, *r*)| is the number of *k*-mers that *t* and *r* share and T and R are the sets of matched tax or read IDs.

Additionally, a second score called ‘Relative Score’ is calculated before writing the results into a file. It puts the *k*-mer Score in relation to the total number of *k*-mers of a taxon *t* present in the index (also called frequency of *t*) and the length of the read *r*:

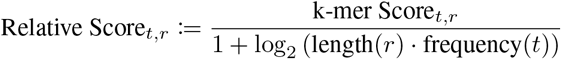

This formula is inspired by the calculation of the E-value in BLAST although in this case, a higher score indicates a better hit. The Relative Score can be used to determine the significance of a matched taxon. From our experience, for a read of length 100 everything with a Relative Score smaller than 0.4 can be seen as insignificant and for example sorted out during decontamination. For the output, the resulting scores are sorted in decreasing order by this relative score so that a read can have multiple identified taxa but the leftmost one has the highest value.

The taxonomic profile that kASA computes consists of the names, taxonomic IDs and relative frequencies as well as the number of matched *k*-mers of all taxa found in a dataset. The number and relative frequencies are printed for each *k* and for unique and non-unique matches as well as relative to all *k*-mers in the input. This is necessary since the database used to form the index can be redundant for several reasons: For example an entry is named differently although it is identical to an existing one, or a subspecies is identical to another one on the amino acid level. kASA also offers a value describing the degree of redundancy or ambiguity present in the index which helps deciding which frequency to use. Note however, that for comprehensive databases this value may be overshadowed by large genomes.

Unique relative frequencies are calculated by dividing the number of uniquely matched *k*-mers for a taxon t by the total number of uniquely matched *k*-mers:

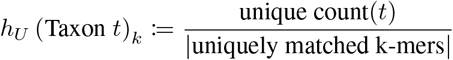

If a *k*-mer *x* matched multiple taxa, the hit count (how often that *k*-mer matched) is divided by the number of matched taxa. The sum of these floating point numbers is divided by the total amount of matched *k*-mers at the end. The formula is as follows:

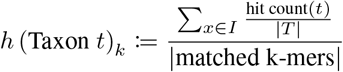

If for example *x* was found five times in three different taxa, each of these three would receive 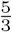 to its dividend.

The so called “overall relative frequency” takes the same numerator as the non-unique relative frequency but divides it by the total amount of *k*-mers in the input file. This is particularly useful when a file contains a lot of unknown DNA (from the index’ point of view), as it prevents over-interpretation of high-ranking taxa in the other relative frequencies.

In conclusion, the identification file offers an answer to the question which organisms were found for each read, creating a basis for further studies. The taxonomic profile provides a broad overview of which organisms are present in the NGS dataset and how much DNA was contributed by them.

## Materials

The database used for the experiments in [19] contained species which were no longer inside our version of the NCBI taxonomy and nt database. We manually fetched these entries via the E-utilities from the NCBI [28] and created a custom ‘content file‘ containing the respective links to the taxonomic IDs. It can be found together with the results in supplementary file D.

Genomes came from the Refseq database as of2019-10-01. This database (ca 1.1 TB in size) and index (1.4 TB in size for kASA) were used for profiling the real data in Table 2, additional file I, and Table 1. To ensure maximum performance of kASA, the indices were shrunken via methods described in the README file which can be found in our GitHub repository (e.g. the large index by 50% during building and then lossless in half afterwards). All tests were conducted on an HPCC for a comparative basis, specifications can be found in additional file A.

## Abbreviations

RAM: Random Access Memory
LCA: Lowest Common Ancestor
SSD: Solid State Drive
HDD: Hard Disk Drive
I/O: In- and Output
NGS: Next Generation Sequencing
HPCC: High Performance Computing Cluster

## Declarations

### Acknowledgements

We thank Jan Grau and Matthew T. Weirauch for valuable conversations and Yvonne Poeschl-Grau for recognizing problems where we least expected them. We would also like to thank the German Centre for Integrative Biodiversity Research (iDiv) Halle-Jena-Leipzig for their support, especially in the initial phase of the project.

### Availability of data and materials

kASA is freely available on https://github.com/SilvioWeging/kASA and licensed under Boost Version 1.0. The Snakemake pipelines are available on https://github.com/SilvioWeging/kASA_snakemake.

### Author’s contributions

SW and AGD designed the algorithms, SW implemented and tested them. Experimental design and test data creation was performed by SW and IG. SW, AGD, and IG wrote the manuscript. All authors read and approved the final manuscript.

### Ethics approval and consent to participate

Not applicable.

### Consent to publish

Not applicable.

## Additional Files

**Additional file A – Formulas for sensitivity, precision and specificity, versions of used tools as well as system specifications and further plots. Also the proof of our almost lossless coding.**

**Additional file B – Table of results for simulated experiments**

**Additional file C – Table with results from run time and memory consumption tests.**

**Additional file D – Table with results from the Lindgreen et. al. quantification test.**

**Additional file E — HTML file created by OPAL to visualize the results for the first CAMI Challenge**

**Additional file F – Study of accuracy with shrunken indices.**

**Additional file G – Results of Kraken2 and kASA from data sampled from human saliva.**

**Additional file H — HTML file created by Krona [29] for results of data from human saliva**

**Additional file I – Profiles of data from the Nanopore-Whole-Genome-Sequencing-Consortium**

