## Additional file A for "kASA: Taxonomic Analysis of Metagenomic Data on a Notebook"

### Supplemental file 1 for kASA

Silvio Weging

May 20, 2020

#### Contents

**1 Evaluation of synthetic data** **2**

**2 Pseudocode of the identification algorithm** **6**

**3 Versions and system specifications** **11**

**4 Information loss of the amino-acid-like encoding** **12**

### 1 Evaluation of synthetic data

We evaluated the identification quality for our synthetic tests from two perspectives: Read and Taxon.

The former is done by checking whether the original taxon of a read was identified correctly by ID. If the ID matched the one defined for that read, it was marked as a correctly identified and assigned read otherwise only as assigned (decreasing both sensitivity and precision). Only the best hits were considered. For kASA the json array for every read containing the "Top hits" was used. This array is calculated by normalisation of the $k$-mer scores to $[0, 1]$ and including everything with a normalised value $\geq 0.8$. This value seemed to correspond best to what would intuitively be considered "relevant" when reporting several results for one read. If two or more taxonomic IDs matched the read or the LCA-based algorithms gave a higher taxonomic rank as result (while containing the correct ID) the read was additionally considered ambiguous but correctly assigned. If the taxonomic path given by backtracking the LCA-path did not contain the correct ID, it was considered incorrectly assigned. We added genomic reads from species not inside the database/indices to test every tools ability to "ignore" reads. This is measured with the specificity via checking, if a nonassignable read was correctly not assigned. The formulas are as follows:

$$\text{Sensitivity} := \frac{|\text{Correctly assigned reads}|}{|\text{Reads}|}$$

$$\text{Precision} := \frac{|\text{Correctly assigned reads}|}{|\text{Assigned Reads}|}$$

$$\text{F1 score} := 2 \cdot \frac{\text{Sensitivity} \cdot \text{Precision}}{\text{Sensitivity} + \text{Precision}}$$

$$\text{Specificity} := \frac{|\text{Correctly unassigned reads}|}{|\text{Nonassignable Reads}|}$$

Because of this setup, tools reporting everything from their index with the same score would get an artificially high F1 score. To counter this, we added the perspective of each taxon which can be done via a binary classification. The "original read taxon" is the one we know, the "reported taxon" is the one the tool returns for that read.

- True positives (TP) - The original read taxon did expect the reported taxon and got it.

- True negatives (TN) - The original read taxon did not expect the reported taxon and it was (correctly) not reported.
- False positives (FP) - The original read taxon did not expect the reported taxon but got it anyway.
- False negatives (FN) - The original read taxon did expect the reported taxon but it was not reported.

With this, we can calculate the four values for every expected taxon and derive the Matthews correlation coefficient:

$$\text{MCC} := \frac{(TP * TN - FP * FN)}{\sqrt{(TP + FP) * (TP + FN) * (TN + FP) * (TN + FN)}}$$

We then averaged these MCCs for every file and got our measure how often a tool reports only what's necessary.

The evaluation was performed by a script for each output, because no standard for tool-outputs exists. Mutations include insertions, deletions and single point mutations all with the same probability. Indices where these mutations occur per read are drawn randomly via python. The seed is fixed for the purpose of recreating data. Read length was fixed to 100 and per genome, $\frac{|\text{bases}|}{100}$ reads were generated. Positions in the genome from which reads were sampled were also drawn randomly.

Please see our GitHub site for further details on how to recreate our results: `https://github.com/SilvioWeging/kASA_snakemake`.

The following figures show the results of our benchmark for all tested tools and $k$'s from 7 to 12 for kASA.

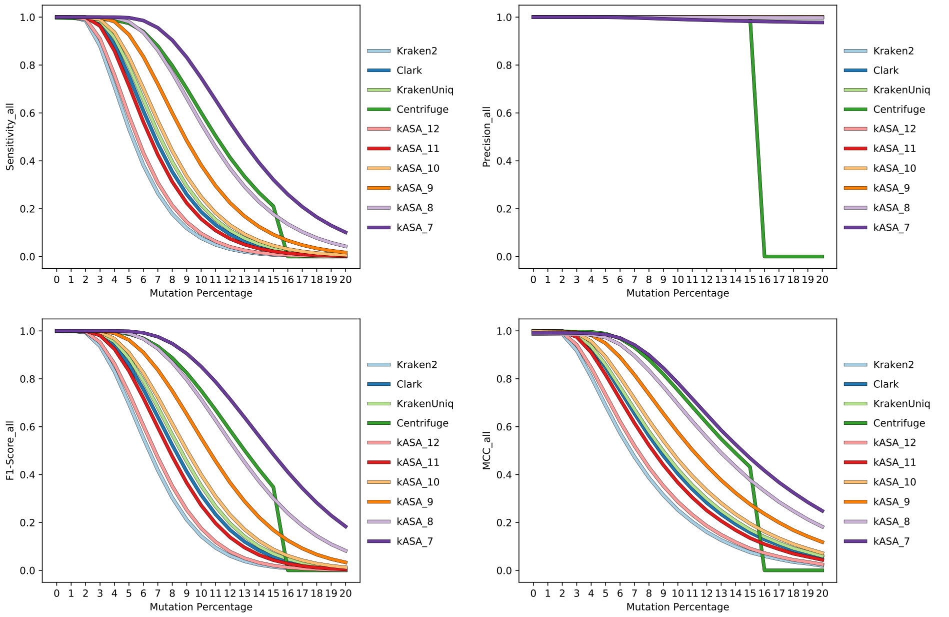

Since for a $k$ of 7, kASA performs best in the above figures, we tested how the specificity is affected by our approach. The next figures show, that it is inversely correlated with mutation percentage. This is due to short similarities with negatives in our benchmark. However, if a threshold is applied, specificity returns to acceptable values. For the test data with zero mutations, a threshold of 0.4 on the relative score was able to yield almost perfect specificity without lowering sensitivity or precision. However for data sets with a higher number of mutations, the ROC is still acceptable but sensitivity suffers visibly.

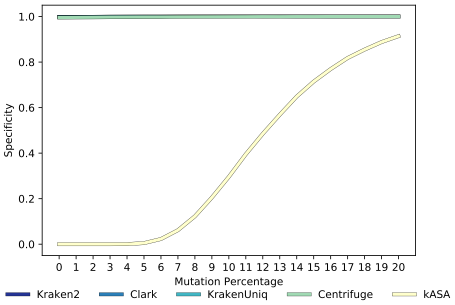


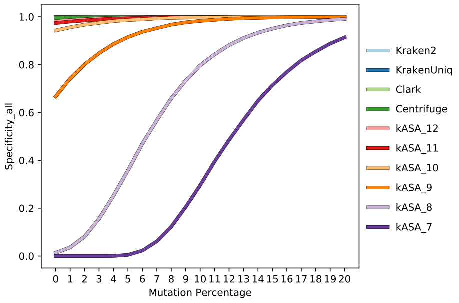


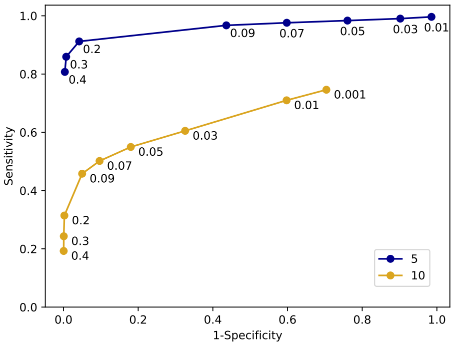

#### 2 Pseudocode of the identification algorithm

Customised set intersection algorithm used for a comparison of NGS data with the index. It computes both the profile and the identification file per read. After the pseudocode, an example is shown in 1 which displays the workings of the algorithm.

**Algorithm 1:** Identify

**input :** The index and the sorted input with converted $k$-mers and ranges
**output:** Scores and counts

```
// Part One
```
list of matched read IDs per k: *rIDs_k* ← [];
list of matched tax IDs per k: *tIDs_k* ← [];
list of last known k-mer per k: *known_k* ← [];

**for** Every *entry* with *range* in input **do**
    reset all lists;
    $k\text{-}mers_{[lower,upper]}$ = gather all k-mers with the same range;
    **for** All k-mers in $k\text{-}mers_{[lower,upper]}$ **do**
        $x$ ← current k-mer;
        **if** range invalid **then**
            `continue`;
        *currKMerShifted* ← x with lowest value for k;
        *start* ← *lower*;
        *end* ← *upper*;
        **do once**
            **if** *currKMerShifted* is in *range* **then**
                use binary search to find the *start*;
            **else**
                `continue`;
        **if** $x$ has been *seen before* **then**
            **for** all $k$'s **do**
                add *read ID* to *rIDs_k* if $x_k$ matches entry in $known_k$;
            `continue`;
        **else**
            *seen before* ← $x$;
        `// Part Two`
        `// see below`
    `// Part Three`
    `// see below`
**return** Scores and counts

```
// Part One
// see above
for Every entry with range in input do
    for All k-mers in k-mers[lower,upper] do
        // Part Two
        for y in index from start to end do
            for all k's from lowest to highest do
                x_k ← x shortened for this k;
                y_k ← y shortened for this k;
                if x_k < y_k then
                    for all remaining k's from k to highest do
                        add read ID to rIDs_k if x_k matches entry in
                          known_k but avoid duplicates;
                    break out of the y loop and get next x;
                else
                    if x_k == y_k then
                        if x_k matches entry in known_k then
                            add tax ID from y to tIDs_k and read ID to
                              rIDs_k but avoid duplicates;
                        else
                            // all possible matches for old x_k
                               found -> save
                            for all entries in tIDs_k do
                                count unique or non-unique match;
                                for all entries in rIDs_k do
                                    save match of read ID and tax ID;
                                reset rIDs_k and add the current read ID;
                                reset tIDs_k and add the current tax ID;
                                known_k ← x_k;
                    else
                        // x_k > y_k
                        iterate y further through index and add tax IDs
                          to tIDs_k if y_k matches entry in known_k;
                        stop if no full match occurs;
                        break out of k loop;
            if highest k was reached then
                get next y;
        // Part Three
        // see below
```

```
for Every entry with range in input do
    for All k-mers in k-mers_[lower,upper] do
        // Part One and two
        // <see above>
        // Part Three
        // look for any remaining y in case there are no
           more x left
        for any y left in range do
            for k from  lowest to highest do
                if y_k matches entry in known_k then
                    add tax ID to tIDs_k;
                else
                    break;
            if at least one k matched then
                get next y;
            else
                break;
        // all possible matches for old x_k found -> save
        for all entries in tIDs_k do
            if size of tIDs_k == 1 then
                save unique match;
            else
                save non-unique match;
            for all entries in rIDs_k do
                save match of read ID and tax ID;
return Scores and counts
```

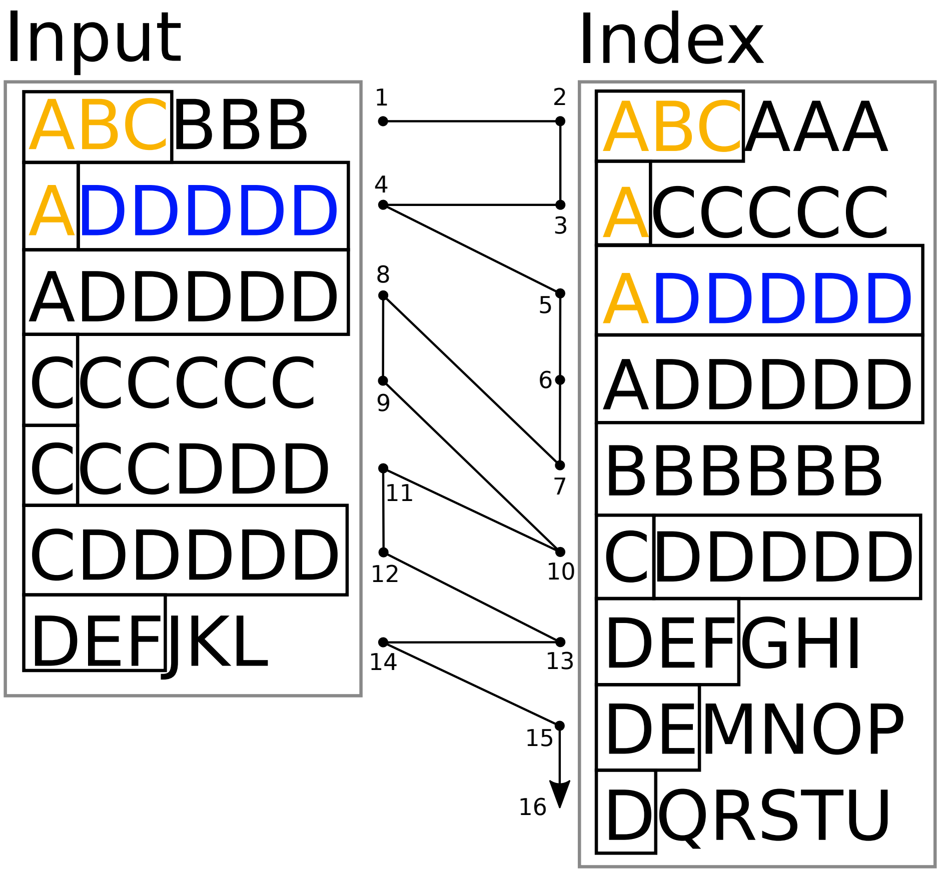


Figure 1: Schematic of the identification algorithm with a more complex example. The arrow in the middle together with the numbers show the order of execution. Rectangles around the letters mean matching letters. Color is used in the first $k$-mers to show how known $k$-mers are matched (step 3 matches with the "A" from step 2).

### 3 Versions and system specifications

Table 1: Versions of used tools.

| Tool | Version or date |
|---|---|
| Clark | 1.2.5 |
| Centrifuge | 1.0.4 2020-10-02 |
| Kraken2 | 2.0.8 2019-25-04 |
| Kraken | 1.1.1 |
| KrakenUniq | 0.5.8 |

**HPCC - iDiv EVE**

- DELL R640
- CPU: 2x 20-Core Intel(R) Xeon(R) Gold 6148 CPU @ 2.40GHz
- RAM: 1480GB DDR4 up to 2993MT/s
- Connected via InfiniBand(TM)

**Desktop**

- CPU: Intel(R) Core(TM) i7 7700K
- RAM: 16GB DDR4 - 2800 MHz
- SSD: Samsung 970 EVO(M.2)

**Laptop**

- CPU: Intel(R) Core(TM) i7-6500U
- RAM: 8GB DDR3 - 1600 MHz
- SSD: Samsung T5(USB 3.1G2)

### 4 Information loss of the amino-acid-like encoding

Let $\mathcal{A}$ be an alphabet of size $n$ with $1 \leq n \leq 28$, $n \in \mathbb{N}$ and $\mathcal{B} := \{A, C, G, T\}$. Let furthermore $S$ be a word consisting of at least three letters from $\mathcal{B}$, so $S \in \mathcal{B}^*$ and $|S| \geq 3$.

Let $\texttt{code} : B \times B \times B \to \mathcal{A}$ be a function with the following property:

$$\texttt{code}((b_0, b_1, b_2)) = code((c_0, c_1, c_2)) \Rightarrow b_1 = c_1,$$
$$(b_0, b_1, b_2),\ (c_0, c_1, c_2) \in \mathcal{B} \times \mathcal{B} \times \mathcal{B}. \qquad (\star)$$

`code` is usually neither surjective nor injective.

$\texttt{translate} : \mathcal{B}^* \to \mathcal{A}^* \times \mathcal{A}^* \times \mathcal{A}^*$ is now a function for translating the DNA sequence $S$ into three amino acid-like sequences with iterative application of `code`. If `code` is applied with a shifted start, we get the conversion in three frames described in the paper. So the resulting words $w_0$, $w_1$, $w_2 \in \mathcal{A}^*$ are as follows:

$$w_j := \bigoplus_{i=j}^{\lfloor \frac{|S|-j}{3} \rfloor} \texttt{code}(S\,[j + 3 \cdot i, j + 3 \cdot i + 3]),\ j = 0, 1, 2$$

where $\bigoplus$ is the string concatenation.

**Observation 4.0.1.** *$w_0$, $w_1$, $w_2$ are created from overlapping triplets, which means that for a DNA sequence $b_0$, $b_1$, $b_2$, $b_3$, $b_4$, ... the triplet of the first frame $b_0$, $b_1$, $b_2$ shares two letters $b_1$, $b_2$ with the second frame and one letter $b_2$ with the third frame. Furthermore, the second frame $b_1$, $b_2$, $b_3$ shares $b_2$, $b_3$ with the third frame which starts with $b_2$, $b_3$, $b_4$. $b_3$ and $b_4$ are now again the bases forming the first and second letter in the first frame.*

**Lemma 4.1.** *Apart from the first and last letter of S, no information loss occurs when using* `translate`.

*Proof.* To show that no information loss occurs, except for the first and last letter of $S$, we construct an appropriate inverse function $\texttt{translate}^{-1}$. We assume without loss of generality, that $\{w_0,\ w_1,\ w_2\} \in \text{im}(\texttt{translate})$.

$\texttt{code}^{-1}$ is created by determining each triplet $b \in B \times B \times B$ associated with the respective letter $a \in \mathcal{A}$ and storing it in a dictionary with $a$ as key. This means, that e.g. for $w_0 = a_0, ..., a_l$ with $0 \leq l < \lfloor \frac{|S|}{3} \rfloor$, $\texttt{code}^{-1}(a_0)$ is a set of ordered triples with the same middle component according to prerequisite $(\star)$.

Following observation 4.0.1, the sets created by applying $\texttt{code}^{-1}$ to the letters of $w_0$, $w_1$ and $w_2$ must contain at least one triplet, where the bases match in the described positions (see Figure 2).

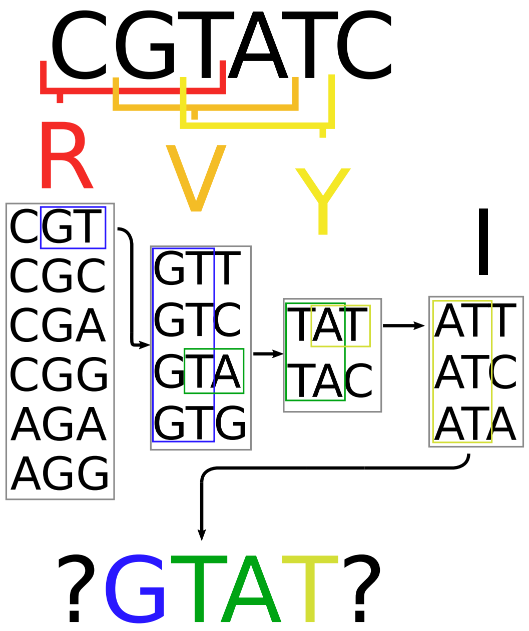


Figure 2: Translation and inversion.

Repeating this chained matching and checking reveals all interior bases of $S$. Since the first base in the first frame and the last base of the last frame are not necessarily unique and cannot be checked by the other frames, they are considered ambiguous. Therefore the sequence $S$ can be reconstructed with just the amino-acid-like encoded frames except for the first and last base. □

**Remark 4.1.1.** *The standard codon table cannot be constructed with* `code`*, because prerequisite ($\star$) is not satisfied ("S" has two middle bases). To fix that, one must first split the amino acid "S" into two letters ("AGT" and "AGC") and second give the stop codon "TGA" an additional letter as well. The default codon table used in kASA implements the additional stop codon but does not introduce a new letter to split "S" to be compatible with already converted amino acid sequences. Benchmark results (see Figure 2 in the paper) for both the standard codon table and one with a split "S" did not differ noticeably so an approximation of* `code` *(by the standard codon table) is sufficient.*

**Remark 4.1.2.** *One can reconstruct the first and last base of $S$ if we additionally restrict the alphabet $\mathcal{A}$ further: The first base of every triplet must be* *unique to the assigned letter $a \in \mathcal{A}$. Secondly, six instead of three frames are used (so the reverse complement is translated as well). A possible function* `code` *using a 16-letter alphabet maps each combination of the first two bases in a triplet to a unique letter, so*

$$\texttt{code}_{16}((b_0, b_1, b_2)) = code_{16}((c_0, c_1, c_2)) \Rightarrow b_0 = c_0 \text{ and } b_1 = c_1,$$

$$(b_0, b_1, b_2),\ (c_0, c_1, c_2) \in \mathcal{B} \times \mathcal{B} \times \mathcal{B}.$$

**Remark 4.1.3.** *During the proof we made the assumption that the ordering of the frames are as* `translate` *created them. However, should this ordering be disturbed we can try to use the constructed* $\texttt{translate}^{-1}$ *anyway because the reconstruction will fail if the ordering is not correct. This is because no other combination would generate $S$ in full length in the reconstruction process.*

An implementation of the constructive proof can be found in `https://github.com/SilvioWeging/kASA/tree/master/scripts/reconstructDNA.py`.
