## Supplementary figures and images for "kASA: Taxonomic Analysis of Metagenomic Data on a Notebook"

### heatmap_bar.png

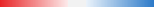

### plot_shannon.pdf

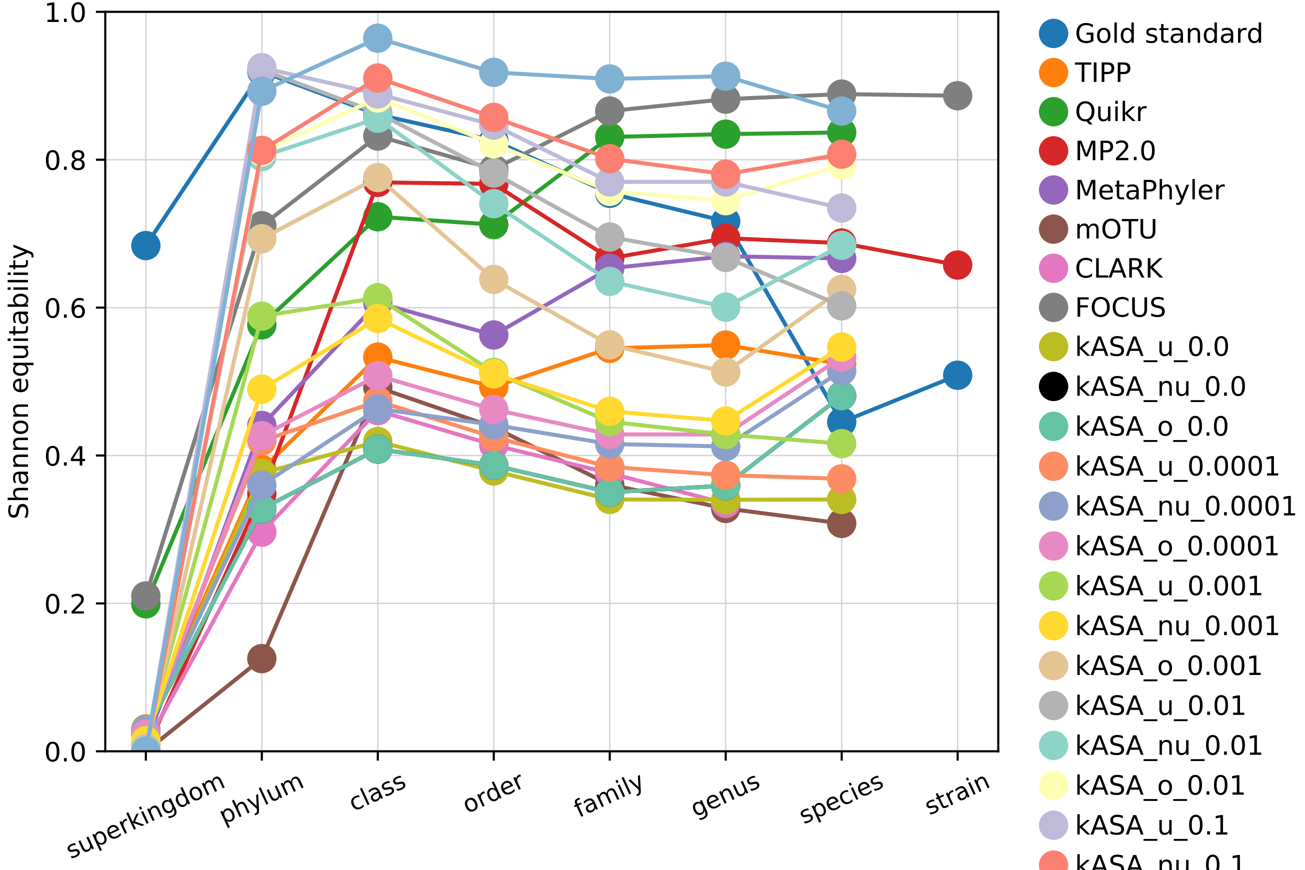

### plot_shannon.png

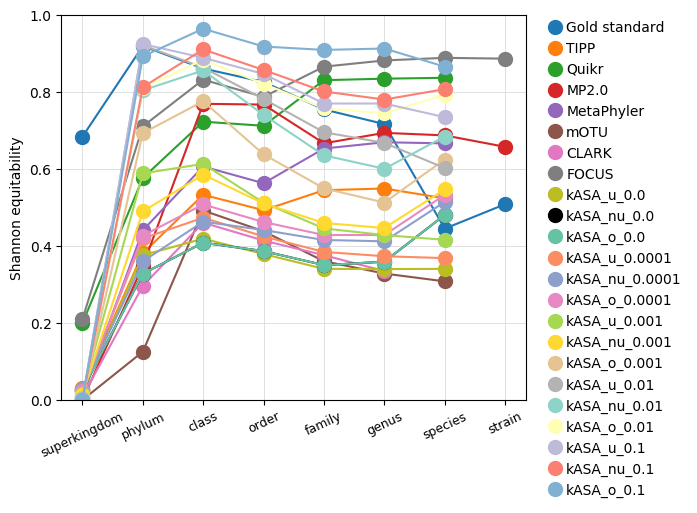

### plot_shannon_diff.pdf

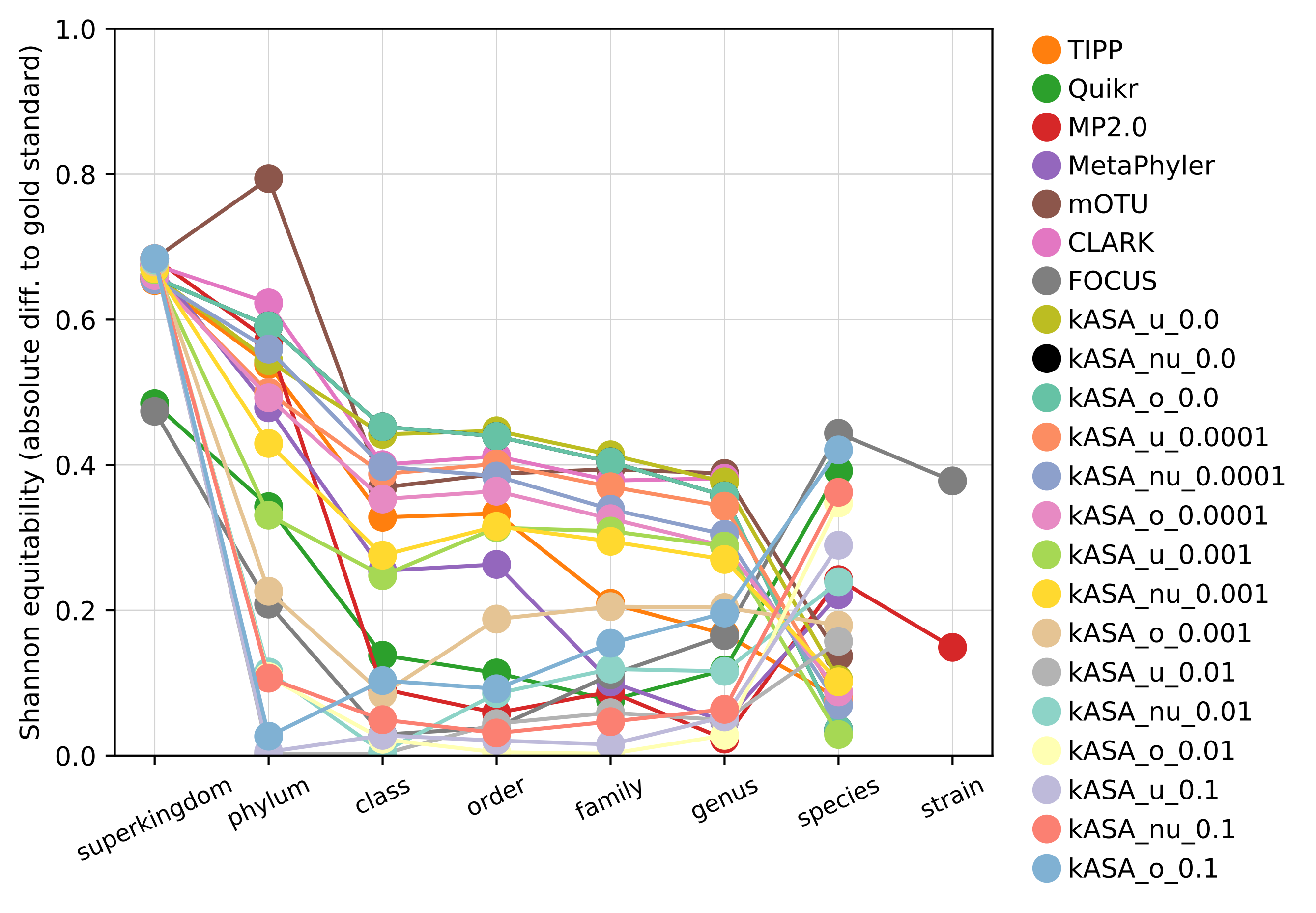

### plot_shannon_diff.png

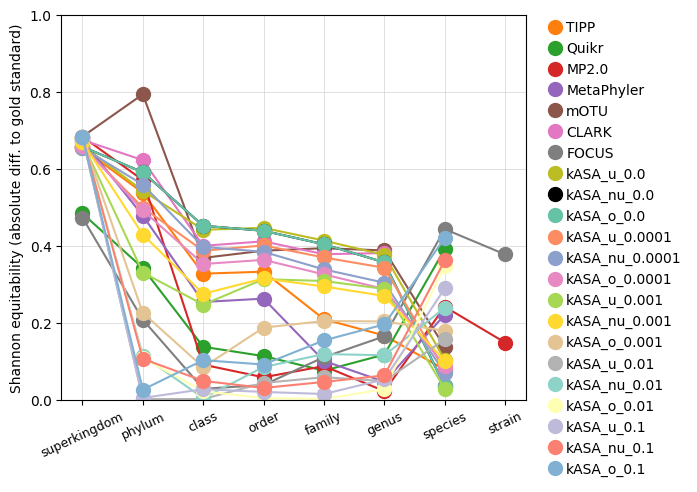

### rarefaction_curves.pdf

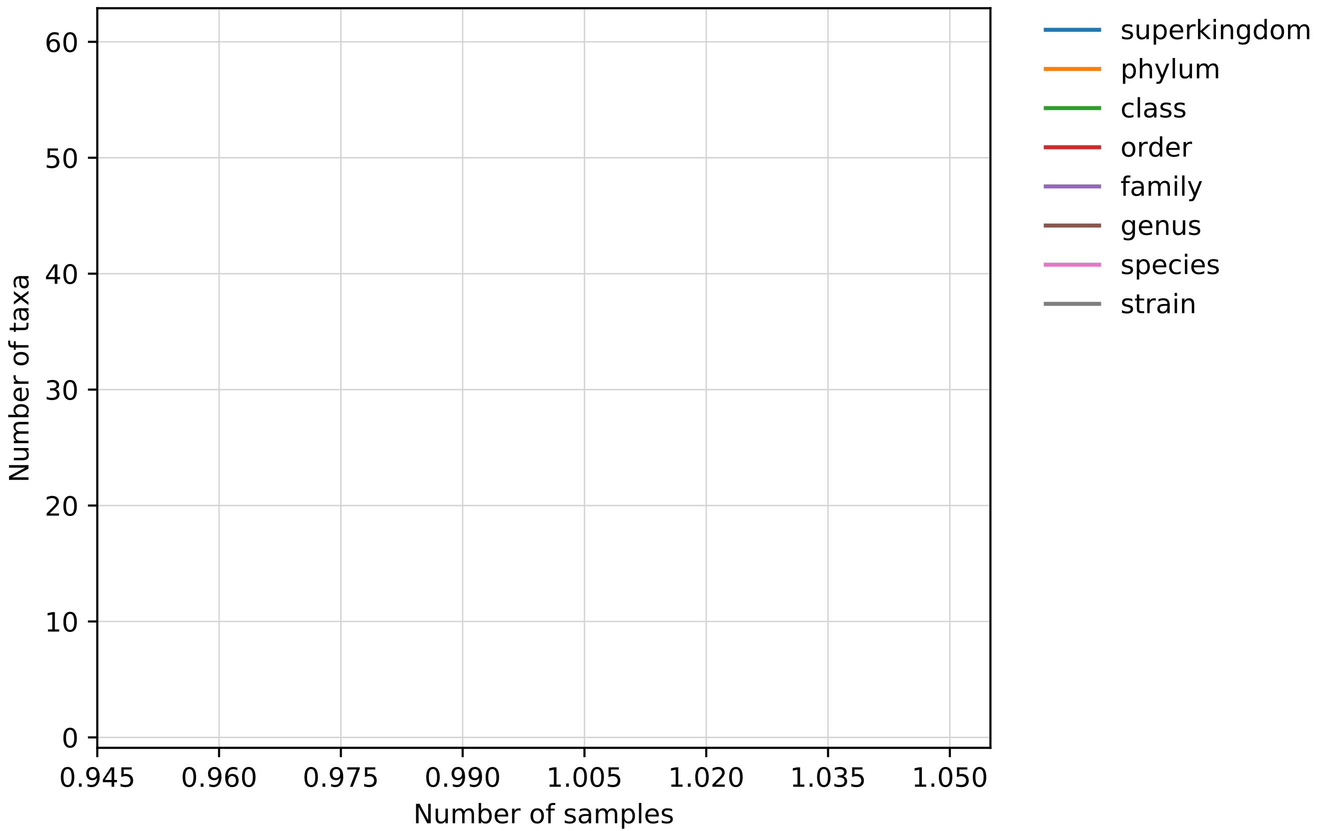

### rarefaction_curves.png

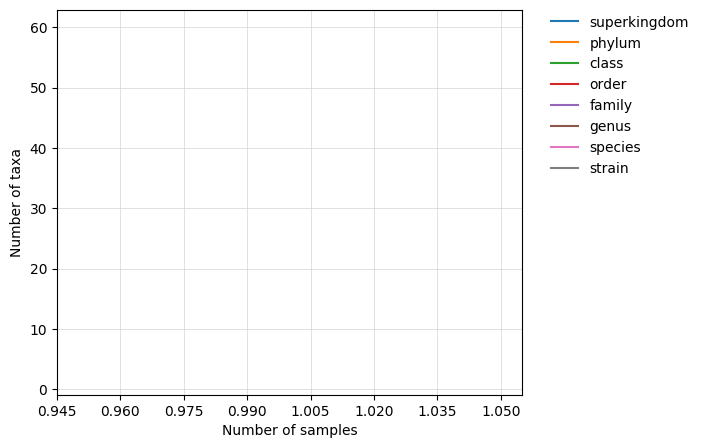

### rarefaction_curves_log_scale.pdf

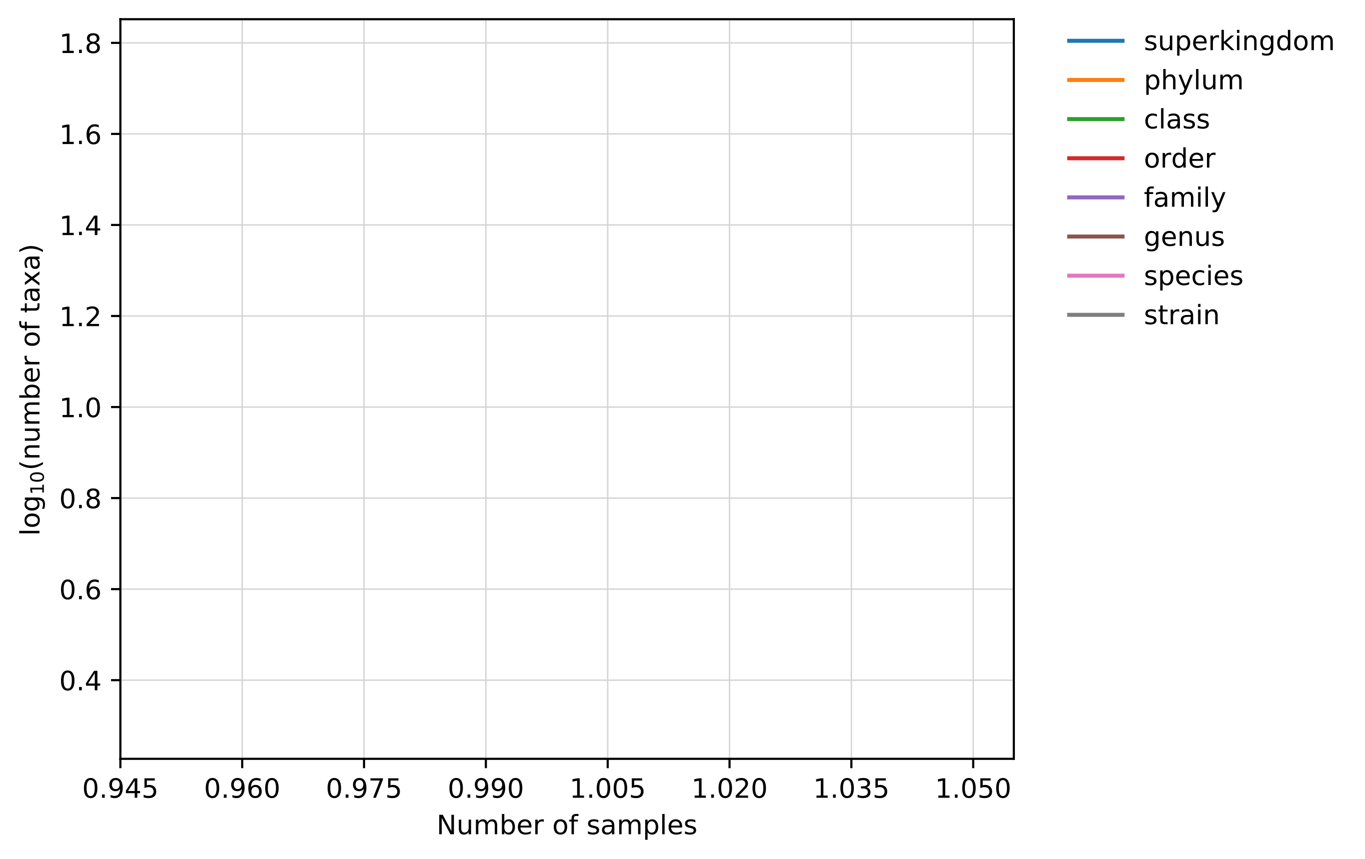

### rarefaction_curves_log_scale.png

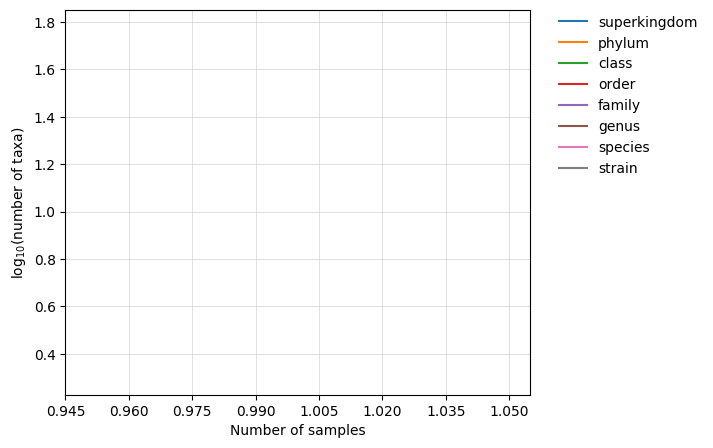

### Shrink_profile.pdf

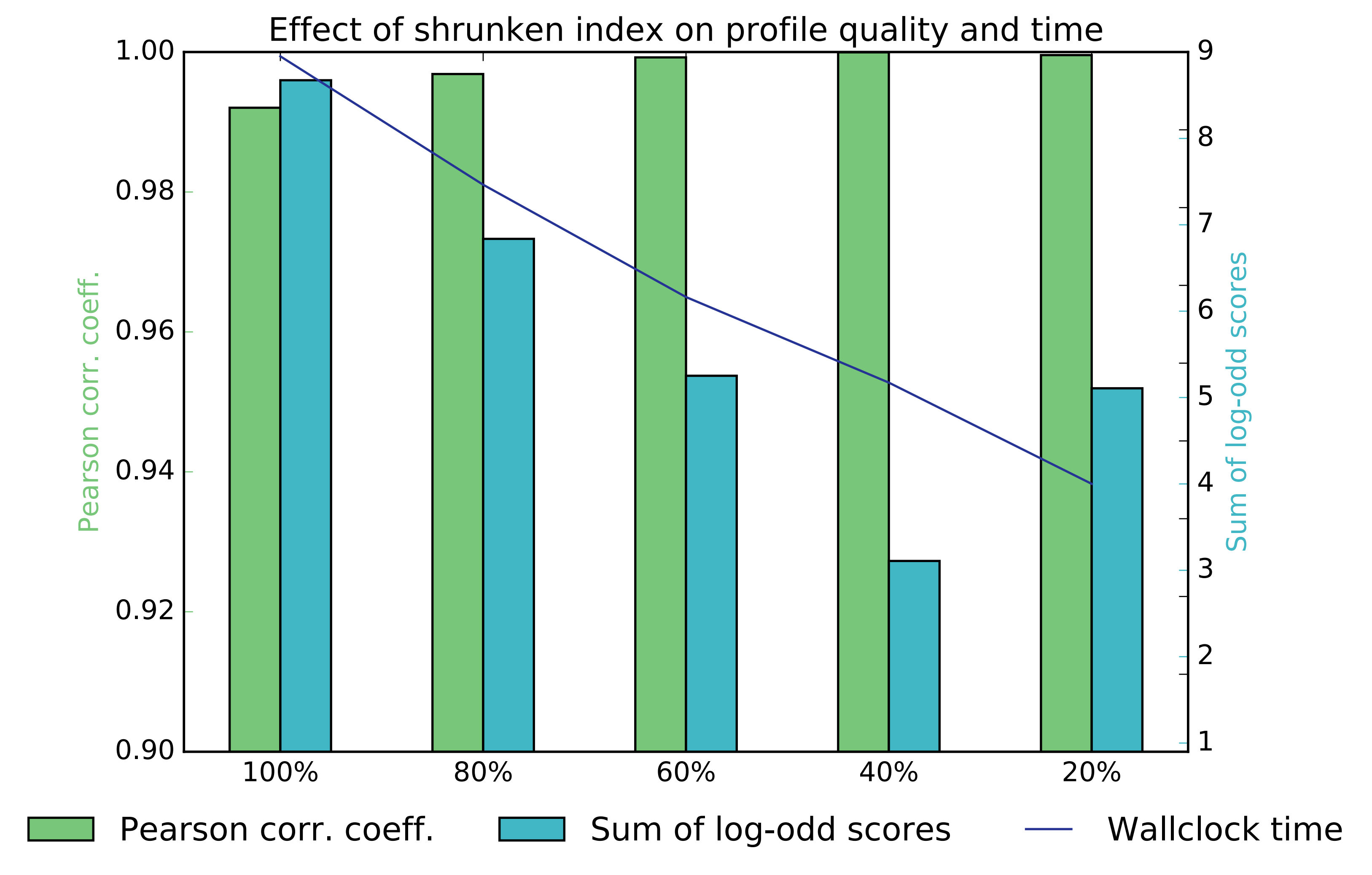

### spider_plot.pdf

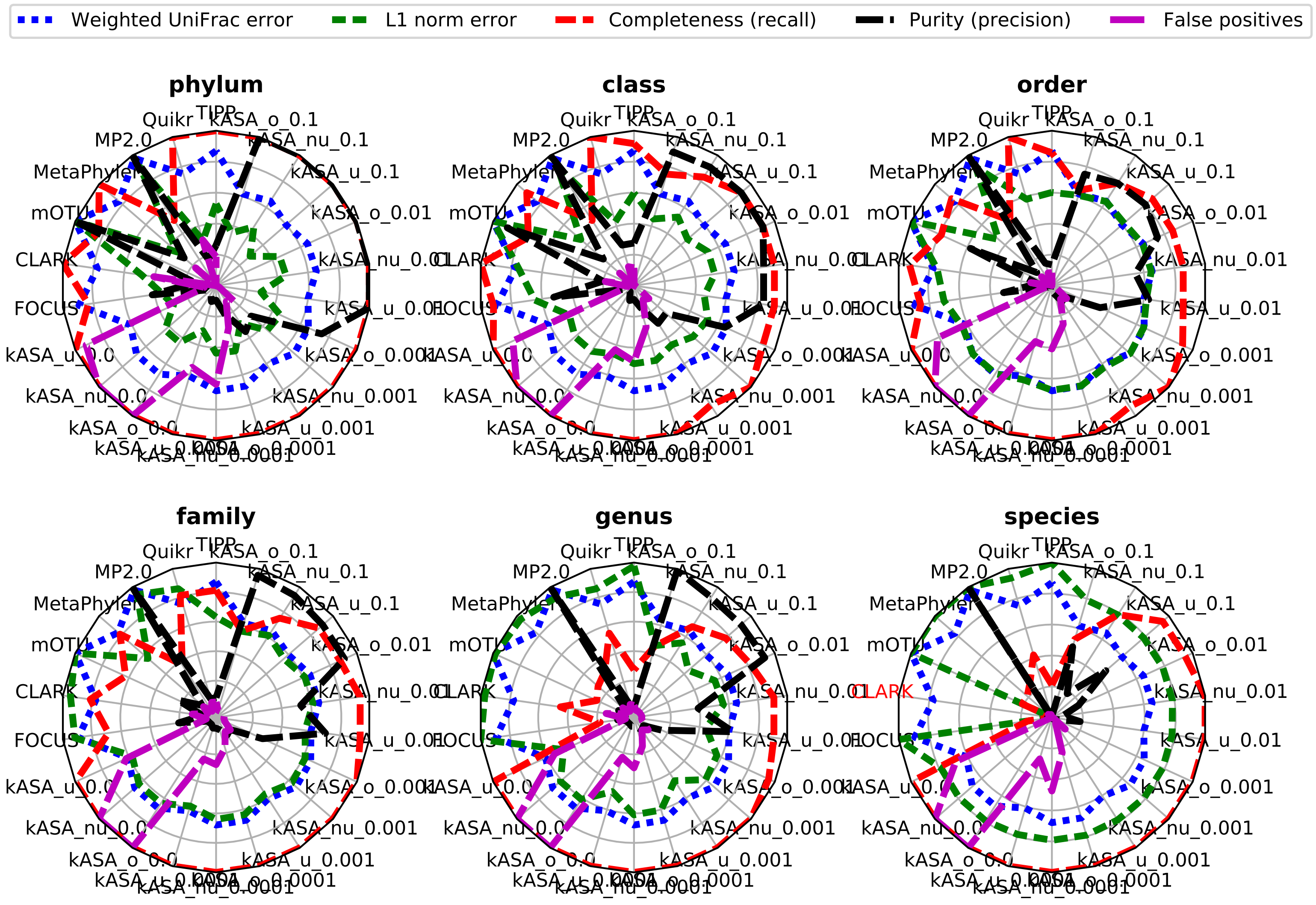

### spider_plot.png

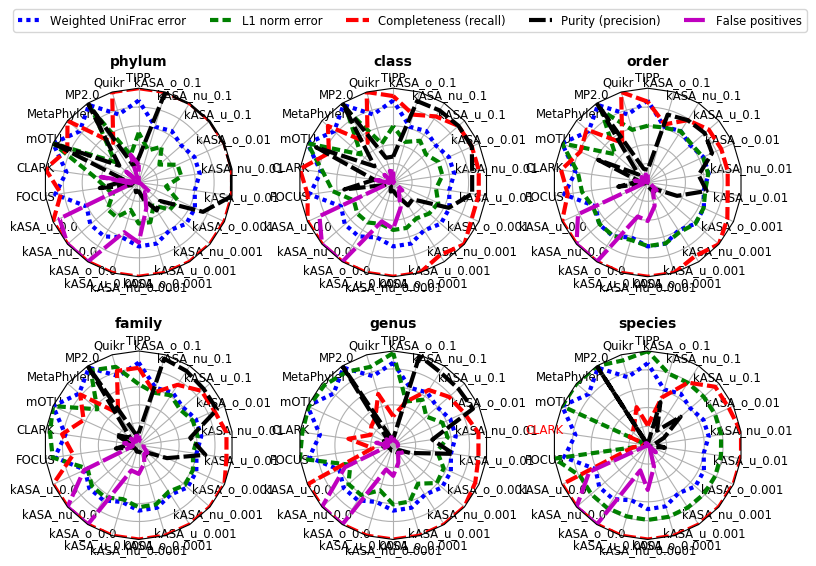

### spider_plot_recall_precision.pdf

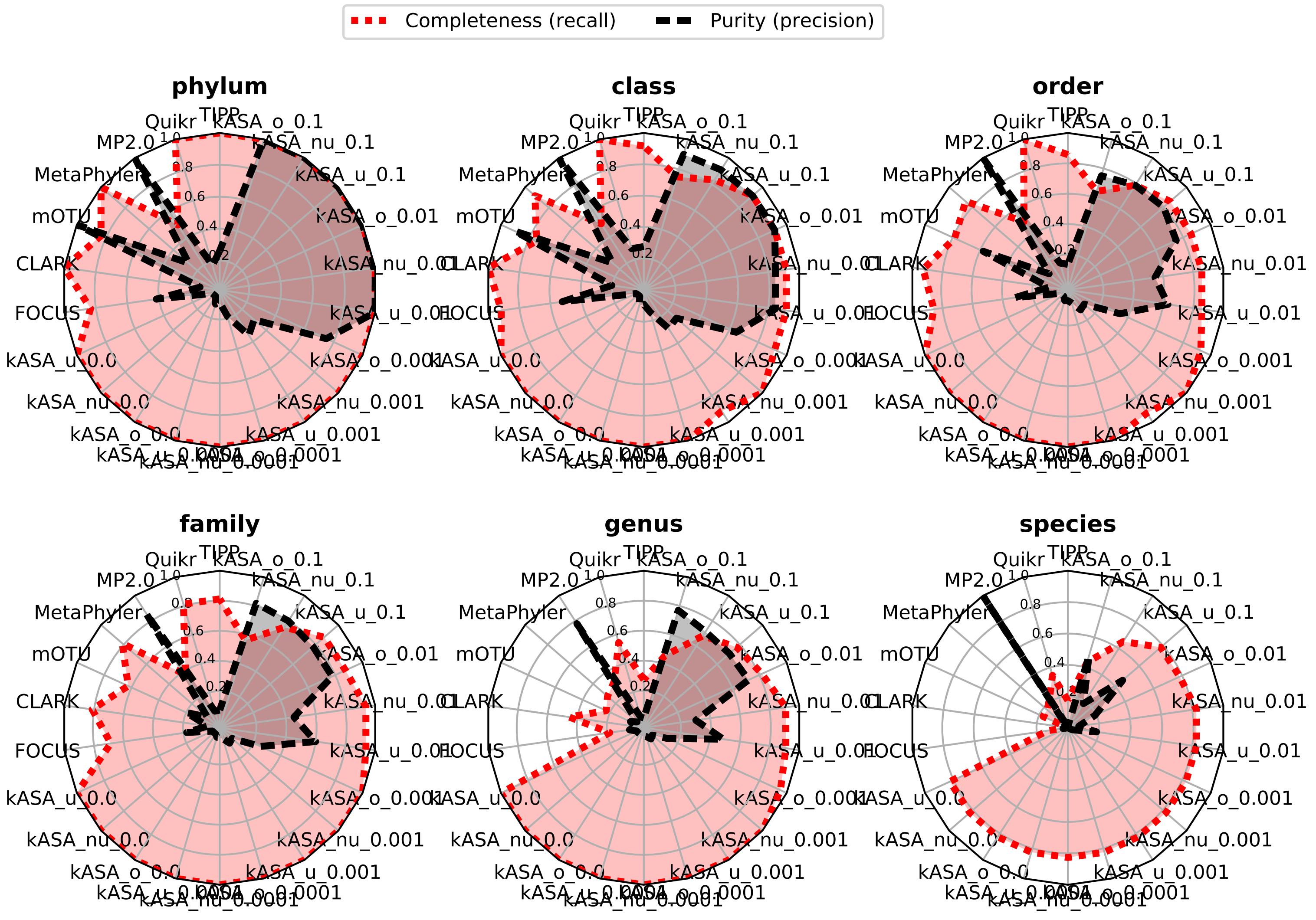

### spider_plot_recall_precision.png

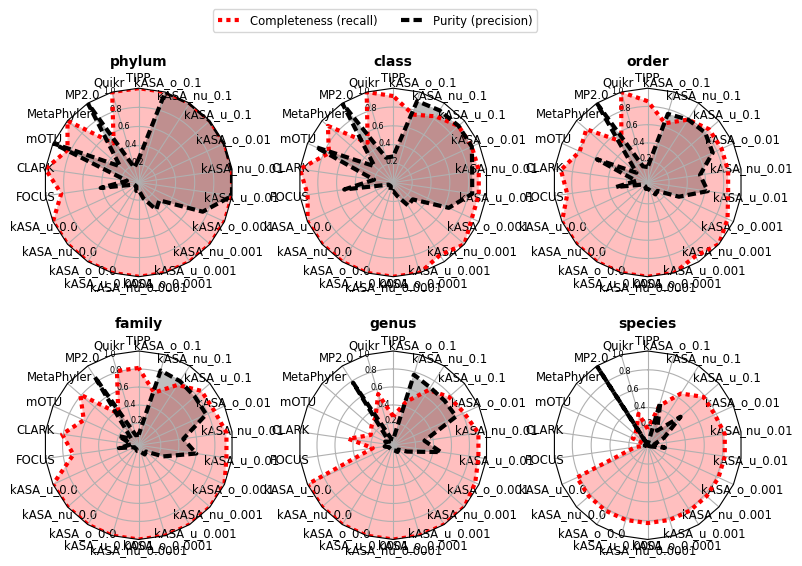
