## Additional file H for "kASA: Taxonomic Analysis of Metagenomic Data on a Notebook": Suppl_H.html

Javascript must be enabled to view this page.

magnitude
magnitudeUnassigned

real

666.874842412043

0.0949387472676
0.423936784054361

3.39594e-07

3.25227e-06
1.08409e-06

1.08409e-06
2.16818e-06

1.08409e-06

1.9667825e-06
5.9003475e-06

1.467248e-06
2.934496e-06

3.29836e-07

4.28693e-07

5.93505e-07

1.15214e-07

6.5149e-08
3.25745e-08

3.25745e-08

9.3392e-07
4.6696e-07

1.62808e-07

3.04152e-07

3.74473e-07

1.70046e-08

4.15325e-08

2.69961e-07

6.09143e-08

1.24597e-07

5.8187186e-07
1.74561558e-06

1.16374372e-06

1.277261e-07
2.554522e-07

8.94546e-08

1.86896e-08

1.95819e-08

3.93722e-07
7.87444e-07

1.96861e-07

1.96861e-07

1.2084752e-07
6.042376e-08

2.27889e-08

3.60676e-08

1.56726e-09

8.42144e-07

3.31925e-09

9.37413e-07

3.31433e-08

2.9790075e-06
8.9370225e-06

3.3226e-08
1.6613e-08

1.6613e-08

2.9623945e-06
5.924789e-06

4.57675e-07

2.76488e-07

4.15325e-08

8.3065e-08

6.35024e-07

5.51959e-07

8.33586e-07

8.3065e-08

1.29212e-07

1.12663e-08

9.775639e-07
3.895257e-07

1.154475e-07
2.30895e-07

2.03445e-08

7.849e-08

1.6613e-08

8.3065e-08
1.6613e-07

8.3065e-08

3.73792e-08

1.53634e-07

2.76883e-08

1.6613e-07

2.76883e-08

1.25856e-07

3.93563e-09

1.2881394432e-05
4.511942544e-06

8.3065e-08

5.57524e-07

3.236224e-06
6.472448e-06

3.317e-07

7.72405e-07

4.86166e-07

4.10657e-07

8.02961e-07

4.32335e-07

1.38442e-08

6.21285344e-07
1.242570688e-06

4.84344e-10

6.20801e-07

1.245975e-07
4.15325e-08

4.15325e-08
8.3065e-08

4.15325e-08

6.6452e-08

1.6613e-08
4.9839e-08

3.3226e-08
1.6613e-08

1.6613e-08

1.24597e-07

1.95201e-07

1.24597e-07

1.18664e-08

1.25389e-07

1.69741e-07

5.53766e-10
1.661298e-09

1.107532e-09
5.53766e-10

5.53766e-10

0.000181704

2.21507e-08
6.64521e-08

4.43014e-08
2.21507e-08

2.21507e-08

8.75123e-09

2.49195e-07
8.3065e-08

1.6613e-07
8.3065e-08

8.3065e-08

3.49565e-07

1.74322e-06

0.00127716

4.15325e-08

7.99376e-07

1.12996e-05

2.05435e-07

9.49357e-08

1.8766378e-05
4.7027286e-05

3.59311e-06

1.17468e-07

2.46721e-06

3.38769e-06
6.77538e-06

3.38769e-06

3.09406e-06

1.221368e-05
6.10684e-06

6.10684e-06

9.9678e-08

2.53545e-07

0.000183775

7.50754e-08

2.12673e-06

9.69091e-08

2.76883e-08

2.49195e-07

2.6544453e-06
7.9633359e-06

2.26014e-06
1.13007e-06

1.13007e-06

2.76883e-08
5.53766e-08

2.76883e-08

9.85641e-07
1.971282e-06

5.586e-07

4.27041e-07

5.11046e-07
1.022092e-06

5.11046e-07

1.6613e-07

2.311632e-07
6.934896e-07

4.15324e-08
2.07662e-08

2.07662e-08

4.20794e-07
2.10397e-07

2.10397e-07

1.10753e-07

1.9942e-07

1.24597e-07

9.90409e-06

1.16291e-07

4.15325e-08

3.3226e-07

2.07987e-09

4.15325e-08

1.85293e-07

0.0741855077223
0.262439225077298

1.67186e-08

3.22129e-07

4.15325e-08

0.000867163

1.77312e-07

3.52397e-08

5.13673e-07

0.00780101280867
0.022515259547434

6.36808e-06
3.18404e-06

1.55049e-06

1.63355e-06

1.0441e-07

5.53351e-07

3.82964e-07

1.19309e-05

2.90197e-07

1.20576e-07

9.45716e-07

1.6613e-08

4.0779253e-06
8.1558506e-06

1.8797e-07

3.68171e-07

4.42844e-07

4.42844e-07

3.93531e-07

3.18859e-07

1.62801e-08

4.42844e-07

7.26608e-08

2.41893e-08

4.69639e-07

3.93531e-07

3.92345e-08

7.26608e-08

3.80344e-07

1.23228e-08

7.4267e-08

1.41016e-06

2.94602e-07

2.20545e-07

2.15751e-06

7.4267e-08

2.171124e-06
1.085562e-06

3.19147e-07

3.4143e-07

4.24985e-07

5.85274e-07

3.56373e-06

9.98657e-07

1.61376e-08

8.68721e-08

1.6373e-07

1.1152e-07

2.13462e-06

1.66534e-08

2.84822e-06

9.03312e-08

6.995938e-06
1.3991876e-05

3.76745e-06

3.42556e-07

3.42556e-07

2.20082e-06

3.42556e-07

6.36831e-08

3.29296e-07

1.07984e-06

2.7705e-07

1.27017e-06

2.20705e-07

0.000788761564796

0.00037327457621
0.00074654915242

4.82221e-09

5.51359e-06

0.000205064

4.4164e-08

0.000162648

2.1106206188e-05
4.2212412376e-05

9.3226e-08

3.00456e-08

3.5788e-08

1.23111e-07

5.16613e-08

7.47585e-08

1.55553e-08

2.01858e-08

8.78634e-06

3.40843e-08

7.71174e-07

1.23408e-08

8.73354e-06

2.96213e-08

1.16296e-06

1.41363e-07

2.37754e-08

2.93096e-08

1.76603e-07

9.37086e-08

3.15598e-07

1.47911e-08

6.19888e-10

1.46352e-08

9.88125e-08

2.22598e-07

1.94502e-06

1.59347e-06

1.17777e-06
2.35554e-06

1.17777e-06

1.73141e-06

0.00065033

1.98101e-06

5.8241e-07
2.91205e-07

2.91205e-07

7.84473e-06

5.59333e-07

3.05666e-05

2.10889e-06

1.60902e-07

5.28277e-07

3.15382e-07

1.403932e-06
7.01966e-07

1.73032e-07

2.53237e-07

2.75697e-07

7.4267e-08

1.7011e-07

3.6992e-07
7.3984e-07

1.95692e-07

1.74228e-07

2.77091e-06

5.53766e-07
1.107532e-06

2.33336e-07

3.2043e-07

4.58201e-08

7.44854e-08

1.39736218e-05

3.646382e-06
7.292764e-06

1.69616e-07

8.24472e-07

4.91414e-07

9.7763e-07

1.18325e-06

1.4368818e-06
7.184409e-07

1.83533e-07

1.42e-07

1.42e-07

1.83533e-07

6.73749e-08

5.243976e-06
2.621988e-06

1.61195e-06

3.53315e-07

5.41939e-07

1.14784e-07

2.05375e-07

1.39987e-06

1.984389e-06
3.968778e-06

1.41504e-07

8.17612e-07

4.8083e-07

2.28757e-07

1.14617e-07

2.01069e-07

9.97598e-08

7.43587e-06

3.32854e-07
1.66427e-07

1.66427e-07

9.71776e-06
1.943552e-05

3.41873e-06

3.00816e-06

3.29087e-06

3.66938e-07

4.39937e-07

2.97534e-07

3.29308e-07

2.92565e-06

1.4944e-07

5.67268e-07

2.6161e-06

1.14342e-07

7.47054e-07

4.84995e-07

1.103282e-05
5.51641e-06

3.21573e-06

2.01583e-06

2.8485e-07

6.28411e-08

7.14621e-07

2.80879e-07

4.16549e-07

9.7561e-07

1.15415e-06

1.67808e-08

7.0314424e-05

1.66734e-07

4.588036e-06
9.176072e-06

8.23292e-07

2.89992e-06

8.64824e-07

2.66389e-06

2.5698e-05
1.2849e-05

4.01107e-06

4.41638e-06

4.42155e-06

6.10299e-07

5.269946e-06
2.634973e-06

2.25722e-06

3.77753e-07

3.77795e-07

7.62451e-07
1.524902e-06

7.62451e-07

3.951193e-06
7.902386e-06

4.24833e-07

3.52636e-06

8.4622e-06
1.69244e-05

1.81971e-07

4.12292e-07

1.98859e-06

4.02744e-07

3.27399e-07

2.73769e-07

2.01813e-06

5.76482e-07

8.25789e-07

1.13272e-06

1.61157e-07

1.61157e-07

3.90894e-08

1.71001e-07

7.4267e-08

0.0125066484262

9.09966e-07
4.54983e-07

2.70112e-07

7.7566e-08

1.07305e-07

9.650916e-05
0.00019301832

2.3648e-06

1.61944e-06

1.82856e-05

3.47723e-06

7.35333e-06

5.53503e-05

4.78361e-06

3.27485e-06

1.649348e-05
3.298696e-05

1.4507e-05

1.98648e-06

2.12337e-05
1.061685e-05

8.45453e-06

2.16232e-06

1.30989e-06

1.39807e-07

0.00015190768
7.595384e-05

2.70161e-05

2.04658e-06

2.19546e-05

2.34582e-06

2.38704e-06

2.02037e-05

0.0059979596126
0.0119959192252

0.000224521

0.000198011

9.50087e-07

0.000487305

7.37202e-08

0.00017238

0.000150917

0.000142395

6.17491e-08

0.000186337

8.21899e-07

4.89396e-06

6.90099e-08

8.81837e-05

2.69758e-07

1.49571e-07

2.40196e-06

1.55442e-06

0.000114501

5.06983e-07

0.000339824

1.23004e-07

1.79717e-06

2.96043e-06

2.11743e-07

0.000182885

0.00012183

1.43809e-07

1.84176e-07

0.000111938

0.000149003

0.000892222

0.00033232

0.000386247

0.000162978

2.05795e-07

0.000402778

0.000156914

1.88867e-06

0.000272009

1.28197e-05

9.42223e-05

4.35772e-06

2.44403e-06

7.71855e-07

1.7948e-06

3.71867e-07

1.65196e-06

2.48408e-05

0.000133818

1.94052e-06

2.91545e-07

0.000156266

2.28797e-06

0.000152732

6.81664e-08

1.99082e-07

5.11715e-07

8.04968e-07

0.000110998

4.49636e-06
8.99272e-06

1.01385e-06

1.70096e-06

1.78155e-06

4.96705e-07

7.32668e-06
3.66334e-06

2.17275e-06

1.49059e-06

2.991289e-05
5.982578e-05

2.35482e-06

4.1525e-06

1.04074e-05

2.28357e-06

1.07146e-05

2.0067986e-05
1.0033993e-05

6.71513e-06

1.46521e-06

1.02273e-06

8.30923e-07

1.57424e-06

8.32247e-07

1.010652e-05
5.05326e-06

2.48716e-06

2.5661e-06

3.29087e-06

1.16083e-07

8.3065e-08

4.82221e-09

2.61819e-07

1.85937e-06

1.01645e-06

1.08526e-06

2.80658e-07

4.82221e-09

1.05813e-06

3.26327e-08

1.831590798e-06
3.663181596e-06

3.98764e-08

1.08228e-06

1.53398e-10

1.45364e-07

1.8524e-07

3.78677e-07

4.66955e-08

8.50549e-08
1.701098e-07

6.84419e-08

1.6613e-08

1.47753e-06
2.95506e-06

1.47753e-06

1.92991e-06

9.77234e-08

1.48523e-07

4.193e-07

2.34643e-06

7.73336e-07
1.546672e-06

3.46104e-08

3.46104e-08

3.46104e-08

3.46104e-08

3.46104e-08

3.00418e-07

3.46104e-08

3.46104e-08

3.46104e-08

3.46104e-08

3.46104e-08

1.26814e-07

5.1204e-07

4.84032e-07
9.68064e-07

4.84032e-07

5.85911e-09

9.27706e-07
4.63853e-07

4.63853e-07

6.90935e-07

7.15142e-08

6.24086e-08

3.24896e-07
6.49792e-07

2.64397e-07

6.0499e-08

3.29816e-07

5.08268e-09

1.74987e-08

5.27952e-06
2.63976e-06

1.34216e-06

1.2976e-06

3.35441e-06

4.91375e-06

1.775056e-07
3.550112e-07

9.49716e-08

8.2534e-08

2.9522e-06

1.67652e-06

5.06651e-07

1.815222e-06
9.07611e-07

2.68241e-07

3.82117e-07

2.57253e-07

5.78347e-08

7.8216e-08

3.37179e-06

6.5945904e-07
3.2972952e-07

3.61152e-09

3.26118e-07

1.77066e-07

5.81239e-07

1.460730512e-06

1.456435e-07
2.91287e-07

3.12895e-08

1.14354e-07

1.1616096e-06
5.808048e-07

4.72635e-08

4.72635e-08

8.8796e-08

8.8796e-08

8.8796e-08

8.51948e-08

4.72635e-08

4.72635e-08

4.0168e-08

2.22422e-09
4.44844e-09

2.22422e-09

1.692736e-09
3.385472e-09

8.46368e-10

8.46368e-10

4.61448e-08

1.05151e-06

2.35165e-08

4.07194e-07

4.27381e-07

3.293488e-06
6.586976e-06

9.38568e-07

2.35492e-06

4.3304e-07
8.6608e-07

4.3304e-07

1.01172e-06

1.27953e-08

8.68526e-08

1.24136e-07

3.07982e-06

2.8096e-06
1.4048e-06

1.4048e-06

8.3065e-08

9.67858e-08

2.0432e-06

6.4101403e-07
1.28202806e-06

5.30693e-09

1.18664e-08

4.98806e-07

8.3065e-08

2.37328e-08

1.82369e-08

1.063e-07

1.30169e-07

2.37328e-08
4.74656e-08

1.18664e-08

1.18664e-08

2.39263e-07

2.82421e-06

3.40566e-07

1.75516e-06

4.69245e-07

5.62213e-06

1.38497e-07

6.47564e-07

1.66965e-09

8.26498e-07
4.13249e-07

4.13249e-07

6.23906e-07

8.26694e-07

1.71394e-05

2.78756e-07

1.61288e-07

1.32295e-07

3.25638e-07

5.09213e-07

2.45394e-06

4.16549e-07

2.85846e-07
5.71692e-07

2.85846e-07

7.05572e-07

5.47085e-07

2.85547e-07

1.31945e-05

1.933548e-06
3.867096e-06

9.66774e-07

9.66774e-07

2.63039e-08

9.61906e-07

1.509967e-06
3.019934e-06

2.56117e-07

1.25385e-06

4.74303e-07

2.28216e-07

1.477374e-06
7.38687e-07

4.80097e-07

2.5859e-07

9.79008e-07

8.66148e-08

9.44924e-07

8.69685e-08

4.53559e-06

4.9424e-06

1.6613e-07

4.35964e-06

1.65914e-07

1.80673e-06

8.2534e-08

3.08848e-07

9.883575e-06
1.976715e-05

7.22395e-06

7.51542e-07

4.8935e-07

9.14972e-07

5.03761e-07

1.23111e-07

1.02993e-06

9.87577e-08

8.52513e-07

2.833422e-06
1.416711e-06

1.37614e-07

3.08075e-07

3.06378e-07

1.14257e-07

3.06378e-07

1.20913e-07

1.23096e-07

4.79389e-07

4.90467e-06

0.00031888943

1.77023e-06
8.85115e-07

2.33145e-07

1.02615e-07

2.33145e-07

3.1621e-07

0.0001585596
0.0003171192

6.38852e-05

3.21276e-05

6.25468e-05

2.15035e-06

3.90678e-07

2.86757e-07

4.43013e-08

1.19937e-07

8.58611e-08

0.129356570481599
0.0484795586562

3.19143e-08

7.38957e-07

0.000943253

2.00982e-07

1.52738e-08

1.19088e-08

1.03142e-05

7.16378e-07
3.58189e-07

1.24597e-07

2.33592e-07

3.61777e-07

2.77489e-06

1.02706e-06

7.97501e-06

1.4375e-07

0.0001031942
0.0002063884

2.01113e-05

1.60997e-05

3.40784e-05

1.75401e-05

1.53647e-05

2.10771e-07

6.55723e-07

5.63995e-07

2.11793e-06

2.18019e-07
4.36038e-07

2.18019e-07

1.4117e-06

1.42071e-05

3.20735e-06

8.89486034e-06
1.778972068e-05

6.36012e-07

6.68026e-07

5.90179e-08

7.54255e-07

4.00857e-06

6.46932e-07

4.37184e-09

3.62322e-07

2.54792e-07

3.71365e-07

4.36035e-07

9.88595e-08

3.07648e-08

3.92449e-07

4.99893e-08

1.21099e-07

1.16874e-07

2.61317e-07

4.43157e-06

1.75984e-06

4.89734e-07
2.44867e-07

1.11248e-07

1.33619e-07

1.63664e-06

1.71668e-07

1.33155e-06

7.72054e-05

0.00214297

8.0628e-07
4.0314e-07

1.57914e-07

2.45226e-07

1.27016e-06

1.72382e-06

1.4053712e-05

6.11778e-06
1.223556e-05

6.11778e-06

9.09076e-07
1.818152e-06

9.09076e-07

8.3065e-08

2.37752e-07

4.57275e-05

1.0224e-06

1.06798e-07

1.45084e-06

9.42853e-07

1.6613e-07

6.57966e-06
1.315932e-05

1.64772e-06

1.60956e-06

1.66999e-06

1.65239e-06

3.19757e-07

2.96265e-07

3.39505e-08

2.98533e-06

3.61015e-07

3.3583e-05

1.17251e-06
2.34502e-06

5.85649e-07

5.86861e-07

8.00473e-08

1.32904e-06
2.65808e-06

6.6452e-07

6.6452e-07

2.33221e-08

1.18805e-06

1.10835e-07

6.209066e-06

1.5143e-06
7.5715e-07

7.5715e-07

4.694766e-06
2.347383e-06

7.24693e-07

1.62269e-06

3.02932e-06
1.51466e-06

4.12535e-07

4.12535e-07

6.8959e-07

3.11148e-07

1.45018e-06

5.13272e-06

2.1376e-07

5.69291e-07

3.05588e-07
1.52794e-07

8.23302e-08

7.04638e-08

1.87588e-06

4.03458e-08

7.16425e-07

3.35299e-08

1.5299e-07

7.68454e-07

0.000247654

1.18069e-07

7.21867e-06

4.99651e-07

2.20779e-06

1.634247e-07
3.268494e-07

3.20157e-08

3.20157e-08

3.20157e-08

3.89734e-08

2.84042e-08

1.7872e-07

7.13822e-06

1.26003e-05

1.17147e-06

3.3364e-05

3.04692e-07

2.21506e-07
4.43012e-07

1.10753e-07

1.10753e-07

3.2243e-05

9.30328e-07

1.06087e-06

6.59599096e-05
3.29799548e-05

4.67186e-07

1.58673e-06

1.01502e-06

1.93616e-07

1.29041e-07

4.63277e-08

8.29142e-08

1.02517e-07

1.91904e-06

8.49806e-07

1.24752e-07

1.88824e-06

1.791e-06

7.17901e-07

4.70005e-07

1.04578e-06

1.45097e-06

2.11537e-07

1.17603e-07

1.06501e-06

4.51766e-07

2.50274e-06

1.72943e-07

4.79033e-07

1.84589e-08

4.64839e-07

1.05003e-06

2.23986e-07

1.2675e-08

5.58402e-07

2.28641e-07

1.1714e-07

4.70005e-07

5.04683e-07

1.15691e-07

2.45651e-06

8.49806e-07

1.00592e-06

1.878e-06

5.32306e-07

4.0321e-07

5.3673e-07

3.70586e-07

1.09849e-06

9.34867e-07

2.67501e-07

3.72316e-07

9.9044504e-05
0.000198089008

3.46032e-06

2.07671e-06

1.76091e-06

7.78782e-07

1.59549e-06

1.10872e-06

2.53746e-06

1.88224e-06

2.81773e-06

1.61074e-06

7.73537e-07

1.54418e-06

2.02022e-06

1.38424e-06

1.42938e-06

9.40706e-07

1.82625e-06

1.54876e-06

1.04014e-06

8.31754e-07

1.71684e-06

1.05785e-06

1.10108e-06

1.12638e-06

3.90003e-07

8.40685e-07

1.48242e-06

1.06398e-06

1.5091e-06

1.18581e-06

1.29758e-06

1.19912e-06

2.2695e-06

2.81055e-06

8.07057e-07

9.4507e-07

6.35637e-07

1.23036e-06

3.01723e-06

2.40423e-06

2.6533e-06

1.8741e-06

9.82837e-07

9.6775e-07

1.20756e-06

1.35438e-06

1.93727e-06

6.05482e-07

8.04858e-07

1.32112e-06

2.37021e-06

5.31353e-07

2.89672e-06

1.87659e-06

7.86452e-07

7.3985e-07

6.73791e-07

1.81783e-06

1.25843e-05

0.0002010417843
0.00010052089215

2.42567e-06

1.29244e-06

8.60878e-07

1.93623e-06

1.8256e-05

1.51357e-06

5.32277e-09

7.03467e-07

2.46648e-07

7.15397e-07

2.44271e-06

9.11265e-07

9.6269e-07

1.20735e-07

4.46051e-07

7.09938e-09

3.20445e-07

1.18617e-05

2.28092e-05

2.28526e-06

1.2871e-07

1.89267e-06

2.49036e-06

1.22504e-06

2.7262e-06

7.72034e-07

2.11631e-05

3.36766e-08

6.85236e-07

1.89369e-07

8.3065e-08

1.17021e-05

2.76883e-08

6.96644e-06

1.6613e-07

1.60708e-06

7.95843e-07

8.3065e-08

0.00017855108
0.00035710216

3.8791e-05

6.61847e-05

6.37088e-06

2.55862e-05

2.88067e-05

1.28116e-05

8.18873e-07

4.3205e-07

2.43273e-07

1.52738e-08

1.04098e-07

1.36218e-06

2.30736e-07

7.2128e-08

5.932e-07

4.42014e-07

8.97347e-08

1.60546e-06

3.00878e-06

3.28799e-08

1.70119e-07
8.50595e-08

8.50595e-08

5.2551325267e-05
0.000105102650534

1.10097e-08

3.11539e-07

5.15746e-08

1.65139e-10

1.78307e-07

7.56723e-08

2.86487e-05

3.40833e-08

4.02083e-08

1.95805e-07

1.10097e-08

1.06343e-07

4.91141e-08

8.41958e-08

7.88366e-08

1.10097e-08

1.50034e-07

6.52468e-08

3.31425e-07

4.56005e-07

1.64158e-07

7.35062e-08

4.56005e-07

1.53824e-09

1.18252e-07

2.01073e-08

5.06251e-08

3.43381e-07

8.2409e-09

5.75626e-08

4.4684e-08

1.23388e-06

1.2043e-07

5.06251e-08

1.32395e-07

3.76223e-07

1.2548e-08

4.27854e-08

1.52199e-06

8.2409e-09

9.39458e-09

5.29423e-08

1.41741e-08

2.48539e-08

3.40833e-08

5.84754e-08

1.66e-06

1.69429e-08

6.17717e-08

3.56245e-07

1.84569e-06

5.7138e-07

1.14278e-07

3.11539e-07

2.55696e-08

2.40306e-07

1.32582e-08

9.47879e-07

1.71472e-07

2.48539e-08

3.38443e-08

5.84754e-08

1.32635e-07

2.49195e-07

1.74703e-08

1.43652e-06

4.49935e-08

1.3152e-07

1.53407e-07

1.42475e-07

1.41741e-08

3.69862e-07

1.10097e-08

6.16137e-08

2.93987e-07

1.10097e-08

1.08446e-08

3.39228e-07

6.1515e-08

1.10097e-08

1.74703e-08

1.93251e-08

2.48539e-08

6.18747e-07

1.26302e-07

2.74576e-08

5.75626e-08

7.37131e-07

2.01073e-08

1.25412e-07

5.46128e-07

7.05454e-07

7.86104e-10

2.49181e-07

1.14278e-07

1.10097e-08

1.20316e-08

2.64432e-07

3.40833e-08

7.86104e-10

2.34411e-07

2.14846e-06

1.78195e-07

3.39228e-07

7.1133e-08

6.54261e-07

2.949e-09

6.23649e-07

1.1554614e-05
5.777307e-06

2.2211e-06

2.67326e-06

8.82947e-07

4.27855e-07

9.78477e-09
1.956954e-08

2.07662e-09

7.70815e-09

1.38442e-07

1.01664e-05

1.76513e-07

7.08042e-08

6.325501744e-05

2.34434228e-05
4.68868456e-05

9.88451e-08

9.88451e-08

7.68351e-06

8.92948e-08

1.6613e-08

7.68351e-06

8.92948e-08

7.68351e-06

9.9678e-07

1.536702e-05
7.68351e-06

7.68351e-06

4.37184e-09
2.18592e-09

2.18592e-09

9.4049e-07

0.000238875

3.01634e-06

2.37643e-05

2.14889e-09

3.27052e-06

4.9719e-07

1.47832e-08

3.32089e-06

1.22e-07

4.008222e-07
2.004111e-07

1.55088e-07

4.53231e-08

1.03831e-07

0.000100768

3.16518e-05

1.5299e-07

1.10003e-06

2.49195e-07

1.745054e-05
8.72527e-06

8.72527e-06

2.58934e-06

3.77154e-08

1.06475e-06

3.68917e-05

3.63494e-08

1.37717e-06
2.75434e-06

1.37717e-06

3.29126e-06

2.36081e-06

1.74771e-06

1.01299e-09

2.0219493e-06
4.0438986e-06

1.76047e-07

6.66364e-08

2.83263e-08

1.42172e-07

6.82154e-08

1.77365e-07

1.4379e-07

1.3934e-07

2.83263e-08

1.72003e-07

4.2989e-08

1.64891e-07

1.49723e-07

9.55338e-08

4.76037e-08

1.89531e-07

1.21241e-07

6.82154e-08

1.72298e-06

1.25651e-06

4.21574e-07

8.41689e-08
1.683378e-07

8.41689e-08

5.81583e-06

3.61188e-07

8.20482e-09

2.16797e-06

1.38547e-06

0.00404264

1.81871e-07

1.224e-06

8.76446e-08

4.67919e-05

1.67145e-06

1.600023e-05

1.42604e-06
2.85208e-06

1.42604e-06

8.68592e-07
4.34296e-07

4.34296e-07

1.19284e-06

5.543359e-06
1.1086718e-05

8.91617e-07

5.37357e-07

5.1307e-07

9.57578e-07

6.71706e-07

1.20603e-06

7.66001e-07

1.8812296e-05
3.7624592e-05

2.06181e-06

2.10321e-06

1.12614e-06

1.11195e-06

9.59379e-07

1.04898e-05

9.60007e-07

1.04422e-06

5.43127e-07

3.2303e-07

1.72642e-07

6.35011e-08

2.81576e-09

2.85544e-07

5.52628e-07

1.42438e-08

4.77932252e-05
9.55864504e-05

4.65385e-06

3.65096e-06

4.26997e-06

4.29152e-06

3.86185e-06

4.26997e-06

4.26997e-06

4.26997e-06

1.81034e-06

1.64152e-08

4.29659e-06

3.86185e-06

4.26997e-06

5.56898e-06

4.71721e-06

5.00077e-06

1.03675e-06

5.09465e-08

1.02318e-07

0.0197461
0.0394922

0.0197461

3.2585e-05

1.91677e-07

7.30797e-07

1.82549e-06

2.97914e-06

1.255062e-06
6.27531e-07

3.62717e-07

2.64814e-07

2.16039e-07

2.393762e-06
4.787524e-06

1.10624e-07

3.28705e-07

7.02111e-07

1.4758e-07

3.49471e-07

3.12092e-07

1.03073e-07

3.40106e-07

5.502624e-05
2.751312e-05

3.97417e-06

4.35634e-06

4.15759e-06

6.76243e-06

3.97417e-06

4.28842e-06

1.49517e-06

3.90713e-07

5.53766e-07

1.35154e-07

3.72082e-07

0.000115791

9.12713e-06

1.17309e-05

9.60958e-07

3.12385e-06

4.31921e-07

1.67143e-07

8.3065e-08

1.4303894e-06
7.151947e-07

9.34481e-08

5.1442e-07

8.3065e-08

2.42616e-08

0.000395451

1.82143e-07

5.361578e-07
2.680789e-07

2.24238e-07

4.38409e-08

2.3812e-07

4.83622e-08
2.41811e-08

2.41811e-08

5.377968e-07
1.0755936e-06

5.55998e-08

1.28123e-07

1.36314e-07

2.1776e-07

0.02341982
0.01170991

0.00515436

0.00655555

1.72075e-06

1.36218e-06

3.50704e-07

9.08986e-06

2.67343e-05

3.3226e-07

1.69007e-07

2.21577e-07

6.27912e-08

5.11029e-05

8.3065e-08
1.6613e-07

4.15325e-08

4.15325e-08

1.91723e-06

3.13182e-07
1.56591e-07

1.56591e-07

4.65131e-07

3.44422e-06

3.223597e-07
6.447194e-07

2.21491e-07

4.55076e-08

5.53611e-08

5.59423e-07

8.05944e-07

2.79379e-06

1.60511e-06

6.88045e-07

9.95163e-06

1.541274e-05
7.70637e-06

7.70637e-06

0.000348436

3.16607e-05

1.33155e-06

1.974932e-06
3.949864e-06

9.87466e-07

9.87466e-07

4.52699e-07

5.10334e-08

1.74138e-06

6.8972e-08

6.10108e-05

1.69545e-07

3.62163e-07
7.24326e-07

3.62163e-07

3.3226e-08

2.04971e-07

1.38442e-08

3.10548e-06

6.25222e-06

2.95748e-07

3.77568e-09

6.12182e-08

3.2134263e-06
6.4268526e-06

4.65974e-08

2.27274e-06

7.49869e-08

8.19102e-07

9.7179143e-06
1.94358286e-05

1.04919e-08

2.2523e-06

2.17135e-06

4.73584e-09

2.91682e-06

2.16924e-06

8.87062e-08

3.41136e-09

6.57871e-08

3.50719e-08

4.36458e-06

2.73186e-06
1.36593e-06

1.36593e-06

4.7648e-08

3.98489e-07

3.3226e-07

5.9413642e-06
2.9706821e-06

1.3276e-07

3.94617e-07

2.84182e-08

2.84182e-08

1.32558e-07

2.84182e-08

1.58407e-08

5.53766e-08

1.55254e-06

7.12174e-08

1.58407e-08

7.22654e-08

7.10157e-08

3.71396e-07

7.78301e-06

1.39271e-08

0.000221067

1.15397e-06

1.93555e-07

6.8177e-06

1.37967e-06

6.126611e-06
1.2253222e-05

7.5676e-07

1.37445e-06

7.5676e-07

6.65207e-07

6.89614e-07

1.88382e-06

2.28109e-08

3.32565e-07

7.4739886e-06

1.22688e-06
2.45376e-06

1.22688e-06

8.20706e-07
1.641412e-06

8.20706e-07

3.3788166e-06
1.6894083e-06

1.58336e-07

7.3064e-07

5.81203e-08

7.42312e-07

3.77465e-05

6.87709e-07
1.375418e-06

5.75751e-07

1.11958e-07

2.2741e-07

4.46942e-06

3.48641e-08

1.476054e-08
2.952108e-08

2.28107e-09

2.28107e-09

3.35519e-09

2.28107e-09

2.28107e-09

2.28107e-09

2.0122e-07

3.00366e-05

5.18798e-08

1.77157e-07

2.81841e-07

1.89863e-08

2.47337e-05

2.05959e-09

1.74971e-06
3.49942e-06

1.74971e-06

3.37611e-07
6.75222e-07

1.8206e-08

1.07934e-07

1.17729e-07

9.3742e-08

6.37939e-06

5.58069e-06

8.52908e-08
4.26454e-08

4.26454e-08

5.7361e-06

1.05803e-07

4.222474e-07
8.444948e-07

6.92208e-08

6.92208e-08

2.14585e-07

6.92208e-08

1.53824e-09

4.85018e-06

4.05383e-07

4.494387e-05
2.2471935e-05

3.86602e-07

4.08622e-07

5.86379e-07

2.96489e-06

1.43023e-06

5.83232e-07

3.19999e-06

3.85848e-07

1.16128e-06

1.1967e-06

6.53476e-07

3.35986e-06

2.9632e-06

5.13561e-07

4.60001e-07

1.67942e-07

9.38934e-07

3.74865e-07

7.36323e-07

4.07367e-07

5.46191e-06

7.64455e-08

2.90727e-07

6.267e-07

0.000243776

5.66074e-06
1.132148e-05

5.66074e-06

3.56126e-06
1.78063e-06

1.04407e-06

7.3656e-07

2.51804e-06
1.25902e-06

2.88429e-07

1.45573e-07

2.86661e-07

2.45195e-07

2.93162e-07

1.47832e-08

4.81363e-08

2.1417e-05

4.15325e-09

8.79587e-06

0.0013521

5.5465291e-05

1.0480551e-06
2.0961102e-06

6.70069e-07

6.82099e-08

2.40083e-07

6.96932e-08

8.424994e-07
1.6849988e-06

6.78966e-08

1.27998e-07

1.01123e-07

7.86843e-08

1.16051e-07

8.72785e-08

1.62357e-07

1.01111e-07

1.6610313e-05
3.3220626e-05

4.07307e-07

1.39458e-05

1.42304e-07

3.28403e-07

5.09743e-07

2.45897e-07

1.74251e-07

8.56608e-07

1.460486e-05
7.30243e-06

3.66212e-06

3.64031e-06

3.858696e-06
1.929348e-06

8.75525e-07

7.76721e-07

2.77102e-07

1.52879e-07

2.949e-09

0.000696567

1.7872e-07

3.22699e-05

4.7255e-05
2.36275e-05

1.16999e-05

1.19276e-05

1.86767e-07
9.33835e-08

9.33835e-08

1.6613e-07

8.3332373e-07
1.66664746e-06

2.91939e-07

3.04973e-09

2.7443e-08

3.36521e-07

1.74371e-07

1.8996712e-05
9.498356e-06

4.63914e-07

4.53942e-07

4.51499e-07

4.53942e-07

3.68434e-07

4.36529e-07

4.53942e-07

4.53942e-07

5.34564e-07

4.51499e-07

4.63914e-07

4.53942e-07

4.53942e-07

4.38972e-07

4.51499e-07

4.51499e-07

4.53942e-07

4.51499e-07

4.53942e-07

4.51499e-07

4.51499e-07

5.85911e-06

4.50506e-06
9.01012e-06

4.50506e-06

8.41803e-07
1.683606e-06

5.81455e-08

5.81455e-08

5.81455e-08

6.09221e-07

5.81455e-08

5.31409e-06

1.01245e-05

3.9619e-08
1.98095e-08

1.98095e-08

1.0450922e-05
5.225461e-06

1.35242e-06

5.17753e-07

2.33346e-07

1.29219e-06

6.4707e-07

2.47882e-07

9.348e-07

9.22944e-09

0.00327041

1.2193e-07
2.4386e-07

1.2193e-07

4.82694e-06

0.00023846

1.406262e-06
7.03131e-07

7.03131e-07

2.79607e-07

1.22148e-07

1.4828983e-05
2.9657966e-05

5.41384e-08

2.73063e-07

3.17791e-07

2.21767e-07

2.10698e-08

5.26065e-07

7.50057e-07

1.06857e-06

1.18664e-08

5.76624e-08

3.17791e-07

4.6587e-08

2.54861e-07

5.0813e-07

2.82032e-07

1.01219e-06

4.23303e-07

1.02309e-08

3.17791e-07

1.92088e-06

8.83416e-08

5.16399e-07

1.04113e-07

1.28308e-07

5.2842e-07

3.29657e-07

5.18199e-07

3.78872e-07

1.5385e-07

3.82092e-07

5.84535e-08

6.73533e-08

4.05009e-07

1.31325e-07

1.44485e-07

1.86267e-08

4.6587e-08

9.42412e-07

2.05595e-07

1.38468e-07

2.05125e-07

3.59323e-07

2.00031e-07

2.05595e-07

1.76497e-07

3.32897e-08

1.6613e-07

2.70878e-06
5.41756e-06

1.31774e-06

1.39104e-06

8.50194e-08
1.700388e-07

8.50194e-08

1.7645e-06

6.34039e-08

1.92269e-06

4.53992e-08

5.93321e-09

7.37122e-09

2.11176e-08
4.22352e-08

2.11176e-08

1.02565e-06

2.3496049e-06

7.11985e-09
1.42397e-08

7.11985e-09

7.963405e-07
1.592681e-06

4.15325e-08

2.56418e-07

4.9839e-07

2.110686e-07
1.055343e-07

5.6882e-08

4.86523e-08

5.316156e-07
2.658078e-07

8.86026e-08

8.86026e-08

8.86026e-08

2.67775e-06

5.26906e-07
2.63453e-07

6.12182e-08

3.56598e-08

1.66575e-07

2.62035e-10

1.53824e-09

2.42213e-06

1.19983e-06

2.749056e-05
1.374528e-05

2.95737e-06

2.89273e-06

2.08574e-06

5.33195e-06

4.7749e-07

5.77078e-06

3.60675e-07

1.70986e-07

1.53824e-09

1.34335e-06

1.47832e-08

5.54412e-08

1.50252e-06

6.40956e-09

3.52828e-07

9.18223e-07

6.11851e-06

6.51005e-07
1.30201e-06

6.51005e-07

1.9277473e-06
3.8554946e-06

3.55993e-08

1.7876e-06

1.04548e-07

6.92119e-06

4.83438e-06

1.45018e-06

1.430774e-06
7.15387e-07

3.41531e-07

3.73856e-07

2.56241e-05

1.0266332e-05
5.133166e-06

1.93818e-07

3.11573e-07

3.74703e-07

1.80884e-07

3.74703e-07

3.74703e-07

2.76883e-07

4.22978e-07

3.74703e-07

3.74703e-07

3.74703e-07

3.74703e-07

3.74703e-07

3.74703e-07

3.74703e-07

8.05944e-07

7.45419e-07

4.00078e-07

3.49489e-05

1.6392578e-06
8.196289e-07

2.76883e-08

2.76883e-08

2.76883e-08

7.36564e-07

1.28748e-06

5.04854e-07

9.60271e-06

2.76883e-08

7.04857e-07

4.69256e-07
2.34628e-07

1.20246e-07

1.14382e-07

1.40749e-07

2.11604e-05

3.93264e-07

4.2586e-08
8.5172e-08

4.2586e-08

3.4116e-08

3.78091e-07

1.03831e-08

3.91592e-08

0.000238294

2.89469e-08

1.52803495e-05
5.4379812e-06

1.28165e-07

1.0162566e-06

4.848054e-07
2.424027e-07

6.60977e-08

1.76305e-07

5.314512e-07
2.657256e-07

3.72376e-08

2.28488e-07

8.82627e-07

2.28021e-08

7.7925176e-06

1.9461044e-06
3.8922088e-06

2.06557e-07

7.84355e-07

6.69445e-07

2.06557e-07

7.91904e-08

1.9501544e-06
3.9003088e-06

7.91904e-08

7.14782e-08

7.18858e-07

5.97394e-08

5.97394e-08

7.13321e-07

7.91904e-08

1.06879e-07

6.17586e-08

5.13673e-07

3.46935e-06

1.53384e-06

5.69588e-09

3.57579e-06

1.24597e-07

6.27382e-07

0.0170202851573
0.035487393857386

2.76883e-08

3.08033e-07

6.68949e-07

3.81568e-08

5.28126e-07

0.00041286006324

4.49935e-08

3.18438e-07

0.000161893288
0.000323786576

7.88911e-07

9.47354e-06

7.88911e-07

1.65962e-06

4.13381e-06

2.1111e-05

1.20856e-05

1.36625e-06

1.65962e-06

4.11394e-06

1.73408e-06

1.38034e-06

1.66903e-05

1.38034e-06

7.75066e-07

7.88911e-07

4.14717e-06

4.83111e-06

9.05164e-06

7.02197e-06

2.31318e-06

1.20523e-06

9.24574e-07

4.11394e-06

1.36625e-06

1.13636e-05

3.89256e-06

7.88911e-07

4.86594e-07

7.02197e-06

7.02197e-06

1.769e-06

1.38034e-06

9.17162e-06

1.53502e-06

1.36537e-06

1.19103e-06

1.9179078e-06
9.589539e-07

9.38977e-07

1.99769e-08

7.38969e-06
1.477938e-05

1.21459e-06

1.21459e-06

1.31842e-06

1.19026e-06

1.21459e-06

1.23724e-06

1.77363e-09
3.54726e-09

1.77363e-09

1.009762266e-05
5.04881133e-06

1.20444e-06

1.20444e-06

1.20444e-06

4.15325e-08

5.53766e-08

9.54473e-09

1.20444e-06

4.15325e-08

8.3065e-08

3.08840912e-06
1.54420456e-06

8.3065e-08

5.53766e-08

2.65249e-07

1.77833e-07

2.58948e-08

8.3065e-08

5.53766e-08

5.19156e-09

1.08656e-07

6.84497e-07

4.11394e-06

4.13381e-06

3.79476e-06

1.14709e-08

1.6613e-07

4.14717e-06

3.60896e-05

4.14717e-06

2.219138e-06
1.109569e-06

6.4133e-08

8.32362e-08

4.2765e-08

9.50211e-08

4.54054e-08

3.34273e-07

7.62303e-08

3.2574e-07

4.2765e-08

5.92223e-07

7.45491e-07

2.061797e-06
4.123594e-06

4.02422e-07

7.43099e-07

5.5852e-07

3.57756e-07

2.79904e-06

2.86113e-07

3.6172961e-05

2.888236e-06
1.444118e-06

6.05306e-07

8.38812e-07

2.5910725e-06
5.182145e-06

8.45877e-08

9.60048e-08

1.11101e-06

3.5209e-07

5.77175e-07

3.70205e-07

2.810258e-05
1.405129e-05

2.20498e-06

2.10896e-06

2.65513e-06

1.64173e-06

2.18753e-06

1.51334e-06

1.73962e-06

1.32228e-05

7.1282455603e-05
0.000142564911206

5.01903e-10

1.93957e-05

1.17232e-07

5.52382e-06

2.7246e-05

5.52381e-08

2.66741e-06

5.72526e-07

5.26319e-06

3.9769e-08

3.9769e-08

6.03779e-07

9.73224e-06

2.52806e-08

4.84045e-08

6.28255e-06

4.37184e-09

3.01424e-08

6.198524e-07
3.099262e-07

1.03831e-07

3.71962e-08

1.68899e-07

3.3931e-09

5.19156e-09

1.33694e-06

1.1783732e-05
5.891866e-06

6.39925e-07

5.51031e-07

7.11255e-07

1.66302e-07

5.51031e-07

4.96831e-07

7.66639e-07

1.58012e-06

1.05012e-07

1.19454e-07

2.04266e-07

6.38677e-07

2.056763e-06
4.113526e-06

4.56857e-07

6.03127e-07

8.3065e-08

4.56857e-07

4.56857e-07

2.1642e-07

1.20523e-06

2.32186e-05

2.36349e-07

1.8078e-08

6.92209e-07
3.461045e-07

4.15325e-08

3.04572e-07

3.06271e-07

0.00049076

3.46104e-07

3.03966e-07

1.77103e-06

4.78343e-08

1.552481e-05
3.104962e-05

7.71541e-06

7.8094e-06

3.46935e-06

2.17692e-05

3.12353e-07

4.67677e-07

8.43914e-06

1.77844e-07

4.36481e-07
8.72962e-07

4.36481e-07

0.00015093898
7.546949e-05

2.03495e-06

2.03673e-06

3.26445e-05

2.07233e-06

3.26449e-05

2.03673e-06

1.99935e-06

0.0167881058198

0.002059
0.0010295

0.0010295

8.3065e-08

1.65259748e-05
8.2629874e-06

1.18664e-08

4.56857e-07

1.38442e-07

1.38442e-07

7.51738e-06

9.9678e-07
4.9839e-07

4.9839e-07

0.0147115

5.19156e-09

6.57598e-08

2.07662e-07

3.80714e-08

2.49195e-07

3.8156154e-06
1.9078077e-06

1.68322e-06

7.48065e-08

8.9528e-08

6.02532e-08

6.5585e-06

3.82099e-06

2.07662e-07

8.82027e-07

2.31455e-07

1.46341e-07

1.03831e-07

2.03129e-07

5.89761e-06

2.48584e-05

5.28823e-08

5.19156e-09

2.25307e-06

2.1642e-07

3.11494e-08

1.951875e-05
3.90375e-05

1.00163e-05

1.00865e-06

1.00865e-06

1.00865e-06

1.10002e-06

1.00865e-06

4.36783e-06

3.93583e-06

6.498534e-07
1.2997068e-06

1.13418e-07

1.13418e-07

8.27634e-08

1.13418e-07

1.13418e-07

1.13418e-07

2.07662e-08

2.68081e-06

9.81226e-06

1.86211e-07

9.55247e-06
1.910494e-05

9.55247e-06

3.19445e-05

7.06052e-07

3.64204e-05

1.52506e-08

4.41773e-06

1.99769e-08

3.9176676e-06
7.8353352e-06

3.87155e-06

4.61176e-08

1.38442e-07

2.59121e-06
5.18242e-06

1.30154e-06

1.28967e-06

7.88911e-07

1.93818e-07

4.15325e-08

1.55508e-06

8.55821e-08

8.3065e-08
4.15325e-08

4.15325e-08

1.64234e-05

8.3065e-08

1.78485e-07

7.51392e-07

1.50308e-07
7.5154e-08

7.5154e-08

5.12881e-06

1.790512e-06
8.95256e-07

4.58011e-07

4.37245e-07

1.8078e-08

1.15909e-07

1.27792e-08

2.26541e-08

8.97793e-09

2.5308e-05

3.19722e-05

8.52509e-09

5.31616e-08

6.12703e-07

4.5835e-06

8.43087e-07

9.16375125e-06
3.05458375e-06

8.3065e-08
1.6613e-07

8.3065e-08

2.97151875e-06
5.9430375e-06

8.00925e-07

8.74499e-07

1.85299e-07

3.74756e-07

5.73825e-07

9.84657e-08

7.91095e-09

1.6613e-08

3.92251e-08

1.24597e-07

2.76883e-08

0.00105407

1.10753e-07
3.32259e-07

1.10753e-07
2.21506e-07

1.10753e-07

0.000210804

6.1109201e-07
1.83327603e-06

1.94871e-09
3.89742e-09

1.94871e-09

8.3065e-07
4.15325e-07

4.15325e-07

1.6613e-07
3.3226e-07

1.6613e-07

2.76883e-08
5.53766e-08

2.76883e-08

2.99374e-09

3.042325e-06
1.084252e-06

1.747642e-06
8.73821e-07

8.73821e-07

2.10431e-07

0.000175586

9.052087e-06
3.017408175e-06

8.32886e-08
4.16443e-08

4.16443e-08

2.61622e-09
1.30811e-09

1.30811e-09

7.44124e-08
1.488248e-07

7.44124e-08

8.3065e-08
1.6613e-07

8.3065e-08

2.81684084e-06
5.63368168e-06

4.15325e-08

6.22987e-08

4.46956e-08

2.18985e-06

8.3065e-08

2.76883e-08

3.69488e-08

1.28347e-08

3.32934e-09

1.24597e-07

8.3065e-09

8.31649e-08

8.3065e-08

1.54645e-08

1.37525e-10

1.24597e-07

1.21639324e-07
3.64917972e-07

3.3125644e-08
6.6251288e-08

9.22944e-10

3.22027e-08

1.089736e-08
5.44868e-09

3.18277e-09

2.26591e-09

1.6613e-07
8.3065e-08

8.3065e-08

4.0387002e-08
1.21161006e-07

1.825604e-09
9.12802e-10

9.12802e-10

1.69683e-08
3.39366e-08

1.69683e-08

2.25059e-08
4.50118e-08

2.25059e-08

2.05126e-07

2.593557e-06
8.64519e-07

1.21841e-06
6.09205e-07

3.04006e-07

1.96988e-07

1.08211e-07

5.10628e-07
2.55314e-07

2.55314e-07

2.90727e-07

8.3065e-08

0.000125323

1.6851e-07

1.38442e-08

3.3236e-08

2.05795e-07

5.8833862e-06
2.5479351e-06

9.69091e-08

6.10306e-07

3.7212e-07

5.594e-07

1.21684e-07

7.87516e-07
1.575032e-06

1.7721e-07

6.10306e-07

6.322187e-06
2.813279e-06

2.11765e-06

1.391258e-06
6.95629e-07

6.95629e-07

2.21507e-07

1.90008e-06

8.3065e-08

4.444671e-06
1.481557e-06

2.03748e-06
1.01874e-06

1.01874e-06

4.62817e-07
9.25634e-07

4.62817e-07

9.65821e-06

2.76883e-08

6.41196e-07

3.334464e-06
8.969036e-07

1.9312596e-06
6.437532e-07

1.2875064e-06
6.437532e-07

2.07662e-08

3.3226e-07

2.90727e-07

2.531504e-07
5.063008e-07

6.32876e-08

6.32876e-08

6.32876e-08

6.32876e-08

3.324345061e-07
1.661881687e-07

5.81687e-11
1.163374e-10

5.81687e-11

1.6613e-07

1.36852e-08

5.73148e-07

1.6613e-07

4.04750568e-05
1.48843211e-05

1.28374e-06

1.57958892e-05
7.8979446e-06

2.45042e-07

9.48294e-07

5.19669e-07

1.63462e-06

9.94606e-07

1.0505e-06

7.44383e-07

5.53766e-08

9.54148e-07

7.51306e-07

4.15325e-08

2.80847e-06
5.61694e-06

1.25972e-06

1.1363e-06

2.50473e-07

1.61977e-07

1.29736e-06

8.23013e-07

7.32261e-07

3.9317347e-07
1.17556494e-06

7.78436e-07
3.89218e-07

1.94609e-07

1.94609e-07

3.95547e-09

2.41887e-06

2.28463e-06

1.19517634e-06
4.0461618e-07

3.69178e-08
7.38356e-08

3.69178e-08

1.86722e-08

6.9805236e-07
3.4902618e-07

5.90684e-08

1.32904e-07

4.88617e-09

6.38961e-09

7.16921e-08

7.40859e-08

9.84925e-07

1.6613e-07

1.29433497e-06
3.00602091e-06

1.6613e-08

8.60371e-07

4.1735097e-07
8.3470194e-07

8.3065e-08

8.3065e-08

8.3065e-08

2.02597e-09

1.6613e-07

7.15473e-07

3.81214764e-06
1.01533566e-06

5.1694566e-07
1.55083698e-06

1.03389132e-06
5.1694566e-07

6.20066e-09

5.10745e-07

8.3065e-08
1.6613e-07

8.3065e-08

4.9839e-07
1.6613e-07

1.6613e-07
3.3226e-07

1.6613e-07

2.49195e-07
5.81455e-07

8.3065e-08
1.6613e-07

8.3065e-08

1.6613e-07

1.33933e-08

8.96241e-07

8.374586e-08
2.5123758e-07

1.6613e-07
8.3065e-08

8.3065e-08

1.36172e-09
6.8086e-10

6.8086e-10

1.01299e-09

5.68014e-08

4.99393e-10

2.272923785e-05
8.54137806e-05

8.2951e-07

5.11169e-08

2.526788e-06
6.672005e-06

4.33591e-07

3.236858e-06
1.618429e-06

1.03831e-07

1.10753e-07

6.08913e-07

7.94932e-07

4.74768e-07

1.011251e-06
3.033753e-06

1.011251e-06
2.022502e-06

4.87341e-07

1.43979e-07

3.79931e-07

1.311838515e-05
3.935515545e-05

2.1031016e-06
1.0515508e-06

8.99265e-07

8.3065e-08

6.92208e-08

8.67182e-07
4.33591e-07

4.33591e-07

8.28616e-07
4.14308e-07

4.14308e-07

1.680733e-06
3.361466e-06

7.74654e-07

9.06079e-07

1.00537e-06
2.01074e-06

1.00537e-06

8.88256e-07
4.44128e-07

4.44128e-07

4.05272e-07
8.10544e-07

4.05272e-07

2.265908e-06
4.531816e-06

8.69836e-07

3.65486e-07

4.52737e-07

5.77849e-07

1.50171e-06
7.50855e-07

7.50855e-07

5.53766e-08
2.76883e-08

2.76883e-08

1.180262e-06
5.90131e-07

5.90131e-07

1.849716e-06
3.699432e-06

8.40526e-07

1.00919e-06

5.1071e-07
1.02142e-06

5.1071e-07

2.75444e-06
1.37722e-06

9.79512e-07

3.97708e-07

8.3065e-08
4.15325e-08

4.15325e-08

2.47955e-09
4.9591e-09

2.47955e-09

2.67192e-07
5.34384e-07

2.67192e-07

2.060856e-06
1.030428e-06

6.7048e-07

3.59948e-07

7.615809e-06
2.538603e-06

8.07095e-07
1.61419e-06

8.07095e-07

1.637224e-06
8.18612e-07

8.18612e-07

1.825792e-06

8.09065e-07
1.61813e-06

8.09065e-07

2.07662e-07
1.03831e-07

1.03831e-07

1.6613e-07

8.3065e-08

2.8171424e-06
1.3739608e-06

1.30474e-06

1.384416e-07
6.92208e-08

4.15325e-08

2.76883e-08

7.72165e-07

6.88253e-08

0.00127537

8.3065e-08

1.24597e-07

7.6912e-09

9.146271603e-06
2.7438814809e-05

9.198568186e-06
4.599284093e-06

6.63193e-10

5.89996e-08

6.04722e-08

5.52071e-08

1.50532e-07

1.87336e-06

1.21614e-06

1.63051e-07

1.69463e-07

8.51396e-07

1.31849e-09
2.63698e-09

1.31849e-09

9.09133804e-06
4.54566902e-06

3.19481e-09

1.10753e-07

1.43031e-08

3.55158e-07

8.3065e-08

2.90727e-07

5.53766e-08

3.95085e-08

2.79637e-07

2.57875e-08

6.6452e-07

1.08752e-07

1.08995e-07

3.1343e-08

1.94694e-08

3.35371e-08

1.78489e-07

6.64922e-07

3.12533e-07

1.3152e-07

8.06917e-08

1.36064e-08

8.97991e-09

4.66974e-07

4.0982e-08

4.22844e-07

4.48551e-07

8.44294e-07

9.22944e-09

0.000289932

1.24597e-07

8.460454e-07
2.889372e-07

5.36342e-07
2.68171e-07

2.68171e-07

2.07662e-08

1.81279e-06

1.0065e-06

1.06841e-08

4.23362e-07
8.46724e-07

4.23362e-07

9.13715e-08

8.3065e-08

3.73668e-06

7.50754e-08

3.42147e-07

5.26537e-08

7.38039e-07

3.48518e-08

2.58755e-06

4.99393e-10

1.24597e-07

3.638754826e-05
9.42321469e-06

4.8566e-07
1.373915e-06

8.3065e-08

8.0519e-07
4.02595e-07

4.02595e-07

3.20054e-07

2.581068e-06
8.60356e-07

8.60356e-07
1.720712e-06

8.60356e-07

7.75714469e-06
2.268929657e-05

1.081797e-06

9.25406e-07
4.62703e-07

4.62703e-07

1.56391e-07

1.385035488e-05

1.148006e-06
5.74003e-07

3.99567e-07

1.74436e-07

5.887788e-06
1.1775576e-05

7.83984e-07

2.78492e-07

3.25767e-07

3.48873e-06

2.26831e-07

7.83984e-07

4.15325e-08
8.3065e-08

4.15325e-08

4.15325e-08

3.84214e-07

2.0898069e-07
4.1796138e-07

2.07662e-08

1.03831e-07

8.3065e-08

1.31849e-09

1.48605e-06

2.07662e-08

1.90099e-08

4.81536e-09

4.585358e-06
1.1235202e-05

1.24189e-06
2.48378e-06

1.24189e-06

1.06166e-07

1.61682e-06

1.645192e-06
8.22596e-07

4.78919e-07

3.43677e-07

7.97886e-07

0.000140023

1.6613e-07

1.39236681e-06
5.56702415000001e-06

9.452817e-07
2.8358451e-06

1.8905634e-06

7.874582e-07
1.5749164e-06

1.38442e-08

7.73614e-07

4.15325e-08
8.3065e-08

4.15325e-08

8.3065e-08
1.6613e-07

8.3065e-08

3.3226e-08
1.6613e-08

1.6613e-08

3.3226e-08
1.6613e-08

1.6613e-08

4.4708511e-07
1.33881224e-06

2.44309e-09

4.4464202e-07
8.8928404e-07

2.44309e-09

2.44309e-09

2.44309e-09

2.44309e-09

2.44309e-09

2.44309e-09

2.44309e-09

2.44309e-09

3.34703e-07

2.44309e-09

2.44309e-09

2.44309e-09

2.44309e-09

2.44309e-09

2.44309e-09

2.44309e-09

2.44309e-09

2.44309e-09

2.44309e-09

2.44309e-09

4.15325e-08

2.44309e-09

2.44309e-09

2.44309e-09

2.44309e-09

2.44309e-09

2.44309e-09

2.44309e-09

2.44309e-09

2.44309e-09

5.55e-10
1.85e-10

1.85e-10
3.7e-10

1.85e-10

8.97008e-07
2.691024e-06

2.61244e-07
5.22488e-07

2.61244e-07

6.35764e-07
1.271528e-06

6.35764e-07

1.01397e-07

4.29898e-08

9.69091e-08

6.610083e-06
2.203361e-06

2.094846e-06
4.189692e-06

4.86735e-07

5.24401e-07

5.24048e-07

5.59662e-07

1.08515e-07
2.1703e-07

1.08515e-07

2.12673e-06

0.0001696696088
6.19926226e-05

5.99131e-07

2.50963e-06
5.01926e-06

2.50963e-06

1.66386e-06

5.70015e-05
2.850075e-05

5.2806e-06

8.84106e-06

1.51711e-06

2.69648e-06

6.06428e-06

4.10122e-06

7.144497e-06
1.4288994e-05

1.35637e-07

3.21847e-07

1.92646e-07

3.23414e-07

1.78909e-07

5.07564e-07

5.7026e-07

8.38285e-07

2.82356e-06

4.78262e-07

7.74113e-07

6.51057e-06

6.46904e-06

2.86915e-07

5.89116e-06
1.178232e-05

5.89116e-06

3.2766532e-06
1.6383266e-06

1.62212e-06

1.62066e-08

5.95543e-07

1.832e-07

5.03127562e-06
1.76039454e-06

2.85484308e-06

1.668222e-07
8.34111e-08

8.34111e-08

8.3065e-08
1.6613e-07

8.3065e-08

2.15968e-06
1.07984e-06

1.07984e-06

3.6221088e-07
1.8110544e-07

9.51459e-09

1.6613e-07

5.46085e-09

1.6613e-07

1.6613e-07
8.3065e-08

8.3065e-08

2.49908e-07

1.6613e-07

4.15325e-08

8.42925e-08

4.15325e-07

2.00208e-07

1.48337e-06

4.57199e-09

1.938183e-07
6.46061e-08

1.292122e-07
6.46061e-08

6.46061e-08

1.14486e-06

6.08373836e-07
1.666257021e-06

4.51202e-08

5.98947e-09

8.38451e-09

2.88352e-08

3.99929e-08

4.53907e-10

3.00883e-08

8.99018698e-07

1.11356e-09
5.5678e-10

5.5678e-10

1.15203e-07
2.30406e-07

1.15203e-07

8.3065e-08
4.15325e-08

4.15325e-08

2.0187242e-08
4.0374484e-08

1.38442e-08

5.94079e-09

4.02252e-10

1.15259e-07
2.30518e-07

1.15259e-07

2.366094e-07
1.183047e-07

4.15325e-08

7.67722e-08

3.40034e-08
1.70017e-08

1.70017e-08

2.1464427e-08
4.2928854e-08

7.16077e-10

4.48325e-09

1.62651e-08

1.6613e-07

1.10753e-07

1.16983e-06

9.89601e-07

4.15325e-08

5.36089e-08

8.3065e-08

1.54316e-07

7.28416e-08
3.64208e-08

3.64208e-08

3.64208e-08

1.68728e-07
5.06184e-07

1.68728e-07
3.37456e-07

1.68728e-07

2.49195e-07

8.06456e-10

3.21455e-06

8.3589861416e-05
2.7930908758e-05

3.02774e-10

2.75911e-05
5.51822e-05

1.74184e-05

1.01727e-05

1.82524e-07

5.38789e-08
1.077578e-07

5.38789e-08

8.3065e-08
1.6613e-07

3.46104e-08

3.46104e-08

1.38442e-08

3.02774e-10

9.35221e-09

1.03831e-08

1.12484e-06

1.24597e-07

8.3065e-08

3.04829e-07

3.346067e-06
1.0038201e-05

2.7763e-06
5.5526e-06

2.7763e-06

4.86702e-07
9.73404e-07

4.86702e-07

8.3065e-08
1.6613e-07

8.3065e-08

8.3065e-08

6.82119e-07

8.89982e-08

1.0800972e-06
2.1601944e-06

1.38442e-08

4.44622e-07

6.21631e-07

1.23768e-06

4.74163e-07

7.94317e-09

6.09143e-08

1.13061e-07

3.709323648e-05
9.74682112e-06

4.55114082e-06
1.365342246e-05

4.34901e-08
8.69802e-08

4.34901e-08

7.35421744e-06
3.67710872e-06

3.29491e-07

8.63013e-07

4.61472e-09

2.47999e-06

8.30542e-07
1.661084e-06

8.30542e-07

8.63959e-07

8.3065e-08

2.7349173e-06
8.2047519e-06

8.75483e-07
1.750966e-06

8.75483e-07

1.71818e-06
8.5909e-07

8.5909e-07

1.0003443e-06
2.0006886e-06

8.3367e-08

2.76883e-08

8.89289e-07

1.513739e-06
4.541217e-06

3.027478e-06

2.88596e-06
1.44298e-06

1.44298e-06

4.30707e-08
8.61414e-08

4.30707e-08

5.53766e-08
2.76883e-08

2.76883e-08

1.22154e-09

2.07662e-08

1.6613e-07
4.15325e-07

1.6613e-07
8.3065e-08

8.3065e-08

8.3065e-08

5.99254e-07

1.59208e-07

2.30806e-07

7.013661661e-05
2.503438293e-05

3.498025e-06

1.30944e-06

7.620325e-07
1.524065e-06

1.16149e-07

2.09124e-07

6.44295e-08

3.59948e-07

1.2382e-08

6.6452e-07
3.3226e-07

3.3226e-07

8.43835e-08

4.112672618e-05

1.34256e-06
2.68512e-06

1.34256e-06

1.28335e-07

2.9517e-06

8.3065e-08
4.15325e-08

4.15325e-08

1.236585e-06
2.47317e-06

5.5215e-08

1.18137e-06

9.5799e-08

8.5229138e-06
4.2614569e-06

2.29161e-07

2.46594e-08

3.45421e-07

2.93948e-07

3.31998e-06

4.82875e-08

6.3319e-08
3.16595e-08

3.16595e-08

1.52844e-06
3.05688e-06

1.52844e-06

4.47900445e-06
8.9580089e-06

1.02112e-07

7.38278e-07

7.80994e-07

2.812e-07

5.85016e-08

8.19587e-07

1.39904e-07

5.33872e-07

1.0179e-06

6.65585e-09

3.77568e-09

1.216968e-07
6.08484e-08

6.08484e-08

2.020147e-07
4.040294e-07

2.92131e-08

9.84701e-08

7.43315e-08

1.10834876e-05
5.5417438e-06

1.41623e-06

9.72138e-08

3.35678e-06

5.09161e-07

1.62359e-07

2.47713e-07
4.95426e-07

1.38442e-07

1.09271e-07

1.62945e-07

2.30154e-07

1.24129e-08

1.7740503e-05
5.913501e-06

2.894081e-06
5.788162e-06

2.2162e-06

3.12533e-07

3.65348e-07

3.01942e-06
6.03884e-06

3.01942e-06

8.3065e-08

2.22344e-07

5.48229e-07
1.82743e-07

3.65486e-07
1.82743e-07

1.6613e-07

1.6613e-08

6.2522e-10

6.895042436e-05
2.302934812e-05

1.3762e-07

1.33626296e-05
6.6813148e-06

5.30685e-07

1.51239e-07

1.44734e-08

4.71165e-08

1.61987e-07

1.47331e-08

1.56628e-07

3.08933e-07

1.21559e-07

5.4765e-07

1.07984e-07

1.82846e-06

1.89255e-06

1.05164e-07

1.50555e-07

2.41554e-07

2.70183e-07

2.98608e-08

3.52757e-07
7.05514e-07

3.52757e-07

1.585765632e-05
3.171531264e-05

3.43502e-07

1.85309e-07

5.06939e-07

2.45015e-07

6.89756e-07

3.81116e-07

4.45756e-07

9.28907e-08

1.74168e-07

4.38284e-07

6.46611e-07

7.27472e-09

3.7167e-08

1.43905e-07

1.35716e-07

1.43723e-07

9.52469e-07

2.11884e-06

1.13381e-07

2.82021e-07

3.35967e-07

1.49018e-06

1.20831e-07

8.45531e-08

1.07708e-07

4.25528e-07

2.41202e-07

1.77417e-07

1.38442e-08

6.17964e-07

1.57306e-07

2.43777e-07

1.10372e-06

3.72501e-07

2.7659e-07

1.37491e-07

3.69319e-07

1.35539e-08

4.54657e-08

1.24032e-07

2.75077e-07

1.32773e-07

3.27959e-07

3.17993e-07

2.61061e-07

2.68608e-07

1.70076e-06

1.6613e-07

1.654692152e-05
7.06928762e-06

4.65164e-06

4.81669256e-06

1.1683724e-07
5.841862e-08

7.04794e-09

7.04794e-09

7.04794e-09

3.72748e-08

2.33618e-06
4.67236e-06

2.33618e-06

1.374766e-08
2.749532e-08

3.54546e-09

1.02022e-08

9.30134e-09

3.03437e-06

7.97836e-07

1.02865e-08

1.93015e-07

0.0150591009605
0.060227446834

1.924933e-05
5.774799e-05

1.349588e-05
2.699176e-05

2.48909e-06

4.5397e-06

2.50922e-06

3.95787e-06

9.408296e-06
4.704148e-06

9.94908e-07

3.70924e-06

2.098604e-06
1.049302e-06

4.40159e-07

1.93818e-07

4.15325e-07

0.0150384715695
0.0451064577005

0.0013649985902

4.0645706e-06
8.1291412e-06

6.10546e-07

2.00987e-07

5.0271e-07

1.17826e-06

8.21715e-07

3.59936e-08

7.14359e-07

0.00067486293
0.00134972586

0.000665822

2.49277e-06

6.54816e-06

7.80527e-07

2.780614e-06
1.390307e-06

1.96982e-07

5.55551e-07

2.48217e-07

3.89557e-07

2.3028e-07

3.352168e-06
1.676084e-06

1.1811e-06

4.94984e-07

0.028599257081

4.11212e-06

0.02854013004
0.01427006502

0.0140435

3.31479e-05

0.000138591

6.0132e-07

5.42248e-05

2.05791e-07

1.9826822e-05
9.913411e-06

7.63931e-07

1.98245e-06

7.16703e-06

6.710407e-06
1.3420814e-05

2.46487e-07

4.88713e-06

1.57679e-06

3.62829e-06

1.7933204e-05
8.966602e-06

1.08232e-06

2.90643e-07

2.69278e-06

3.07863e-06

9.14118e-07

2.02242e-07

7.05869e-07

0.0001037304598

3.828282e-07
1.914141e-07

3.75001e-08

1.53914e-07

2.9734738e-05
5.9469476e-05

2.88218e-06

4.7296e-06

5.19028e-06

1.58913e-06

2.55882e-06

7.17986e-07

1.06025e-06

1.22174e-06

4.98882e-07

2.93855e-06

6.34732e-06

3.9072687e-06
7.8145374e-06

1.98822e-06

6.3534e-07

2.61282e-07

4.56747e-08

7.9769e-07

1.79062e-07

4.43014e-08
2.21507e-08

2.21507e-08

1.80096584e-05
3.60193168e-05

3.68991e-06

1.08953e-06

1.38442e-07

2.32447e-07

1.29026e-06

4.84342e-08

1.63448e-07

2.25462e-08

6.89163e-06

3.86517e-06

1.60119e-07

4.17722e-07

1.380061e-06
4.140183e-06

1.81781e-07
3.63562e-07

1.81781e-07

1.19828e-06
2.39656e-06

1.19828e-06

1.6613e-07
3.3226e-07

1.6613e-07

6.929328e-06
2.309776e-06

4.619552e-06
2.309776e-06

2.26831e-07

8.48725e-07

7.74684e-07

4.59536e-07

1.59208e-07

6.521528e-07
2.6086112e-06

6.521528e-07
1.9564584e-06

1.3043056e-06
6.521528e-07

1.6613e-07

1.6613e-07

6.65958e-08

2.53297e-07

8.018776e-05
4.009388e-05

7.29248e-06

1.81304e-05

1.4671e-05

1.79974e-07

9.09771e-07
3.03257e-07

3.67864e-07
1.83932e-07

1.83932e-07

1.19325e-07
2.3865e-07

1.19325e-07

8.0841e-05

1.6613e-08

2.39267e-08

1.16291e-07

1.422646242e-05
4.174779726e-05

4.09134e-07

1.412835324e-05

3.44128e-08
1.72064e-08

1.72064e-08

1.233560044e-05
6.16780022e-06

1.20962e-09

2.41799e-06

7.98545e-07

5.0381e-07

2.34919e-06

9.70556e-08

8.7917e-07
1.75834e-06

5.51525e-07

3.27645e-07

1.21168e-07

1.24613916e-05

6.1338406e-06
1.22676812e-05

1.89665e-06

1.03505e-06

5.53766e-08

2.63039e-07

2.49483e-06

3.88895e-07

9.68552e-08
1.937104e-07

9.68552e-08

4.01288e-07

1.81199e-06

0.000245595

8.3065e-08

2.07662e-08

6.2522e-10

1.24597e-07

1.04648e-08

8.80489e-06

1.59208e-07

3.09107e-07

2.76383e-08

1.80995e-07

4.61347e-07

2.76883e-08

1.02844e-06

1.03831e-07

2.75785878e-06
1.06260709e-06

1.0462242e-06
5.231121e-07

8.47481e-08

4.38364e-07

1.095325e-07
2.19065e-07

1.68088e-08

9.27237e-08

1.64316e-07

9.88869e-09

6.5628e-09

1.6613e-07

8.3065e-08

0.01339111943
0.09373783601

0.0746262

0.0746262
0.0124377

0.0621885

0.0621885

0.0124377
0.0621885

0.0497508

0.0497508
0.0124377

0.0373131

0.0373131

0.0373131

0.0373131

0.0124377
0.0373131

0.0248754

0.0124377
0.0248754

0.0124377

0.00572051658

0.00572051658

0.001127418
0.000187903

0.000939515

0.000187903
0.000939515

0.000187903
0.000751612

0.000187903
0.000563709

0.000375806
0.000187903

0.000187903

0.00076551643
0.00459309858

0.00382758215

0.00382758215
0.00076551643

0.00076551643
0.00306206572

0.000758606
0.002275818

0.000758606
0.001517212

0.000758606

6.91043e-06
2.073129e-05

1.382086e-05
6.91043e-06

6.91043e-06

0

95.7438320757
666.188276002336

0.1339123631
0.0275086792

0.000115904

8.20319e-05
0.0003281276

8.20319e-05
0.0002460957

0.0001640638
8.20319e-05

8.20319e-05

0.0249891896

0.0043204278
0.0169547848

0.0002600289
8.66763e-05

0.0001733526
8.66763e-05

8.66763e-05

0.0001885497
6.28499e-05

6.28499e-05
0.0001256998

6.28499e-05

0.0007127876
0.0021383628

0.0014255752
0.0007127876

7.24984e-05

9.7549e-05

6.97923e-05

5.8869e-05

9.82605e-05

8.33499e-05

5.92973e-05

7.33338e-05

9.98374e-05

0.0009769641
0.0003256547

0.0004761282
0.0002380641

7.99172e-05

7.62766e-05

8.18703e-05

0.0001751812
8.75906e-05

8.75906e-05

8.74549e-05

0.0017392338
0.0052177014

0.0002690854
0.0001345427

6.44471e-05

7.00956e-05

9.65807e-05
0.0001931614

9.65807e-05

0.0006214672
0.0012429344

0.000124566

0.000160828

5.82526e-05

6.61666e-05

7.916e-05

0.000132494

0.0001616662
8.08331e-05

8.08331e-05

0.000232252
0.000116126

0.000116126

0.0008679652
0.0004339826

6.61025e-05

0.00010894

7.59601e-05

0.000116509

6.6471e-05

0.0001842052
0.0003684104

7.57132e-05

0.000108492

0.0001429926
7.14963e-05

7.14963e-05

7.60083e-05

0.0036892869
0.0012297623

0.000534174
0.000267087

9.03633e-05

9.36765e-05

8.30472e-05

0.0006431124
0.0003215562

8.97773e-05

8.87471e-05

7.05051e-05

7.25267e-05

0.0001384776
6.92388e-05

6.92388e-05

0.0006497028
0.0003248514

8.42068e-05

0.00016336

7.72846e-05

0.0002470289
0.0004940578

7.99489e-05

0.00016708

0.0020410621
0.0080344048

0.0019640646
0.0006546882

0.000168894
8.4447e-05

8.4447e-05

0.0001196134
5.98067e-05

5.98067e-05

0.0002399694
0.0004799388

8.82227e-05

6.93975e-05

8.23492e-05

0.0001567504
7.83752e-05

7.83752e-05

0.0001248944
6.24472e-05

6.24472e-05

0.0002592854
0.0001296427

5.89702e-05

7.06725e-05

5.97076e-05
0.0001791228

0.0001194152
5.97076e-05

5.97076e-05

6.49218e-05

0.0016896858
0.0005632286

0.0011264572
0.0005632286

7.35031e-05

6.28515e-05

6.9145e-05

0.000357729

0.000204636
0.000613908

0.000409272
0.000204636

8.3234e-05

0.000121402

0.0014816397
0.0004938799

5.5693e-05
0.000111386

5.5693e-05

0.0001327332
0.0002654664

7.06413e-05

6.20919e-05

0.000300086
0.000150043

0.000150043

0.0001276024
6.38012e-05

6.38012e-05

9.16095e-05
0.000183219

9.16095e-05

0.0002150778
0.001075389

0.0008603112
0.0002150778

0.0002150778
0.0006452334

0.0004301556
0.0002150778

9.07008e-05

0.000124377

0.0372297224
0.0093074306

0.000620463
0.000206821

0.000206821
0.000413642

0.000105228

0.000101593

0.0046234616
0.0138703848

0.0007225094
0.0003612547

0.000101525

0.000122544

6.61877e-05

7.0998e-05

9.9997e-05
0.000199994

9.9997e-05

0.0002562944
0.0001281472

6.33093e-05

6.48379e-05

8.0444e-05
0.000160888

8.0444e-05

0.0079072374
0.0039536187

7.90633e-05

7.07852e-05

6.74564e-05

4.78083e-05

5.58214e-05

8.26936e-05

5.60307e-05

5.61911e-05

7.54709e-05

4.89219e-05

5.61588e-05

6.73738e-05

7.23529e-05

6.24632e-05

5.76656e-05

7.12295e-05

8.03257e-05

5.99294e-05

9.87201e-05

8.99902e-05

7.18283e-05

5.51254e-05

4.76881e-05

0.000104211

6.16129e-05

7.99477e-05

5.30587e-05

7.16944e-05

0.000105781

6.13869e-05

6.75239e-05

5.59987e-05

6.36434e-05

8.75646e-05

6.56622e-05

7.51045e-05

9.35507e-05

4.93314e-05

6.22988e-05

7.85804e-05

6.4562e-05

5.42365e-05

5.44514e-05

6.79339e-05

5.61419e-05

6.88823e-05

5.59556e-05

6.82286e-05

4.8283e-05

8.64447e-05

7.09366e-05

6.48063e-05

0.000126647

6.8277e-05

5.00012e-05

5.98282e-05

6.34851e-05

5.64724e-05

0.0048327561
0.0016109187

0.0001392392
6.96196e-05

6.96196e-05

0.000225554
0.000112777

0.000112777

9.23529e-05
0.0001847058

9.23529e-05

0.001264244
0.002528488

8.90311e-05

0.000153756

8.66005e-05

9.43735e-05

0.000121596

0.000116323

7.89879e-05

7.66143e-05

0.000124215

8.0889e-05

8.88115e-05

6.68046e-05

8.62416e-05

7.19252e-05
0.0001438504

7.19252e-05

0.0048940341
0.0016313447

0.0032626894
0.0016313447

4.7485e-05

4.13422e-05

4.79019e-05

4.96649e-05

5.81872e-05

4.93414e-05

3.75757e-05

4.65989e-05

4.40983e-05

4.94786e-05

4.34724e-05

3.91645e-05

4.42018e-05

4.7485e-05

5.63917e-05

5.23986e-05

4.8779e-05

0.000384099

4.9647e-05

3.84876e-05

4.49641e-05

6.01926e-05

2.36769e-05

4.63246e-05

3.85444e-05

3.81302e-05

4.07182e-05

2.48681e-05

3.81249e-05

0.0005441901
0.0001813967

0.0001813967
0.0003627934

9.45956e-05

8.68011e-05

0.0003375145
0.0010125435

5.84428e-05
0.0001168856

5.84428e-05

0.0005581434
0.0002790717

7.05583e-05

7.10949e-05

6.68196e-05

7.05989e-05

0.0007159734
0.0021479202

0.0001682816
8.41408e-05

8.41408e-05

0.0002859486
0.0001429743

6.79479e-05

7.50264e-05

0.0009777166
0.0004888583

7.59958e-05

7.9177e-05

8.34125e-05

8.83419e-05

8.03726e-05

8.15585e-05

6.50686e-05

0.0023720124
0.0005930031

7.1531e-05
0.000214593

0.000143062
7.1531e-05

7.1531e-05

0.0003098377
0.0009295131

7.04356e-05
0.0001408712

7.04356e-05

0.0004788042
0.0002394021

9.54666e-05

7.94182e-05

6.45173e-05

0.0001455921
0.0004367763

0.0001455921
0.0002911842

7.69373e-05

6.86548e-05

6.60423e-05
0.0001981269

0.0001320846
6.60423e-05

6.60423e-05

0.0006720876
0.0001680219

0.0001680219
0.0005040657

0.0001680219
0.0003360438

9.39193e-05

7.41026e-05

5.64267e-05

0.0372627812
0.0093544656

0.0038811906
0.0012937302

0.0001283126
6.41563e-05

6.41563e-05

7.84556e-05
0.0001569112

7.84556e-05

0.0011511183
0.0023022366

9.21391e-05

9.98756e-05

8.43754e-05

7.21235e-05

7.91792e-05

8.43703e-05

8.6238e-05

8.1623e-05

6.04765e-05

6.97103e-05

6.81648e-05

6.20531e-05

5.22239e-05

8.09085e-05

7.76571e-05

0.0030985532
0.00929565960000001

0.0009597068
0.0004798534

0.000135017

6.77038e-05

6.57728e-05

6.28311e-05

7.82477e-05

7.0281e-05

6.31993e-05
0.0001263986

6.31993e-05

0.004356001
0.0021780005

7.93883e-05

6.25798e-05

0.000435602

0.000103264

7.03578e-05

6.77246e-05

5.11811e-05

8.03234e-05

5.79914e-05

7.62337e-05

8.27242e-05

7.65734e-05

7.31181e-05

7.53513e-05

6.70566e-05

8.92476e-05

6.47046e-05

6.8715e-05

7.98451e-05

6.63055e-05

6.33881e-05

6.14544e-05

7.43618e-05

9.08625e-05

5.96462e-05

0.000597314
0.000298657

6.23451e-05

0.000174496

6.18159e-05

0.000157686
7.8843e-05

7.8843e-05

0.005709669
0.001903223

0.0001170802
0.0002341604

5.62771e-05

6.08031e-05

0.000166681
0.000333362

0.000166681

0.0001365812
6.82906e-05

6.82906e-05

0.0001580314
0.0003160628

7.1453e-05

8.65784e-05

0.0001978981
0.0003957962

6.35371e-05

0.000134361

0.0003870728
0.0001935364

0.000149461

4.40754e-05

0.001867392
0.000933696

0.000933696

0.0001360186
6.80093e-05

6.80093e-05

6.06902e-05
0.0001820706

0.0001213804
6.06902e-05

6.06902e-05

0.000109463
0.000328389

0.000109463
0.000218926

0.000109463

0.0004102932
0.0001367644

0.0001215792
6.07896e-05

6.07896e-05

0.0001519496
7.59748e-05

7.59748e-05

0.0004446604
0.0013339812

0.0004446604
0.0008893208

0.000120628

7.69174e-05

6.25639e-05

8.70781e-05

9.7473e-05

0.0032959581
0.0010986527

0.0010986527
0.0021973054

7.58993e-05

0.000694411

7.16534e-05

7.32027e-05

6.57353e-05

0.000117751

0.0033935637
0.0011311879

0.001826239
0.0009131195

0.000139982

7.62369e-05

0.000360964

7.75575e-05

6.85704e-05

6.51116e-05

5.73705e-05

6.73266e-05

6.14317e-05
0.0001228634

6.14317e-05

0.0001318112
6.59056e-05

6.59056e-05

0.0001814622
9.07311e-05

9.07311e-05

7.75406e-05

8.96751e-05

7.89137e-05

6.41315e-05

0.000101171

0.0013962876
0.0003490719

0.0010472157
0.0003490719

0.0005646164
0.0002823082

8.23076e-05

6.35388e-05

6.41203e-05

7.23415e-05

6.67637e-05
0.0001335274

6.67637e-05

6.37829e-05

8.83988e-05

0.000144568

6.63272e-05

0.000143719

0.000153787

0.0409554776
0.00755957

0.000116762

0.000134038

0.0005136044
0.0017122302

7.96428e-05
3.98214e-05

3.98214e-05

0.000103479

0.000549792
0.000137448

0.000412344
0.000137448

0.000137448
0.000274896

0.000137448

0.000465712
0.000232856

0.000232856

6.44564e-05

0.0021003215
0.0095547375

0.0069704
0.0019793175

0.0049910825
0.0019793175

0.000386429
0.000772858

0.000133782

0.00012931

0.000123337

0.000159045

0.00019866

0.000122261

0.000157008

0.000156807

0.000152627
0.000305254

0.000152627

0.000380393
0.0001901965

9.12977e-05

9.88988e-05

0.000187127
0.000374254

0.000187127

0.000232136
0.000116068

0.000116068

0.000153089

0.000121004
0.000484016

0.000121004
0.000363012

0.000121004
0.000242008

0.000121004

5.43183e-05

7.31127e-05

0.0001633451
0.0008167255

0.0006533804
0.0001633451

0.0001633451
0.0004900353

0.0001633451
0.0003266902

4.65475e-05

4.6074e-05

7.07236e-05

9.5226e-05

0.0042443856
0.020774301

0.0163807064
0.0040951766

0.002570377
0.007711131

0.002570377
0.005140754

0.000253821

0.000155296

0.00016199

0.00016301

0.00183626

0.0002104352
0.0006313056

0.0002104352
0.0004208704

9.19952e-05

0.00011844

0.0012196186
0.0036588558

0.0007544066
0.0015088132

0.00012086

0.000109158

0.000225597

0.000219235

7.95566e-05

0.000602092
0.000301046

0.000301046

0.000328332
0.000164166

0.000164166

0.0002842374
9.47458e-05

9.47458e-05
0.0001894916

9.47458e-05

0.000149209

0.000336549

0.0017284904
0.0098751064

0.008022657
0.0016045314

0.0064181256
0.0016045314

0.0016045314
0.0048135942

0.0004060971
0.0008121942

8.9913e-05

9.36124e-05

7.0008e-05

8.23802e-05

7.01835e-05

0.0015107082
0.0007553541

0.000136631

8.46283e-05

8.66329e-05

8.56746e-05

0.00010202

9.0671e-05

8.03309e-05

8.87654e-05

0.0003432508
0.0006865016

0.000129709

0.000129709

6.38012e-05

6.38012e-05

8.45381e-05

6.52025e-05

6.52025e-05

0.0001996588
9.98294e-05

9.98294e-05

0.000123959

0.010183218
0.001697203

0.001697203
0.008486015

0.001697203
0.006788812

0.000316492
0.000949476

0.000632984
0.000316492

0.000146569

0.000169923

0.001380711
0.004142133

0.002401344
0.001200672

0.000972313

0.000228359

0.000180039
0.000360078

0.000180039

0.0004740618
0.0028443708

0.0004740618
0.002370309

0.0004740618
0.0018962472

0.0004740618
0.0014221854

0.000517908
0.000258954

0.000258954

0.0004302156
0.0002151078

9.33598e-05

0.000121748

0.000156409

0.004296852
0.000716142

0.00358071
0.000716142

0.002864568
0.000716142

0.002148426
0.000716142

0.0002056973
0.0004113946

5.65263e-05

7.35652e-05

7.56058e-05

0.0007777036
0.0003888518

8.27228e-05

6.81791e-05

6.91599e-05

9.65462e-05

7.22438e-05

0.0001252882
6.26441e-05

6.26441e-05

5.89488e-05
0.0001178976

5.89488e-05

0.0647318252148
0.0111273330358

0.00052351
0.000261755

0.000143003

0.000118752

0.000173717

0.00013801

0.0105538510358
0.052769255179

0.02157822412
0.00539455603

0.01618366809
0.00539455603

0.0005629134
0.0011258268

4.48458e-05

4.74216e-05

0.000391102

7.9544e-05

0.00483164263
0.00966328526

3.75853e-05

4.37258e-05

0.000132285

0.000109083

8.37189e-05

0.0004521379

0.000233354

0.000102959

5.34767e-05

6.23482e-05

4.56696e-05

1.14438e-05

3.16276e-05

4.622e-05

5.26532e-05

9.80739e-06

8.31824e-06

0.000116965

5.2223e-05

6.41298e-05

3.8392e-05

1.03914e-05

8.40283e-05

9.03463e-05

3.5524e-05

2.95155e-05

5.36855e-05

4.09357e-05

3.85487e-05

3.38707e-05

5.05761e-05

5.37894e-05

3.81762e-05

4.9565e-05

2.66081e-05

1.21324e-05

6.90795e-05

0.000106964

5.55909e-05

5.08096e-05

0.000279961

3.04412e-05

3.93929e-05

8.18566e-05

3.88086e-05

0.000285913

5.7643e-05

4.18517e-05

4.39882e-05

6.22218e-05

0.000108286

3.56391e-05

3.38405e-05

3.69418e-05

4.67704e-05

2.89184e-05

4.34169e-05

8.71462e-05

7.58522e-05

0.000119836

4.19949e-05

6.57069e-05

4.05067e-05

3.29689e-05

3.62044e-05

5.00709e-05

1.83115e-05

0.000113282

3.34319e-05

3.02762e-05

3.52307e-05

4.33553e-05

4.15987e-05

5.11736e-05

4.23915e-05

0.000101064

3.85931e-05

7.53805e-05

3.20055e-05

4.58308e-05

1.74163e-05

0.0098409252232
0.0024602313058

0.0024602313058
0.0073806939174

0.0017611836
0.0035223672

0.000266361

0.000204814

0.000150487

0.000173327

0.000241388

0.000143741

5.67276e-05

0.000140728

0.000132987

0.000250623

0.0013980954116
0.0006990477058

7.668691e-05

2.92981e-05

3.80722e-05

9.31661e-06

4.46689e-05

0.0002796667244

1.10753e-07

5.33182e-05

3.69612e-05

1.23714e-08

5.24797e-05

5.85519e-05

2.26226e-05

5.561e-05

4.42132e-05

4.42132e-05

1.87105e-05

1.87105e-05

2.20972e-05

2.20972e-05

4.09946e-05

4.09946e-05

1.23714e-08

0.0001719973

3.41617e-05

3.84668e-05

4.09775e-05

2.87818e-05

2.96095e-05

0.0107962548
0.0026990637

0.0047952645
0.0015984215

0.003196843
0.0015984215

5.05047e-05

5.64841e-05

6.44303e-05

4.42708e-05

5.8255e-05

6.05925e-05

8.55894e-05

6.32424e-05

0.000107875

6.42663e-05

6.41231e-05

6.15912e-05

5.55164e-05

5.42111e-05

5.85502e-05

8.42784e-05

5.70937e-05

5.88702e-05

9.03166e-05

4.58214e-05

5.55602e-05

4.7963e-05

6.97525e-05

6.43722e-05

7.48908e-05

0.0011006422
0.0033019266

0.0007027143
0.0014054286

5.80837e-05

0.000109914

5.79284e-05

8.35137e-05

6.63398e-05

5.55114e-05

5.49848e-05

5.46468e-05

4.53031e-05

7.30989e-05

4.33897e-05

0.0003979279
0.0007958558

4.7729e-05

9.57927e-05

4.62863e-05

9.96753e-05

4.47819e-05

6.36627e-05

9.06163e-08

4.97077e-07

0.000181187
0.000724748

0.000543561
0.000181187

0.000362374
0.000181187

0.000181187

155.322976269954
25.9012293426

0.000163778

8.42026554744999

1.65344955659
8.26511385955

0.001101244
0.000550622

0.000138487

0.00014719

0.000149584

0.000115361

1.75318e-05

0.000323181
0.000646362

0.000156415

0.000166766

1.65158969829
6.60602507116

0.01766384129
0.05299152387

0.03334700258
0.01667350129

9.49693e-05

8.66563e-05

0.000118952

0.000199711

4.63839e-06

0.000238026

7.7273e-05

0.000292749

0.000105553

9.87642e-05

0.000103857

0.00132833

0.000105837

0.000104926

0.000219125

0.000106795

0.000110053

0.000901142

0.000223548

0.000195753

0.000179164

9.19683e-05

0.000132873

0.00010757

0.000222108

9.71771e-05

0.000119708

0.000118326

0.000160282

8.52501e-05

0.00011428

0.000124257

0.000125248

0.000112624

0.00010888

0.00017257

0.000107902

9.48546e-05

0.000241634

0.000103368

0.000167146

0.000121253

0.000158973

0.000114095

0.00032617

0.000112111

0.000462143

0.000118493

0.00416058

0.00015836

0.00249112

0.000398498

0.000112058

0.000198214

9.5397e-05

0.000142188

0.001236126
0.000618063

0.000184123

0.000132134

0.000149118

0.000152688

0.000167856
0.000335712

0.000167856

0.000204421
0.000408842

0.000204421

0.000153316
0.000306632

0.000153316

4.900776405
1.633592135

0.005853696
0.002926848

0.000235292

0.000239454

0.000414148

0.000233706

0.000238554

0.000254919

0.000254434

0.000271337

0.000238526

0.000277585

0.000268893

1.626164303
3.252328606

0.000166069

0.00050454

0.000231161

0.000265849

0.000160287

0.000140719

0.000231797

0.000160402

0.000234484

0.000300363

0.000253651

0.000243821

0.000234235

0.000167265

0.000152008

0.000151271

0.000240167

0.000231386

0.000206786

0.000256075

0.000254236

0.00600604

0.000274112

0.00016136

0.000260628

0.000243123

0.000211037

0.000154395

0.00851312

0.000146814

0.00016832

0.000150589

0.000252288

0.00023806

0.0034484

0.00133864

0.000765867

0.000231939

0.000155209

0.000221613

0.000206595

0.000199898

0.000256872

0.000141936

0.0001646

0.00017569

0.000218005

0.000174821

0.000145547

0.000719966

0.000157695

0.000155801

0.000235255

0.000150208

0.000256717

0.000159873

0.12837

0.000169911

0.000454071

0.000239421

0.00023157

0.000175412

0.000239256

0.000249627

0.000227377

0.000156395

0.000186641

0.000226047

0.00037165

0.00015747

0.00677084

0.000550434

0.013842

0.000371402

0.00151598

0.00475267

0.000243252

0.000211485

0.000258466

0.000336068

0.0533087

0.000149083

0.0002174

0.000230066

0.000171895

0.000159727

0.000234465

0.000252931

0.00014422

0.000164001

0.000117712

0.0002312

0.000336915

0.00163391

0.000351464

0.000362184

0.000230267

0.139711

0.00023469

0.000453503

0.000238361

0.000168154

0.000152569

0.000238402

0.000214801

0.000247486

0.000260098

0.00207991

0.000157133

0.000234926

0.000245279

0.000255704

0.000242755

0.000275712

0.00115511

0.0001524

0.000217951

0.000251874

0.000119825

0.000237397

0.000159357

0.000135592

0.000156782

0.000247742

0.000168972

0.000161888

1.22357

0.0037394

0.000290669

0.000137671

0.004500984
0.009001968

0.000493163

0.000290661

0.000175948

0.000220724

0.000220139

0.000109708

0.00013184

0.000362944

0.000111918

0.000240094

0.000158809

0.000129463

0.000123076

0.000260978

0.000116266

0.000124446

0.000136665

0.000157372

0.000150416

0.000372102

0.000159449

0.000140346

0.000114457

0.000180406
0.000360812

0.000180406

0.0009685235
0.003874094

0.0009685235
0.0029055705

0.000253692
0.000126846

0.000126846

0.0006924075
0.001384815

9.60492e-05

0.000113782

9.40788e-05

9.6598e-05

9.64801e-05

0.000102594

9.28254e-05

0.00014927
0.00029854

0.00014927

0.0313785545
0.1551516879

0.0359582104
0.0089895526

0.0006570266
0.0019710798

9.01697e-05
0.0001803394

9.01697e-05

0.0011337138
0.0005668569

9.53965e-05

9.27595e-05

9.83219e-05

0.000139957

0.000140422

0.0015373137
0.0005124379

0.0010248758
0.0005124379

0.000106563

0.000148693

7.75147e-05

0.000105752

7.39152e-05

0.0004443702
0.0013331106

0.0001451442
7.25721e-05

7.25721e-05

7.67447e-05
0.0001534894

7.67447e-05

0.0002873592
0.0001436796

8.09109e-05

6.27687e-05

6.18931e-05
0.0001237862

6.18931e-05

0.0001789614
8.94807e-05

8.94807e-05

0.0073757179
0.0221271537

0.00072493
0.000362465

0.000126885

0.00023558

0.0111871938
0.0055935969

9.77536e-05

0.000102544

0.000103321

8.65009e-05

0.000105935

0.000115153

7.94317e-05

9.80997e-05

0.000179276

0.000207081

7.91717e-05

0.000101818

0.000126414

0.000109518

9.47975e-05

0.000107962

9.06013e-05

0.000115013

0.000170139

0.00011346

0.00012687

8.45267e-05

0.000154117

0.000172586

0.000159974

6.81571e-05

8.58985e-05

9.08726e-05

0.000114674

0.000115205

0.000438392

0.000106546

9.13746e-05

0.00011646

0.000316151

0.000115845

0.000186159

9.31751e-05

0.000274315

0.000104672

0.000106701

0.000106351

8.05839e-05

0.000776184
0.000388092

0.000109767

0.000163087

0.000115238

0.000117666
0.000235332

0.000117666

0.000353896
0.000176948

0.000176948

8.1195e-05
0.00016239

8.1195e-05

0.000655755
0.00131151

0.000120519

0.000191489

0.000120414

0.000122528

0.000100805

0.000645741
0.002582964

0.000645741
0.001937223

0.0007254652
0.0003627326

9.63065e-05

8.7968e-05

9.35773e-05

8.48808e-05

0.0005660168
0.0002830084

0.000105556

9.16991e-05

8.57533e-05

3.18828e-05

0.0045654245
0.018261698

0.0088452015
0.0029484005

0.000234382
0.000117191

0.000117191

0.000150161
0.000300322

0.000150161

0.000119913
0.000239826

0.000119913

0.00021109
0.00042218

0.000104856

0.000106234

0.000588846
0.000294423

0.000117317

9.06312e-05

8.64748e-05

0.0002924921
0.0005849842

9.52731e-05

0.000197219

0.0003927574
0.0001963787

0.000105532

9.08467e-05

0.00024158
0.00012079

0.00012079

9.95775e-05
0.000199155

9.95775e-05

0.000111445
0.00022289

0.000111445

0.000901592
0.000450796

9.7101e-05

0.000353695

0.000537414
0.000268707

0.000103938

0.000164769

0.0005912942
0.0002956471

8.54921e-05

0.000115235

9.492e-05

0.000129419
0.000258838

0.000129419

0.0001807402
9.03701e-05

9.03701e-05

0.001617024
0.004851072

0.000259308
0.000129654

0.000129654

0.000168509
0.000337018

0.000168509

0.000212888
0.000106444

0.000106444

0.000125695
0.00025139

0.000125695

0.00022115
0.000110575

0.000110575

0.000310854
0.000155427

0.000155427

0.00164144
0.00082072

0.000215236

0.000227905

0.00014119

0.000112174

0.000124215

5.18616e-05

2.23174e-05

3.78772e-05
7.57544e-05

3.78772e-05

0.005716395
0.02286558

0.003885961
0.011657883

0.000687554
0.000343777

0.00022008

0.000123697

0.006300836
0.003150418

0.000132053

0.000813357

0.000132496

0.000157117

0.000130615

0.000136242

0.000145368

0.000161364

0.000128154

0.000136305

0.000124991

0.00015032

0.00023137

0.000165996

0.000128123

0.000113109

0.000163438

0.000249152
0.000498304

0.0001268

0.000122352

0.000285228
0.000142614

0.000142614

0.005491302
0.001830434

0.000279336
0.000139668

0.000139668

0.000267178
0.000133589

0.000133589

0.00138328
0.00069164

0.000134652

0.000133124

0.000159999

0.000105406

0.000158459

0.001731074
0.000865537

0.000154718

0.000137636

0.000126426

0.000156472

0.000130152

0.000160133

0.000116826

0.037970103
0.009593886

0.0085323303

0.0025538921
0.0076616763

0.00370392
0.00185196

0.00185196

0.000660917
0.001321834

0.000239828

0.000421089

4.10151e-05
8.20302e-05

4.10151e-05

0.000870654
0.000290218

0.000290218
0.000580436

0.000290218

0.0037412217

0.0009175347
0.0003058449

0.0006116898
0.0003058449

3.66129e-05

0.000269232

0.000941229
0.002823687

0.000379833
0.000759666

0.000379833

0.000561396
0.001122792

0.000561396

0.000405441
0.000810882

0.000405441

0.015291783

0.004824327
0.001608109

0.002816642
0.001408321

0.000448621

0.000126917

0.000290493

0.000174427

0.000183853

0.00018401

0.000199788
0.000399576

0.000199788

0.006903984
0.002301328

0.0011981
0.00059905

0.000196584

0.000193471

0.000208995

0.000194034
0.000388068

0.000194034

0.001088304
0.000544152

0.000365971

0.000178181

0.000176353
0.000352706

0.000176353

0.000519432
0.000259716

0.000259716

0.001056046
0.000528023

0.000369382

0.000158641

0.001023983
0.003071949

0.001023983
0.002047966

0.00036145

0.000189219

0.000241674

0.00023164

0.000163841
0.000491523

0.000327682
0.000163841

0.000163841

0.000291672
9.7224e-05

9.7224e-05
0.000194448

9.7224e-05

0.000109666

0.0009542665
0.003817066

0.0012618972
0.0004206324

0.00025844
0.00051688

0.000122258

0.000136182

0.0001522862
7.61431e-05

7.61431e-05

8.60493e-05
0.0001720986

8.60493e-05

0.0004498219
0.0013494657

7.95631e-05
0.0001591262

7.95631e-05

0.0003780176
0.0001890088

0.000127412

6.15968e-05

0.00018125
0.0003625

9.9474e-05

8.1776e-05

0.0002514366
8.38122e-05

0.0001676244
8.38122e-05

8.38122e-05

0.001452528
0.000363132

0.000363132
0.001089396

0.000112125
0.00022425

0.000112125

0.000127256
0.000254512

0.000127256

0.000247502
0.000123751

0.000123751

8.25019e-05
0.0001650038

8.25019e-05

0.00181939
0.000363878

0.001455512
0.000363878

0.001091634
0.000363878

0.000727756
0.000363878

0.000195719

0.000168159

0.588303635867
2.93074436747321

8.27182e-05

0.0447647713117
0.1777478614468

5.83813e-05

0.1268745765351
0.0426476425117

0.000960594
0.000480297

0.000100349

0.000110504

0.000165269

0.000104175

0.0001507037
0.0003014074

5.73623e-05

9.33414e-05

9.14477e-05

4.05025e-05

0.0001988993
0.0003977986

5.38384e-05

4.05473e-05

4.25125e-05

6.20011e-05

9.65877e-05
0.0001931754

9.65877e-05

9.07972e-05
4.53986e-05

4.53986e-05

0.0002005752
0.0001002876

5.38139e-05

4.64737e-05

4.65268e-05
9.30536e-05

4.65268e-05

5.71808e-05

0.00296069
0.001480345

4.93426e-05

5.99105e-05

4.741e-05

5.45583e-05

4.69195e-05

5.08703e-05

6.1623e-05

7.56837e-05

0.000114601

0.000126427

3.90162e-05

0.000145821

9.4165e-05

5.60102e-05

5.95679e-05

4.18677e-05

7.1107e-05

4.7673e-05

4.0484e-05

4.7455e-05

7.17467e-05

7.80854e-05

0.0007932908
0.0003966454

3.80315e-05

0.000109813

4.06227e-05

4.28799e-05

3.95488e-05

5.42572e-05

7.14923e-05

0.0029684014
0.0014842007

5.25684e-05

6.83777e-05

6.08804e-05

6.26382e-05

0.00106725

0.000172486

0.0001367812
6.83906e-05

6.83906e-05

0.0001690265
0.000338053

0.000119159

4.98675e-05

0.0013926514
0.0027853028

7.37519e-05

0.00073565

3.77828e-05

6.16099e-05

0.000100893

4.76263e-05

0.000241444

4.25717e-05

5.13218e-05

0.00295394312
0.00147697156

4.85444e-05

6.4917e-05

8.58051e-05

5.19658e-05

1.68266e-06

4.69731e-05

4.82911e-05

3.87387e-05

5.49216e-05

6.72218e-05

4.45541e-05

5.50325e-05

0.000371563

9.47309e-05

5.09651e-05

5.39681e-05

5.09722e-05

5.15433e-05

5.73629e-05

4.3189e-05

5.05978e-05

4.34314e-05

0.0001620006
8.10003e-05

8.10003e-05

0.00010351
5.1755e-05

5.1755e-05

4.54e-05

0.0001860996
9.30498e-05

9.30498e-05

8.36881e-05
0.0001673762

8.36881e-05

0.0001903132
9.51566e-05

9.51566e-05

0.000316934
0.000158467

0.000158467

0.0002950218
0.0001475109

8.99439e-05

5.7567e-05

0.0037781132517
0.0075562265034

6.28867e-05

0.000211653

7.44523e-05

8.10659e-05

7.05888e-05

0.000133502

8.72166e-05

4.66083e-05

9.62761e-05

0.000107177

0.000430616

7.78482e-05

0.000163126

5.91671e-05

6.47633e-05

8.0793e-05

4.87501e-05

8.03783e-05

1.83043e-07

2.21137e-08

8.37026e-05

0.000104224

7.60157e-05

9.60912e-05

8.27278e-05

0.000294143

6.39451e-05

7.0684e-05

7.18748e-05

6.61841e-05

7.68144e-05

7.708e-05

0.000117935

9.34942e-05

8.3007e-05

0.000169614

5.69788e-05

6.42405e-05

1.48995e-07

5.21342e-05

4.22085e-05

8.72487e-05
0.0001744974

4.34228e-05

4.38259e-05

4.95157e-05
9.90314e-05

4.95157e-05

0.0002689398
0.0001344699

8.18619e-05

5.2608e-05

0.0001054587
0.0002109174

4.43536e-05

6.11051e-05

6.49782e-05
0.0001299564

6.49782e-05

0.0001173986
5.86993e-05

5.86993e-05

0.000185011
9.25055e-05

4.54911e-05

4.70144e-05

6.78637e-05
0.0001357274

6.78637e-05

0.0001147854
5.73927e-05

5.73927e-05

0.0004116774
0.0002058387

8.38417e-05

4.63314e-05

7.56656e-05

0.000450052
0.000900104

0.000100681

5.23537e-05

6.07915e-05

0.000115785

4.96401e-05

7.08007e-05

9.28854e-05
0.0001857708

9.28854e-05

0.0004118048
0.0002059024

4.91617e-05

0.000107003

4.97377e-05

0.000198885
9.94425e-05

5.19997e-05

4.74428e-05

0.0002856572
0.0001428286

4.5345e-05

5.6642e-05

4.08416e-05

0.0005291014
0.0002645507

0.000110332

9.51824e-05

5.90363e-05

4.34807e-05

0.0008194362
0.0004097181

6.38962e-05

0.000295827

4.99949e-05

0.0007075705
0.001415141

4.83671e-05

4.57603e-05

5.75982e-05

0.000107286

0.000135592

6.35329e-05

0.000125018

0.000124416

0.0001881888
0.0003763776

4.96001e-05

7.17156e-05

6.68731e-05

9.52875e-05
0.000190575

4.91836e-05

4.61039e-05

0.000689803
0.0003449015

7.61409e-05

6.71546e-05

0.000201606

0.003609086
0.001804543

0.000607461

5.73787e-05

6.57996e-05

5.14137e-05

0.000113766

8.44805e-05

8.66389e-05

6.77047e-05

6.866e-05

0.000432171

5.54181e-05

6.37097e-05

4.99411e-05

5.45465e-05
0.000109093

5.45465e-05

0.0020858871
0.0041717742

9.11796e-05

0.000371338

0.000119838

9.98979e-05

5.43737e-05

6.73332e-05

5.63685e-05

9.65576e-05

8.28072e-05

0.000207931

7.39731e-05

0.000180376

5.56885e-05

8.28003e-05

0.000110942

8.45274e-05

5.80705e-05

0.000114396

7.74886e-05

0.0004087976
0.0002043988

5.05695e-05

4.71497e-05

5.14987e-05

5.51809e-05

0.000309357
0.0001546785

6.1228e-05

9.34505e-05

6.815e-05

0.00011834
5.917e-05

5.917e-05

5.3822e-05

7.14999e-05
0.0001429998

7.14999e-05

0.000154579
7.72895e-05

7.72895e-05

5.01188e-05
0.0001002376

5.01188e-05

0.0002468902
0.0001234451

4.04073e-05

8.30378e-05

4.23305e-05
8.4661e-05

4.23305e-05

0.0009988264
0.0004994132

4.07398e-05

3.88904e-05

4.79448e-05

5.66322e-05

7.83433e-05

5.24356e-05

4.56003e-05

5.10817e-05

4.74632e-05

4.02819e-05

9.16934e-05
4.58467e-05

4.58467e-05

0.000255087

0.0002181053
0.0004362106

6.4928e-05

9.87027e-05

5.44746e-05

0.0006899936
0.0003449968

4.11222e-05

6.45406e-05

0.000108673

6.64816e-05

6.41794e-05

0.0001881868
0.0003763736

6.29938e-05

0.000125193

4.46135e-05
8.9227e-05

4.46135e-05

8.87186e-05
4.43593e-05

4.43593e-05

0.0012666636
0.0006333318

0.000164264

0.000184192

7.44862e-05

0.000116031

9.43586e-05

0.000632572
0.000316286

0.00011102

0.000205266

0.0015922612
0.0031845224

5.28777e-05

5.18451e-05

5.45948e-05

4.01125e-05

5.58025e-05

0.000228356

4.3348e-05

4.32814e-05

6.16874e-05

4.59152e-05

4.97417e-05

8.36467e-05

0.000678064

4.42767e-05

5.87115e-05

8.26798e-05
4.13399e-05

4.13399e-05

6.59247e-05

0.0002020786
0.0001010393

4.95363e-05

5.1503e-05

0.0003105202
0.0001552601

9.04768e-05

6.47833e-05

3.79445e-05
7.5889e-05

3.79445e-05

4.89128e-05
9.78256e-05

4.89128e-05

4.05454e-05
8.10908e-05

4.05454e-05

0.0009463536
0.0004731768

0.000149082

6.48477e-05

0.000195161

6.40861e-05

7.63336e-05
0.0001526672

7.63336e-05

8.15192e-05
4.07596e-05

4.07596e-05

0.000509225
0.0002546125

4.57715e-05

4.03865e-05

3.88401e-05

5.12833e-05

7.83311e-05

6.48506e-05

0.0002641685
0.000528337

0.000144096

5.87729e-05

6.12996e-05

0.0011742652
0.0005871326

4.88609e-05

0.000104757

5.97068e-05

6.25334e-05

7.89953e-05

7.69977e-05

6.70421e-05

8.82394e-05

4.35053e-05

0.000478874
0.000239437

0.00012186

0.000117577

6.08619e-05
0.0001217238

6.08619e-05

0.0001504649
0.0003009298

4.41108e-05

6.11743e-05

4.51798e-05

0.0007317148
0.0014634296

6.6051e-05

0.000232142

6.90175e-05

0.000121393

5.68483e-05

6.8e-05

5.90898e-05

5.91732e-05

0.0013850662
0.0027701324

4.01462e-05

4.21302e-05

7.42196e-05

7.52658e-05

5.67541e-05

3.77606e-05

4.75507e-05

4.05127e-05

5.82679e-05

2.76309e-05

5.09096e-05

4.18866e-05

5.27419e-05

3.81286e-05

3.93988e-05

5.07932e-05

0.000103342

2.59314e-05

4.90408e-05

3.59497e-05

4.60522e-05

5.42386e-05

4.107e-05

3.82454e-05

4.29266e-05

8.5453e-05

4.74322e-05

4.12869e-05

0.0008161633
0.0016323266

6.74719e-05

0.000119999

5.5668e-05

7.40311e-05

0.000408354

9.06393e-05

4.69408e-05

0.004881504
0.002440752

3.71845e-05

8.1079e-05

4.26067e-05

0.00136488

0.000118253

0.000105645

4.70067e-05

0.000329042

5.00275e-05

5.21386e-05

0.000212889

0.0001297025
0.000259405

6.62874e-05

6.34151e-05

0.000124192
0.000248384

0.000124192

0.0013333094
0.0006666547

5.62256e-05

0.000141132

7.09022e-05

0.000100376

0.000119224

4.92666e-05

6.43972e-05

6.51311e-05

0.0006881248
0.0003440624

5.53592e-05

4.53391e-05

5.00812e-05

4.90856e-05

7.62947e-05

6.79026e-05

0.0001243418
6.21709e-05

6.21709e-05

0.0001084449
0.0002168898

6.29541e-05

4.54908e-05

0.000128966
0.000257932

0.000128966

0.000216634
0.000108317

4.95223e-05

5.87947e-05

0.000136976
0.000273952

0.000136976

6.61565e-05
0.000132313

6.61565e-05

0.000411511
0.0002057555

6.05516e-05

4.71529e-05

4.21588e-05

5.58922e-05

7.43181e-05
0.0001486362

7.43181e-05

0.000178554
8.9277e-05

8.9277e-05

8.26416e-05
0.0001652832

8.26416e-05

0.0024243262
0.0012121631

5.85516e-05

5.60258e-05

4.34518e-05

6.16774e-05

5.39352e-05

5.39744e-05

5.52644e-05

7.48749e-05

4.23275e-05

5.38208e-05

5.99343e-05

6.15282e-05

4.73774e-05

4.90125e-05

7.47522e-05

4.45508e-05

5.39337e-05

4.74543e-05

4.76833e-05

6.60089e-05

4.53806e-05

6.06431e-05

5.96749e-05

0.0001679408
8.39704e-05

8.39704e-05

5.05803e-05
0.0001011606

5.05803e-05

0.0001004842
5.02421e-05

5.02421e-05

0.0003394288
0.0001697144

5.83524e-05

0.000111362

6.56263e-05
0.0001312526

6.56263e-05

0.0003825573
0.0007651146

7.36829e-05

6.76428e-05

7.5739e-05

4.90401e-05

6.6496e-05

4.99565e-05

4.72017e-05
9.44034e-05

4.72017e-05

9.69002e-05
4.84501e-05

4.84501e-05

0.0001431448
7.15724e-05

7.15724e-05

6.2462e-05
0.000124924

6.2462e-05

0.0004480292
0.0002240146

7.08831e-05

4.72482e-05

6.39943e-05

4.1889e-05

0.000110633
5.53165e-05

5.53165e-05

0.000260314
0.000520628

0.000129861

0.000130453

4.86361e-05
9.72722e-05

4.86361e-05

0.0001947022
9.73511e-05

4.6298e-05

5.10531e-05

3.82922e-05
7.65844e-05

3.82922e-05

0.0001588254
0.0003176508

7.73847e-05

8.14407e-05

8.57096e-05
4.28548e-05

4.28548e-05

0.000268684
0.000134342

0.000134342

0.000845909
0.0004229545

0.000208086

4.41602e-05

8.71546e-05

8.35537e-05

4.47724e-05
8.95448e-05

4.47724e-05

8.10462e-05
0.0001620924

8.10462e-05

6.92707e-05
0.0001385414

6.92707e-05

0.001357949
0.002715898

9.26566e-05

8.46879e-05

6.04412e-05

6.61174e-05

8.4476e-05

7.81935e-05

5.41922e-05

5.50876e-05

5.87315e-05

6.73367e-05

7.02412e-05

6.67651e-05

5.63005e-05

5.93777e-05

6.94853e-05

6.12112e-05

6.4882e-05

7.53516e-05

5.75789e-05

7.48349e-05

0.000339081
0.000678162

0.000110765

0.000228316

8.08744e-05
4.04372e-05

4.04372e-05

4.66644e-05

4.03227e-05
8.06454e-05

4.03227e-05

0.000186632
9.3316e-05

9.3316e-05

0.0001177787
0.0002355574

4.63031e-05

7.14756e-05

4.35111e-05

8.40257e-05
0.0001680514

8.40257e-05

0.0001023002
5.11501e-05

5.11501e-05

0.0002811097
0.0005622194

6.53729e-05

4.91216e-05

4.52008e-05

4.39684e-05

7.7446e-05

0.0003219898
0.0006439796

4.77055e-05

3.76321e-05

4.56093e-05

4.95261e-05

5.52427e-05

3.93378e-05

4.69363e-05

0.0059870772
0.0019956924

0.0001457042
7.28521e-05

7.28521e-05

0.0002253253
0.0004506506

6.055e-05

5.00575e-05

5.99587e-05

5.47591e-05

0.0018541062
0.0009270531

6.4405e-05

5.24584e-05

9.47387e-05

5.61179e-05

5.05743e-05

5.82801e-05

5.90531e-05

7.72494e-05

5.96084e-05

6.1134e-05

5.92302e-05

6.73348e-05

5.7057e-05

5.47573e-05

5.50545e-05

0.0002467224
0.0004934448

6.38411e-05

5.44979e-05

6.75992e-05

6.07842e-05

0.0001835449
0.0003670898

6.60429e-05

6.129e-05

5.6212e-05

0.0001093792
5.46896e-05

5.46896e-05

0.000126424
6.3212e-05

6.3212e-05

0.0001451577
0.0002903154

6.99665e-05

7.51912e-05

0.0001542706
7.71353e-05

7.71353e-05

6.30551e-05

8.31795e-05
0.000166359

8.31795e-05

4.77386e-05

7.21127e-05

0.0003025735
0.001210294

0.0003025735
0.0009077205

0.000605147
0.0003025735

0.000103341

7.47935e-05

0.000124439

0.418225666324
0.104614353256

0.075488098768
0.025194226756

9.45815e-05

0.0005896182
0.0002948091

5.1959e-05

6.86071e-05

5.19463e-05

7.67508e-05

4.55459e-05

0.000382668
0.000191334

0.000191334

0.000140432
0.000280864

0.000140432

0.0002755396
0.0001377698

5.26868e-05

8.5083e-05

0.008660320516
0.004330160258

4.58182e-05

4.64053e-05

0.000126276

7.67062e-05

6.41927e-05

7.04704e-05

0.000146029

5.45345e-05

5.11573e-05

5.93065e-05

5.43622e-05

5.18462e-05

5.87618e-05

7.43076e-05

6.86892e-05

8.37952e-05

4.48453e-05

4.50493e-05

5.22396e-05

6.36258e-05

6.30807e-05

4.75928e-05

5.38562e-05

5.45499e-05

4.92597e-05

0.00066611

4.41438e-05

2.95158e-07

5.39607e-05

5.62471e-05

6.0974e-05

4.52657e-05

5.38897e-05

4.98795e-05

6.08223e-05

4.22611e-05

9.66168e-05

0.000121077

7.35494e-05

5.67561e-05

0.000121289

6.955e-05

0.000166826

5.83813e-05

8.3477e-05

8.17158e-05

4.99532e-05

7.03455e-05

5.25535e-05

5.06362e-05

0.000304814

6.65081e-05

6.55047e-05

0.0002115176
0.0004230352

6.21346e-05

5.44976e-05

9.48854e-05

0.00650509
0.003252545

5.01181e-05

0.000144604

6.76557e-05

4.27242e-05

4.65968e-05

5.11384e-05

4.27334e-05

7.85559e-05

0.000122997

7.53567e-05

5.14853e-05

7.90635e-05

7.35582e-05

4.38914e-05

5.746e-05

4.62567e-05

4.34375e-05

6.66786e-05

7.22636e-05

7.54743e-05

5.28793e-05

5.81165e-05

6.53454e-05

6.24375e-05

5.00954e-05

4.54789e-05

7.60079e-05

5.35588e-05

6.2375e-05

6.66541e-05

0.000134301

0.00010743

0.000101744

0.000140912

5.98917e-05

5.79386e-05

8.00667e-05

8.7266e-05

6.12381e-05

6.0578e-05

4.79418e-05

5.36723e-05

6.79012e-05

5.60017e-05

5.57211e-05

8.89761e-05

6.59666e-05

0.011388163198
0.022776326396

8.32376e-05

5.9657e-05

6.71085e-05

5.83521e-05

8.55278e-05

5.31824e-05

5.7267e-05

5.56406e-05

0.000166519

7.83217e-05

6.38011e-05

5.74805e-05

6.28341e-05

6.46748e-05

7.66031e-05

0.00011232

6.97285e-05

5.86152e-05

6.14952e-05

6.09319e-05

6.88218e-05

0.00011267

5.4568e-05

4.70435e-05

6.73116e-05

5.27065e-05

5.64821e-05

5.04188e-05

5.99189e-05

9.66284e-05

0.000135009

5.65018e-05

5.95329e-05

7.52954e-05

5.92402e-05

8.96827e-05

6.16199e-05

6.53053e-05

8.28512e-05

7.83952e-05

0.000119008

8.72373e-05

6.13955e-05

7.86624e-05

6.75587e-05

7.234e-05

8.09581e-05

7.73616e-05

9.24854e-05

7.5734e-05

1.74338e-07

5.86888e-05

6.48533e-05

0.00014945

7.17367e-05

9.44814e-05

0.000207597

5.42523e-05

0.000117301

6.536e-05

8.92971e-05

7.1188e-05

6.05949e-05

5.34394e-05

4.95333e-05

7.20006e-05

7.84335e-05

6.14693e-05

5.02868e-05

5.46411e-05

7.81995e-05

4.71912e-05

5.93099e-05

6.35352e-05

5.31848e-05

6.43127e-05

7.80154e-05

5.02927e-05

6.06759e-05

7.65365e-05

7.86562e-05

7.17964e-05

0.000235351

6.31498e-05

6.96413e-05

5.75532e-05

0.000239217

0.000142443

7.28652e-05

5.46411e-05

0.000327995

1.61836e-06

5.75996e-05

8.78351e-05

5.60955e-05

6.8543e-05

7.7894e-05

7.01751e-05

5.26986e-05

5.78811e-05

3.401e-05

6.12638e-05

0.000142697

6.41712e-05

0.000118331

9.28286e-05

5.78253e-05

0.000160119

5.82561e-05

6.53415e-05

8.42227e-05

6.01225e-05

8.3049e-05

6.97679e-05

8.36558e-05

6.05328e-05

5.93922e-05

6.88043e-05

5.5927e-05

6.15518e-05

6.13259e-05

5.50049e-05

0.000100801

6.35087e-05

0.00010072

5.55009e-05

5.34974e-05

8.62018e-05

8.22951e-05

7.04229e-05

5.1373e-05

7.108e-05

6.95617e-05

5.58648e-05

0.00010527

8.70212e-05

7.55471e-05

5.80388e-05

7.32297e-05

7.43058e-05

5.96384e-05

5.30963e-05

5.01356e-05

7.4292e-05

5.93425e-05

0.000115281

5.8894e-05

5.22418e-05

7.70769e-05

0.0016399388
0.0008199694

3.75251e-05

3.84767e-05

3.72444e-05

3.03345e-05

3.1263e-05

2.82577e-05

3.45127e-05

3.66228e-05

3.9399e-05

2.98808e-05

3.20627e-05

3.86395e-05

3.65637e-05

3.76814e-05

3.82993e-05

5.76586e-05

5.4271e-05

4.77215e-05

3.71983e-05

6.2382e-05

3.39747e-05

0.0009235197
0.0018470394

4.59416e-05

5.60572e-05

4.73596e-05

4.38135e-05

5.55769e-05

5.68328e-05

4.61107e-05

0.000476349

4.95033e-05

4.59751e-05

0.0001696258
8.48129e-05

8.48129e-05

0.0066492246
0.0033246123

9.42002e-05

6.44793e-05

7.53801e-05

7.11696e-05

5.39527e-05

8.37848e-05

0.0001022

6.91706e-05

8.73158e-05

8.10529e-05

0.00010396

6.24011e-05

5.23395e-05

5.7211e-05

4.2372e-05

8.23532e-05

4.31555e-05

9.5003e-05

6.01836e-05

6.49103e-05

4.22397e-05

8.94305e-05

4.21822e-05

5.49583e-05

4.23305e-05

7.0079e-05

9.65331e-05

8.82601e-05

0.000203882

6.59834e-05

5.68402e-05

4.18733e-05

7.45813e-05

5.21054e-05

4.88104e-05

7.18698e-05

5.85483e-05

4.22812e-05

5.67492e-05

4.81872e-05

7.34115e-05

5.59921e-05

7.57536e-05

8.68094e-05

5.52893e-05

5.66351e-05

7.9225e-05

4.7176e-05

0.0793748731
0.2380779609

0.0006320934
0.0003160467

6.82219e-05

5.36502e-05

6.41916e-05

5.79068e-05

7.20762e-05

4.66584e-05

0.0016616547
0.0033233094

5.37171e-05

3.9328e-05

5.40664e-05

6.36439e-05

5.4523e-05

5.02696e-05

4.92684e-05

4.66824e-05

4.35097e-05

4.43234e-05

5.5622e-05

4.59769e-05

4.91288e-05

6.96227e-05

5.70168e-05

4.42246e-05

3.68694e-05

4.53237e-05

4.24232e-05

4.89843e-05

3.9568e-05

3.88623e-05

4.52675e-05

4.8141e-05

6.79405e-05

4.20617e-05

6.81685e-05

0.000111987

8.91319e-05

0.000116002

0.0006061466
0.0012122932

4.78969e-05

5.51192e-05

4.80857e-05

4.10958e-05

5.42232e-05

4.63807e-05

5.07295e-05

5.01483e-05

4.33451e-05

5.65122e-05

4.47522e-05

6.78578e-05

0.0767443667
0.1534887334

5.76631e-05

5.14383e-05

6.1258e-05

0.000107965

6.5451e-05

6.40209e-05

7.48595e-05

9.62897e-05

6.64295e-05

4.53103e-05

5.90366e-05

3.69925e-05

0.0757262

3.69413e-05

5.8202e-05

0.000136309

4.52534e-05

0.000158431
7.92155e-05

7.92155e-05

0.0014301544
0.0003575386

0.0003575386
0.0010726158

0.0003575386
0.0007150772

5.85271e-05

0.000160482

6.50685e-05

7.3461e-05

0.00336699618
0.00092534996

0.00232929922
0.00086917646

1.49899e-05

6.3964e-06

4.0225e-05

2.30896e-05

1.85231e-05

2.19851e-05

0.0005909463
0.0011818926

3.33704e-05

4.25478e-05

0.000386303

4.14064e-05

4.49939e-05

4.23248e-05

1.24683e-05

5.0884e-05

1.10245e-05

9.77446e-06

1.79113e-05

5.09585e-05

0.000112347
5.61735e-05

5.61735e-05

0.000229435
0.00091774

0.000229435
0.000688305

0.000238794
0.000119397

0.000119397

0.000220076
0.000110038

0.000110038

0.0017152097
0.0004397337

0.0002922702
9.74234e-05

9.84226e-05
4.92113e-05

4.92113e-05

9.64242e-05
4.82121e-05

4.82121e-05

0.0004351811
0.0001596354

4.37251e-05

0.0001159103
0.0002318206

3.67994e-05

2.3097e-05

2.7528e-05

2.84859e-05

0.0001826749
0.0005480247

0.0002526442
0.0001263221

5.90069e-05

6.73152e-05

0.0001127056
5.63528e-05

5.63528e-05

5.78726e-05

0.000315471
0.001261884

0.000946413
0.000315471

6.05047e-05
0.0001210094

6.05047e-05

0.0002953286
0.0001476643

7.13858e-05

7.62785e-05

0.000214604
0.000107302

0.000107302

9.06657e-05

8.99075e-05

0.000203514
0.000814056

0.000203514
0.000610542

0.0002591
0.00012955

0.00012955

0.000147928
7.3964e-05

7.3964e-05

5.52186e-05

0.000466064
0.000233032

0.000113761

0.000119271

6.26633e-05

6.35757e-05
0.0001271514

6.35757e-05

5.50531e-05

0.0001556128
7.78064e-05

7.78064e-05

5.98533e-05

0.0854460686
0.0218352213

0.000630423
0.001260846

0.00052058

0.000109843

0.000165093

0.0321851645
0.0108122237

0.0001330412
6.65206e-05

6.65206e-05

0.004801798
0.002400899

0.000178786

0.000104007

0.000126146

0.000156769

0.000118663

0.000449436

0.000549425

0.000103553

0.000163418

0.00012396

0.000326736

0.000225484
0.000112742

0.000112742

0.0001601878
8.00939e-05

8.00939e-05

8.8621e-05
0.000177242

8.8621e-05

4.83103e-05

0.000237335
0.00047467

0.000102495

0.00013484

0.000102659
0.000205318

0.000102659

0.000378843
0.000757686

0.000155723

0.00022312

0.0001864472
9.32236e-05

9.32236e-05

0.000342805
0.00068561

0.000171154

0.000171651

9.42109e-05
0.0001884218

9.42109e-05

0.00053074
0.00026537

0.00026537

7.20973e-05

0.000572822
0.000286411

0.000144369

0.000142042

0.0001612204
0.0003224408

8.29424e-05

7.8278e-05

0.000456334
0.000912668

0.000456334

5.10148e-05
0.0001020296

5.10148e-05

0.000457104
0.000228552

0.000228552

0.000103126
0.000206252

0.000103126

7.93363e-05
0.0001586726

7.93363e-05

0.0002448646
0.0004897292

8.7386e-05

7.68482e-05

8.06304e-05

0.0005094906
0.0002547453

7.15187e-05

7.27446e-05

0.000110482

0.000680355
0.00136071

0.000680355

0.000147416
7.3708e-05

7.3708e-05

0.0017123858
0.0034247716

7.0765e-05

6.78052e-05

7.95893e-05

9.17448e-05

0.00016392

5.71962e-05

6.11749e-05

6.16351e-05

5.95415e-05

0.00011933

0.000106806

6.73584e-05

4.62052e-05

6.37797e-05

5.8918e-05

6.0804e-05

6.18129e-05

0.000113525

6.4298e-05

6.99616e-05

8.56349e-05

8.05801e-05

7.20131e-05
0.0001440262

7.20131e-05

0.002176483
0.0010882415

0.000104996

0.000167461

8.48698e-05

0.000231747

0.00010412

0.000115757

7.94367e-05

8.889e-05

0.000110964

0.000131099

0.000648884
0.000324442

0.000179833

0.000144609

0.0002857926
0.0001428963

8.33103e-05

5.9586e-05

0.000675496
0.000337748

0.000337748

0.0287321539
0.0096076093

6.69522e-05
0.0001339044

6.69522e-05

0.001925538
0.003851076

0.000171947

0.000133699

0.000112839

0.000123254

0.000243095

0.000262059

0.000293628

0.000169908

0.000114215

0.000139581

0.000161313

0.0010518233
0.0021036466

4.76411e-05

9.74373e-05

0.00013075

0.00022515

5.63922e-05

5.21457e-05

0.000107035

5.07606e-05

0.000132118

4.74094e-05

0.000104984

0.0005286426
0.0002643213

7.69492e-05

7.88931e-05

0.000108479

6.60972e-05
0.0001321944

6.60972e-05

6.29711e-05
0.0001259422

6.29711e-05

0.001887428
0.000943714

5.02854e-05

0.000178022

0.000152284

6.69348e-05

6.06396e-05

0.000167527

6.17152e-05

6.70694e-05

8.44192e-05

5.48174e-05

0.0001684877
0.0003369754

0.000102523

6.59647e-05

0.0001027424
5.13712e-05

5.13712e-05

0.0004353514
0.0008707028

0.000136065

8.03719e-05

7.64852e-05

5.36154e-05

8.88139e-05

0.000471602
0.000943204

0.000218136

0.000128751

0.000124715

0.000111035
0.00022207

0.000111035

0.000105897
0.000211794

0.000105897

0.0001589159
0.0003178318

5.19469e-05

0.000106969

5.86621e-05
0.0001173242

5.86621e-05

0.00022519
0.000112595

0.000112595

0.0001931426
9.65713e-05

9.65713e-05

8.05726e-05
0.0001611452

8.05726e-05

0.0004090154
0.0002045077

4.80121e-05

4.85715e-05

5.2204e-05

5.57201e-05

0.0023677042
0.0047354084

0.000163516

5.10506e-05

5.2145e-05

0.000169557

0.000224531

0.000163277

0.000172264

5.24745e-05

0.000134606

0.000134606

6.23604e-05

0.00010964

4.52304e-05

0.000104668

6.44541e-05

0.000114424

8.4075e-05

4.19347e-05

5.49214e-05

0.000147517

0.00011067

8.93798e-05

8.65512e-05

6.84571e-05

9.0674e-05

0.0005704761
0.0011409522

7.48932e-05

0.000157184

7.24272e-05

8.63921e-05

9.47864e-05

8.47932e-05

0.000283538
0.000141769

0.000141769

0.000137335
0.00027467

0.000137335

0.000175386

0.0006096939
0.0002032313

0.0001480746
7.40373e-05

7.40373e-05

0.000258388
0.000129194

0.000129194

0.00010392
0.00020784

0.00010392

0.000207136
0.000103568

0.000103568

0.000120366
6.0183e-05

6.0183e-05

0.0297660629
0.007475014

0.007475014
0.0222910489

5.95877e-05

0.0007563159
0.0015126318

9.63297e-05

7.35332e-05

0.0001181

9.53632e-05

6.96137e-05

8.10176e-05

9.30628e-05

6.9517e-05

5.97787e-05

0.0064289652
0.0032144826

8.36206e-05

7.19683e-05

0.000109747

0.000108224

8.21111e-05

6.6064e-05

7.54431e-05

8.69957e-05

9.796e-05

0.00010167

8.38111e-05

0.000104807

0.000103498

5.54942e-05

0.000402104

0.000340148

4.59207e-05

0.000121042

0.000125046

9.21613e-05

0.00014015

0.000117871

5.79839e-05

0.000128533

0.000116883

0.000114078

9.7126e-05

8.40216e-05

0.0001175306
5.87653e-05

5.87653e-05

0.0001045446
5.22723e-05

5.22723e-05

7.44054e-05

0.0014538202
0.0007269101

8.12045e-05

6.45838e-05

6.87979e-05

0.000119828

7.81479e-05

9.19615e-05

0.000124383

9.80035e-05

0.0050645494
0.0025322747

6.24435e-05

6.05786e-05

5.65507e-05

5.68348e-05

6.95081e-05

6.77083e-05

6.67124e-05

0.000285133

8.83763e-05

0.000111157

0.00018626

5.24721e-05

9.28393e-05

6.72424e-05

6.12469e-05

6.94133e-05

0.000106802

6.44178e-05

9.64464e-05

5.37023e-05

6.11724e-05

6.45754e-05

0.000102785

6.04088e-05

5.71439e-05

6.58924e-05

8.68749e-05

0.000114846

6.54905e-05

7.72402e-05

1.5350322786554
0.384537145139

0.000118734

5.31847e-05

0.0022649745
0.0067949235

0.0004382296
0.0008764592

0.000141165

0.000118961

8.5598e-05

9.25056e-05

0.0006466624
0.0003233312

0.000133293

9.70589e-05

9.29793e-05

0.0005836442
0.0002918221

7.89321e-05

0.00021289

0.0005750865
0.001150173

5.55647e-05

7.68868e-05

8.42002e-05

6.17768e-05

7.46493e-05

5.8233e-05

6.10197e-05

0.000102756

0.0006365051
0.0012730102

6.8159e-05

0.000107133

6.99779e-05

0.000192855

6.85688e-05

6.01012e-05

6.97102e-05

8.71012e-05

0.0021226486
0.0063679458

8.3998e-05
4.1999e-05

4.1999e-05

0.0001696068
8.48034e-05

8.48034e-05

0.00011469
0.00022938

0.00011469

5.93416e-05
0.0001186832

5.93416e-05

8.98114e-05
0.0001796228

8.98114e-05

9.2482e-05
0.000184964

9.2482e-05

7.42702e-05
0.0001485404

7.42702e-05

0.003130502
0.001565251

0.0014509

0.000114351

0.0057721392
0.0019515672

0.000435422
0.000217711

0.000114253

0.000103458

8.25624e-05

0.000592716
0.000296358

9.28573e-05

0.000104324

9.91767e-05

0.000331002
0.000165501

0.000165501

0.0003720232
0.0001860116

0.000101123

8.48886e-05

0.0001877052
9.38526e-05

9.38526e-05

0.000239964
0.000119982

0.000119982

0.0015791772
0.0007895886

8.68016e-05

0.000702787

5.78963e-05
0.0001736889

0.0001157926
5.78963e-05

5.78963e-05

0.0503737947
0.0167912649

0.000274538
0.000137269

0.000137269

0.0319191382
0.0159595691

7.31278e-05

8.03576e-05

0.000186737

6.64471e-05

0.000123365

0.000559762

0.000123937

7.65868e-05

0.00117811

0.00117811

6.90277e-05

0.000284351

7.83621e-05

0.000124104

0.000152906

0.000115215

0.000143839

8.70547e-05

0.000113254

0.000441605

6.26261e-05

0.00015944

0.00289232

0.000223041

6.1719e-05

8.91781e-05

7.35876e-05

0.000102144

8.86462e-05

0.000120079

0.000168228

0.00017728

4.53781e-05

7.34353e-05

8.5103e-05

0.00116136

6.98021e-05

8.13119e-05

7.19466e-05

7.27241e-05

0.00012888

0.000138984

9.9159e-05

0.000115491

7.93235e-05

0.000244528

6.91283e-05

0.000118443

8.28733e-05

0.000104299

0.000358581

0.000157941

0.000151466

6.4935e-05

0.000115413

6.84822e-05

0.000146852

7.35252e-05

7.36393e-05

7.43769e-05

0.000119306

9.604e-05

0.00014578

7.28157e-05

0.00012947

0.000134025

0.000107113

9.21647e-05

0.000101544

0.000105294

0.000116608

8.04588e-05

0.000661644

0.000111573

7.68263e-05

7.8385e-05

0.000398702

0.00011234

0.000439578

0.000106313

0.000123271

0.000126498

0.0013888536
0.0006944268

7.65105e-05

6.83863e-05

0.000114683

7.44875e-05

9.01335e-05

0.000167991

0.000102235

7.01075e-05
0.000140215

7.01075e-05

0.0001790274
8.95137e-05

8.95137e-05

6.75443e-05

0.000467029
0.001401087

0.000467029
0.000934058

0.000138318

0.000118098

0.000109326

0.000101287

6.12435e-05

0.000105398

0.085399370949
0.028466456983

0.000115472
0.000230944

0.000115472

0.0061585652
0.0030792826

5.16148e-05

5.09359e-05

4.25284e-05

4.33832e-05

4.4531e-05

4.32862e-05

4.90976e-05

4.37367e-05

6.08285e-05

5.87618e-05

4.99218e-05

0.0009975062

8.79579e-05

5.5355e-05

7.47881e-05

4.92859e-05

4.42767e-05

6.1807e-05

0.000412705

5.41713e-05

0.000100999

5.61603e-05

4.23593e-05

5.13322e-05

4.29586e-05

5.07738e-05

4.86611e-05

4.75207e-05

4.45495e-05

6.21954e-05

0.000275185

3.99242e-05

5.65988e-05

4.60669e-05

5.9271e-05

4.41287e-05

4.98534e-05

4.95169e-05

6.22919e-05

5.23791e-05

7.88011e-05

4.45142e-05

8.32127e-05

0.000211056

0.0004378532
0.0008757064

7.45432e-05

0.000108625

0.000161444

9.3241e-05

0.016036102166
0.008018051083

0.00543333

5.92571e-05

6.23487e-05

7.29421e-05

6.5921e-05

1.45583e-07

0.000152424

5.38522e-05

5.18282e-05

6.0417e-05

0.000952778

5.12862e-05

7.95969e-05

0.000162939

0.000377158

7.61099e-05

7.87258e-05

0.000110165

5.42865e-05

6.25399e-05

0.0009596745
0.001919349

9.09112e-05

5.89881e-05

6.67318e-05

6.08485e-05

9.88699e-05

0.000339819

5.35845e-05

7.49544e-05

5.90154e-05

5.59517e-05

0.000333993
0.0001669965

5.43868e-05

4.9988e-05

6.26217e-05

0.0122329183
0.0244658366

5.53813e-05

5.56932e-05

0.000393927

6.21071e-05

5.59203e-05

4.73146e-05

6.72189e-05

5.83177e-05

5.09854e-05

7.70317e-05

5.01212e-05

4.48772e-05

6.31523e-05

6.08811e-05

0.000168273

8.71519e-05

6.70549e-05

9.59372e-05

0.000104959

7.67311e-05

0.000316447

4.91094e-05

7.53466e-05

6.54175e-05

5.64963e-05

5.34504e-05

4.03984e-05

4.84559e-05

4.99512e-05

4.84236e-05

5.39751e-05

6.73564e-05

5.19015e-05

5.75095e-05

4.79935e-05

5.50576e-05

8.84357e-05

0.000103309

4.84586e-05

4.05214e-05

4.87335e-05

4.87189e-05

4.68334e-05

7.86308e-05

0.000102936

4.41034e-05

5.41263e-05

4.22852e-05

5.70393e-05

5.92246e-05

5.17797e-05

4.8546e-05

6.57502e-05

5.10737e-05

4.8434e-05

6.65706e-05

7.20806e-05

5.77929e-05

6.37883e-05

6.36258e-05

4.86402e-05

0.000183563

0.000194445

7.60262e-05

6.03877e-05

5.51445e-05

6.1227e-05

4.86061e-05

5.27717e-05

0.000280136

5.38247e-05

4.44475e-05

4.54444e-05

7.31908e-05

0.000123386

4.90929e-05

5.85387e-05

6.28366e-05

4.76131e-05

4.52898e-05

4.80675e-05

4.58534e-05

5.32174e-05

4.12647e-05

9.99498e-05

6.82599e-05

5.59305e-05

5.99397e-05

4.96145e-05

5.76521e-05

5.01126e-05

5.54215e-05

6.50882e-05

4.94318e-05

5.31577e-05

7.28831e-05

6.8663e-05

6.00086e-05

4.56972e-05

5.2097e-05

5.53894e-05

7.86659e-05

0.000736006

4.65838e-05

4.66435e-05

6.27728e-05

4.63169e-05

6.31374e-05

5.36257e-05

6.2624e-05

5.86286e-05

4.96924e-05

5.74381e-05

0.000139018

7.57852e-05

5.20196e-05

7.76778e-05

5.59735e-05

4.49562e-05

6.64027e-05

4.90929e-05

5.26263e-05

4.72085e-05

8.60816e-05

4.86061e-05

5.75512e-05

5.07841e-05

4.45142e-05

5.65126e-05

4.25952e-05

4.05628e-05

6.4624e-05

5.44391e-05

5.6116e-05

5.35526e-05

0.000247094

4.32823e-05

4.7575e-05

5.85862e-05

4.99617e-05

6.36764e-05

5.2218e-05

5.45361e-05

4.59233e-05

5.69643e-05

4.52638e-05

5.22664e-05

5.85295e-05

7.52869e-05

0.000114724

5.84291e-05

5.04467e-05

7.89152e-05

6.28721e-05

6.39833e-05

7.45083e-05

5.29805e-05

4.67499e-05

5.03531e-05

8.36756e-05

6.10475e-05

8.32131e-05

7.09287e-05

5.59458e-05

0.000176965

0.000110806

4.88194e-05

5.21204e-05

0.0026699685
0.005339937

5.7877e-05

5.12115e-05

4.54141e-05

5.85454e-05

9.87641e-05

5.49819e-05

5.14981e-05

5.45293e-05

9.80899e-05

5.23161e-05

0.000161591

4.89176e-05

0.000411461

6.91936e-05

0.000103053

4.80509e-05

5.65442e-05

5.87946e-05

4.83706e-05

5.98785e-05

5.98694e-05

6.97491e-05

6.49404e-05

9.67608e-05

6.05831e-05

5.33314e-05

4.64694e-05

7.90247e-05

5.99147e-05

6.18333e-05

4.97678e-05

4.97622e-05

5.4666e-05

6.57349e-05

5.40108e-05

5.44681e-05

0.0004271509
0.0008543018

8.48931e-05

0.000146911

7.58748e-05

0.000119472

0.0007181788
0.0003590894

4.72154e-05

4.8977e-05

9.76817e-05

5.89173e-05

0.000106298

0.876336802099999
0.2921566088

0.2894530337
0.5789060674

6.52072e-05

3.9579e-05

6.14026e-05

0.000110201

6.69489e-05

5.00563e-05

5.49285e-05

6.16935e-05

7.1188e-05

3.88024e-05

6.63873e-05

5.76016e-05

3.86186e-05

3.76519e-05

5.25492e-05

3.79748e-05

4.10562e-05

4.39576e-05

4.48005e-05

5.24111e-05

5.16528e-05

7.1408e-05

5.55895e-05

3.64165e-05

6.01639e-05

5.38176e-05

8.76873e-05

5.51912e-05

0.00020416

5.02391e-05

4.93287e-05

4.47801e-05

3.88075e-05

4.25307e-05

5.82373e-05

6.25121e-05

5.33664e-05

0.000138656

5.49344e-05

5.34452e-05

7.2463e-05

5.35026e-05

4.52656e-05

8.7495e-05

4.98026e-05

6.45514e-05

3.82199e-05

3.86672e-05

8.82351e-05

5.16715e-05

6.0422e-05

4.79336e-05

3.66363e-05

4.32325e-05

5.86955e-05

4.8017e-05

4.9777e-05

4.94682e-05

6.37858e-05

7.19964e-05

5.36131e-05

3.44691e-05

7.48527e-05

5.71472e-05

3.43874e-05

6.91748e-05

0.0295438

0.000136976

3.61817e-05

5.4283e-05

8.32181e-05

0.0957198

4.60514e-05

6.17597e-05

8.52611e-05

5.52547e-05

5.14262e-05

5.77104e-05

5.71073e-05

4.74206e-05

6.17189e-05

3.51428e-05

0.000260243

0.00027727

5.88283e-05

0.00286573

4.99785e-05

0.000159562

4.22455e-05

3.67494e-05

3.81348e-05

5.94749e-05

5.03946e-05

3.78508e-05

4.4508e-05

4.84531e-05

5.03979e-05

7.42053e-05

0.000105966

6.06337e-05

5.93846e-05

0.00190238

7.22953e-05

4.35635e-05

5.01102e-05

4.89861e-05

4.75414e-05

6.85278e-05

6.09914e-05

0.00162556

7.27112e-05

0.1506

3.56686e-05

4.21112e-05

5.20632e-05

6.21982e-05

4.72999e-05

7.81985e-05

7.3856e-05

4.09175e-05

5.55084e-05

0.000853495
0.0004267475

4.96171e-05

0.000112759

8.59518e-05

6.04399e-05

6.05088e-05

5.74709e-05

0.0002284948
0.0001142474

6.50033e-05

4.92441e-05

0.0006477645
0.001295529

6.04838e-05

4.99067e-05

6.40882e-05

5.31743e-05

5.50626e-05

6.42278e-05

5.11062e-05

9.02889e-05

6.04912e-05

4.67113e-05

5.22235e-05

6.48013e-05

0.000366147
0.000732294

7.02212e-05

6.62971e-05

9.87457e-05

0.000130883

7.1743e-05
0.000143486

7.1743e-05

0.0012079966
0.0006039983

8.27404e-05

0.000109809

9.24019e-05

6.39148e-05

7.10245e-05

8.52802e-05

9.88275e-05

0.0001272513
0.0002545026

6.10357e-05

6.62156e-05

6.8223e-05

0.0002823076
0.0001411538

7.95525e-05

6.16013e-05

7.1498e-05
0.000142996

7.1498e-05

0.0153607227
0.0051202409

0.000131094
6.5547e-05

6.5547e-05

0.006019516
0.003009758

3.86619e-05

2.80776e-05

3.04445e-05

2.93823e-05

4.36262e-05

6.6986e-05

2.85016e-05

2.78728e-05

6.60162e-05

4.0947e-05

3.32889e-05

7.2458e-05

3.16804e-05

2.78481e-05

3.50408e-05

3.7442e-05

3.25496e-05

3.26094e-05

3.29846e-05

6.70187e-05

2.85114e-05

3.61024e-05

3.2445e-05

0.00111126

4.00833e-05

3.6333e-05

4.13035e-05

3.48258e-05

3.08517e-05

3.47882e-05

2.91895e-05

3.98771e-05

2.81824e-05

3.09552e-05

2.91832e-05

3.57768e-05

3.01995e-05

3.63352e-05

2.83248e-05

2.88978e-05

3.17851e-05

4.11812e-05

2.91349e-05

3.74252e-05

3.11344e-05

5.8346e-05

2.78985e-05

4.22619e-05

8.90293e-05

4.52298e-05

2.94693e-05

0.0002388028
0.0001194014

6.22374e-05

5.7164e-05

0.003851069
0.0019255345

5.28581e-05

4.50059e-05

4.09805e-05

6.84781e-05

9.50544e-05

5.76181e-05

0.000123116

5.17519e-05

0.000144352

4.86949e-05

4.56253e-05

4.58768e-05

4.66857e-05

0.000118448

5.36022e-05

4.53557e-05

5.35453e-05

4.59673e-05

4.41632e-05

4.51577e-05

9.60878e-05

7.13682e-05

6.28098e-05

4.89548e-05

4.42617e-05

4.63036e-05

5.56533e-05

4.89945e-05

4.20378e-05

4.66186e-05

9.01073e-05

8.69185e-05

0.0003724607
0.0011173821

0.0002943714
0.0005887428

6.72666e-05

6.72036e-05

9.09751e-05

6.89261e-05

0.0001561786
7.80893e-05

7.80893e-05

0.0042898227
0.0127829477

0.0001588722
0.0003177444

8.07644e-05

7.81078e-05

0.0003359082
0.0006718164

0.00011843

6.27752e-05

0.000154703

0.0058826956
0.0029413478

7.92293e-05

0.000147971

5.78575e-05

0.000178333

7.36451e-05

0.000111195

0.00101333

0.000464155

0.000122372

0.000140824

0.000244065

0.000123503

6.58819e-05

0.000118986

0.0007671741
0.0015343482

9.53247e-05

8.52455e-05

0.000128693

0.000147225

0.000117119

0.000113057

8.05099e-05

8.65204e-05

0.00011837
0.00023674

0.00011837

4.83753e-05

0.000700211
0.000257807

7.321e-05

0.000369194
0.000184597

0.000184597

4.23136e-05

5.4282e-05

0.020759663788
0.062231387364

0.0004288248
0.0002144124

4.96882e-05

6.0252e-05

4.70102e-05

5.7462e-05

0.0001329556
6.64778e-05

6.64778e-05

0.000125039
0.000250078

0.000125039

0.0024586522
0.0049173044

7.68576e-05

0.00140576

0.000109474

4.57717e-05

4.11613e-05

4.48483e-05

5.15643e-05

0.000132589

4.05939e-05

4.83078e-05

4.379e-05

4.3732e-05

4.89225e-05

6.37177e-05

5.69398e-05

4.42122e-05

6.66105e-05

9.37996e-05

0.0027763845
0.005552769

8.95809e-05

7.81544e-05

8.72146e-05

0.000138797

0.000409072

8.01858e-05

0.000125871

8.71453e-05

0.00010055

8.55335e-05

0.000228093

0.000177185

7.42074e-05

9.51787e-05

8.79822e-05

8.61913e-05

0.000137106

9.37516e-05

7.50662e-05

6.10206e-05

0.000108722

0.000115767

6.83633e-05

8.56457e-05

4.7604e-05

0.0009743688
0.0019487376

7.13919e-05

0.000766846

6.9919e-05

6.62119e-05

0.013895650188
0.027791300376

7.85746e-05

6.90416e-05

7.60749e-05

7.34219e-05

6.73498e-05

6.65067e-05

0.000171609

0.00011643

9.48318e-05

7.55528e-05

0.000163316

0.000293741

6.07993e-05

8.75547e-05

7.54533e-05

8.30608e-05

8.44953e-05

6.95773e-05

7.27248e-05

6.32253e-05

0.00015315

8.21483e-05

5.30116e-05

7.87729e-05

0.000110266

7.94416e-05

0.000246231

7.00172e-05

4.18851e-05

0.000170931

0.000447243

0.000178498

6.95766e-05

0.000114194

6.13797e-05

8.30972e-05

8.75483e-05

8.5444e-05

6.99727e-05

6.90145e-05

6.20182e-05

7.50558e-05

0.00012394

6.16227e-05

0.00268493

6.0617e-05

7.32838e-05

6.69643e-05

6.96312e-05

8.42415e-05

5.43588e-07

7.00709e-05

8.65598e-05

7.68182e-05

0.000103655

6.18402e-05

8.95377e-05

0.000113855

4.70252e-05

7.61015e-05

0.000120015

7.5151e-05

9.11187e-05

6.71961e-05

7.17067e-05

7.98492e-05

0.000122409

0.000134458

0.000186341

0.000100413

0.000112685

0.000101266

9.71857e-05

9.22273e-05

6.67643e-05

0.000197465

5.56358e-05

7.34373e-05

5.9404e-05

8.49359e-05

6.44958e-05

9.98233e-05

0.000119274

7.21903e-05

0.000375821

9.23766e-05

8.38932e-05

0.000155053

6.80657e-05

8.31737e-05

7.08415e-05

0.00170392

6.73041e-05

8.36979e-05

7.48992e-05

7.20879e-05

5.9729e-05

0.000110057

6.58078e-05

0.0004021498
0.0002010749

6.91763e-05

6.80494e-05

6.38492e-05

0.0001657378
8.28689e-05

8.28689e-05

0.0024206715
0.0008068905

0.0002406884
0.0004813768

7.94114e-05

0.000161277

0.0002063568
0.0004127136

5.55418e-05

8.27595e-05

6.80555e-05

0.0003955704
0.0001977852

5.138e-05

8.46291e-05

6.17761e-05

0.000213734
0.000106867

0.000106867

5.51931e-05
0.0001103862

5.51931e-05

0.0001819442
9.09721e-05

9.09721e-05

0.0066153866334
0.0022051288778

0.0022051288778
0.0044102577556

6.12275e-05

3.4028e-05

3.34145e-05

4.05945e-05

4.1196e-05

3.18798e-05

0.000180892

6.60921e-05

3.32098e-05

2.91778e-08

4.07028e-05

3.16137e-05

5.04532e-05

3.17112e-05

3.71916e-05

7.73868e-05

3.14303e-05

8.39584e-05

3.85663e-05

4.47994e-05

0.000111978

4.43118e-05

3.83828e-05

8.29851e-05

4.09244e-05

3.43796e-05

3.15279e-05

8.3482e-05

4.55564e-05

6.80162e-05

5.81661e-05

4.47855e-05

4.52098e-05

9.56242e-05

4.3347e-05

3.55185e-05

3.66777e-05

3.32343e-05

9.20354e-05

6.66667e-05

8.19424e-05

0.000103618

0.0001800612
9.00306e-05

9.00306e-05

0.0003981641
0.0007963282

8.91926e-05

5.81541e-05

5.02653e-05

5.20241e-05

8.7196e-05

6.1332e-05

0.01374605007
0.00458201669

0.0020727622
0.0010363811

7.92415e-05

0.000169713

7.30385e-05

6.69679e-05

7.7086e-05

6.88896e-05

6.41002e-05

0.000109819

0.000100102

7.05608e-05

7.97736e-05

7.7089e-05

0.0003022784
0.0001511392

5.91988e-05

9.19404e-05

0.0021962289
0.0043924578

6.49266e-05

7.01846e-05

5.47416e-05

0.000129663

7.88904e-05

6.2018e-05

7.43994e-05

6.75975e-05

5.85655e-05

0.000109203

8.3286e-05

8.31528e-05

9.79051e-05

5.6565e-05

6.99603e-05

7.0938e-05

8.61396e-05

6.4414e-05

8.34532e-05

0.000108422

7.17035e-05

6.89872e-05

6.21765e-05

9.35783e-05

5.71312e-05

5.78428e-05

6.16085e-05

5.21189e-05

9.66564e-05

0.000195671
0.000391342

0.000195671

0.00013471369
0.00026942738

3.29569e-06

0.000131418

0.000175376
0.000350752

0.000175376

0.0003441991
0.0006883982

0.000148812

6.62511e-05

0.000129136

5.51488e-05
0.0001102976

5.51488e-05

8.00101e-05
0.0001600202

8.00101e-05

0.0002131488
0.0004262976

5.43656e-05

4.95502e-05

0.000109233

0.0001918554
9.59277e-05

9.59277e-05

5.51942e-05
0.0001103884

5.51942e-05

0.0208739572
0.0833211472

0.0572295105
0.0190765035

4.17305e-05
8.3461e-05

4.17305e-05

0.0020166238

0.0010083119
0.0020166238

2.94907e-05

2.31943e-05

2.38959e-05

0.000155471

2.24347e-05

2.34593e-05

2.68906e-05

4.76725e-05

2.51669e-05

2.93821e-05

2.34806e-05

2.3233e-05

2.82233e-05

2.58491e-05

5.89477e-05

2.57065e-05

2.57959e-05

2.7446e-05

2.65805e-05

2.22201e-05

4.78159e-05

2.4461e-05

2.51219e-05

2.45403e-05

2.63583e-05

2.52378e-05

2.56491e-05

2.61248e-05

3.94932e-05

2.62852e-05

2.26837e-05

0.0012471475
0.002494295

3.38497e-05

0.0011816579

7.01025e-05

6.86938e-05

4.86441e-05

0.000950533

4.36845e-05

3.16399e-05

0.033311308
0.016655654

3.01519e-05

8.13853e-05

2.87168e-05

0.0165154

0.0165154

0.0002473192
0.0001236596

2.53416e-05

3.13122e-05

2.85573e-05

3.84485e-05

0.0004488024
0.0001496008

4.89472e-05
9.78944e-05

4.89472e-05

9.97368e-05
4.98684e-05

4.98684e-05

0.0001015704
5.07852e-05

5.07852e-05

9.38072e-05
4.69036e-05

4.69036e-05

6.3889e-05

0.0015370603
0.0046111809

0.0029903116

0.0013466292
0.0026932584

7.78195e-05

5.28634e-05

2.49561e-05

3.77294e-05

2.43757e-05

3.23989e-05

9.60224e-05

6.85332e-05

2.74892e-05

2.47516e-05

4.56515e-05

4.73986e-05

0.000930699

2.54312e-05

2.5592e-05

2.28489e-05

2.42033e-05

2.30606e-05

2.30124e-05

2.37092e-05

2.36339e-05

5.16708e-05

7.77325e-05

2.53096e-05

4.86905e-05

2.35582e-05

2.38318e-05

2.27598e-05

2.63077e-05

2.25236e-05

2.77005e-05

2.73564e-05

3.87991e-05

3.87991e-05

2.33266e-05

3.8715e-05

2.41499e-05

2.47737e-05

4.86771e-05

5.13074e-05

2.77531e-05

3.99996e-05

2.22383e-05

2.20263e-05

2.97826e-05

3.83842e-05
7.67684e-05

3.83842e-05

0.0002202848
0.0001101424

7.69534e-05

3.3189e-05

8.3809e-05
4.19045e-05

4.19045e-05

0.000633952
0.00294219

0.000113785
0.00022757

0.000113785

0.002080668
0.000520167

0.001560501
0.000520167

0.001040334
0.000520167

0.000102964

0.000112773

0.00010442

0.00020001

0.0013099107716
0.006549553858

0.0013099107716
0.0052396430864

0.000133809
0.000401427

0.000267618
0.000133809

0.000133809

0.0035283053148
0.0011761017716

0.0011761017716
0.0023522035432

0.000233644

0.000233238

0.000268848

9.29008e-05

8.51647e-05

1.52716e-08

0.000262291

0.0059843615
0.0011968723

0.0022398488
0.0005599622

0.0016798866
0.0005599622

0.0011199244
0.0005599622

0.000396995

7.25287e-05

9.04385e-05

0.0006369101
0.0025476404

0.0006131817
0.0002043939

0.0002043939
0.0004087878

9.97229e-05

0.000104671

0.0004325162
0.0012975486

0.0004325162
0.0008650324

6.9482e-05

9.57337e-05

8.9926e-05

9.43956e-05

8.29789e-05

81.3347157189991
16.2720144571

0.000668791
0.002675164

0.000668791
0.002006373

0.000668791
0.001337582

0.000171088

0.00015991

0.000174765

0.000163028

0.000109526

0.0012039476
0.0006019738

0.000123032

9.74872e-05

9.85135e-05

9.26009e-05

9.4002e-05

9.63382e-05

0.000480726
0.000240363

0.000240363

0.000212552

63.7157198860001
15.928994303

0.027222164
0.081666492

0.000651206
0.000325603

0.000325603

0.000760915
0.00152183

0.000368824

0.000392091

0.001877638
0.000938819

0.000938819

0.000678402
0.000339201

0.000339201

0.000697096
0.001394192

0.000697096

0.000514893
0.001029786

0.000514893

0.00171209
0.00342418

0.00060182

0.00111027

0.000625706
0.000312853

0.000312853

0.000683982
0.000341991

0.000341991

0.000389332
0.000778664

0.000389332

0.00102409
0.00204818

0.000565277

0.000458813

0.003063032
0.006126064

0.000556398

0.000722324

0.00108888

0.00069543

0.000856348
0.000428174

0.000428174

0.002293814
0.001146907

0.000513669

0.000633238

0.02522332
0.01261166

0.000555263

0.000570177

0.000587409

0.000585401

0.000503766

0.00111138

0.000661468

0.00090734

0.000603721

0.0011348

0.000554503

0.00124437

0.000613088

0.00148851

0.000602543

0.000887921

0.001302762
0.000651381

0.000321315

0.000330066

0.001285559
0.002571118

0.000702333

0.000583226

0.001357136
0.000678568

0.000384133

0.000294435

47.7050590910001
15.901772139

0.00868894
0.01737788

0.00868894

0.000257326

0.3276751
0.16383755

0.00701774

0.00608849

0.00619274

0.00579381

0.00633488

0.0529791

0.00742796

0.00975099

0.0114509

0.00623197

0.00575985

0.0101133

0.00637322

0.0223226

0.47833086
0.23916543

0.0164982

0.00791991

0.00623132

0.208516

0.00124479
0.000622395

0.000622395

0.000404561
0.000809122

0.000404561

15.23116822
30.46233644

0.198124

0.00937719

0.204287

0.0689946

0.105081

0.0129378

0.00520143

0.183204

0.0678817

0.0706751

0.00874629

0.316922

0.197614

0.205154

0.247471

0.105815

0.124039

0.176843

0.0124895

0.0236115

0.157672

0.00384993

0.0754667

0.247387

0.790335

0.206553

0.376811

0.0277578

0.0050679

0.0284946

0.192861

0.113183

0.0124203

0.00986903

0.122212

0.35246

0.255799

0.0107032

0.0243551

1.35597

0.209616

0.173197

0.078144

0.056004

0.0744698

0.0376466

0.28831

0.751942

0.167993

1.33245

0.160012

0.0544235

0.0863032

0.368968

0.20741

0.210027

0.0662037

0.00956206

0.0109089

0.0660275

0.00705738

0.203467

0.0670821

0.196043

0.125022

0.187582

0.191638

0.355358

0.0133135

0.00692639

0.0580413

0.0561469

0.0435421

0.189386

0.215549

0.632545

0.241833

0.211097

0.193738

0.00615446

0.101763

0.561429

0.00778846

0.17767

0.0576807

0.01259306
0.00629653

0.00324015

0.00305638

0.0089272
0.0178544

0.00198416

0.00133195

0.00561109

0.0326698
0.0653396

0.0326698

0.01105778
0.00552889

0.00274298

0.00278591

0.005354777
0.010709554

0.0019964

0.00151265

0.000158587

0.00168714

0.39399
0.196995

0.196995

0.00185552
0.00371104

0.00185552

0.000210375

0.000616073

0.0673718892
0.0172379705

0.0017229201
0.0051687603

0.000484264
0.000242132

0.000242132

0.002864876
0.001432438

0.000585016

0.000192917

0.000210487

0.000444018

9.67002e-05
4.83501e-05

4.83501e-05

0.00071001
0.00213003

0.00071001
0.00142002

0.000209861

0.000500149

0.017564484
0.005915326

0.000229132
0.000458264

0.000229132

0.003509174
0.001754587

0.000164677

0.000182883

0.00042921

0.000379691

0.000179112

0.000217864

0.00020115

0.000551558
0.001103116

0.000170785

0.000201165

0.000179608

0.000181494

0.000299542
0.000599084

0.000134213

0.000165329

0.001661574
0.000830787

0.000205097

0.000187445

0.000244167

0.000194078

0.002068226
0.004136452

0.000206849

0.000182273

0.000178182

0.000181579

0.000174529

0.000192646

0.00017867

0.000199365

0.000191723

0.000200831

0.000181579

0.000114481

0.00453375
0.00151125

0.0004108
0.0002054

0.0002054

0.000419596
0.000839192

0.000212706

0.00020689

0.000477682
0.000238841

0.000238841

0.000221519
0.000443038

0.000221519

0.000851788
0.000425894

0.000194103

0.000231791

0.000108199

8.93569e-05

0.006679215
0.020037645

0.002281078
0.004562156

0.00012994

0.000131195

0.000134461

0.000118737

0.000119134

0.000115668

0.000120895

0.000134253

0.000388938

0.000127121

0.000129738

0.000249683

0.000126269

0.000130345

0.000124701

0.007768284
0.003884142

0.000135262

0.00028344

0.000247214

0.00013597

0.000141805

0.000216963

0.000139838

0.000135983

0.000328761

0.000135871

0.000139812

0.000129952

0.000246814

0.000229546

0.000129511

0.000196075

0.000145219

0.000227392

0.000138305

0.000274718

0.000125691

0.000513995
0.00102799

0.000122901

0.000125414

0.00011776

0.00014792

0.000202718

9.51376e-05

8.93569e-05

0.311433233275
1.2307629070994

0.001303102
0.002606204

0.000210423

0.000154977

0.00017957

0.00018315

0.00021868

0.000182888

0.000173414

0.001521356
0.000760678

0.000414582

0.000346096

0.001042064
0.000521032

0.000234978

0.000286054

0.001198067
0.002396134

0.000145436

0.000146952

0.000165572

0.000195809

0.000202559

0.000207293

0.000134446

0.03486158
0.10458474

0.01485541
0.02971082

0.00015545

0.000156399

0.000318544

0.00015325

0.00031185

0.00489935

0.000238737

0.000176441

0.00017547

0.000342986

0.000166522

0.000182982

0.00303979

0.000339873

0.000763294

0.000163177

0.000318787

0.000217786

0.000397349

0.000263603

0.000349417

0.00056854

0.000155645

0.000224472

0.000225075

0.000182547

0.000193551

0.000174523

0.000522739
0.001045478

0.000297342

0.000225397

0.000784798
0.000392399

0.000193556

0.000198843

0.011572485
0.02314497

0.00043228

0.000153518

0.000215667

0.000151778

0.000187038

0.00127411

0.000141367

0.000176948

0.00021164

0.000364404

0.000154859

0.000163593

0.00017919

0.000160375

0.000221054

0.00016605

0.000379377

0.000169349

0.000226739

0.000189896

0.000149315

0.000411287

0.00015704

0.000159

0.00337559

0.000168535

0.000150512

0.000163216

0.000383536

0.000269817

0.00021261

0.000163801

0.000176686

0.000145485

0.000166823

0.000352854
0.000176427

0.000176427

0.00070563
0.000352815

0.000173866

0.000178949

0.000622323
0.001244646

0.000304176

0.000318147

0.000474617
0.000949234

0.000474617

0.00018846
0.00037692

0.00018846

0.001039982
0.000519991

0.000234084

0.000285907

0.001115778
0.000557889

0.000228437

0.000329452

0.001333119
0.002666238

0.000211

0.000462645

0.000212496

0.000446978

0.001671526
0.000835763

0.000197808

0.000156463

0.000166847

0.000170824

0.000143821

0.000344902
0.000172451

0.000172451

0.000904898
0.001809796

0.000563282

0.000182654

0.000158962

0.001126034
0.000563017

0.000278419

0.000284598

0.001245432
0.000622716

0.000164309

0.000200574

0.000257833

0.000194061
0.000388122

0.000194061

0.00017143

0.000635534
0.000317767

0.000317767

0.001187218
0.000593609

0.00020254

0.000224045

0.000167024

0.000352813
0.000705626

0.000172897

0.000179916

0.000180088

0.024739429
0.0738758129999999

0.002902284
0.001451142

0.00020745

0.000209328

0.000170964

0.000185351

0.000143097

0.000142816

0.000218026

0.00017411

0.007672829
0.015345658

0.000144628

0.000135954

0.000149578

0.000249

0.000132991

0.000166806

0.000156797

0.000153048

0.000156136

0.000126853

0.000124195

0.000135152

0.00042124

0.000127187

0.000156309

0.000129314

0.000165733

0.000125091

0.000140932

0.000145411

0.00014814

0.000139135

0.000148597

0.000163446

0.000357662

0.000172918

0.000212218

0.000151654

0.000145904

0.000129109

0.000167296

0.000127989

0.000168714

0.000217505

0.000127134

0.000160125

0.000197062

0.000140478

0.000128532

0.000130635

0.000150394

0.00014752

0.000157768

0.000153193

0.000148735

0.000139542

0.000199069

0.000173832
0.000347664

0.000173832

0.000945994
0.000472997

0.000148159

0.000182522

0.000142316

0.00839585
0.004197925

0.000190189

0.000171985

0.000192138

0.000271624

0.000159021

0.000151459

0.00017741

0.000174407

0.000184258

0.000166031

0.000167362

0.000161127

0.000162143

0.000332321

0.000160885

0.000193509

0.000211835

0.000169562

0.000190287

0.00019196

0.000157821

0.000260591

0.000155446

0.000412554
0.000825108

0.000412554

0.005238728
0.002619364

0.000150737

0.000179044

0.000160596

0.000155353

0.000183822

0.000142324

0.000212415

0.00014865

0.000181981

0.000223963

0.000162836

0.00017694

0.000196495

0.000188545

0.000155663

0.000187028

0.000384628
0.000192314

0.000192314

0.012175104
0.006087552

0.000259857

0.000186069

0.00021676

0.000201817

0.00017881

0.000223642

0.000213476

0.000145075

0.000240842

0.000214607

0.000168671

0.000152647

0.000234484

0.000381004

0.000207914

0.000184287

0.000384441

0.000187754

0.0001819

0.000193756

0.000222276

0.000217952

0.000215426

0.00020527

0.000220507

0.000209716

0.000212826

0.000225766

0.000175892
0.000351784

0.000175892

0.000940554
0.001881108

0.000168546

0.00020588

0.000162137

0.000213277

0.000190714

0.000828808
0.001657616

0.000206285

0.000218611

0.000195955

0.000207957

0.000156458

0.18665371296
0.55951373388

0.109125
0.21825

0.109125

0.002516044
0.005032088

0.000113787

0.000110449

0.000114782

0.000122409

0.000112078

0.000111336

0.000121512

0.000128824

0.000128213

0.000116797

0.000112468

0.000125808

0.000138729

0.000129171

0.00011675

0.000119983

0.000108243

0.000131271

0.000124698

0.000120808

0.000107928

0.008417966
0.004208983

0.000286639

0.000179528

0.000156168

0.000164538

0.000150138

0.000185034

0.000148941

0.000664523

0.000420307

0.000148896

0.000185768

0.00100596

0.000173867

0.00019165

0.000147026

0.000518984
0.000259492

0.000110296

0.000149196

0.000116838

0.009558716
0.004779358

0.000418348

0.000256931

0.00016867

0.000209245

0.000153842

0.000169444

0.000190614

0.000153547

0.000145019

0.000194894

0.000152296

0.000136574

0.000142685

0.000139156

0.00015439

0.000154575

0.000150238

0.00017865

0.000159156

0.000175196

0.000183099

0.000129384

0.000199816

0.000230116

0.000262315

0.000171158

0.000340772
0.000170386

0.000170386

0.003033601
0.006067202

0.000155791

0.000158385

0.000163709

0.000168071

0.000178428

0.000101323

0.000147972

0.000239821

0.000156907

0.00031745

0.000170852

0.000142603

0.000413383

0.000182304

0.000176077

0.000160525

0.000416674
0.000208337

0.000208337

0.000146025

0.01271065
0.0254213

0.00012236

0.000130402

0.000327161

0.000125327

0.000156948

0.000125841

0.000384683

0.000296908

0.000166269

0.0001393

0.000170991

0.000310077

0.000338032

0.000161033

0.000245464

0.000304419

0.000135448

0.000182214

0.000157237

0.000137244

0.0012297

0.000253039

0.00016592

0.000157551

0.000149727

0.000160412

0.000310026

0.000295962

0.00130924

0.000343912

0.00013249

0.000173737

0.000145807

0.000135496

0.000153652

0.000125097

0.000176645

0.000170275

0.000188097

0.000122636

0.000231055

0.000164839

0.000378964

0.000114177

0.000152552

0.000240116

0.000118414

0.00022586

0.000320598

0.000146039

0.000136802

0.000284623

0.000179832

0.001044163
0.002088326

0.000377817

0.000348059

0.000318287

0.04004982296
0.08009964592

0.000128591

0.000139363

0.000176737

0.007256197

0.000221808

0.000152813

0.000206327

0.000146852

0.000130393

0.00236544

0.000172251

0.000201615

0.000128645

0.000204131

0.00021976

0.000253319

0.00014014

0.000192685

0.000401855

0.00037674

0.000149861

0.000153338

0.000132442

0.000127311

0.000667989

0.000200113

0.000186241

0.000124128

0.000125697

0.000135675

0.000140582

0.00011504

0.000123213

0.000149802

0.00012743

0.000176015

0.00012408

0.000122171

0.000169665

0.000149284

0.000141066

0.000157237

0.000173452

0.000153646

0.000138887

0.000177074

0.000154999

0.000151768

0.000227955

0.000547802

0.000146537

0.000190123

0.000131717

0.000143627

0.00011092

0.000152197

0.000139873

0.000148003

0.000140492

0.000152626

0.000133783

0.000161722

0.000109154

0.000147913

0.000133567

0.000111542

0.000129898

0.000138629

0.000157005

0.000158878

0.000140143

0.00016414

0.00015153

0.00011937

0.000171386

0.000131034

0.000154391

0.000143736

0.000114108

0.000148191

0.000139443

0.000159473

0.000166808

0.000132677

0.000143505

0.000122857

0.000127087

0.00016461

0.000139536

0.00011597

0.000123402

0.000136024

3.57126e-06

0.000130343

0.000239116

0.000229886

0.00082397

0.000129671

0.000161169

0.000147566

0.000155759

0.000142443

0.00017704

0.000184704

0.000328056

0.000120166

0.00012189

0.000123402

0.000155178

0.000148187

0.000240578

0.000162161

0.000178102

0.000142616

0.000171457

0.000136898

0.000165134

7.89797e-05

0.000128476

0.00149843

0.000185418

0.000123632

0.000175082

0.000144382

0.000175662

0.000144207

0.015345436

0.000509696

0.000151778

0.000256125

0.000261475

0.00118452

0.000321028

0.00055657

0.000168786

0.000233673

0.000141586

0.00190207

0.000454006

0.000477407

0.000144458

0.000140036

0.000556096

0.000253596

0.000139535

0.000139794

0.0001287

0.000123282

0.000116572

0.000246724

0.000130143

0.000504645

0.00031419

0.000519121

0.000147259

0.000187695

0.000129673

0.000328737

0.00173564

0.000151949

0.000904433

0.00124763

0.000187406

0.000118735

0.000130667

0.016200942
0.008100471

0.000179418

0.000231007

0.00122944

0.000243329

0.000181674

0.0002078

0.000162037

0.000173564

0.000361068

0.000470482

0.000198998

0.000183552

0.000207128

0.000164147

0.000228608

0.000440952

0.000533162

0.000404468

0.000179047

0.000151953

0.000225543

0.000200474

0.00021245

0.000176524

0.000225582

0.000189802

0.000364595

0.000183336

0.000190331

0.000184542

0.000180184

0.002914928
0.001457464

0.000165961

0.000247319

0.00014866

0.0001262

0.000210346

0.000150975

0.000125579

0.000125728

0.000156696

0.000253047
0.000506094

4.8029e-05

0.000205018

0.001666926
0.005000778

0.000386966
0.000193483

0.000193483

0.001760952
0.000880476

0.000194275

0.000174475

0.000511726

0.000421494
0.000210747

0.000210747

0.000180604
0.000361208

0.000180604

0.000403232
0.000201616

0.000201616

0.000223881
0.000447762

0.000223881

0.000312899
0.000625798

0.000163869

0.00014903

0.000163031

0.00098733
0.000493665

0.000154603

0.000175703

0.000163359

0.000190173

0.0529691643148
0.1564901389444

0.000197617

0.003945444996
0.007890889992

0.00015708

0.000311337

0.000234848

0.000486852

0.000179025

0.000142659

0.000621445

0.00023402

0.00016671

0.000938142

7.01996e-07

0.000156976

0.00017056

0.000145089

0.001745588
0.000872794

0.000281661

0.000591133

0.001328172
0.000664086

0.000167474

0.000216478

0.000280134

0.000165376

0.000426332
0.000213166

0.000213166

0.000130259

0.000136201

0.005443787
0.010887574

0.00016543

0.00231475

0.00280795

0.000155657

0.001325448
0.002650896

0.000169904

0.000194559

0.000145207

0.000141596

0.000199823

0.000233985

0.000240374

0.000748181
0.001496362

0.000158728

0.000120644

0.000146964

0.000143353

0.000178492

0.004979412162
0.002489706081

5.66081e-07

0.000160167

0.000133501

0.000144266

0.000157597

0.000175618

0.0004797

0.000333953

0.000172938

0.000152081

0.000181003

0.000148727

0.000249589

0.000444547
0.000889094

0.000270364

0.000174183

0.000190201

0.000440192
0.000880384

0.000223457

0.000216735

0.008789399
0.017578798

0.00028763

0.000128401

0.000148542

0.000135734

0.000157899

0.000154744

0.000113054

0.000138739

0.000271581

0.000146889

0.000138323

0.000205861

0.000131345

0.000162029

0.000137854

0.0001221

0.000125093

0.000332789

0.000155712

0.000208446

0.000139639

0.000115323

0.000116231

0.000146471

0.00012568

0.000104264

0.000228041

0.000126105

0.000154456

0.000152395

0.000127754

0.000121887

0.000135343

0.000128658

0.000133383

0.000143417

0.000143242

0.000132723

0.000134778

0.000130628

0.000113753

0.000142892

0.00020354

0.000153929

0.000144129

0.000706276

0.000229735

0.000262155

0.000180484

0.000163515

0.000215936

0.000129872

0.00019565

0.000363264
0.000181632

0.000181632

0.000180992

0.004202376
0.002101188

0.000127225

0.00024322

0.000178592

0.000147881

0.000161509

0.000147336

0.000138437

0.000157995

0.000167147

0.000171316

0.000119505

0.000198473

0.000142552

0.000611236
0.000305618

0.000165081

0.000140537

0.000334961
0.000669922

0.000165471

0.00016949

0.000492663
0.000985326

0.000199072

0.000293591

0.000138299

0.000589922
0.000294961

0.000146118

0.000148843

0.000623575
0.00124715

0.000224314

0.000211655

0.000187606

0.007879296
0.003939648

0.000154729

0.000166265

0.000126502

0.000184093

0.000130487

0.00182809

0.000197722

0.000206675

0.000170372

0.000160057

0.000157692

0.000203269

0.000140475

0.00011322

0.000111531
0.000223062

0.000111531

0.000608058
0.000304029

0.000155778

0.000148251

0.0082561822
0.0041280911

0.000112731

0.000127258

0.000118403

0.000118267

0.000127824

0.000116505

0.000111266

0.000244632

0.000125725

0.000122849

0.000239131

0.000124483

0.000121621

0.000129345

0.000121718

0.000125746

0.000114613

0.000130405

0.000105517

0.000116817

0.000137497

0.00012622

9.73281e-05

0.000134469

0.00012273

0.000131796

0.000110514

0.000117592

0.000128889

0.000130721

0.000126504

0.000108975

0.00094582
0.00047291

0.00031129

0.00016162

0.000188857

0.000221493

0.001249713
0.002499426

0.000152105

0.000241655

0.000123342

0.000149367

0.000145785

0.000152054

0.000143131

0.000142274

0.000323756
0.000647512

0.000158742

0.000165014

0.0079708501378
0.0159417002756

0.000127299

0.00017735

0.00107751

0.000200509

0.00020769

0.000158631

0.000221697

0.000165106

0.000154055

0.000186652

0.000165696

0.000160299

0.000130433

0.000191084

0.00019177

0.000189134

0.000176502

0.000401291

0.000140213

0.00013345

0.000146642

0.000194439

5.41378e-08

0.000163681

0.000184518

0.000144709

0.000147871

0.000147681

0.000165259

0.000144682

0.000199233

0.000195889

0.000227822

0.000199799

0.000160481

0.00017382

0.000195804

0.0001339

0.000175854

0.000153432

0.000158909

0.000119055
0.00023811

0.000119055

0.000215069
0.000430138

0.000215069

0.000709107
0.001418214

0.000709107

0.000200647

0.000206741

0.000152371
0.000304742

0.000152371

0.00126767
0.000633835

0.000198486

0.000250954

0.000184395

0.001020992
0.000510496

0.000187686

0.000158285

0.000164525

0.000124458

0.000140563

0.000179008

0.001410434
0.000705217

0.00025349

0.000279594

0.000172133

7.39736e-05
3.69868e-05

2.1944e-05

1.50428e-05

5.51843e-05
0.0001103686

5.51843e-05

0.000167862

0.000125497

1.81824e-05
3.63648e-05

1.81824e-05

0.000113648

0.042284939
0.010746373

0.007676522
0.02280066

0.002380632
0.004761264

0.000232966

0.000310356

0.000247575

0.000249816

0.000266764

0.000270045

0.000305129

0.000238218

0.000259763

0.000228906

0.000531928
0.000265964

0.000265964

0.008683724
0.004341862

0.000238927

0.000458835

0.00022294

0.00025323

0.000219301

0.000209237

0.000228171

0.000240049

0.000291661

0.000216885

0.00020068

0.000230527

0.000289964

0.000235315

0.000238255

0.00027034

0.000297545

0.000459158
0.000918316

0.000260179

0.000198979

0.003132489
0.001044163

0.000656718
0.001313436

0.000218212

0.000222929

0.000215577

0.00077489
0.000387445

0.000387445

0.005605417
0.002025688

0.00046527
0.000232635

0.000232635

0.000233598

0.000694348
0.000347174

0.000347174

0.000459928
0.000919856

0.000241845

0.000218083

0.000238049

0.000228716
0.000457432

0.000228716

0.000285588
0.000571176

0.000285588

0.000181326

0.000244237

0.000132199

7.3522825484
36.7184298210748

0.037322452
0.009330613

0.027991839
0.009330613

0.000674784
0.000337392

0.000337392

0.00073471
0.000367355

0.000367355

0.008625866
0.017251732

0.000279309

0.00395595

0.00027926

0.000274899

0.000243505

0.000269841

0.00069769

0.0023207

0.000304712

0.00050223
0.000251115

0.000131973

0.000119142

0.020717543906
0.082677066624

0.056421098718
0.018807032906

0.001632306
0.000816153

0.000164696

0.000138223

0.000198124

0.000169264

0.000145846

0.00073286
0.00146572

0.000202469

0.000302035

0.000228356

0.00148683
0.000743415

0.000152621

0.000221125

0.000198746

0.000170923

0.001009534
0.000504767

0.00024633

0.000258437

0.032019675812
0.016009837906

0.000185672

2.86906e-07

0.000325783

0.000361492

0.000175794

0.000164229

0.000447303

0.000163896

0.000150207

0.000163538

0.000195431

0.000795119

0.000155794

0.000200577

0.000205902

0.000368394

0.000178121

0.000177606

0.000225872

0.000357874

0.000204399

0.000152651

0.000251471

0.000161369

0.000163975

0.00015697

0.000160669

0.000156837

0.000176283

0.000185436

0.000196672

0.000175876

0.000186999

0.000167973

0.000200139

0.000189857

0.000356668

0.000178382

0.000241805

0.000163282

0.000185943

0.000123752

0.000327229

0.000189275

0.000163472

0.000162652

0.00053701

0.000699626

0.00015028

0.000208964

0.000176188

0.000155244

0.00086339

0.000286509

0.0018123

0.000176821

0.000151497

0.00015295

0.000217551

0.000137726

0.000150759

0.000154096

0.001910511
0.005538424

0.000193109

0.00060882
0.00121764

0.00060882

0.000172646
0.000345292

0.000172646

0.000247252
0.000494504

0.000247252

0.00048314
0.00024157

0.00024157

0.000894228
0.000447114

0.000211278

0.000235836

8.75686e-05

0.00021008
0.00042016

0.00021008

9.34156e-05

0.0071186597
0.0276784606

0.002840955
0.008522865

0.000370974
0.000185487

0.000185487

0.000238264
0.000119132

0.000119132

0.000995854
0.001991708

0.000220522

0.000132932

0.000119713

0.000134587

0.000127281

0.000132425

0.000128394

0.000125882
0.000251764

0.000125882

0.000429008
0.000214504

0.000114393

0.000100111

0.000359659
0.000719318

0.000126851

0.000111116

0.000121692

0.0001784
0.0003568

0.0001784

0.0010977
0.00054885

0.000117025

0.000131209

0.000300616

0.000113187
0.000226374

0.000113187

0.0006325037
0.0002697743

8.83574e-05

8.84618e-05

9.29551e-05
0.0001859102

9.29551e-05

0.0048197049
0.0016065683

0.0032131366
0.0016065683

0.000125802

0.000155857

0.000149979

0.000111997

9.97205e-05

0.000107353

0.000108732

9.58797e-05

0.000151313

8.32942e-05

0.000118672

8.37479e-05

9.5152e-05

0.000119069

0.0034566297
0.0012914328

0.0001479716
7.39858e-05

7.39858e-05

9.75277e-05

9.11198e-05

0.0001800336
9.00168e-05

9.00168e-05

0.000816672
0.000408336

8.8139e-05

0.000105376

9.1949e-05

0.000122872

0.0001956608
9.78304e-05

9.78304e-05

0.00022111
0.000110555

0.000110555

8.16712e-05

0.0001860802
9.30401e-05

9.30401e-05

0.00014735

0.0011099293
0.0031280976

0.000329748
0.000659496

0.000104762

0.000116401

0.000108585

0.000207104
0.000103552

0.000103552

0.000103029

9.86613e-05

0.000481824
0.000240912

0.000109022

0.00013189

0.000468054
0.000234027

0.000119346

0.000114681

0.06830670669
0.27156053276001

0.0059461318
0.0178383954

0.0097647816
0.0048823908

0.000161662

0.000158453

0.000148855

0.00116103

0.00041739

0.000170855

9.25748e-05

0.00016555

0.000231687

0.000209716

0.000165201

0.000256773

0.000144486

0.00016197

0.000115935

0.00016308

0.000148409

0.00016073

0.000346718

0.00015079

0.000150526

0.00059951
0.000299755

0.000148165

0.00015159

0.000763986
0.001527972

0.000294705

0.000144709

0.000165368

0.000159204

0.003268126
0.009626723

0.000177655

0.006180942
0.003090471

0.000178687

0.000132851

0.000179284

0.000145008

0.000201558

0.000140532

0.000207228

0.000169596

0.000122207

0.000138846

0.000150044

0.000140951

0.000143357

0.000125253

0.000150365

0.000128443

0.000167617

0.000157868

0.000151213

0.000159563

0.000158516

0.05266712217001
0.01755570739

0.01725045739
0.03450091478001

0.000129235

0.000116743

0.000102113

0.000127161

0.000128033

0.000117457

0.000325816

0.000116866

0.000109438

0.000251089

0.000134211

0.000227885

0.000102573

0.000176029

0.00011344

0.000110662

0.000338508

9.8315e-05

0.000125715

0.00017518

0.000125058

0.00021749

0.000111406

0.00011473

0.00011249

0.000127283

0.000100126

0.000108037

0.000116536

0.000121223

0.000136944

0.000145034

0.000111552

0.000211632

0.00023852

0.000117635

0.000119361

7.36901e-09

0.000144932

0.000209335

0.000160393

0.000178249

0.000134935

9.60313e-05

9.21794e-05

0.000220773

0.000124584

0.000134134

0.000134034

0.000223976

0.000155278

0.000129176

0.000180205

0.000110909

0.00012266

0.000140343

0.000138564

0.00011276

0.000243078

0.000119472

0.000115889

0.000125047

0.000137956

0.000122648

0.000176683

0.000109451

0.000151346

0.000135289

0.000101726

0.000112731

0.000311671

9.35678e-05

0.000132671

0.000124098

0.000188787

0.000103988

0.00021415

0.000117817

0.000140223

0.00012137

0.000154482

0.000102617

0.000105672

0.000170204

0.000133735

0.000111946

9.47909e-05

0.000145913

0.000173428

0.00083491

0.000116183

0.000113239

0.00011337

0.000134807

0.000123173

0.000110259

0.000145354

0.000136669

0.000106352

0.000165714

4.28121e-07

0.000108909

0.000125522

0.000130025

9.54145e-05

0.000105376

0.00010298

0.000106698

0.000103685

0.000125925

0.000102928

0.000100955

0.00028667

0.000124187

0.000111194

0.000107472

0.000107415

0.000201023

0.00020619

0.000267774
0.000133887

0.000133887

0.000171363
0.000342726

0.000171363

0.000109452

0.014387847
0.043163541

0.028775694
0.014387847

0.000166467

0.000169334

0.000145919

0.000182636

0.000145162

0.000134558

0.000165251

0.000392579

0.000283008

0.000162278

0.000170836

0.000151421

0.000159281

0.000254051

0.000157103

0.000144319

0.00017029

0.000194563

0.000151924

0.000144364

0.000160873

0.000131297

0.000276984

0.000172977

0.000163304

0.000175208

0.000169885

0.000176838

0.000149743

0.000137902

0.000425419

0.000135094

0.000413057

0.000168057

0.000164967

0.000144233

0.000169049

0.000164024

0.000174882

0.000145293

0.000311864

0.00017767

0.000311396

0.000182324

0.000169241

0.000165319

0.00016034

0.000132334

0.000155396

0.00014999

0.00013364

0.000172703

0.000141241

0.000141628

0.000164752

0.000152431

0.000128795

0.000157929

0.000170478

0.00017745

0.000196729

0.00014824

0.000150697

0.00297083

0.000114903

0.000160067

0.001411499
0.004234497

0.000775602
0.000387801

0.00016403

0.000223771

0.002047396
0.001023698

0.000147792

0.0002399

0.000139353

0.000165512

0.000141818

0.000189323

0.0169067845
0.0503175905

0.000247486
0.000123743

0.000123743

0.0001692756
8.46378e-05

8.46378e-05

0.000733128
0.001466256

0.000367055

0.000197866

0.000168207

0.000812164
0.001624328

0.000174845

0.000151362

0.000162712

0.000160084

0.000163161

0.000349857
0.000699714

0.000165215

0.000184642

0.000745624
0.000372812

0.00024609

0.000126722

0.008695406
0.004347703

0.000197346

0.000154526

0.000133501

0.000131928

0.00017134

0.00016026

0.000127346

0.000257292

0.000131262

0.000140201

0.000127324

0.000126039

0.000149328

0.000123846

0.000350034

0.000171029

0.000131465

0.000130395

0.000266551

0.000183678

0.000132097

0.000144248

0.000560332

0.000146335

0.000149374
0.000298748

0.000149374

0.000164381
0.000328762

0.000164381

0.000555778
0.000277889

0.000147055

0.000130834

0.000395634
0.000791268

0.000188214

0.00020742

0.000168861

0.000102843
0.000205686

0.000102843

0.0125987714
0.0062993857

0.000168328

8.6783e-05

0.000233027

0.000147397

0.000100664

0.000103356

8.00677e-05

9.77759e-05

8.84996e-05

0.00011604

8.71873e-05

0.000120327

8.96754e-05

0.000106453

8.4792e-05

0.000123149

9.44785e-05

0.000130972

9.18491e-05

9.75808e-05

8.85685e-05

8.84719e-05

0.000126009

0.000102866

9.64387e-05

7.80136e-05

0.000101825

7.95746e-05

8.13818e-05

0.000174465

0.000117451

8.49202e-05

8.43331e-05

7.94805e-05

0.000131787

9.0195e-05

9.87814e-05

9.7408e-05

9.00129e-05

9.99081e-05

8.35128e-05

9.01333e-05

9.1273e-05

0.000101656

0.000188096

8.71476e-05

9.08501e-05

7.99387e-05

9.50714e-05

9.55775e-05

8.97975e-05

0.000108604

9.14744e-05

8.2495e-05

8.91065e-05

8.24195e-05

0.000108686

8.30838e-05

0.000105072

0.000106489

0.000108608

0.000107876

0.000126026

0.000171391
0.000342782

0.000171391

0.002455556
0.001227778

0.000151886

0.000218311

0.000135688

0.000141635

0.000136581

0.000166295

0.000133335

0.000144047

0.000891301
0.001782602

0.000184326

0.000128665

0.00016896

0.000154441

0.000125056

0.000129853

0.00129141
0.00387423

0.00258282
0.00129141

0.00032896

0.000235457

0.000179153

0.000183905

0.000181461

0.000182474

0.006996263
0.020988789

0.006996263
0.013992526

0.000175696

0.000159748

0.000163795

0.00533953

0.000117228

0.000163911

0.000152027

0.000195642

0.000190367

0.000174294

0.000164025

6.39736e-05

9.61377e-05

0.000125886

0.000122717

0.721982862158
2.88786172683278

0.15886406018
0.47659218053984

0.0305312220929
0.06106244418582

0.000128552

0.000157228

0.00015664

0.000112515

0.000131017

0.0002458

0.000117076

0.000115182

0.000136274

0.000192193

0.000132636

0.000124652

0.000643885

0.000138369

0.000134567

0.000139931

0.000128324

0.00012844

0.000162262

0.000256436

0.000131585

0.00015508

0.000115537

0.000125285

0.000124909

0.000119595

0.000176882

0.000139261

0.0236011

0.000133467

0.000142085

0.000156167

0.0001247

0.000132267

0.000120531

0.000115581

0.000124916

0.000183861

9.35371e-07

0.000109235

0.000141268

0.000127533

0.000108809

0.000129801

4.66601e-08

0.000129999

0.000107932

7.01359e-08

0.000126038

0.000144762

4.92592e-09

0.000224844
0.000449688

0.000107596

0.000117248

0.000706302
0.000353151

0.000123955

0.000125206

0.00010399

0.01725938721992
0.00862969360996

0.000206828

0.000206029

0.000246389

0.000186794

0.000233709

0.000462001

0.000418474

0.000243779

0.000320246

0.000229368

0.000183769

1.30498e-09

0.00053388

0.000233792

0.000216289

1.30498e-09

0.000191179

0.00232259

0.000228475

0.000193373

0.000192341

0.000347318

0.00025626

0.000976808

0.2382502989541
0.119125149477

0.000142321

0.000117792

0.000140343

0.000327331

0.000103756

0.000100191

8.4755e-05

0.00723617

0.000230119

0.000101878

0.000257887

0.000109264

0.000147272

0.000129435

0.000145131

0.000265145

0.000102263

0.000115272

0.00011368

0.000182925

2.99714e-07

0.000129627

9.98942e-05

0.000108115

9.29681e-05

0.000118234

0.000100439

0.000109184

9.17633e-05

0.000184537

0.000106798

0.000589566

0.000114159

9.37435e-05

0.00075108

0.000109196

0.000213365

0.000284189

0.000199629

0.000116805

0.000192671

0.00011132

0.000138296

9.67333e-05

0.000239971

0.000107099

9.01625e-05

9.65778e-05

9.62113e-05

0.000508782

0.000109046

0.000243045

9.35774e-05

0.000102314

0.000161045

9.51876e-05

8.92566e-05

0.000104441

9.59963e-05

0.000100454

0.000120728

0.000116305

9.83684e-05

0.000101107

0.000131428

0.000103953

0.000119536

0.000130244

0.000105474

0.000123943

0.000105339

0.000117987

0.000103155

0.00011256

0.000141952

0.000104682

0.000118283

0.000136941

0.000117333

0.000198809

0.000137296

0.000111171

0.000103403

0.000108497

0.000114236

0.000104194

9.61905e-05

0.0830711081

0.00010739

0.000176586

0.00014035

0.000711474

0.000114439

9.35429e-05

0.000104528

0.000431438

0.000122977

0.000103551

0.000102392

0.000103297

9.69206e-05

0.000103831

0.000114629

8.81237e-05

0.000101559

0.000101357

0.000104032

0.000116106

9.61164e-05

0.000107381

0.000109326

0.000137321

0.00010744

8.98951e-05

0.000110006

0.000102876

0.000100274

0.000103503

0.000107102

0.000123

0.000110533

0.000102578

0.000110363

9.91688e-05

0.000566582

8.45437e-05

8.87259e-05

9.58748e-05

0.00010061

9.88429e-05

0.000277431

0.000119687

0.000238034

9.3874e-05

9.70223e-05

0.00010881

9.70427e-05

9.33947e-05

0.000123277

0.000138744

9.65474e-05

0.000132038

0.000103938

9.74998e-05

0.0744987

9.50349e-05

0.000104701

0.000107109

0.000114107

9.13095e-05

0.000103848

0.000119708

0.000128665

0.000130171

9.1415e-05

9.06549e-05

0.000101226

9.38374e-05

0.000117329

0.000122526

0.000123027

9.34535e-05

0.000105315

0.000107522

0.000109218

0.000110383

0.000105902

0.000275496

0.000314878

0.000120593

0.000130962

0.000115365

0.000127027

0.000120915

0.000280287

0.000275289

0.000131332

6.09143e-08

3.253e-06

9.69405e-05

0.000369507

0.000152592

0.000114

0.000112727

0.000112533

0.000145259

0.000130006

0.00011135

0.000117527

0.000101322

0.000198036

0.000124558

0.0043812

9.88503e-05

0.000127156

0.000129157

8.7682e-05

0.000113502

0.000158267

0.000114916

0.00011364

0.000120784

0.000105262

0.000120023

0.000115215

9.93611e-05

0.000106218

0.000253584

0.000113043

2.95488e-08

0.000125015

9.25544e-05

0.000196341

0.000134024

0.000561185

0.000121023

0.000118008

0.000219099

0.000270779

0.000103434

0.000127521

0.000158373

0.00011635

0.000120342

0.000141622

8.86483e-05

0.000109137

0.000133006

0.000235813

0.000132558

0.000122356

0.000101121

0.000145269

0.000143392

0.000106949

0.000103692

0.000109032

0.000105162

0.000234644

0.000114083

0.562996974978
1.68892120313494

0.00022675
0.000113375

0.000113375

1.12284877835694
0.561424389178

6.91553e-05

6.48303e-05

7.96045e-05

6.98468e-05

4.83156e-05

0.000142234

6.98132e-05

7.12144e-05

0.000132546

0.000522944

9.99827e-05

8.11824e-05

7.87769e-05

6.98934e-05

8.04742e-05

6.40964e-05

8.28171e-05

0.000216139

5.94972e-05

7.60845e-05

8.73307e-05

0.000175098

6.10409e-05

7.75136e-05

6.09432e-05

6.81113e-05

7.19114e-05

6.31633e-05

6.03832e-05

6.48866e-05

0.000116717

6.801e-05

8.80846e-05

5.65408e-05

0.000149375

5.63628e-05

7.58418e-05

0.000104754

7.39742e-05

6.54175e-05

8.22471e-05

0.00024048

8.2697e-05

0.000105114

0.000198013

9.81055e-05

5.66778e-05

8.67016e-05

6.80868e-05

5.66129e-05

6.63012e-05

0.000105255

6.13674e-05

2.11892e-06

9.43994e-05

6.47798e-05

7.52382e-05

5.65175e-05

6.16606e-05

7.26464e-05

6.11853e-05

7.31353e-05

7.53266e-05

6.74116e-05

7.37695e-05

8.31357e-05

6.0589e-05

8.12189e-05

6.23556e-05

0.000108122

0.000253584

6.15004e-05

7.82964e-05

6.70481e-05

5.20491e-05

5.62928e-05

6.69435e-05

6.63576e-05

0.000149158

0.000138955

0.000130909

8.21876e-05

9.18998e-05

6.39179e-05

6.31605e-05

0.000100001

7.37642e-05

5.59723e-05

5.61645e-05

6.00825e-05

6.8871e-05

7.0981e-05

0.000118283

9.9094e-05

5.4071e-05

5.41415e-05

5.64239e-05

6.16928e-05

0.000133969

0.000114867

6.40366e-05

5.73516e-05

9.69268e-05

0.000113983

5.98322e-05

6.76534e-05

7.27637e-05

6.20482e-05

6.21602e-05

5.81649e-05

6.11795e-05

9.57969e-05

6.60232e-05

6.80184e-05

7.38735e-05

9.94324e-05

6.29514e-05

5.41026e-05

6.2756e-05

6.16687e-05

6.50042e-05

8.78624e-05

0.00011087

8.17773e-05

6.99556e-05

5.29984e-05

5.56824e-05

6.92636e-05

0.000225297

5.93866e-05

6.47161e-05

7.47073e-05

0.000112297

8.50445e-05

5.66682e-05

0.000125482

6.41231e-05

6.63691e-05

5.48711e-05

5.9687e-05

6.34305e-05

6.33634e-05

7.57254e-05

5.75272e-05

0.000104823

6.31258e-05

7.91247e-05

6.8681e-05

0.0004273

0.000109551

5.70788e-05

6.30429e-05

0.000107082

5.31607e-05

7.14872e-05

8.05051e-05

0.000101907

6.96871e-05

6.95915e-05

7.49702e-05

0.000108592

0.000107174

6.78895e-05

7.06672e-05

5.75854e-05

7.11517e-05

5.48251e-05

0.0181257586

0.000216604

0.000410826

0.0148723

8.52415e-05

0.000844176

0.00029312

0.000176398

0.000374658

0.000325762

0.000325762

0.000433351

0.000110943

9.75791e-05

0.000171637

0.00024626

0.000311079

6.03655e-05

6.45971e-05

6.34555e-05

0.000227858

8.91136e-05

7.26153e-05

6.9209e-05

7.76166e-05

0.000122016

0.00013519

9.03293e-05

5.219e-05

5.58294e-05

6.3854e-05

6.82949e-05

5.59007e-05

7.2068e-05

6.7576e-05

6.95229e-05

5.80314e-05

6.57678e-05

5.94062e-05

7.23339e-05

6.04186e-05

5.9371e-05

5.48227e-05

5.95841e-05

5.65715e-05

1.36344e-09

5.69068e-05

6.00729e-05

5.49801e-05

5.69203e-05

5.42249e-05

5.5698e-05

7.89083e-05

6.35583e-05

7.95861e-05

6.09765e-05

0.0185446

7.22185e-05

5.68012e-05

6.719e-05

6.98038e-05

6.628e-05

0.00011914

7.55083e-05

7.01419e-05

5.6347e-05

4.78044e-05

0.000799688

6.60926e-05

6.90504e-05

7.86367e-05

6.34031e-05

7.6984e-05

6.82829e-05

5.92658e-05

8.84503e-05

5.63816e-05

7.06004e-05

5.76734e-05

7.34532e-05

5.22124e-05

6.82416e-05

0.000125481

7.89434e-05

7.85845e-05

5.61108e-05

9.91469e-05

7.27839e-05

5.71511e-05

0.00011759

7.90101e-05

7.79114e-05

6.85463e-05

6.4679e-05

6.15613e-05

6.26088e-05

0.000117449

6.7187e-05

6.91013e-05

5.99748e-05

8.3218e-05

6.90719e-05

5.72385e-05

0.000107107

7.30519e-05

5.06412e-05

7.93347e-05

6.29452e-05

9.37083e-05

0.000110682

0.001446532

0.000127139

0.000302528

0.000472713

0.000544152

5.2631e-05

6.3218e-05

0.000128402

6.93624e-05

6.44787e-05

6.3306e-05

6.14914e-05

5.82815e-05

6.45618e-05

8.09383e-05

6.43846e-05

0.000156405

7.75856e-05

5.76375e-05

6.99623e-05

6.8443e-05

8.08002e-05

8.33534e-05

9.96564e-05

6.13202e-05

5.89839e-05

6.42907e-05

7.75307e-05

5.6098e-05

8.69256e-05

6.5814e-05

9.14402e-05

5.47703e-05

6.6424e-05

5.91805e-05

5.45956e-05

5.87608e-05

0.000113846

0.000102472

7.31395e-05

5.43914e-05

7.15164e-05

0.000155221

7.99504e-05

1.10524e-08

6.81218e-05

8.74237e-05

5.94195e-05

0.000161676

5.89996e-05

8.08987e-05

0.000115316

0.000420897

0.000167673

0.000253224

7.93037e-05

4.59853e-05

6.62205e-05

6.86624e-05

7.30562e-05

6.76448e-05

0.00214142

0.000282206

0.000229897

0.0014884

0.00010176

0.00138664

0.000140917

6.50433e-05

5.50007e-05

0.00010354

0.00012164

9.7357e-05

6.2337e-05

0.00273901

6.20343e-05

8.29514e-05

5.66578e-05

0.000247324

7.42763e-05

8.67648e-05

7.68772e-05

8.36757e-05

6.89381e-05

7.2101e-05

7.7083e-05

7.23713e-05

7.00235e-05

8.12852e-05

6.50049e-05

9.00637e-05

7.67338e-05

6.94591e-05

7.19734e-05

5.78249e-05

9.43056e-05

9.21244e-05

7.63271e-05

5.94233e-05

8.27839e-05

6.62447e-05

6.4739e-05

0.000377027

7.85253e-05

8.27511e-05

9.38597e-05

7.20863e-05

7.97752e-05

5.2939e-05

5.90326e-05

7.55012e-05

6.42622e-05

8.10977e-05

5.70551e-05

8.7815e-05

7.32899e-05

6.23865e-05

5.76318e-05

6.46465e-05

7.35688e-05

7.8734e-05

0.000147853

0.000148194

6.98672e-05

6.92579e-05

7.53083e-05

0.000107518

6.43112e-05

8.43277e-05

5.38325e-05

8.35493e-05

8.28864e-05

5.01344e-05

5.19397e-05

5.63596e-05

0.000167036

6.33416e-05

7.21952e-05

0.0116079047

0.00013645

0.000173404

0.000112194

9.02031e-05

0.000175778

0.000141112

8.64041e-05

7.90605e-05

7.13625e-05

0.000102857

0.000187203

0.00781808

0.000153691

0.00017911

0.000262314

0.000162412

0.000232017

6.42514e-05

0.000123326

0.000155608

0.000152854

9.59561e-05

0.000131796

0.0002171

0.000400875

0.000102486

5.20467e-05

6.47274e-05

9.2428e-05

0.000108728

6.17135e-05

7.20772e-05

9.76337e-05

5.07114e-05

6.5603e-05

8.27289e-05

6.26195e-05

5.47035e-05

0.000118228

5.61041e-05

1.14483e-07

7.03569e-05

7.38515e-05

7.31362e-05

5.92899e-05

9.46415e-05

8.87707e-05

5.29983e-05

6.22633e-05

7.38112e-05

7.56312e-05

6.76672e-05

5.71299e-05

6.6502e-05

8.82217e-05

8.18511e-05

6.67855e-05

6.04558e-05

6.14262e-05

5.9274e-05

6.0917e-05

4.95994e-05

5.1488e-05

5.95636e-05

6.05614e-05

8.6146e-05

5.96951e-05

7.09482e-05

5.74024e-05

6.20889e-05

5.6752e-05

7.90441e-05

7.97778e-05

5.92693e-05

7.03602e-05

6.4053e-05

5.71554e-05

8.96141e-05

7.35199e-05

6.98182e-05

7.88504e-05

5.95527e-05

6.21805e-05

5.62747e-05

5.40154e-05

6.5427e-05

6.00451e-05

6.39977e-05

0.000207862

7.41655e-05

5.76619e-05

5.98668e-05

6.95414e-05

5.61669e-05

6.83219e-05

0.000278704

9.37829e-05

6.46164e-05

8.53723e-05

5.81999e-05

6.8028e-05

6.22204e-05

5.68322e-05

0.000243767

6.78535e-05

6.51638e-05

6.07297e-05

9.03194e-05

0.000168391

8.67585e-05

8.8862e-05

1.38218e-06

8.09539e-05

5.51513e-05

7.5497e-05

5.96152e-05

6.48661e-05

7.07325e-05

0.000167668

5.11966e-05

5.12973e-05

6.54438e-05

5.64025e-05

5.84227e-05

6.96846e-05

6.12361e-05

6.75318e-05

6.45417e-05

0.000170696

0.000136776

0.000139609

8.16388e-05

6.73543e-05

6.62068e-05

8.04607e-05

6.8321e-05

7.22595e-05

7.56722e-05

7.33292e-05

5.94285e-05

5.7833e-05

0.000261479

6.98805e-05

5.67547e-05

5.39773e-05

6.41225e-05

0.000117782

5.67635e-05

8.14028e-05

5.64735e-05

6.19852e-05

8.20288e-05

5.28792e-05

7.6225e-05

6.39976e-05

9.53603e-05

6.14304e-05

7.52815e-05

7.66307e-05

8.10147e-05

7.30111e-05

0.000110381

6.90899e-05

7.73511e-05

5.85159e-05

8.09473e-05

6.95165e-05

9.86384e-05

0.000104108

8.04796e-05

6.44815e-05

8.4175e-05

5.68475e-05

6.14727e-05

7.99933e-05

6.9385e-05

9.70679e-05

7.18769e-05

5.81617e-05

6.70606e-05

5.37716e-05

5.25297e-05

0.000136728

7.9991e-05

0.000124985

6.50376e-05

5.66695e-05

0.000112411

7.31508e-05

8.95633e-05

0.000345255

6.0171e-05

6.00689e-05

0.000105996

7.36248e-05

0.000136839

0.000108404

8.56632e-05

0.000100695

7.12628e-05

6.64302e-05

7.5186e-05

7.41864e-05

5.97312e-05

7.07775e-05

0.000168663

0.000163147

7.3649e-05

9.83007e-05

6.08688e-05

6.25781e-05

7.76177e-05

7.03901e-05

9.17567e-05

7.31771e-05

8.58649e-05

6.99766e-05

6.59753e-05

8.06796e-05

6.24299e-05

6.76617e-05

5.71269e-05

9.73553e-05

8.53756e-05

7.64139e-05

7.12357e-05

6.15265e-05

5.72996e-05

6.76554e-05

7.72447e-05

7.64292e-05

9.33634e-05

0.000132064

8.8935e-05

9.73144e-05

5.98166e-05

8.1819e-05

6.2567e-05

6.69914e-05

0.000137006

5.67563e-05

5.82993e-05

5.68287e-05

5.42379e-05

6.58062e-05

7.10204e-05

6.31653e-05

0.00012628

8.26191e-05

9.38743e-05

7.47331e-05

6.62242e-05

7.56855e-05

6.46478e-05

6.85852e-05

6.56037e-05

0.000597723

7.5499e-05

7.74567e-05

5.34943e-05

6.21606e-05

7.13083e-05

0.000123269

6.93803e-05

8.42708e-05

6.83101e-05

0.000688037

6.21005e-05

0.000112612

8.5489e-05

6.26626e-05

9.46876e-05

7.739e-05

6.2216e-05

5.67831e-05

6.44924e-05

6.93321e-05

7.68932e-05

5.60032e-05

6.00401e-05

0.000125343

2.76064e-07

0.000178631

7.00596e-05

6.33897e-05

0.000106209

0.000120514

6.68872e-05

8.08703e-05

6.84487e-05

6.42517e-05

7.81528e-05

4.79328e-05

7.23169e-05

7.49144e-05

5.95367e-05

6.12042e-05

7.36082e-05

6.38135e-05

6.45119e-05

6.00055e-05

6.40958e-05

0.4348533505

0.000278742

0.0055650642

8.80625e-05

5.88497e-05

0.000167973

0.00491161

0.000338569

6.51499e-05

0.000151099

0.000112691

6.85245e-05

0.00012611

8.62767e-05

0.00022269

7.20734e-05

0.000386463

0.427319

0.427319

7.31128e-05

0.000106371

0.000219983

0.000138085

5.6301e-05

5.00896e-05

6.73504e-05

8.29373e-05

6.66712e-05

8.16518e-05

6.1969e-05

8.42385e-05

0.000104676

7.42132e-05

0.000107184

8.34521e-05

0.000118715

6.86701e-05

8.00836e-05

0.000102188

0.000234332

7.63207e-05

5.502e-05

0.000109397

7.37049e-05

6.52913e-05

0.000116288

6.77132e-05

6.52319e-05

0.000122086

7.60577e-05

0.000139263

6.14997e-05

7.49666e-05

5.67654e-05

9.49294e-05

7.28225e-05

6.22635e-05

0.000115718

5.53866e-05

6.89428e-05

0.000103975

0.003720501

0.000256714

0.000282149

0.00224522

0.000326879

0.000345857

0.000263682

6.74357e-05

6.69945e-05

6.86134e-05

0.000130211

6.51619e-05

6.39766e-05

9.80941e-05

0.000104526

8.15755e-05

0.000101612

5.85612e-05

7.64279e-05

7.71884e-05

7.67937e-05

0.000109361

8.78875e-05

7.17241e-05

8.90395e-05

7.46402e-05

6.46305e-05

8.07782e-05

6.4072e-05

4.90941e-05

8.51003e-05

5.78341e-05

0.000148433

0.000176668

6.84465e-05

5.87941e-05

6.91724e-05

7.08978e-05

8.08294e-05

6.11936e-05

7.71688e-05

7.74084e-05

5.8347e-05

9.5095e-05

7.27222e-05

5.87597e-05

5.49526e-05

7.34996e-05

6.59349e-05

8.94801e-05

0.000129466

5.77252e-05

6.30183e-05

6.75829e-05

6.14685e-05

7.92419e-05

7.69879e-05

5.69418e-05

7.97261e-05

5.87744e-05

8.0784e-05

7.60299e-05

5.90847e-05

0.000101669

8.06143e-05

7.30031e-05

0.000109684

8.21121e-05

6.06037e-05

7.09395e-05

8.08932e-05

0.00010685

0.000109142

6.22343e-05

6.73472e-05

7.80121e-05

5.87987e-05

6.71677e-05

7.75709e-05

8.2428e-05

7.75309e-05

0.000162493

9.10028e-05

6.47317e-05

9.43525e-05

8.86654e-05

7.40594e-05

8.49182e-05

6.73963e-05

6.28594e-05

7.11514e-05

6.11794e-05

0.000104027

6.73413e-05

5.60832e-05

5.70659e-05

5.09108e-05

5.89967e-05

7.10663e-05

7.94491e-05

0.000102562

5.96836e-05

6.91218e-05

6.91477e-05

7.86629e-05

6.4911e-05

6.13119e-05

0.000267147

0.000134092

6.92766e-05

0.000112085

5.68777e-05

5.63356e-05

0.000126177

5.7992e-05

0.00013981

5.80546e-05

0.000235598

8.1096e-05

0.000110501

5.94244e-05

0.000100326

8.56651e-05

6.45202e-05

4.58161e-08

7.03204e-05

5.35955e-05

8.94544e-05

8.88695e-05

7.27146e-05

0.000113776

0.00173084

0.00173084
0.00086542

0.000224848

0.000318906

0.000321666

6.97218e-05

0.00046852
0.00023426

0.00023426

0.000579618
0.000289809

0.000289809

0.000365481
0.000121827

0.000243654
0.000121827

0.000121827

0.000129568

0.000134934

0.00065222
0.00032611

0.00016111

0.000165

0.0006823342
0.0003411671

0.000149887

9.57428e-05

9.55373e-05

0.021105212
0.005276303

0.000350277
0.000116759

0.000233518
0.000116759

0.000116759

0.000526302
0.000175434

0.000350868
0.000175434

0.000175434

0.00498411
0.01495233

0.000235656
0.000117828

0.000117828

0.000115962
0.000231924

0.000115962

0.000745208
0.000372604

0.000110245

0.000129229

0.00013313

0.000255896
0.000127948

0.000127948

0.000449302
0.000224651

0.000113559

0.000111092

0.000346219
0.000692438

0.000149157

0.000197062

0.000135936
0.000271872

0.000135936

0.00038695
0.0007739

0.000138128

0.000126565

0.000122257

0.001502948
0.000751474

0.000102304

0.000125274

0.000307192

0.000103801

0.000112903

0.001652533
0.003305066

0.000334235

0.000108274

0.000137865

0.000360613

0.000120256

0.000112096

0.000134666

0.00011678

0.000107123

0.000120625

0.000262814
0.000131407

0.000131407

0.000178365
0.00035673

0.000178365

0.000336892
0.000168446

0.000168446

0.000273787
0.000547574

0.000273787

0.0150794751
0.0601867884

0.0002772996
9.24332e-05

9.24332e-05
0.0001848664

9.24332e-05

0.0165291972
0.0055534364

0.00011905
0.0002381

0.00011905

0.000307046
0.000153523

0.000153523

0.00023974
0.00011987

0.00011987

0.000397392
0.000198696

0.000198696

0.000292671
0.000585342

0.000148218

0.000144453

0.000149426
0.000298852

0.000149426

0.000543238
0.000271619

0.000161023

0.000110596

0.000337336
0.000168668

0.000168668

0.000164957
0.000329914

0.000164957

0.00035942
0.00071884

0.000194835

0.000164585

0.000140102
0.000280204

0.000140102

0.000331682
0.000165841

0.000165841

0.000661386
0.000330693

0.000154316

0.000176377

0.000427634
0.000213817

0.000213817

0.000131112

0.003041414
0.001520707

0.000150717

0.000156864

0.000152992

0.000148032

0.000144953

0.00015803

0.00016907

0.000134607

0.000135482

0.00016996

0.000265174
0.000132587

0.000132587

0.001058898
0.000529449

0.000178384

0.000138506

0.000212559

0.0007824568
0.0003912284

9.91204e-05

0.000189878

0.00010223

0.0013652304
0.0004550768

0.0009101536
0.0004550768

9.15479e-05

0.00013652

9.15479e-05

0.000135461

0.0087131647
0.0261394941

0.001447946
0.000723973

0.000137309

0.000131538

0.000113773

0.000122387

0.000108015

0.000110951

0.000103866
0.000207732

0.000103866

0.00025844
0.00012922

0.00012922

0.000138435
0.00027687

0.000138435

0.0001910082
9.55041e-05

9.55041e-05

0.000248086
0.000124043

0.000124043

0.000103751
0.000207502

0.000103751

8.95923e-05
0.0001791846

8.95923e-05

0.0001644081
0.0003288162

8.29938e-05

8.14143e-05

0.0001854132
9.27066e-05

9.27066e-05

0.000208466
0.000104233

0.000104233

0.0122520452
0.0061260226

7.6796e-05

9.05153e-05

0.000214395

6.61138e-05

7.96585e-05

7.01994e-05

7.67962e-05

7.69374e-05

0.000154767

7.2802e-05

6.68065e-05

7.74143e-05

8.09834e-05

7.43782e-05

8.1777e-05

0.000109451

0.000121325

7.57426e-05

9.12812e-05

0.000102061

7.9775e-05

8.47192e-05

7.21237e-05

7.82395e-05

7.49359e-05

0.000142818

7.67911e-05

6.9692e-05

8.07905e-05

7.79071e-05

0.000116046

6.98102e-05

6.67035e-05

7.66083e-05

5.9671e-05

7.16356e-05

6.68245e-05

7.92868e-05

8.4255e-05

8.73199e-05

7.66196e-05

7.25745e-05

7.43886e-05

8.68695e-05

6.22307e-05

9.04647e-05

6.92272e-05

8.70863e-05

7.0209e-05

7.78661e-05

7.21302e-05

7.41712e-05

7.26018e-05

6.87295e-05

6.39226e-05

7.30199e-05

7.8422e-05

7.38571e-05

7.65511e-05

6.92028e-05

6.73986e-05

0.000103728

6.52226e-05

7.34395e-05

8.38035e-05

7.2283e-05

7.7288e-05

7.95648e-05

7.54231e-05

7.56889e-05

7.92403e-05

7.27049e-05

8.47524e-05

7.12597e-05

7.79273e-05

0.000344186
0.000172093

0.000172093

0.000142774
0.000285548

0.000142774

0.000402543
0.000805086

0.000123245

0.000279298

0.000360645
0.000120215

0.00024043
0.000120215

0.000120215

0.000145149
0.000435447

0.000290298
0.000145149

0.000145149

0.000386463
0.000128821

0.000257642
0.000128821

0.000128821

0.000268516
0.000134258

0.000134258

0.20096936005592
0.050740501239

0.0212904432
0.0071501054

0.000247758
0.000123879

0.000123879

0.00029348
0.00014674

0.00014674

0.0056630018
0.0028315009

9.22332e-05

0.00018367

8.11115e-05

9.14317e-05

0.000140208

9.40184e-05

0.000105031

8.55762e-05

0.000109653

8.48378e-05

0.000113707

0.000102807

0.000105249

0.000115815

0.000107661

9.84891e-05

9.292e-05

0.000102108

0.000287059

0.00011232

0.000123846

9.46718e-05

0.000121309

0.000101001

8.47672e-05

0.000159873

0.000265243
0.000530486

9.3283e-05

0.00017196

0.00081582
0.00040791

0.000170961

0.000111552

0.000125397

0.000116551
0.000233102

0.000116551

0.000105639
0.000211278

0.000105639

0.0028956716
0.0014478358

0.000147539

8.14936e-05

9.66976e-05

0.000113198

0.000127188

0.000113981

9.27157e-05

0.000123627

0.000113077

0.000230352

9.10509e-05

0.000116916

0.000102873
0.000205746

0.000102873

0.0012748137
0.0025496274

0.000324969

0.00013961

0.00013481

9.68357e-05

0.000112942

0.000212381

0.000129193

0.000124073

0.000334494
0.000167247

0.000167247

0.12893841561692
0.043590395839

0.000130274

0.000469254
0.000234627

0.000129702

0.000104925

0.000127587

0.000248284

0.0026258635
0.005251727

0.000100162

9.15605e-05

9.06989e-05

0.000123666

9.44022e-05

0.000126362

0.00010545

0.000109512

8.4357e-05

0.000186611

8.64111e-05

0.000335842

8.90118e-05

0.000319642

0.000120261

0.000561914

0.000100573

0.000135875

8.41766e-05

0.000134685

0.0076737226
0.0038368613

9.12753e-05

0.000201062

0.000112259

8.90322e-05

0.000134899

0.000225696

0.000111411

0.000513395

0.000103593

0.00010036

0.000353729

0.000130808

0.000364102

0.000129475

0.000121639

0.000108928

0.0001143

9.22778e-05

9.75174e-05

0.000107759

0.000107913

0.00010319

9.88006e-05

8.8434e-05

0.000135006

7.80779e-05

0.0005878182
0.0002939091

7.47421e-05

0.000219167

0.000122766

0.001574106
0.000787053

0.000104385

0.000102301

0.000142892

0.000114745

0.000192174

0.000130556

0.000108723
0.000217446

0.000108723

0.000147533

0.01971885807792
0.00985942903896

9.31679e-05

8.22896e-05

9.97393e-05

8.81724e-05

0.000108906

8.20748e-05

7.85864e-05

9.36173e-05

0.000102323

0.000993857

9.6787e-05

0.000100904

0.000194253

7.86269e-05

7.81517e-05

9.20049e-05

0.000111104

0.000138348

0.00018283

0.000103492

9.0506e-05

9.11114e-05

0.000263648

0.000127408

0.000129205

8.19503e-05

9.52193e-05

0.000101031

9.70222e-05

0.000101033

9.63445e-05

0.000238763

0.000101765

7.74231e-05

8.38986e-05

8.18112e-05

0.000102891

3.03896e-09

0.000121708

0.00322578

9.86566e-05

0.000116863

7.66711e-05

8.19323e-05

8.01524e-05

0.000118817

9.7209e-05

9.23498e-05

0.000176174

8.12358e-05

0.000175363

7.70114e-05

8.17486e-05

0.000100052

9.74362e-05

0.000120947

0.000183441

8.95754e-05

0.002481751
0.004963502

0.000106741

0.00083447

0.000150075

0.000124645

0.000110109

0.000860526

0.000165805

0.00012938

0.000128977

0.043058814
0.021529407

0.0025832

8.25667e-05

8.52099e-05

0.000238661

0.00012808

0.000935729

0.000153723

0.000103502

0.000519907

8.77435e-05

0.000203358

7.29978e-05

0.000103976

0.000400198

0.000207754

0.000264156

0.00953333

8.14456e-05

9.13459e-05

7.7222e-05

0.000122279

0.000118736

0.000397564

9.67261e-05

7.69929e-05

0.000549371

7.91358e-05

0.000127058

0.000177588

0.00014208

0.000868292

0.000868292

0.000498737

8.93727e-05

8.25709e-05

0.000151976

9.66782e-05

0.000130871

0.000224774

7.7559e-05

0.00010064

0.000865925

0.00014186

0.000220919

0.000137596

8.5118e-05

0.000225865
0.00090346

0.000225865
0.000677595

0.00045173
0.000225865

0.00010477

0.000121095

0.000260885

0.0003555028
0.0001777514

9.47906e-05

8.29608e-05

9.41853e-05

8.43798e-05

0.075097903424
0.299960610696

0.000145776

0.224716931272
0.074952127424

0.003199086
0.001599543

0.000583784

0.000189524

0.000188042

0.000442016

0.000196177

0.100126613688
0.050063306844

0.000158826

0.000126513

0.000197351

0.000145452

0.000495149

0.000170064

0.0001776

0.000119318

0.000130236

0.000154717

0.000166593

0.000120706

0.000198403

0.000117176

0.000119808

0.001604038

0.000336059

0.000409851

0.000152504

0.000162408

0.000224304

0.000162458

0.000156454

0.000420015

0.000319147

0.00019576

0.000152104

0.000140004

5.87744e-05

0.000484917

0.000159269

0.000120213

0.00019233

0.000163153

0.00013914

0.000182787

0.000138923

0.000160962

0.000133681

0.000175326

0.00015676

0.000158498

0.000126394

0.000166909

0.000388205

0.000207149

0.000140945

0.000194284

0.000153452

0.000162677

0.000116266

0.000137264

0.000211636

0.000150434

0.000119316

0.000281095

0.000320333

0.000416448

0.000173533

0.000508088

0.000139894

0.00011481

0.000139165

0.000226974

0.000185473

0.000140713

0.00011692

0.000313835

0.000152636

0.000163238

0.000149008

0.000112595

0.00012173

0.00013699

0.000149921

0.000129812

0.000122555

0.0002329

0.000140394

0.000159286

0.000167873

0.000212057

0.00012035

0.000334145

0.000121028

0.00015825

0.000150949

0.000114238

0.000141283

0.000168452

0.000135775

0.000195253

0.000388924

0.000158591

0.000190055

0.000142385

0.000168252

0.000117181

0.00200973

0.000114496

0.000175971

0.0002616

0.000165884

0.000137233

0.000119934

0.000121743

0.000162784

0.000145987

0.000445805

0.00023208

1.31992e-06

0.000150403

0.000269451

0.000114957

0.000200746

0.000119073

0.000160828

0.000451022

0.000145623

0.000162512

0.000147918

0.000187609

0.000173971

0.000169689

0.000129403

0.000350078

0.000119814

0.000144556

0.000183552

0.000118744

0.000125739

0.000134129

0.000340282

0.000119939

0.000150077

0.000149308

0.00014429

0.000445925

0.000165314

0.000133976

0.000342665

0.00227923

0.000118261

0.000158205

0.000115432

0.000152013

0.00050877

0.000120021

0.000883435

0.000171299

0.000476349

0.000153053

0.0003201

0.000167389

0.000160823

0.00014656

0.000194917

0.000692416

0.00036092

6.59224e-07

0.000118228

0.000310519

0.00033219

0.000196262

0.000126976

0.000277862

0.000119025

0.000120736

0.000165664

0.000115541

0.000113348

0.000141484

0.000487475

0.000153755

0.000141469

0.000122589

0.009680296

0.000705853

0.00381234

0.000367663

0.000307346

0.000659436

0.000177261

0.000140086

0.000466451

0.00033451

0.000124967

0.000303522

0.000755954

0.000674179

0.000850728

0.000109407

6.61523e-05

0.000554768

0.000139456

0.000198973

0.000416326

0.000150293

0.000158667

0.000150627

0.000207175

0.000139451

0.003235504
0.001617752

0.000169575

0.000277693

0.000168182

0.000725084

0.000277218

0.010207016
0.005103508

0.000152708

0.000146528

0.000157059

0.000146324

0.000153547

0.000387209

0.000150345

0.000143964

0.000164249

0.000144981

0.00014696

0.000140816

0.000147138

0.000147232

0.00024022

0.000148383

0.000171843

0.000299429

0.000154698

0.00016569

0.000226898

0.000152842

0.000157001

0.000147075

0.000150865

0.000156308

0.000145928

0.000146754

0.000148809

0.000161705

0.0004450232
0.0002225116

8.14756e-05

0.000141036

0.02668409996
0.01334204998

0.000172522

0.000195688

0.000194707

0.000211319

0.000417036

0.000232137

0.000578329

0.000162688

0.000200285

0.000163219

0.000518361

0.000293064

0.000201858

0.000273783

0.000489643

0.000181665

0.000193096

0.000152028

0.00026092

0.00019732

0.00017602

0.000211149

0.000291854

0.000196779

0.000347584

0.000415912

0.000503158

1.44198e-06

0.000329855

0.000161749

0.00212491

0.000197997

0.000184796

0.000780443

0.000352726

0.000170509

0.000331671

0.000316882

0.000202594

0.000169397

0.000178923

0.000194839

0.000211193

0.000316818
0.000158409

0.000158409

0.004566674
0.002283337

0.000169039

0.000417738

0.000185198

0.000305904

0.000168768

0.000161851

0.000328161

0.000546678

0.000422259
0.000844518

0.000205304

0.000216955

0.000938148
0.001876296

0.000170268

0.000172442

0.000426768

0.00016867

0.00016984
0.00033968

0.00016984

0.000253672
0.000126836

0.000126836

0.22249376317283
0.0556861980432

0.034903593
0.011634531

0.000130859
0.000261718

0.000130859

0.000476024
0.000238012

0.000238012

0.004250866
0.008501732

0.000153107

0.000148778

0.000120989

0.000129782

0.000142299

0.000136941

0.000233982

0.000123851

0.000136128

0.000180849

0.000124668

0.000143833

0.000255306

0.000407944

0.000615278

0.000124974

0.000137427

0.000128091

0.000806639

0.006809856
0.003404928

0.000130266

0.000114383

0.000191894

0.00011927

0.000123342

0.000130729

0.000106227

0.000178239

0.00219779

0.000112788

0.000276414
0.000138207

0.000138207

0.001010444
0.000505222

0.000123951

0.000141542

0.000132613

0.000107116

0.000226784
0.000453568

0.000111397

0.000115387

0.000596475
0.00119295

0.00018797

0.000172694

0.000128096

0.000107715

0.000247006
0.000123503

0.000123503

0.00141778
0.00070889

0.000118843

0.000135277

0.000112212

0.000104655

0.000237903

0.000268684
0.000134342

0.000134342

0.000488224
0.000244112

0.000116891

0.000127221

0.000229284
0.000114642

0.000114642

0.000159375
0.00031875

0.000159375

0.000260362
0.000130181

0.000130181

0.000254972
0.000127486

0.000127486

0.000400647
0.000801294

0.00011448

0.000135547

0.00015062

0.002393427
0.000797809

0.000354109
0.000708218

0.000116478

0.000120942

0.000116689

0.00015994
0.00031988

0.00015994

0.00056752
0.00028376

0.000150148

0.000133612

0.005979339
0.001993113

0.003986226
0.001993113

0.000123521

0.000143869

0.000117255

0.000123777

0.000148119

0.000122239

0.000126126

0.000149934

0.000166662

0.000120626

0.000141923

0.000159917

0.000120275

0.00022887

0.001917363
0.000639121

0.000394498
0.000788996

0.000124485

0.000144553

0.00012546

0.000489246
0.000244623

0.000244623

0.0301367484432
0.09031015232963

5.5034e-05
0.000110068

5.5034e-05

0.0163558392
0.0327116784

8.66747e-05

0.000106021

9.26452e-05

0.000102676

8.45223e-05

0.0158833

9.99458e-05
0.0001998916

9.99458e-05

1.19235e-05
2.3847e-05

1.19235e-05

0.0113599836432
0.02271996728643

0.000101453

0.000106356

0.000109516

0.000140335

0.000106307

0.000101108

0.000108862

0.000110128

0.000156757

0.00016125

8.12497e-05

7.34609e-05

0.000110383

6.73193e-05

9.09881e-05

9.14323e-09

0.000134572

0.000125825

0.000119246

0.000196365

0.000124334

9.69663e-05

8.85085e-05

9.75875e-05

8.35344e-05

9.2875e-05

8.22575e-05

0.000104906

0.000101969

0.000106153

0.000142628

8.62955e-05

8.18177e-05

0.000121338

0.000188713

0.000105755

7.83255e-05

0.00011169

0.000114963

8.14947e-05

8.28976e-05

0.000118452

0.000105246

0.000136551

9.06506e-05

0.000210641

9.11544e-05

7.16491e-05

0.000116625

0.000105998

6.78949e-05

9.72435e-05

0.000108501

7.37506e-05

9.30398e-05

8.56005e-05

8.40342e-05

0.000112478

9.74199e-05

0.000102643

8.09717e-05

0.000120907

0.000144119

0.00016503

0.000127317

0.000102732

9.31987e-05

0.000145764

0.000103682

8.04134e-05

0.000111823

8.60636e-05

0.000101865

8.87514e-05

0.000100699

8.73673e-05

9.44256e-05

0.000146223

0.00012173

7.34375e-05

0.000165654

8.45837e-05

0.000120371

0.000104798

0.000100775

9.4006e-05

0.000165847

9.36685e-05

8.80871e-05

0.00010108

9.29987e-05

7.97409e-05

0.00070401

7.11202e-05

0.000106327

9.22635e-05

0.000179944

0.000101108

0.000115381

0.000175628

8.68718e-05
0.0001737436

8.68718e-05

0.000100093

0.0011630266
0.0005815133

9.31612e-05

9.05406e-05

9.89066e-05

8.88539e-05

0.000108588

0.000101463

0.0011540832
0.0005770416

9.18476e-05

0.000141489

0.00010632

0.000118472

0.000118913

0.000205544
0.000102772

0.000102772

0.000889521
0.0004447605

7.57947e-05

0.000128206

9.89758e-05

0.000141784

3.43491e-05
6.86982e-05

3.43491e-05

0.000443622
0.000221811

0.000221811

0.00010481
0.00020962

0.00010481

0.00063441
0.00190323

0.00126882
0.00063441

0.000133898

0.000114687

0.000138612

0.000133696

0.000113517

0.0108189228
0.0036063076

0.000122233
0.000244466

0.000122233

0.0065706118
0.0032853059

8.56265e-05

8.82667e-05

8.37929e-05

7.46652e-05

0.000180017

9.16948e-05

0.000144549

9.11035e-05

9.18043e-05

0.000108943

0.000100703

8.49393e-05

0.000100746

0.000619804

0.00010339

8.53613e-05

0.00011888

0.000103142

9.64486e-05

9.04434e-05

8.73768e-05

7.75206e-05

0.000154146

8.96054e-05

0.000123904

0.000120912

8.75206e-05

0.0001987687
0.0003975374

0.000101777

9.69917e-05

0.006093222
0.018279666

0.002631584
0.001315792

0.000127898

0.000156595

0.000124163

0.000282079

0.000227811

0.000251627

0.000145619

0.00477743
0.00955486

0.00477743

0.000301872
0.000150936

0.000150936

0.000144309

0.000171257

0.0641521928
0.0161733887

0.000470052
0.000156684

0.000156684
0.000313368

0.000156684

0.0045852535
0.0136457665

0.001858976
0.000929488

0.000135538

0.000123548

0.000137757

0.000137288

0.000151195

0.000115512

0.00012865

0.000109994

0.000438875
0.00087775

0.000112895

0.00032598

0.002078992
0.001039496

0.000251069

0.000145439

0.000138566

0.000117084

0.000106166

0.000146787

0.000134385

0.000277514
0.000138757

0.000138757

0.000894197
0.001788394

0.000152778

0.000112365

0.000106447

0.000108108

0.00012041

0.00015138

0.000142709

0.001028207
0.0005141035

8.9557e-05

8.80568e-05

0.000167934

8.52388e-05

8.33169e-05

0.000520343
0.001040686

0.00014

0.000112333

0.000117038

0.000150972

0.000130746
0.000261492

0.000130746

0.004500429
0.001500143

0.000843802
0.000421901

0.00026945

0.000152451

0.000322874
0.000161437

0.000161437

0.000117595
0.00023519

0.000117595

0.000503819
0.001007638

0.000175017

0.000168542

0.00016026

0.000590782
0.000295391

0.000295391

0.000150311

0.0006157722
0.0002052574

0.0001825828
9.12914e-05

9.12914e-05

0.000113966
0.000227932

0.000113966

0.0092583668
0.0277751004

0.0183215132
0.0091607566

9.0335e-05

0.000105321

9.88097e-05

9.64693e-05

0.000157648

9.90505e-05

0.00771442

0.000111105

0.000116122

0.000239559

0.000136938

9.60613e-05

9.89178e-05

9.76102e-05
0.0001952204

9.76102e-05

0.000559881
0.000186627

0.000186627
0.000373254

0.000186627

0.000138856

0.000119983

0.000242934
0.000121467

0.000121467

0.000210019

0.000371596
0.001486384

0.001114788
0.000371596

0.000124405
0.00024881

0.000124405

0.000137016
0.000274032

0.000137016

0.00022035
0.000110175

0.000110175

0.000156411

0.000259228

0.0067054098
0.0267775384

0.0009033675
0.0026660017

0.0007852674
0.0003926337

0.000239668

4.447e-05

5.22383e-05

5.62574e-05

9.21148e-05
0.0001842296

9.21148e-05

0.0003988444
0.0001994222

9.67332e-05

0.000102689

0.000350192
0.000175096

6.9635e-05

0.000105461

4.41008e-05

0.0058020423
0.0174061269

0.000785062
0.000392531

0.000123465

0.000121847

0.000147219

0.0005083114
0.0002541557

0.000160012

9.41437e-05

0.0103107112
0.0051553556

8.83101e-05

7.67237e-05

8.50122e-05

7.83195e-05

0.000155384

8.17147e-05

7.27125e-05

9.10996e-05

0.000105769

7.74444e-05

7.87551e-05

7.32328e-05

6.56231e-05

8.11303e-05

7.6243e-05

0.000104372

0.000170609

0.000104513

0.000116134

2.64324e-05

0.000123674

8.87501e-05

0.000133662

0.000100313

8.6302e-05

8.11146e-05

0.000108702

8.45293e-05

0.000706844

9.33683e-05

7.88496e-05

8.66849e-05

0.000104444

0.000169809

9.37591e-05

8.1654e-05

0.000120844

8.06161e-05

8.42289e-05

0.000130619

0.00014914

0.00014182

0.000131748

8.27616e-05

9.03287e-05

0.000111255

22.42411903014
5.60952795121

16.81459107893
5.60952795121

0.426368
0.213184

0.0405388

0.0471456

0.0612663

0.0642333

0.00790982
0.01581964

0.00425501

0.00365481

0.00164583
0.00329166

0.00164583

0.03353287651
0.06706575302

8.4651e-07

0.00546981

0.0149069

0.0015098

0.00607463

0.00371507

0.00185582

0.0123132
0.0061566

0.0061566

0.0119088
0.0059544

0.0059544

0.00320028

0.02407185
0.0481437

0.0031269

0.00265935

0.00620207

0.00800973

0.0040738

0.00481332
0.00240666

0.00240666

0.00907462
0.01814924

0.00545975

0.00361487

0.00766745
0.0153349

0.00766745

10.13552658
5.06776329

0.0311244

0.0187608

0.23319

0.0203721

0.0634306

1.74934

0.00259015

0.181084

0.285096

0.672912

0.235739

0.0913315

0.280304

0.157581

0.186057

0.00435646

0.0386321

0.0712377

0.0399692

0.195736

0.0145206

0.00408897

0.243856

0.0345959

0.0187752

0.00355261

0.18953

0.00641864
0.00320932

0.00320932

0.0450671
0.0901342

0.00395018

0.00364076

0.00310922

0.00670041

0.0037976

0.00362803

0.00600545

0.00668713

0.00754832

0.00437033
0.00874066

0.00437033

0.0077645

0.13240398
0.26480796

0.0170494

0.0200538

0.0200079

0.012682

0.00744992

0.0118592

0.00753234

0.00789547

0.00907627

0.00715158

0.0116461

6.6447e-06

6.6447e-06

0.02805892
0.01402946

0.00691445

0.00341057

0.00370444

0.00302135

0.03417518
0.01708759

0.00505712

0.00350701

0.00177806

0.00359115

0.00315425

2.99752e-05

0.629466433949
2.51130066119305

0.001361146
0.004083438

0.000852608
0.000426304

0.000426304

0.000466258
0.000932516

0.000466258

0.000468584
0.000937168

0.000468584

0.09431666705669
0.0316644330189

0.000179591

0.018620468
0.009310234

0.00103496

0.000226996

0.000224876

0.000489611

0.000297127

0.000966491

0.000350062

0.000463132

0.000697517

0.000452034

0.00041494

0.000188216

0.000655611

0.002150173

0.000338268

0.000380534

0.000894639

0.000536732

0.000289085

0.000409403

0.0186841431205
0.03736828624099

0.000171037

0.000154664

0.000146602

0.000171294

0.000157414

0.000151943

0.000183071

0.000146335

0.000156224

0.000190452

0.000148525

0.000253414

0.000279926

0.000207978

0.000178838

0.000184629

0.000147239

0.00018346

0.00015812

0.000577716

0.000223375

0.00015488

0.000153133

1.12049e-09

0.000191993

0.000226525

0.000237029

0.000164411

0.000184243

0.000264128

0.000169098

0.000181625

0.000159802

0.000158487

0.000179222

0.000267735

0.000147038

0.00594946

0.000151363

0.000179873

0.000148465

0.000191581

0.000350824

0.000167493

0.000151476

0.000191122

0.000163296

0.000178728

0.000352784

0.000190493

0.000416078

0.000169675

0.000153553

0.000177902

0.000191251

0.000148634

0.000181775

0.000185628

0.000348122

0.000258812

0.000213532

0.00015711

0.000154615

0.000392547

0.000156345

0.000333816
0.000667632

0.000333816

0.000318493

0.0023861977968
0.0011930988984

0.00041205

5.63996e-08

0.000183207

4.2078e-08

0.000200915

0.000178441

0.000218353

2.08338e-08

1.3587e-08

0.000181213
0.000362426

0.000181213

0.000178548

0.000245116
0.000122558

0.000122558

0.000823024
0.001646048

0.000203263

0.000168119

0.000263724

0.000187918

0.000339714
0.000679428

0.000339714

0.0728331186120001
0.024277706204

0.033944494
0.016972247

0.000193679

0.000185551

0.000154732

0.000796988

0.000162965

0.000221198

0.000174743

0.000153428

0.000164107

0.000160254

0.000163873

0.000254449

0.000184074

0.00015388

0.000169128

0.000153199

0.00018451

0.000236062

0.000174136

0.000395138

0.000182485

0.000159985

0.000219055

0.000197317

0.000164591

0.00014732

0.000392077

0.000199319

0.000149567

0.000158349

0.000158902

0.000187284

0.00027915

0.000167878

0.000187739

0.000143491

0.000197607

0.000320591

0.000515261

0.000150465

0.000146586

0.000205233

0.000190494

0.000168946

0.00224095

0.00224095

0.00017133

0.00250454

0.000299363

0.000214415

0.000159681

0.000118271

0.000262565

0.000175241

0.000206415

0.000357058

0.000212286

0.00015932

0.000160058

0.000169346

0.000135622

0.000402097
0.000804194

0.000402097

0.001240971
0.002481942

0.000242799

0.000176292

0.000193087

0.000207673

0.000205058

0.000216062

0.00033621
0.000168105

0.000168105

0.010762046408
0.005381023204

0.000192329

0.00017643

0.000270242

0.000152835

0.000176259

0.000167346

0.000369236

0.000126492

0.000162197

0.000321008

0.000150273

0.00014963

0.000279549

0.000162214

0.000127615

0.000205718

0.000280299

0.000254255

0.000177032

3.76404e-07

0.000155499

0.000151363

0.000159305

7.68708e-05

0.000156027

0.000153718

0.000174583

0.00011488

0.000177302

0.00016014

0.000113263
0.000226526

0.000113263

0.000192991

0.534384174173
1.59940759391726

0.000399812
0.000199906

0.000199906

0.000989696
0.001979392

0.000399249

0.000590447

0.0003373
0.0006746

0.0003373

0.000444225
0.00088845

0.0002132

0.000231025

0.000230894

0.0001533104
7.66552e-05

7.66552e-05

9.41428e-05
0.0001882856

9.41428e-05

8.36054e-05
0.0001672108

8.36054e-05

0.000195842
0.000391684

0.000195842

0.12741212332
0.06370606166

0.00208767

8.8039e-06

0.00208984

3.19576e-06

0.000269137

0.00571737

0.00210936

0.00209648

0.00228289

0.0119909

0.00217698

0.00215023

0.00213341

0.000271765

0.00796375

0.0105004

0.00212423

0.00772965

0.000147915

0.000181667

0.000173477

0.000192898

0.000163926

0.000268525

0.00015775

0.000986557
0.001973114

0.000173554

0.000161461

0.000295589

0.000168949

0.000187004

0.00025516

7.27503e-05
0.0001455006

7.27503e-05

0.000194258

0.19758158204518
0.0987907910226

0.000127769

0.000176596

0.00018239

0.000139533

0.000119238

0.00014898

0.000148042

0.000178285

0.000150982

0.000299467

0.000154867

0.000181664

0.000172902

0.000143015

0.040441078

0.000155631

0.000135611

0.00016545

0.000209765

0.000164075

0.000166451

0.000200098

0.000183587

0.000185843

0.000170527

0.000150488

0.000169936

0.000184095

0.000168864

0.000170909

0.000196788

0.000156903

0.000173826

0.000189008

0.000169142

0.000160453

0.000145504

0.000146459

0.000192511

0.000160182

0.000167524

0.000164268

0.000142977

0.000169756

0.000151443

0.000182046

0.000141589

0.000175446

0.000190544

0.000186778

0.000156639

0.000526794

0.00648016

0.00014903

0.000194027

0.00014786

0.000160106

0.000163503

0.00015559

0.000156623

0.000164517

0.000168669

0.000154392

0.000153224

0.00154764

0.000156863

0.000190737

0.00041181

0.000139556

0.000194878

0.000162688

0.000187101

0.000153855

0.000177897

0.000185096

0.000157936

0.000159857

0.000185443

0.000159587

0.00167563

0.000149427

0.000188694

0.000158533

0.000167083

0.000146149

0.000186083

0.000170925

0.000148795

0.000145177

0.000173037

0.000168205

0.00532206

0.000711251

0.000166888

0.00016918

0.000190067

0.000144559

0.00015637

0.0030548

0.000167371

0.00016969

0.000161444

0.00573294

0.000331607

0.000141633

0.000164322

0.000162147

0.000167398

0.000150153

0.000188363

0.000165867

0.000151326

0.000147349

0.000165572

0.00015449

0.00015386

0.000181266

0.000175044

0.000154775

0.000189181

0.00181945

0.000148611

0.000175618

0.000143074

0.000171826

0.000180547

0.000206284

0.000204182

0.000172518

0.000160001

3.42658e-09

0.000149772

0.000167471

0.000172425

0.000180958

0.000160711

0.000151975

0.000170737

0.000139703

0.0364949

0.000149591

0.00765299

0.000173475

6.06596e-07

0.000162069

0.00018008

0.000179236

0.000160159

0.00341033

0.000181232

0.000181039

0.000174904

0.000157835

0.000125831

0.000171243

0.000250829

0.00019874

0.000170839

0.49465679584
0.24732839792

0.000443221

0.00344539

0.00045841

0.000516984

0.000450107

0.000731397

0.000483215

0.00325433

0.000682752

0.00312267

0.000654769

0.000187111

0.000459491

0.00195461

0.00146019

0.00207609

0.000717891

0.000475638

0.00227529

0.000446709

0.00241883

0.000716441

0.000450492

0.141922

0.000496786

0.0020389

0.00152542

0.00136768

0.000473286

0.000463309

0.00241262

0.000480895

0.000489482

0.00139496

0.00046161

0.00194039

0.000674703

0.000482874

0.000448631

0.000485576

0.00146312

0.00143284

0.000664368

0.00146712

0.000712193

0.00321161

0.00150223

0.00142641

0.000472431

0.00205462

0.000626945

0.00207012

0.000655489

0.00193892

0.000675905

0.000183431

0.000700403

0.000744455

0.00144053

0.00351405

0.000757264

0.00242426

0.0004864

0.00206272

3.78592e-06

0.00291268

0.00208124

0.000498073

0.0020618

0.00290975

0.000477892

0.000462855

0.000514913

0.000722365

0.000500014

0.00137351

0.000967401

0.000483588

0.000714042

0.000474676

0.000721803

0.000501969

0.00213039

0.00241379

0.000480932

0.00147759

0.00340172

0.00047174

0.000440895

0.000363726
0.000181863

0.000181863

0.00016358

0.0001956556
9.78278e-05

4.36045e-05

5.42233e-05

0.0560307736178
0.0280153868089

0.0272844

4.41384e-07

5.51759e-08

0.000526287

1.06879e-07

0.00020229

1.80637e-06

0.001675492
0.003350984

0.00061315

0.000170129

0.00016538

0.000266376

0.000460457

0.000183113

4.67272e-05
9.34544e-05

4.67272e-05

0.000914554
0.000457277

0.000457277

0.014148547
0.028297094

0.000270071

0.000361486

0.00242355

0.000892865

0.000258085

0.00867403

0.00126846

0.001510096
0.003020192

0.000192014

0.000316722

0.000181406

0.000368491

0.000451463

0.000229481

0.002174835
0.00434967

0.000178433

0.000172429

0.000538579

0.00018736

0.000156726

0.000171359

0.000178471

0.000184584

0.000215771

0.000191123

0.000446053
0.000892106

0.000167731

0.000278322

0.00354362
0.00177181

0.00141626

0.00035555

0.000176978

0.0003275436
0.0006550872

5.1656e-05

5.23199e-05

5.4055e-05

5.18578e-05

5.25799e-05

6.5075e-05

0.000280393
0.000560786

0.000280393

0.001814888
0.003629776

0.00015104

0.000440507

0.000157755

0.000264381

0.000175909

0.000152865

0.000151007

0.00017509

0.000146334

0.0353445468512
0.07068909370237

0.000172887

0.00016799

0.000162107

0.000128338

0.000198298

0.000139581

0.000149682

0.000163269

0.000169325

0.000350113

0.000158619

0.000157538

0.000167913

0.000128319

0.000131397

0.000134398

0.000200498

0.000166167

0.000140922

0.000137752

0.000160366

0.000136077

0.000127983

0.000144283

0.000138785

0.000171918

0.000240315

4.53104e-08

0.000210528

0.0157348

0.000128688

0.000262905

0.000244429

0.000223149

0.00014204

0.00119509

0.000234479

0.000132032

0.000160997

0.000130603

0.000124952

0.000193359

0.000126675

0.000133211

0.000164142

0.00013103

0.000139694

0.000131154

0.000161448

0.000218622

0.000164236

0.000147754

0.000159526

0.000158755

3.54077e-09

0.000140466

0.00330578

0.000161002

0.000162859

0.000129491

0.000131542

0.0012683

0.000186525

0.000161447

0.000128345

0.000168389

0.000141318

0.000171461

0.00016931

0.000962895

0.000161766

0.000134675

0.000166925

0.000242802

0.000142163

0.000134915

0.000133209

0.000129537

0.000200675

0.00018634

0.000745356

0.000177867

0.00018778

0.000234472
0.000468944

0.000234472

0.001346878
0.000673439

0.000481456

0.000191983

0.0001490762
7.45381e-05

7.45381e-05

0.000306896
0.000153448

0.000153448

0.000663082
0.000331541

0.000174468

0.000157073

0.0001749788
8.74894e-05

8.74894e-05

0.000401096
0.000802192

0.000401096

7.0499e-05

0.0170842115095
0.03416842301891

0.010493235

0.000186743

0.000192626

0.000203322

0.000220822

0.000230792

0.000224091

0.000179772

0.000337376

0.000215507

0.000182544

0.000230694

0.000217398

0.000215219

0.000186006

0.000556352

0.000200711

0.000185617

0.00018671

0.00407466

0.000211727

0.000227402

0.000175636

0.000196221

0.000459884

0.000195279

0.000190149

0.00019503

0.000206578

0.000208367

3.54077e-09

0.00072929

0.000206325

0.000181333

0.000200047

3.38871e-08

0.000192953

0.000197075

0.000216725

0.000202394

0.000201006

0.000184099

0.000192069

0.000183713

0.000196694

0.000455665

0.000227389

0.000219317

0.000195963

3.54077e-09

0.000192351

0.000375672

3.54077e-09

0.000182337

0.000241318

0.000326225

0.000196452

0.000413337

0.00022007

0.000261113

0.000212639

0.01776185
0.008880925

0.00025867

0.00648146

0.000174172

0.000377734

0.000167025

0.00016553

0.000280868

0.00041216

0.000168768

0.000394538

0.001849224
0.000924612

0.000166663

0.000181119

0.000160779

0.000416051

8.47149e-05

0.000388514
0.000194257

0.000194257

0.000172952

6.32007e-05

0.000233521

0.013303716
0.004434572

0.002464444
0.004928888

0.000358401

0.000348056

0.000205467

0.000210794

0.000407248

0.000374027

0.000560451

0.001603164
0.003206328

0.000663787

0.000219206

0.000480555

0.000239616

0.000366964
0.000733928

0.000366964

0.000241041
0.000723123

0.000482082
0.000241041

0.000241041

0.000403849
0.000807698

0.000214248

0.000189601

0.0110308505
0.0330925515

0.001150581
0.002301162

0.000375796

0.000266281

0.000264018

0.000244486

0.005029886
0.010059772

0.000430163

0.00119857

0.000355283

0.000184614

0.000424488

0.000177486

0.000161858

0.000157085

0.000855552

0.000187494

0.000171693

0.000197534

0.000180624

0.000169748

0.000177694

0.0010335
0.00051675

0.000266263

0.000250487

0.006937146
0.003468573

0.000323345

0.000219616

0.000185931

0.000318929

0.000337636

0.000912873

0.000396182

0.000393033

0.000381028

0.0008650605
0.001730121

0.000147179

8.45135e-05

0.000182635

0.000238119

0.000212614

0.0603659641581
0.0201219880527

0.0082772701054
0.0041386350527

0.000768981

0.0002594

0.000265071

0.000275521

0.000777709

0.000286759

2.00527e-08

0.00051869

0.000986484

0.00670428
0.01340856

0.000217716

0.000208392

0.000225977

0.000326416

0.000206426

0.000201897

0.000216378

0.000208138

0.000199976

0.000484477

0.00029618

0.000331863

0.000217591

0.000217869

0.000195933

0.000232771

0.00020795

0.000169119

0.000219252

0.000224272

0.00033942

0.000384227

0.000210807

0.00023227

0.000214631

0.000304247

0.000210085

0.00354296
0.00708592

0.000514148

0.000270917

0.000725037

0.000191399

0.00126095

0.000307823

0.000272686

0.005425484
0.002712742

0.000398764

0.000199521

0.00022201

0.000332467

0.000196721

0.000224732

0.000609745

0.000328981

0.000199801

0.000789328
0.000394664

0.000394664

0.003549818
0.001774909

0.00043811

0.000883952

0.000222533

0.000230314

0.001128078
0.000564039

0.000212866

8.7224e-05

0.000263949

0.000579518
0.000289759

0.000289759

0.002707366
0.001353683

0.00103335

0.000320333

0.000101472

0.000116363

0.000390503
0.000781006

0.000221969

0.000168534

8.71612e-05

0.000354904
0.000709808

0.000181685

0.000173219

0.000137833
0.000551332

0.000413499
0.000137833

0.000275666
0.000137833

0.000137833

0.000244648
0.000122324

0.000122324

0.05161201
0.20644804

0.15483603
0.05161201

0.000303823
0.000607646

0.000303823

0.1013408
0.0506704

0.0058223

0.0448481

0.000266809
0.000533618

0.000266809

0.000370978
0.000741956

0.000370978

0.007963388
0.002018167

0.001918034
0.005644822

0.000291077
0.000582154

0.000151055

0.000140022

0.000236112
0.000118056

0.000118056

0.000319591
0.000639182

0.000149433

0.000170158

0.00010928

0.000289418
0.000144709

0.000144709

0.001619148
0.000809574

0.0001411

0.000157169

0.000203557

0.00016502

0.000142728

0.000251494
0.000125747

0.000125747

0.000300399
0.000100133

0.000100133
0.000200266

0.000100133

0.00072354
0.00012059

0.00012059
0.00060295

0.00048236
0.00012059

0.00012059
0.00036177

0.00012059
0.00024118

0.00012059

3.75825e-08

7.93933e-05
0.0004763598

7.93933e-05
0.0003969665

7.93933e-05
0.0003175732

0.0002381799
7.93933e-05

7.93933e-05
0.0001587866

7.93933e-05

1.0426e-07

0.0009826

2.31812e-07

0.000229866

5.5137e-05

1.00724e-08

9.02373e-05

7.55136e-09

0.2050866548
0.6152599644

0.168272
0.336544

0.168272

0.0001517096
7.58548e-05

7.58548e-05

0.0734776
0.0367388

0.0367388

0.27717684
0.13858842

0.00437142

0.134217

0.0047515482
0.0007919247

0.0007919247
0.0039596235

0.0007919247
0.0031676988

0.0023757741
0.0007919247

0.000128893
0.000257786

0.000128893

0.000233778
0.000116889

0.000116889

0.0001976834
9.88417e-05

9.88417e-05

0.000142399
0.000284798

0.000142399

0.000185617
0.000371234

0.000185617

0.000119285
0.00023857

0.000119285

0.001300328
0.007801968

0.001300328
0.00650164

0.005201312
0.001300328

0.001300328
0.003900984

0.000805634
0.000402817

0.000203704

0.000199113

0.000320743
0.000641486

0.000137099

0.000183644

0.000286634
0.000573268

0.000146323

0.000140311

0.000290134
0.000580268

0.000136813

0.000153321

0.007204697161
0.043228182966

0.007204697161
0.036023485805

0.0041349319
0.0165397276

0.0119771907
0.0039923969

0.0039923969
0.0079847938

8.44381e-05

0.000105495

8.99159e-05

9.9476e-05

0.000112481

8.96791e-05

9.17798e-05

0.000138156

0.000118164

0.000103167

0.000206732

0.0001147

7.42254e-05

8.27745e-05

0.000109909

8.02155e-05

0.000129328

0.000195354

0.000127204

0.000147673

8.33696e-05

0.000153016

0.000111474

9.77795e-05

0.000124276

5.97671e-05

0.00010889

0.000127128

9.9228e-05

9.95539e-05

0.000116889

9.67005e-05

0.000193503

0.000145716

7.4239e-05

0.000142535
0.000427605

0.00028507
0.000142535

0.000142535

0.012279061044
0.003069765261

0.003069765261
0.009209295783

0.0002255
0.00011275

0.00011275

0.0017293286
0.0008646643

0.000200841

0.000110728

8.16347e-05

0.000112684

0.000207207

8.53126e-05

6.6257e-05

0.000249336
0.000124668

0.000124668

0.001967682961
0.003935365922

0.000172883

6.86472e-05

9.2895e-05

0.000114482

6.9713e-05

7.9871e-05

1.86059e-07

5.44263e-05

8.04802e-05

0.000100373

0.000152871

5.89558e-05

0.000253798

0.000139575

6.9146e-05

7.44902e-07

8.03282e-05

8.66884e-05

7.62259e-05

0.000215393

0.003521841
0.0198753718

0.000227334
0.000803963

0.000105373
0.000210746

0.000105373

0.000121961
0.000365883

0.000243922
0.000121961

0.000121961

0.000273251
0.001366255

0.000273251
0.001093004

0.000273251
0.000819753

0.000117651
0.000235302

0.000117651

0.0003112
0.0001556

0.0001556

7.99785e-05
0.000159957

7.99785e-05

0.0116122096
0.0024391261

0.0091730835
0.0024391261

0.0067339574
0.0024391261

0.000115992

9.73678e-05

0.0001629778
0.0003259556

8.16308e-05

8.1347e-05

0.0004410912
0.0008821824

8.74502e-05

0.000113781

0.000130237

0.000109623

0.000288902
0.000144451

0.000144451

0.00090061
0.000450305

0.000237834

0.000102245

0.000110226

9.23468e-05

0.0002627134
0.0001313567

8.29996e-05

4.83571e-05

9.02188e-05

0.000106483
0.000212966

0.000106483

0.0001481612
7.40806e-05

7.40806e-05

7.20325e-05

0.000115463

0.00012027
0.00024054

0.00012027

0.000130495
0.00026099

0.000130495

9.41949e-05
0.0001883898

9.41949e-05

0.0005021514
0.0024111462

3.32036e-05

0.0004689478
0.0018757912

0.0003332648
0.0009997944

0.000230328
0.000115164

0.000115164

0.0004362016
0.0002181008

3.05214e-05

0.000100956

8.66234e-05

0.000135683
0.000407049

0.000135683
0.000271366

0.000135683

2.52806e-08

6.01399e-05

0.0040616443
0.0230764005

0.0040616443
0.0190147562

0.0012934653
0.0038803959

0.0008954616
0.0004477308

6.27289e-05

4.64349e-05

5.73851e-05

6.5829e-05

5.19396e-05

5.46345e-05

5.94086e-05

4.93702e-05

0.0001657988
8.28994e-05

8.28994e-05

7.05121e-05
0.0001410242

7.05121e-05

0.000692323
0.001384646

7.37826e-05

7.12311e-05

7.30423e-05

8.2425e-05

0.000391842

0.0082045144
0.0020511286

0.00116477
0.00349431

0.001662179
0.0008310895

3.80423e-05

4.51658e-05

5.00412e-05

4.93947e-05

4.55421e-05

3.82731e-05

3.77166e-05

3.53699e-05

6.58789e-05

5.86373e-05

6.32336e-05

3.53257e-05

6.28735e-05

3.97042e-05

3.53064e-05

8.72863e-05

4.32979e-05

0.0003336805
0.000667361

7.10401e-05

0.000101745

0.000107543

5.33524e-05

0.0026590758
0.0008863586

0.0009184148
0.0004592074

6.30609e-05

6.70599e-05

4.53962e-05

5.50899e-05

9.26413e-05

4.66345e-05

4.43787e-05

4.4946e-05

0.0008543024
0.0004271512

7.64566e-05

6.48308e-05

6.00908e-05

6.30116e-05

0.000103396

5.93654e-05

0.0025738184
0.0006434546

0.0019303638
0.0006434546

0.000283612
0.000567224

8.58922e-05

5.41574e-05

5.38259e-05

8.97365e-05

0.0003598426
0.0007196852

7.22887e-05

6.92058e-05

7.00327e-05

7.35973e-05

7.47181e-05

0.0002943832
7.35958e-05

0.0002207874
7.35958e-05

0.0001471916
7.35958e-05

7.35958e-05

0.001678968
0.00035728

0.00035728
0.001321688

0.000124924
0.000499696

0.000374772
0.000124924

0.000249848
0.000124924

0.000124924

0.000464712
0.000232356

0.000232356

0.0332549502
0.0056620672

0.001482407
0.000439972

7.53283e-05

7.75924e-05

0.000200821
0.000803284

0.000602463
0.000200821

0.000401642
0.000200821

0.000200821

8.62303e-05

0.0052220952
0.026110476

0.0188239292
0.0047059823

0.0103233159
0.0034411053

0.000188762
9.4381e-05

9.4381e-05

9.91057e-05
0.0001982114

9.91057e-05

0.000161706
8.0853e-05

8.0853e-05

8.78934e-05
0.0001757868

8.78934e-05

8.34721e-05
0.0001669442

8.34721e-05

0.0001588029
0.0003176058

6.07048e-05

9.80981e-05

0.000219924
0.000109962

0.000109962

0.0021809
0.00109045

0.00109045

0.0001675386
8.37693e-05

8.37693e-05

0.000588198
0.000294099

0.000107751

0.000186348

0.000202704
0.000101352

0.000101352

0.0001964992
9.82496e-05

9.82496e-05

0.0009571303
0.0019142606

7.74941e-05

0.000113156

9.26087e-05

6.09717e-05

0.000116828

9.59704e-05

6.76741e-05

9.03983e-05

0.000242029

0.00020317
0.000101585

0.000101585

0.000644294
0.001932882

0.00073992
0.00036996

0.000213269

0.000156691

0.000173857
0.000347714

0.000173857

0.000200954
0.000100477

0.000100477

0.000620583
0.001861749

0.000108454
0.000216908

0.000108454

0.000342709
0.000685418

0.000229025

0.000113684

0.00016942
0.00033884

0.00016942

0.0005161129
0.0020644516

0.0015483387
0.0005161129

9.91929e-05
0.0001983858

9.91929e-05

0.0002166
0.0004332

0.000105852

0.000110748

0.00021592
0.00010796

0.00010796

0.00018472
9.236e-05

9.236e-05

0.000422586
0.002535516

0.00211293
0.000422586

0.001690344
0.000422586

0.000206181
0.000618543

0.000412362
0.000206181

0.000206181

0.000649215
0.000216405

0.000216405
0.00043281

0.000216405

0.000350677

0.014864562
0.002477427

0.002477427
0.012387135

0.009909708
0.002477427

0.002477427
0.007432281

0.000152157
0.000304314

0.000152157

0.000887856
0.000443928

0.000128237

0.000177962

0.000137729

0.000942725
0.00188545

0.00012294

0.000219013

0.000199582

0.000165683

0.000134382

0.000101125

0.000652548
0.001305096

0.000138046

0.000285009

0.000116278

0.000113215

0.000572138
0.000286069

0.000125832

0.000160237

34.0126541625
202.283732848405

0.00724528

0.001771873
0.006820279

0.001771873
0.005048406

0.000156103
0.000312206

0.000156103

0.000267213

0.000120169
0.000240338

0.000120169

0.000548204
0.001096408

0.000166413

0.000381791

0.000558606
0.001117212

0.000130544

0.00023048

0.000197582

0.000243156
0.000121578

0.000121578

163.102879582
32.967021911

32.943202688
130.041374333

0.116541072
0.038847024

0.00242322
0.00121161

0.00121161

0.0020856
0.0010428

0.0010428

0.001823366
0.000911683

0.000911683

0.034818056
0.069636112

0.00166237

0.00115394

0.0021752

0.0025299

0.0012233

0.000994475

0.00114442

0.00140867

0.000773676

0.00100935

0.00561778

0.00116098

0.00309332

0.000981947

0.00104718

0.00116236

0.00092203

0.0012874

0.000879841

0.00365464

0.000935277

0.000862875
0.00172575

0.000862875

0.01003083
0.00334361

0.00334361
0.00668722

0.00167431

0.0016693

0.01318854
0.00659427

0.00659427

71.99909898
24.04135043

0.0132625
0.00663125

0.00663125

0.00611509

0.02144176
0.01072088

0.00690907

0.00381181

0.0975003

0.00585742

0.1191909
0.2383818

0.0142845

0.0559528

0.0489536

0.0154795

23.7757773
47.5515546

0.26257

1.4556

0.0110731

0.00549187

0.026764

0.00723245

0.308501

0.00994484

0.00864859

0.0053122

0.0244103

0.00695709

0.284215

0.0407912

0.0325622

0.0595614

0.0137366

0.00581791

0.159354

0.00550728

0.0717309

0.00532821

0.00590713

0.0183969

0.00586512

0.00907855

2.79378

0.350638

0.0303141

0.044355

0.00569887

0.135893

4.71702

0.163461

0.0591935

0.00579719

0.0234361

0.00684451

1.03145

0.11792

0.00727943

0.00580085

0.0139123

1.016

0.00958596

0.00593656

0.0183288

0.0194826

0.00922

0.102703

0.0169139

0.0205453

0.0945021

0.00515475

0.0331962

0.00194982

0.319957

0.0355403

0.0170525

0.719376

0.80073

0.00937768

0.0672867

0.0411816

0.0473815

0.0841421

0.0154613

0.00658226

0.0380148

0.0117361

0.0151091

0.0153906

0.0765605

0.00710588

0.00857222

0.00526794

0.0145559

0.498263

0.0316943

0.0648838

1.6581

2.17522

0.00623537

0.00893173

0.482272

0.0724903

0.732568

0.0099876

0.0273215

0.0899422

0.00579868

1.11316

0.1205

0.242104

0.00515189

0.00617582

0.0324205

0.0228033

0.00695565

0.0054411

0.0738012

0.00552831

0.02339

0.00690459

0.00583937

0.032809

0.0332387

0.0255763

0.0125104

0.00574022

0.0503136

0.00865221

0.00407779
0.00815558

0.00407779

0.00160306

0.04727514
0.01575838

0.00275736
0.00137868

0.00137868

0.0144246
0.0072123

0.00120159

0.00114673

0.00107688

0.00106073

0.00166279

0.00106358

0.0071674
0.0143348

0.00590268

0.00126472

0.450343186
1.351029558

0.44390687
0.88781374

0.011265

0.0032179

0.00041805

0.00280018

0.00294819

0.00368709

0.00278265

0.0200257

0.00357904

0.0125116

0.0112886

0.0101707

0.0169708

0.00279343

0.00345233

0.00256668

0.00708106

0.00182964

0.00309735

0.00314129

0.00505089

0.00330506

0.00535287

0.00783268

0.0184434

0.00414957

0.00272963

0.00334484

0.00477059

0.00266078

0.00341299

0.00703996

0.00422914

0.00946028

0.00457075

0.0024496

0.00496885

0.0038959

0.00299064

0.00865595

0.00279901

0.00304006

0.00303286

0.00572484

0.0051513

0.00311799

0.00258908

0.00325012

0.00411648

0.00421807

0.00282992

0.00272896

0.00237206

0.00303943

0.00233759

0.00476182

0.0044866

0.00552321

0.0026253

0.00244004

0.00250437

0.00291438

0.00279737

0.00423399

0.00481127

0.00407063

0.0246701

0.0026604

0.00368188

0.00482057

0.0071118

0.00267889

0.00338443

0.00459664

0.00313577

0.00474173

0.00514674

0.0142985

0.00398942

0.00229465

0.00863058

0.00237742

0.00492035

0.00235484

0.00344257

0.00851119

0.00595468
0.01190936

0.0031136

0.00284108

0.000481636
0.000963272

0.000481636

0.015032612
0.035412796

0.010695144
0.005347572

0.000891734

0.00360561

0.000850228

0.00161475

0.00235239

0.00193786

0.00163369

0.00214635

0.00412601

0.000758612

0.000960713

0.03896727
0.01298909

0.00789132
0.00394566

0.0019043

0.00204136

0.01808686
0.00904343

0.00216006

0.00219942

0.00256684

0.00211711

0.00234706
0.00117353

0.00117353

0.001891857
0.000630619

0.000630619
0.001261238

0.000630619

0.000785626

0.00181744

0.000525647
0.001051294

0.000525647

0.00417268

0.00094576

0.38486484
0.12828828

0.04897251
0.09794502

0.00225181

0.0282017

0.018519

0.07931577
0.15863154

0.00832878

0.00244258

0.00294027

0.00247918

0.00239183

0.00947595

0.00202502

0.00203812

0.00565063

0.00555403

0.00244854

0.00268109

0.00359957

0.00260253

0.0037048

0.00271051

0.00233201

0.00273268

0.00246354

0.00207495

0.00243159

0.00352292

0.00268465

0.00213582

22.986747425
8.162514135

0.00182459

0.00322114
0.00161057

0.00161057

0.00176586

0.00312281

0.00146675
0.0029335

0.00146675

0.00169068

0.00567202

0.0026377
0.0052754

0.0026377

13.29378192
6.64689096

0.00302419

0.00296719

0.0119499

0.00301688

0.0027176

0.0143086

0.00988223

0.0134231

0.146289

0.00488452

0.0145545

0.499675

0.00143988

0.0128149

0.00507597

1.48719

0.0154744

0.004319

0.00382547

0.0481815

1.98169

0.0079858

2.33532

0.00424117

0.0034592

0.00918096

0.00358974
0.00179487

0.00179487

1.48351

0.00220339
0.00440678

0.00220339

0.00158824

0.00162078

0.00184925
0.000924625

0.000924625

0.00256145
0.0051229

0.00256145

0.00325768
0.00162884

0.00162884

0.002546802
0.000848934

0.000848934
0.001697868

0.000848934

0.06410706
0.02189182

0.0015684

0.03421996
0.01710998

0.00275241

0.00181534

0.00179583

0.00179199

0.00284187

0.00194353

0.00174322

0.00242579

0.00321344
0.00642688

0.00164883

0.00156461

0.0257654

0.094483338
0.023819223

0.003121868
0.00857205

0.00164959
0.000824795

0.000824795

0.000793554

0.003007038
0.001503519

0.000712784

0.000790735

0.025101684
0.008367228

0.004005112
0.002002556

0.00125226

0.000750296

0.000803523
0.001607046

0.000803523

0.000754319
0.001508638

0.000754319

0.001767935
0.00353587

0.000922595

0.00084534

0.001962422
0.000981211

0.000981211

0.004115368
0.002057684

0.00122074

0.000836944

0.035007855
0.011669285

0.00133103
0.00266206

0.00133103

0.001684946
0.000842473

0.000842473

0.00198487
0.00396974

0.00198487

0.00291042
0.00145521

0.00145521

0.00131736
0.00263472

0.00131736

0.00134058
0.00268116

0.00134058

0.001716465
0.00343293

0.000904052

0.000812413

0.001681297
0.003362594

0.000864825

0.000816472

0.001982526
0.000660842

0.001321684
0.000660842

0.000660842

0.086543129
0.017526356

0.069016773
0.017526356

0.051490417
0.017526356

0.000225397
0.000450794

0.000225397

0.002557202
0.001278601

0.000287308

0.000322486

0.000362648

0.000306159

0.000321747

0.000587476
0.000293738

0.000293738

0.000286191
0.000572382

0.000286191

0.014080676
0.007040338

0.000335217

0.000376314

0.000350697

0.000372359

0.000272812

0.000339497

0.000397339

0.000359098

0.000349039

0.000309999

0.000449151

0.000353701

0.000335323

0.000349097

0.000352527

0.000328911

0.000400447

0.000319946

0.000321279

0.000367585

0.000547316
0.000273658

0.000273658

0.00069451
0.00138902

0.000228652

0.00023137

0.000234488

0.001517807
0.003035614

0.000338391

0.00029079

0.000475089

0.000413537

0.000264403
0.000528806

0.000264403

0.000491123
0.000982246

0.000255565

0.000235558

0.000215602

0.000213164
0.000426328

0.000213164

0.000261144
0.000522288

0.000261144

0.000714388
0.000357194

0.000357194

0.000593484
0.001186968

0.000326893

0.000266591

0.001140194
0.000570097

0.000294087

0.00027601

0.000540292
0.000270146

0.000270146

0.000253855

0.000788342
0.001576684

0.000276168

0.000237864

0.00027431

0.0005278
0.0010556

0.0005278

0.000297447

0.000490568
0.000981136

0.000490568

0.919284940832
4.59065267045999

0.000229852

0.918677355832
3.670760144628

0.0001603934
8.01967e-05

8.01967e-05

0.000272723

0.0008312858
0.0024938574

0.0016625716
0.0008312858

0.000107284

9.33696e-05

0.000120049

0.000101857

9.49294e-05

9.28008e-05

0.000117097

0.000103899

0.914188628332
2.739799978996

0.000582294
0.000291147

0.000291147

0.000721947
0.001443894

0.000264547

0.00022031

0.00023709

0.001832064
0.000916032

0.000916032

0.003829802
0.001914901

0.000224199

0.000249442

0.000250498

0.000223998

0.000236494

0.000237149

0.000274396

0.000218725

0.010612396
0.005306198

0.000308709

0.000255012

0.00162527

0.000281628

0.000523723

0.000295431

0.000380544

0.000704736

0.000316206

0.000614939

0.000567328
0.000283664

0.000283664

0.001834624
0.003669248

0.000229976

0.000221446

0.000245842

0.000217027

0.000250008

0.000218995

0.000232981

0.000218349

0.001965202
0.000982601

0.000247675

0.000470636

0.00026429

0.00050319
0.000251595

0.000251595

0.000661538
0.000330769

0.000330769

0.000294683
0.000589366

0.000294683

0.004798154
0.009596308

0.000346934

0.00025737

0.000255114

0.000235522

0.000280822

0.000291652

0.000226775

0.000398216

0.000228255

0.000243481

0.000233491

0.000189321

0.000219461

0.000237531

0.000209011

0.000235061

0.000274601

0.000219332

0.000216204

0.000459119

0.000296758

0.011042066
0.005521033

0.000231547

0.000436039

0.000229338

0.000166444

0.000250845

0.000253501

0.000345756

0.000216719

0.000274515

0.000234257

0.000319404

0.000278349

0.000380188

0.000235857

0.000300731

0.000649886

0.000364975

0.000352682

0.000469756
0.000939512

0.000469756

0.001764934
0.000882467

0.000297129

0.000272959

0.000312379

0.0024997
0.00124985

0.000375488

0.000590731

0.000283631

0.000672484
0.000336242

0.000336242

0.000913928
0.001827856

0.000293114

0.000305875

0.000314939

0.000539334
0.001078668

0.000262454

0.00027688

0.000434823
0.000869646

0.00020188

0.000232943

0.00899805
0.004499025

0.0016854

0.000270117

0.000293882

0.00030107

0.000423241

0.000403146

0.000259678

0.000286961

0.000335688

0.000239842

0.000248686
0.000497372

0.000248686

0.001424314
0.002848628

0.000213625

0.00018446

0.000239671

0.000193431

0.000200237

0.00019277

0.00020012

0.00360748
0.00180374

0.000285214

0.000325031

0.000281196

0.000321792

0.000300174

0.000290333

0.00352567
0.001762835

0.000396361

0.000277527

0.00027955

0.000261106

0.00028103

0.000267261

0.000205672

1.465158702
0.732579351

0.000864222

0.0174952

0.00978064

0.132549

0.00264431

0.010009

0.00936871

0.0241905

0.0259307

0.00120904

0.00253587

0.00420715

0.000907079

0.00966242

0.0765777

0.0100784

0.00951167

0.0148145

0.208368

0.0237213

0.00176441

0.0260295

0.0111281

0.00923496

0.0673416

0.00955867

0.0130967

0.000468024
0.000234012

0.000234012

0.0018988872
0.0009494436

0.00022243

0.000234081

0.000227088

3.80896e-05

0.000227755

0.000578492
0.000289246

0.000289246

0.000433852
0.000867704

0.000433852

0.003707896
0.001853948

0.000332927

0.000304879

0.000294729

0.000279684

0.000337736

0.000303993

0.00024433
0.00048866

0.00024433

0.001910748
0.000955374

0.000327199

0.000325408

0.000302767

0.001819527
0.003639054

0.000291089

0.000460274

0.000239892

0.000287663

0.000321742

0.000218867

0.00147606
0.00073803

0.000404126

0.000333904

0.00298782
0.00149391

0.000255388

0.000313852

0.00025412

0.000196707

0.000253751

0.000220092

0.000357521

0.003122943
0.006245886

0.000294776

0.000273914

0.000266998

0.000282049

0.000397896

0.000287677

0.00039667

0.000276963

0.000359366

0.000286634

0.00119294
0.00238588

0.000297418

0.000268636

0.000299832

0.000327054

0.072142611464
0.036071305732

0.000249435

0.000235962

0.001198

0.000200494

0.000439717

0.00025326

0.000294349

0.000222877

0.000211748

0.000390061

0.000204095

0.00021351

0.000195024

0.000212233

0.000260256

0.000193013

0.000260123

0.000480601

0.000196635

0.000547346

0.000246809

0.00022635

0.000195808

0.000236022

0.000196668

0.000606288

0.000246547

0.000238263

0.000394606

0.000401941

0.000291565

0.000262136

0.00029097

0.000467348

0.000191103

0.000258035

0.000204606

0.000315169

0.000208776

0.000215686

0.000295492

0.000210616

0.000227563

0.000347047

0.000219289

0.000224034

0.000213203

0.000231797

0.00035537

0.000212752

0.000325966

0.000406441

0.000394243

0.00019931

0.00030272

0.000313013

0.000338639

0.000219636

0.000248285

0.000287593

0.000252451

0.000291261

0.000257058

0.000698379

0.000303442

0.000433948

0.000214556

0.000214632

0.000223642

0.000241129

0.000241402

0.000217989

0.000221288

0.000333613

0.000290664

0.000200546

0.000219809

0.000198285

0.000358985

0.000188446

0.000260635

0.0001868

0.000225249

0.000211146

0.000402918

0.000244555

0.000231261

0.000322347

0.000200057

1.32732e-07

0.00017961

0.000220652

0.00021429

0.000350788

0.000255967

0.000272684

0.000217209

0.000251144

0.000294088

0.000288522

0.000240065

0.000238104

0.000418021

0.000339059

0.000250765

0.000191044

0.00023335

0.000193216

0.000226265

0.000232421

0.00023883

0.000182535

0.000387032

0.000253153

0.000330109

0.000208498

0.000213405

0.000212006

0.000235531

0.00030787

0.000225884

0.000210998

0.000520089

0.000204969

0.000211418

0.000199437

0.000224448

0.000467879

0.000220729

0.000296152

0.022974808
0.011487404

0.000436378

0.000672826

0.000308532

0.00129583

0.00120947

0.00639811

0.000267773

0.000626989

0.000271496

0.005138568
0.002569284

0.000225156

0.000253316

0.000252161

0.000447019

0.000263055

0.000300274

0.000268945

0.00022689

0.000332468

0.000232486
0.000464972

0.000232486

0.000725762
0.000362881

0.000362881

0.002000298
0.001000149

0.000237976

0.000280576

0.000244679

0.000236918

0.000335608
0.000671216

0.000335608

0.000586095
0.00117219

0.000586095

0.00504641
0.01009282

0.000241117

0.000253261

0.000272328

0.000220974

0.000263777

0.000296298

0.00025657

0.000251161

0.000226208

0.000390607

0.000266042

0.000251043

0.000283332

0.000404578

0.000207426

0.000262545

0.000250411

0.000227573

0.000221159

0.000721312
0.000360656

0.000360656

0.000231443
0.000462886

0.000231443

0.001679788
0.000839894

0.000839894

0.000748988
0.000374494

0.000374494

0.006738408
0.003369204

0.000284042

0.000293823

0.000253794

0.000330683

0.000374249

0.000272513

0.000352987

0.000269273

0.000294286

0.000379336

0.000264218

0.000480245
0.00096049

0.000234953

0.000245292

0.001441833
0.002883666

0.00021657

0.000256904

0.000262719

0.000457343

0.000248297

0.000231944
0.000463888

0.000231944

0.000339517

0.00067941
0.00135882

0.00067941

0.000372455
0.00074491

0.000372455

0.000319643

0.000509856
0.000254928

0.000254928

0.000317351
0.000634702

0.000317351

0.001113598
0.000556799

0.000271064

0.000285735

0.012090758
0.006045379

0.000228206

0.000225295

0.000231703

0.000240784

0.000345315

0.000221853

0.000224698

0.000399429

0.000220875

0.000238197

0.000263304

0.000234445

0.000260308

0.000198852

0.000344463

0.000240331

0.000266043

0.000273678

0.000238042

0.000204641

0.00022404

0.000286702

0.000194112

0.000240063

0.007857824
0.003928912

0.000256994

0.000416177

0.00032185

0.000233817

0.000679411

0.000271068

0.000252434

0.00022192

0.000245779

0.000295969

0.000229312

0.00027576

0.000228421

0.00290241
0.001451205

0.000280206

0.000415345

0.000266589

0.000259161

0.000229904

0.002258925
0.00451785

0.000634368

0.000481507

0.00114305

0.000453988
0.000907976

0.000218034

0.000235954

0.000514184
0.000257092

0.000257092

0.00027273
0.00054546

0.00027273

0.0003317
0.0006634

0.0003317

0.000693268
0.001386536

0.000227335

0.000229668

0.000236265

0.00044602
0.00022301

0.00022301

0.002130342
0.001065171

0.000253235

0.000251839

0.000256903

0.000303194

0.002139198
0.001069599

0.000217312

0.000203441

0.000205141

0.000215672

0.000228033

0.001373123
0.002746246

0.000268476

0.0002774

0.000262302

0.000256144

0.000308801

0.000584628
0.000292314

0.000292314

0.001267764
0.000633882

0.000316346

0.000317536

0.00058601
0.00117202

0.000314996

0.000271014

0.001910696
0.003821392

0.000214453

0.000227746

0.000248629

0.000220333

0.000215406

0.000271855

0.000221181

0.000291093

0.000230772

0.000350973
0.000701946

0.000350973

0.000254693
0.000509386

0.000254693

0.000546002
0.001092004

0.000292558

0.000253444

0.000281692

0.000275212

0.005080892
0.002540446

0.000261354

0.000213493

0.000214202

0.000232691

0.000220281

0.000242823

0.000255775

0.000514708

0.000385119

0.002752192
0.001376096

0.000202363

0.000235498

0.000260823

0.000283368

0.000200899

0.000193145

0.000284045
0.00056809

0.000284045

0.062181932
0.031090966

0.000212878

0.000236612

0.000254974

0.00024671

0.000212732

0.000299429

0.000292761

0.000231338

0.000188986

0.000199362

0.000281459

0.00021199

0.000188297

0.000254806

0.000214381

0.000239022

0.000278643

0.000240716

0.000258141

0.000257655

0.000397938

0.000212865

0.000218635

0.000200481

0.000210842

0.000405542

0.00019948

0.000184643

0.000281977

0.000210177

0.000238247

0.000403339

0.000249866

0.000203739

0.000341232

0.00023827

0.000200448

0.000284486

0.000246014

0.000273212

0.000203895

0.000473727

0.000201915

0.000251136

0.000318135

0.000263143

0.00023709

0.000223147

0.000250773

0.000321562

0.000342338

0.000206647

0.000259268

0.000222709

0.000218139

0.000225309

0.000247206

0.000274981

0.000318147

0.000217932

0.000246328

0.000238564

0.000278306

0.000281145

0.000272415

0.000300224

0.000199825

0.000222608

0.000288607

0.000210695

0.000212128

0.000361393

0.000194044

0.000358609

0.000881952

0.000174265

0.00025165

0.000224559

0.000400831

0.000321803

0.000231721

0.000207151

0.000219042

0.000251959

0.000243349

0.000232858

0.000254998

0.000196546

0.000270256

0.000191014

0.000312184

0.000232281

0.000261862

0.000221039

0.000261693

0.000257634

0.000218965

0.000228694

0.000203753

0.000180027

0.000242959

0.000257726

0.000230034

0.000207175

0.000243723

0.000250424

0.000199956

0.000243455

0.000314345

0.000309117

0.000182422

0.000230624

0.000214126

0.00028032

0.000272651

0.00133166

0.000203748

0.000686091
0.001372182

0.000224452

0.000217699

0.00024394

0.000324873
0.000649746

0.000324873

0.000253084
0.000126542

0.000126542

0.002016602
0.006049806

0.001264034
0.000632017

0.000306464

0.000325553

0.000358366
0.000716732

0.000358366

0.000623908
0.001247816

0.000332456

0.000291452

0.000804622
0.000402311

0.000402311

0.000215594

0.000423619
0.001270857

0.000847238
0.000423619

0.000423619

0.000554817
0.000184939

0.000184939
0.000369878

0.000184939

0.001011678
0.000337226

0.000337226
0.000674452

0.000337226

0.000176084

0.000201649

0.195807693
0.039672873

0.15613482
0.039672873

0.000190417

0.000167766

0.000340691

0.038973999
0.115763073

0.000434724
0.000869448

0.000434724

0.00038018
0.00076036

0.00038018

0.000865793
0.001731586

0.00045062

0.000415173

0.000399741

0.000339121

0.002001088
0.001000544

0.000365303

0.000303993

0.000331248

0.007002838
0.014005676

0.000336615

0.000348654

0.000310254

0.000326856

0.000348096

0.00033142

0.000309462

0.000321595

0.000339001

0.000379073

0.000387301

0.000373167

0.000545625

0.00031696

0.000367883

0.0003078

0.00042704

0.000311034

0.000298732

0.00031627

0.001739652
0.003479304

0.000348824

0.000487018

0.000385165

0.000518645

0.000610042
0.000305021

0.000305021

0.011579854
0.023159708

0.000384341

0.000388378

0.000394408

0.000586316

0.000385233

0.000457321

0.000385589

0.000415684

0.000373969

0.000519472

0.000306233

0.000344959

0.00038265

0.000412068

0.000373332

0.000446945

0.000490126

0.000344059

0.00035464

0.000390743

0.000438491

0.000316563

0.000340642

0.000418453

0.000402948

0.000371532

0.000381682

0.000306551

0.000466526

0.000420062

0.014506469
0.029012938

0.000378967

0.000278235

0.000316005

0.000364076

0.000365034

0.000286497

0.000292197

0.00027084

0.000315011

0.000294672

0.000242886

0.000674133

0.000298484

0.000303722

0.000301089

0.000156829

0.000279461

0.000279101

0.000541339

0.000304464

0.000290054

0.000356934

0.000314438

0.000292687

0.000251143

0.00028978

0.000322407

0.000341204

0.000279974

0.000278354

0.000296672

0.000281713

0.000265344

0.000291106

0.000292081

0.00033641

0.000313993

0.000322552

0.000523045

0.000354182

0.000348679

0.000286796

0.000430144

0.000327547

0.000276188

0.052020768689
0.258004948445

0.205984179756
0.052020768689

0.010561119
0.031683357

0.000248251
0.000496502

0.000248251

0.000584149
0.001168298

0.000268493

0.000315656

0.000539798
0.001079596

0.000244816

0.000294982

0.000243359
0.000486718

0.000243359

0.005811237
0.011622474

0.000241312

0.000206541

0.000420351

0.00044361

0.000241416

0.000228654

0.000242878

0.00021953

0.000264471

0.000279493

0.000348344

0.000295877

0.000262833

0.000258932

0.000255023

0.000228512

0.000275652

0.000270883

0.000260684

0.000273161

0.00029308

0.001582858
0.000791429

0.000264208

0.00029353

0.000233691

0.000299617
0.000599234

0.000299617

0.000490678
0.000245339

0.000245339

0.000556536
0.000278268

0.000278268

0.000463624
0.000231812

0.000231812

0.00257572
0.00128786

0.000282232

0.000252702

0.000249886

0.00027073

0.00023231

0.002138807967
0.000712935989

0.000403268989
0.000806537978

0.000402513

7.55989e-07

0.000309667
0.000619334

0.000309667

0.00038154
0.00114462

0.000163479
0.000326958

0.000163479

0.000436122
0.000218061

0.00010354

0.000114521

0.0403266621
0.0134422207

0.001458432
0.002916864

0.000279491

0.000291447

0.000265879

0.000320214

0.000301401

0.007897178
0.003948589

0.000291148

0.00027576

0.000293206

0.000539215

0.000320856

0.000327767

0.000273512

0.000429094

0.000297996

0.000331486

0.000268948

0.000299601

0.0072033777
0.0144067554

0.00028162

0.000243579

0.000263726

0.000240048

0.000326808

0.000257261

0.000297601

0.000281798

0.0003565

0.000265291

0.000294455

0.000248362

0.000296526

9.85797e-05

0.000241158

0.000313159

0.000241353

0.000255247

0.000208786

0.000212071

0.000229764

0.000300631

0.000272461

0.000210989

0.000266473

0.00042188

0.000277251

0.001148544
0.000574272

0.000273976

0.000300296

0.0005151
0.00025755

0.00025755

0.000356608
0.001069824

0.000356608
0.000713216

0.000356608

0.000550967
0.001652901

0.000583212
0.000291606

0.000291606

0.000259361
0.000518722

0.000259361

0.000848649
0.000282883

0.000282883
0.000565766

0.000282883

0.000319865
0.000959595

0.00063973
0.000319865

0.000319865

0.000325653

0.000340901
0.000681802

0.000340901

0.049608216
0.016536072

0.000412045
0.00082409

0.000412045

0.000781388
0.000390694

0.000390694

0.000673834
0.000336917

0.000336917

0.000543751
0.001087502

0.000348317

0.000195434

0.001385368
0.000692684

0.000428828

0.000263856

0.000491295
0.00098259

0.000491295

0.000539748
0.000269874

0.000269874

0.002041288
0.001020644

0.000350017

0.000312713

0.000357914

0.000510634
0.000255317

0.000255317

0.000669362
0.000334681

0.000334681

0.000597076
0.000298538

0.000298538

0.000265584
0.000531168

0.000265584

0.006693654
0.003346827

0.000273578

0.000296193

0.00032277

0.00033423

0.000295351

0.00031869

0.000324932

0.000291098

0.000260408

0.000350422

0.000279155

0.003509874
0.001754937

0.000314249

0.000313676

0.000357611

0.00034951

0.000419891

0.000664374
0.001328748

0.000397696

0.000266678

0.000629346
0.001258692

0.000369424

0.000259922

0.001601404
0.000800702

0.000390551

0.000410151

0.006646378
0.003323189

0.000292151

0.000302083

0.000259108

0.000269857

0.000246242

0.000269917

0.000268277

0.000291714

0.00031634

0.000270082

0.000284477

0.000252941

0.000331085
0.00066217

0.000331085

0.000747176
0.000373588

0.000373588

0.000635014
0.000317507

0.000317507

0.000793542
0.000396771

0.000396771

0.000743922
0.002231766

0.000743922
0.001487844

0.000422445

0.000321477

0.006751804
0.019863002

0.000320639
0.000641278

0.000320639

0.001715744
0.000857872

0.000284583

0.000313018

0.000260271

0.00173742
0.00086871

0.000329112

0.000539598

0.000621618
0.000310809

0.000310809

0.000344834
0.000689668

0.000344834

0.00039241

0.000790211
0.001580422

0.000268347

0.000240183

0.000281681

0.000616906
0.000308453

0.000308453

0.001044756
0.002089512

0.000233698

0.000230189

0.000266594

0.000314275

0.00302622
0.00151311

0.000374501

0.000404794

0.000362117

0.000371698

0.00378602

0.000196198

0.00031237

0.003815572
0.018830516

0.015014944
0.003815572

0.002267655
0.000755885

0.000279388
0.000558776

0.000279388

0.000952994
0.000476497

0.000476497

0.006691926
0.00231309

0.002065746
0.004131492

0.000259675

0.000282007

0.0002366

0.000291897

0.000308613

0.000282446

0.000404508

0.000247344

0.000746597
0.002239791

0.000358901
0.000717802

0.000358901

0.000387696
0.000775392

0.000387696

0.0121137956
0.002056177

0.0100576186
0.002056177

0.004181468
0.0010897356

2.59478e-05

2.28708e-05

0.0030029952
0.0010009984

0.0010009984
0.0020019968

2.54519e-05

0.00013189

3.65385e-05

0.000140877

5.69174e-05

3.99005e-05

2.65572e-05

2.8798e-05

0.000261084

3.34233e-05

3.07278e-05

6.69286e-05

2.64645e-05

6.71584e-05

2.82813e-05

3.99186e-05

0.0009664414
0.0038199736

0.0006455626
0.0018908958

0.000568541
0.0002842705

6.67243e-05

6.01104e-05

5.30518e-05

5.28957e-05

5.14883e-05

0.0001450969
0.0002901938

4.26977e-05

4.91604e-05

5.32388e-05

4.5792e-05

0.0001704032
0.0003408064

0.000117483

5.29202e-05

0.0003584391
0.0001194797

0.0001242484
6.21242e-05

6.21242e-05

0.000114711
5.73555e-05

5.73555e-05

0.0001822383
6.07461e-05

6.07461e-05
0.0001214922

6.07461e-05

7.8129e-05
2.6043e-05

5.2086e-05
2.6043e-05

2.6043e-05

0.00011461
0.00034383

0.00022922
0.00011461

0.00011461

0.0189428598
0.0031571433

0.0157857165
0.0031571433

0.0031571433
0.0126285732

0.0094714299
0.0031571433

0.00026581
0.000132905

0.000132905

9.85732e-05
0.0001971464

9.85732e-05

0.000531679
0.001063358

0.000220466

0.000157276

0.000153937

0.000132489
0.000264978

0.000132489

0.000480791
0.000961582

0.000322044

0.000158747

0.000244578
0.000489156

0.000126291

0.000118287

0.0003821884
0.0001910942

9.78308e-05

9.32634e-05

0.000503548
0.000251774

0.000140967

0.000110807

0.000619512
0.000309756

0.00015533

0.000154426

0.000453896
0.000907792

0.000203325

0.000250571

0.0006592158
0.0003296079

6.78555e-05

8.92189e-05

9.49667e-05

7.75668e-05

0.00013412
0.00040236

0.00013412
0.00026824

0.00013412

1.004102317
6.024613902

1.004102317
5.020511585

1.004102317
4.016409268

0.165331619
0.495994857

0.000871728
0.000435864

0.000435864

0.001424994
0.000712497

0.000712497

0.003452362
0.006904724

0.000524102

0.000305111

0.00157467

0.000454598

0.000593881

0.000645439
0.001290878

0.000337217

0.000308222

0.000656094
0.000328047

0.000328047

0.31951482
0.15975741

0.0053906

0.00342602

0.00556478

0.0253268

0.0563794

0.0130772

0.006456

0.00584885

0.00328994

0.00480585

0.00821741

0.0180013

0.00397326

2.516312094
0.838770698

0.00037614
0.00075228

0.00037614

0.836455919
1.672911838

0.00152607

0.00999046

0.0148776

0.015169

0.0261062

0.0137322

0.0049361

0.121787

0.000408698

0.00113168

0.525883

0.00326414

0.00104487

0.00111584

0.00173982

0.00124893

0.00928055

0.0445619

0.000805193

0.00156517

0.0109883

0.000648478

0.0147182

0.00992652

0.00088861
0.000444305

0.000444305

0.001494334
0.002988668

0.000407862

0.000330147

0.000430854

0.000325471

0.000253432
0.000126716

0.000126716

9.47236e-05
0.0001894472

9.47236e-05

0.00026779

0.000103831

0.00135864
0.00022644

0.00022644
0.0011322

0.00022644
0.00090576

0.00022644
0.00067932

0.00045288
0.00022644

0.00022644

0.000148273
0.000444819

0.000148273
0.000296546

0.000148273

0.3402062859
0.0640923833

0.2761139026
0.0640923833

0.0002893808
7.23452e-05

7.23452e-05
0.0002170356

7.23452e-05
0.0001446904

7.23452e-05

0.0700883688
0.0175220922

0.0163482333
0.0490446999

0.000308564
0.000617128

0.00014255

0.000166014

0.000783388
0.000391694

0.000129588

0.000262106

0.000206714
0.000103357

0.000103357

0.00056431
0.00112862

0.000235423

0.000151848

0.000177039

0.0279794098
0.0139897049

0.000625746

0.000246466

0.000686293

0.000369024

0.000496919

0.00135399

0.000198388

0.000185134

0.000924647

0.000560739

0.000128988

0.000766918

6.36283e-05

0.00109696

0.000246006

0.000194517

0.000328358

0.000126555

0.000234843

0.000469901

0.00202852

6.39836e-05

0.000180516

0.00221302

0.000199645

0.0004061784
0.0008123568

7.3406e-05

9.05024e-05

0.000115595

0.000126675

0.000584425
0.00116885

0.00020385

0.000121039

0.000107528

0.000152008

0.0011738589
0.0035215767

0.0011793974
0.0005896987

4.61224e-05

3.29099e-05

4.31863e-05

3.72614e-05

9.93556e-05

4.14223e-05

3.693e-05

6.94952e-05

9.11944e-05

4.54543e-05

4.63669e-05

0.0011683204
0.0005841602

0.000104328

7.51448e-05

3.603e-05

4.81984e-05

3.73149e-05

3.60865e-05

9.16902e-05

4.6691e-05

3.57126e-05

3.77627e-05

3.52011e-05

0.008599728
0.002149932

0.002149932
0.006449796

0.004299864
0.002149932

0.000177315

0.000115206

0.000112977

0.000847588

0.000112764

0.000107116

0.000276595

0.000287096

0.000113275

0.0443480139
0.1330440417

0.0440605839
0.0881211678

7.61609e-05

0.00014636

7.712e-05

8.38616e-05

0.000151223

0.000536736

0.000167758

6.30494e-05

6.25018e-05

7.22346e-05

0.000105216

6.98542e-05

7.31756e-05

5.62536e-05

5.68746e-05

6.44803e-05

0.000104434

6.5396e-05

6.92619e-05

6.08279e-05

5.5813e-05

7.35892e-05

0.000117102

5.83983e-05

0.00131598

6.2145e-05

0.000155733

7.0337e-05

0.000124322

5.72493e-05

7.05488e-05

6.25396e-05

6.79646e-05

8.31694e-05

6.62619e-05

0.000488383

7.85882e-05

6.21313e-05

7.24863e-05

7.74743e-05

5.85997e-05

0.0369541

6.17016e-05

0.000141861

6.01019e-05

0.000552863

7.08183e-05

7.31635e-05

7.77442e-05

0.00011461

6.34917e-05

6.36349e-05

7.39511e-05

6.06976e-05

6.0678e-05

7.03306e-05

5.91839e-05

6.00573e-05

0.000118175
0.00023635

0.000118175

0.000169255
0.00033851

0.000169255

0.0011824962
0.0001970827

0.0001970827
0.0009854135

0.0001970827
0.0007883308

0.0005912481
0.0001970827

0.0003941654
0.0001970827

0.0001228

7.42827e-05

5.39627e-07

0.000909788

0.0002215569
7.38523e-05

0.0001477046
7.38523e-05

7.38523e-05

29.7913861443
177.909141954313

0.000745275
0.000149055

0.000149055
0.00059622

0.000149055
0.000447165

0.00029811
0.000149055

0.000149055

0.000504023

0.000690186
0.000345093

0.000345093

1.00801958
0.202735798

0.805283782
0.202735798

0.202735798
0.602547984

0.000453713

0.000586563

0.018567644
0.009283822

0.000208028

0.0002053

0.00803947

0.000187602

0.000643422

0.000260694
0.000521388

0.000260694

0.000322639
0.000645278

0.000322639

0.00223508

0.000590923

0.000320334
0.000640668

0.000320334

0.001653568
0.000826784

0.00041578

0.000185199

0.000225805

0.000266275

0.000700008
0.000350004

0.000350004

0.000678916
0.000339458

0.000339458

0.001416293
0.002832586

0.000495267

0.000456903

0.000464123

0.000302514

0.005505886
0.002752943

0.000208899

0.000251854

0.000263527

0.000335188

0.000986187

0.000469507

0.000237781

0.16342681
0.32685362

0.16223

0.00119681

0.001508416
0.000754208

0.000754208

0.000457689

0.002449594
0.001224797

0.000197444

0.000298248

0.000729105

0.000418061
0.000836122

0.000418061

0.00424282
0.00848564

0.00424282

0.002317594
0.001158797

0.000728037

0.000168639

0.000262121

0.00033192
0.00066384

0.00033192

0.00139794
0.00279588

0.00139794

0.000842784
0.000421392

0.000421392

0.000429034
0.000858068

0.000429034

0.000927284
0.000463642

0.000463642

0.000842638
0.000421319

0.000421319

0.000465891

0.001687918
0.003375836

0.000471766

0.000306784

0.000377655

0.000242261

0.000289452

0.001957898
0.000978949

0.000633572

0.000345377

0.000300762

0.00384581
0.00769162

0.00384581

90.4962074891895
18.1745271981

0.000141132

69.5296822303998
17.3825902261

0.23017968
0.69053904

0.4233988
0.2116994

0.120404

0.0912954

0.008444583
0.016889166

0.000359602

0.000411876

0.000298061

0.0010302

0.00082463

0.000310027

0.000612264

0.000255902

0.000486488

0.000310071

0.000270841

0.00029722

0.000281757

0.000616636

0.000329674

0.00142648

0.000322854

0.006528606
0.003264303

0.00146712

0.00085406

0.000943123

0.004435465
0.00887093

0.00057605

0.00135278

0.000958476

0.000376098

0.000710772

0.000461289

0.000495184
0.000990368

0.000495184

0.00212528
0.00106264

0.00106264

0.00155621
0.000778105

0.000778105

0.11444152
0.34332456

0.106765422
0.213530844

0.000580591

0.000215716

0.000335935

0.000119987

0.000157294

0.000150526

0.000326285

0.000244203

0.000305545

0.000123933

0.000173574

0.000160265

0.000264789

0.000272368

0.000154806

0.00015327

0.00140326

0.000124191

0.000517797

0.000309484

0.000248289

0.000175574

0.000123828

0.000249355

0.000657424

0.00263108

0.000250482

0.000263285

0.000171402

0.00034756

0.000198217

0.000245926

0.000288246

0.000203968

0.000234257

0.000161373

0.000205512

0.000138264

0.000208784

0.000337961

0.000680306

0.000198676

0.000135161

0.000187421

0.000359143

0.000202198

0.000312591

0.000206374

0.000176535

0.000167295

0.000595217

0.000128762

0.000635782

0.000483691

0.000177194

0.000186707

0.000476889

0.000250922

0.000260258

0.000201026

0.000327933

0.000216271

0.000793662

0.000157854

0.000182303

0.0002637

0.00039475

0.000250187

0.000489285

0.000187447

0.000482361

0.000224283

0.00014921

0.000113249

0.000187587

0.000222354

0.000150761

0.000183971

0.000409678

0.000222676

0.000628541

0.00018109

0.00213143

0.0001704

0.00823649

0.000192245

0.000152609

0.000276797

0.000297123

0.000383745

0.000161511

0.000283575

0.00036061

0.000151821

0.00077844

0.000183615

0.000238637

0.000176725

0.000226932

0.000851765

0.000319029

0.000292988

0.0001847

0.000183923

0.000195589

0.000136056

0.000294719

0.000503157

0.000158234

0.00038401

0.000183321

0.000195461

0.000698965

0.000148067

0.000376854

0.00017883

0.000165617

0.000151765

0.000414648

0.000182684

0.000229585

0.000230278

0.000396571

0.000177606

0.000172657

0.000166169

0.00875479

0.000240651

0.000166358

0.000235196

0.000463589

0.000576407

0.000305389

0.000247071

0.000388569

0.000502827

0.000180225

0.000361217

0.000203237

0.000346515

0.000217462

0.000465557

0.000204769

0.00133548

0.00015844

0.000368249

0.000259405

0.000279998

0.000300839

0.0002034

0.000401216

0.000525523

0.000201645

0.000494294

0.000450988

0.00015252

0.000305144

0.000145994

0.000121103

0.000129262

0.000173306

0.000281812

0.000185984

0.019184498

0.000188905

0.000170709

0.000182806

0.0165118

0.000160407

0.000185387

0.000358277

0.000176093

0.000169469

0.000893953

0.000186692

0.000559014

0.00026458

0.000277067

0.000229208

0.000203973

0.000368488

0.000494355

0.000165468

0.000813126

0.000348765

0.00097217

0.000164176

0.000234432

0.000120654

0.000258388

0.000157026

0.000254389

0.000463518

0.000123157

0.00031709

0.000164637

0.00032014

0.000227569

0.000494915

0.000144186

0.000166484

0.000475689

0.000217112

0.00026103

0.000238885

0.000182157

0.00013716

0.000251448

0.000432303

0.000226233

0.000178452

0.000243278

0.00021168

0.000311514

0.000610215

0.000797846

0.000177979

0.000280272

0.000184303

0.000244168

0.000270862

0.000133247

0.000669719

0.000159015

0.000217619

0.000198012

0.000362584

0.000329196

0.000158562

0.000459259

0.000637807

0.000557657

0.000378118

0.000192896

0.000232509

0.000308412

0.00016046

0.001480468
0.000740234

0.000740234

0.006935864
0.013871728

0.000330955

0.000358634

0.00428509

0.000266665

0.000194037

0.000188888

0.000163524

0.000483325

0.000494945

0.000169801

0.000234264

0.223599573
0.670798719

0.000515208
0.001030416

0.000515208

0.216799598
0.433599196

0.000632939

0.000458222

0.000421225

0.000495603

0.000458645

0.00166593

0.000477603

0.000320185

0.00235068

0.000440122

0.000740365

0.00107391

0.000431941

0.00047841

0.000402077

0.000647556

0.000418713

0.000411368

0.00035864

0.000539914

0.000473538

0.0373733

0.000452806

0.000491686

0.000344899

0.000516632

0.00120714

0.000503863

0.000464654

0.000471586

0.000459535

0.00046664

0.000275128

0.000325606

0.00030669

0.000649738

0.00048668

0.00051443

0.000355528

0.000846082

0.00130269

0.000477695

0.000467256

0.000431442

0.000339842

0.000595862

0.000682592

0.000498073

0.000499385

0.000541746

0.000718354

0.000358058

0.000483901

0.000492313

0.000462772

0.000511671

0.000315065

0.00105929

0.000586489

0.000749365

0.000445285

0.000530702

0.000829565

0.000440092

0.000468453

0.000483916

0.000448678

0.000352619

0.000427876

0.000798333

0.000495707

0.000446716

0.000449862

0.000489192

0.024771

0.000327991

0.000529911

0.000547357

0.000360337

0.000352496

0.000433308

0.000426242

0.000343993

0.000465779

0.00075341

0.000453718

0.000890713

0.000346975

0.00149546

0.000496471

0.000451456

0.000479821

0.000801567

0.00103322

0.000396126

0.000894066

0.000663439

0.000741065

0.000899682

0.000295893

0.000439335

0.000338635

0.000371442

0.00117848

0.0011472

0.000584007

0.00082325

0.0860995

0.000387428

0.000318203

0.00110384

0.000482672

0.000544318

0.000344073

0.00112734

0.000320092

0.00103234

0.00036664

0.000696045

0.000552196

0.000769182
0.000384591

0.000384591

0.004940764
0.002470382

0.000486952

0.000389581

0.000728007

0.000405287

0.000460555

0.004540686
0.002270343

0.000310112

0.00148229

0.000477941

0.000605299
0.001210598

0.000605299

0.001108304
0.000554152

0.000554152

0.016470078
0.049200088

0.009186903
0.018373806

0.000135016

0.000175377

0.000216519

0.000393668

0.00435932

0.000191401

0.000391802

0.000204688

0.00206079

0.000434762

0.000154506

0.000469054

0.004691534
0.009383068

0.000138707

0.000423986

0.000190119

0.000281347

0.000347843

0.000289694

0.000203355

0.000203068

0.000246688

0.000169638

0.00132673

0.000239653

0.000169659

0.000461047

0.001294272
0.002588544

0.00100226

0.000292012

0.000210146

0.002174446
0.001087223

0.000269298

0.000143948

0.000168648

0.000193925

0.000150546

0.000160858

16.5305679071
49.5917037213

0.00200737
0.00401474

0.00102713

0.00098024

0.04081323
0.020406615

0.00247693

0.00657197

0.000695049

0.00166941

0.00092192

0.00276371

0.00188953

0.00164339

0.000759086

0.00101562

16.5081539221
33.0163078442

0.0064209

0.057868

0.0192613

0.00622062

0.00988158

0.466735

0.00228562

0.817494

2.31121e-05

0.00239583

0.00279802

0.13239

0.0502961

0.00230979

0.00313575

0.0557627

0.0640633

1.54437

0.00764213

0.00398509

0.00978144

0.0350943

0.0751548

0.00300901

1.88313

0.00238099

0.00347972

0.00695118

0.0275195

0.138346

0.134107

0.0842012

0.00205475

0.1149356

0.0218831

0.0432132

0.0498393

0.139776

0.0166939

0.00264122

0.401392

0.00523431

0.00253211

0.00725322

0.00309081

0.0622555

0.00236601

0.00205002

0.070946

0.0398129

0.00255869

0.00275259

0.118197

0.00388663

0.032967

0.0069721

0.0360008

0.0421046

0.00246516

0.0306039

0.0806095

0.0879991

0.00330617

0.0327236

0.0451326

0.00721658

0.0029289

0.00257628

0.0481302

0.0349478

0.30990051

0.299987

0.00991351

0.00634333

0.00642838

0.00234511

0.0560037

0.00273723

0.0072528

0.020703

0.00245063

1.26463

0.00211817

0.00632342

0.167026

0.00695485

0.124055

0.00255828

0.0024239

0.0608993

0.132339

0.014193

0.00591556

0.00540379

0.00616761

0.0825591

0.0499353

0.0374664

0.0805364

0.0527073

0.0189964

0.0137557

0.0036206

0.00627655

0.0725494

0.037687

0.00224833

0.00676528

0.0358206

0.0798331

0.00664832

0.260891

0.0719767

0.00240552

0.0022169

0.00211742

0.0485831

0.00259161

0.0454304

0.166732

0.00191861

0.0067073

0.0319254

0.0645166

0.00628273

0.214846

0.150995

0.0527832

0.0699624

0.0508403

0.0573155

0.00334283

0.00278888

0.003965

0.0394622

0.00410619

0.0601009

0.0720844

0.0660666

0.00593421

0.0769114

0.0613354

0.12376

0.00210836

0.00543231

0.132397

0.00233032

0.245209

0.00368012

0.0438914

0.0277471

0.00377815

0.0655135

0.00252373

0.0531535

0.00560638

0.0024514

0.0737815

0.0508476

0.0728785

0.00276262

0.0376343

0.00202576

0.986028

0.00522383

0.00466255

0.0594045

0.0296463

0.00572463

0.00615368

0.00389075

0.131323

0.0666652

0.00361607

0.00914199

0.0633781

0.0060057

0.00257579

0.0702936

0.00290591

0.00639732

0.00555189

0.00316506

0.00867925

0.00313089

0.0038282

0.0306931

0.0425099

0.362805

0.0512684

0.00256019

0.00608072

0.00288316

0.00634087

0.141387

0.00263951

0.0650197

0.143008

0.0311637

0.00275747

0.0167969

0.00287485

0.0212294

0.156877

0.0542591

0.0646668

0.0818161

0.0530404

0.00199937

0.0451691

0.00512182

0.00342244

0.489676

0.0483517

0.179331

0.00242155

0.0692338

0.138691

0.00622093

0.0636116

0.801291612000001
0.267097204

0.000464301
0.000928602

0.000464301

0.003920834
0.001960417

0.000236124

0.000233101

0.000264817

0.000246871

0.000221017

0.00022632

0.000271014

0.000261153

0.000376416
0.000188208

0.000188208

0.002606758
0.001303379

0.000349061

0.000407857

0.000546461

0.00160953
0.000804765

0.000804765

0.001136878
0.000568439

0.000568439

0.001416327
0.002832654

0.000825009

0.000591318

0.000511168
0.000255584

0.000255584

0.000439379
0.000878758

0.000439379

0.000761948
0.000380974

0.000380974

0.000341708
0.000170854

0.000170854

0.016187518
0.008093759

0.000446476

0.000390708

0.000415804

0.000866669

0.000376057

0.000512936

0.0003695

0.000431496

0.000391403

0.000276621

0.000517791

0.00030685

0.000329906

0.000485933

0.00039272

0.000494019

0.00108887

0.000381229
0.000762458

0.000381229

0.24526808
0.49053616

0.0584472

0.0400898

0.00149858

0.0348427

0.0342173

0.0344326

0.0417399

0.004771795
0.00954359

0.000684553

0.000418192

0.000395499

0.000228682

0.000758735

0.000559838

0.000629999

0.000293589

0.00039681

0.000405898

0.000747398
0.000373699

0.000373699

0.00051203
0.000256015

0.000256015

0.791795840007
2.79185692868999

0.015422462984
0.046267388952

0.000856203
0.001712406

0.000135295

0.000154605

0.000144205

0.00012802

0.000294078

0.001121691
0.002243382

0.000179984

0.000392815

0.00041025

0.000138642

0.002894256084
0.005788512168

0.000181998

0.000133698

0.000125396

0.000129323

0.000143692

0.000127974

0.000144001

0.000201181

0.000192736

0.000131274

2.04384e-07

0.00015268

9.13017e-05

0.000140159

0.000141894

0.000132865

0.000351769

0.000150726

0.000221384

0.0018747298
0.0009373649

0.000236226

0.000197143

4.94599e-05

0.000281545

0.000172991

0.001939876
0.000969938

0.000147202

0.000102519

0.000164461

0.000165574

0.00013456

0.000255622

0.00107081
0.000535405

0.000107176

0.000116257

0.000311972

0.000361938
0.000723876

0.000190212

0.000171726

0.000960568
0.000480284

0.000480284

0.000263611
0.000527222

0.000263611

0.00184894
0.00092447

0.000246301

0.000150854

0.000162224

0.000188747

0.000176344

0.000150153
0.000300306

0.000150153

0.00087833
0.000439165

0.000208734

0.000230431

0.000846564
0.001693128

0.000157493

0.00014162

0.000129488

0.000130364

0.000146992

0.000140607

0.00928284
0.00464142

0.000173616

0.000143738

0.000111547

0.000128883

0.000113033

0.000118347

0.000158942

0.000125186

0.000129435

0.00011689

0.000131925

0.000179302

0.000160612

0.000116764

0.000148941

0.000219739

0.000142973

0.000125265

0.000268682

0.000138252

0.000118466

0.000109424

0.000118171

0.000118654

0.000110608

0.000272897

0.000113574

0.000144486

0.000109695

0.000118163

0.000130111

0.000112758

0.000112341

0.0096920175
0.0032306725

0.000545662
0.000272831

0.00014692

0.000125911

0.004355325
0.0021776625

0.000112013

8.22115e-05

0.000158081

0.000196055

0.000123104

0.000122784

0.0002671

0.000136875

0.000103303

0.000108627

0.000132682

0.000176414

0.000101982

0.000130187

0.000107936

0.000118308

0.000501689
0.001003378

0.000150697

0.000186952

0.00016404

0.00027849
0.00055698

0.00015322

0.00012527

0.000158321
0.000474963

0.000158321
0.000316642

0.000158321

0.439582595934
0.146527531978

0.000228817
0.000457634

0.000228817

0.141693873078
0.283387746156

0.000104948

0.000426795

0.000221521

9.40023e-05

0.00024042

0.000125754

0.000353194

9.73796e-05

0.000111901

0.000222583

0.000127392

0.000124787

8.50406e-05

0.000117839

9.52467e-05

9.52125e-05

9.65925e-05

0.000109357

0.000137507

9.80692e-05

1.37937e-06

0.000202149

0.000117594

0.000131037

0.000339042

0.000108945

0.000535691

0.000211815

9.04821e-05

0.000118672

0.000129681

0.000201204

0.000170418

0.000175988

0.000102748

9.61625e-05

0.000126546

8.40749e-05

9.07094e-05

0.000168333

0.00011636

8.38985e-05

0.000127855

0.000755499

0.000112758

9.96373e-05

9.26611e-05

0.000117344

0.000120112

8.835e-05

0.000494368

0.000106557

9.67e-05

0.000121838

0.00100727

9.04478e-05

8.99234e-05

0.000122479

0.000219521

0.000125838

9.65623e-05

9.6651e-05

9.15443e-05

0.000135833

9.60331e-05

0.060314

0.000102692

9.32571e-05

0.00027192

0.000413878

0.00106815

0.000266451

0.000164733

0.000123576

0.000137941

0.000123481

0.000129753

0.00010968

9.42497e-05

0.000115514

0.000108968

0.000127872

9.6693e-05

0.000218817

0.000127245

0.000149142

0.000260007

0.000103905

0.000104812

0.000126949

0.000101033

0.000614664

0.000105916

0.000124636

3.95818e-07

0.000203307

0.000109416

0.000107854

0.000123279

0.000218885

0.000130001

0.00101984

0.000229076

0.000632328

0.000603509

0.000101081

0.000115049

9.35798e-05

0.00010549

0.000111714

0.000134445

0.000284597

0.000105081

0.000100369

0.000117186

9.50014e-05

0.000110588

0.000109516

0.000107884

9.85872e-05

9.47253e-05

0.00116805

9.29885e-05

0.000181396

0.000107371

0.000121571

0.000122748

0.000628108

0.000112286

0.000773634

0.00103262

0.000124107

8.25132e-05

0.00010392

0.000186163

0.000108735

0.00012199

9.54112e-05

0.00019051

0.000279053

0.000131618

0.000333872

7.94563e-05

9.65204e-05

0.000249967

9.72773e-05

0.000106616

0.000118979

0.000194216

9.7146e-05

0.000200048

9.72255e-05

0.000101409

0.00028211

0.00173524

9.09393e-05

9.63951e-05

9.8579e-05

0.000124847

9.36993e-05

8.68565e-05

0.000107852

0.000253838

0.000103969

0.00010984

0.000108632

0.000103486

0.000114751

9.35576e-05

0.000175789

0.000186885

0.000124488

9.02285e-05

0.000109088

0.000302801

0.000113744

0.000122009

0.000120806

0.00017667

0.000111467

0.000138097

0.000188162

0.000115088

0.000123037

0.000115621

0.000133511

0.000192027

0.00172623

0.000118576

0.000128396

0.000242108

8.46566e-05

0.000113579

0.000108617

0.000102542

0.000110744

8.67528e-05

9.62979e-05

0.000101234

9.3739e-05

0.000105776

0.000114289

0.00022488

1.43339e-06

0.000100542

0.000113812

9.13448e-05

0.000118013

0.000103062

0.00011723

0.00010381

9.39988e-05

0.000134966

9.8366e-05

0.0001258

0.000119164

0.000192604

0.000221929

0.000297384

0.000125208

0.000145529

0.000126066

0.000113574

0.000117894

0.000124313

0.000105637

0.000113782

0.000114685

0.000109504

0.00013381

0.00874214

0.00024177

0.0263235

8.43786e-05

0.000172803

0.000193727

0.000314449

9.16385e-05

0.000112354

8.74516e-05

0.000108453

0.000125505

8.91114e-05

0.000129027

0.000113572

0.000116587

9.72529e-05

0.000530079

9.44693e-05

0.000119385

0.000120872

0.000117167

0.000120448

0.000131954

0.000126619

0.000101311

0.000227362

0.0034822018
0.0017411009

0.000163193

0.000123957

0.00018829

9.51709e-05

0.000264792

0.000126417

0.000204817

0.000123989

0.000175612

0.000274863

0.002796418
0.001398209

0.000966863

0.000206056

0.00022529

0.000371436
0.000185718

0.000185718

0.000708597
0.001417194

0.000114698

0.000224353

0.000127555

0.00012694

0.000115051

0.000571217
0.001142434

0.00011351

0.000182488

0.000142673

0.000132546

0.000149373

0.0063582098475
0.012716419695

0.000193328

0.000524126

0.00016581

0.000197477

0.000175757

9.58475e-08

0.000104865

0.00017255

0.00017152

0.000122887

0.000352764

0.000145163

0.000262197

0.00014684

0.000158477

0.000194292

0.000159422

0.000314207

0.000143124

0.000329073

0.000138567

0.000168355

0.000287249

0.000134395

0.000111639

0.000122587

0.000168605

0.000178379

0.000186329

0.000137771

0.000162136

0.000172405

0.000150366

0.000205452

0.012861921
0.004287307

0.00036495
0.0007299

0.00015893

0.00020602

0.000188032
0.000376064

0.000188032

0.00027842
0.00013921

0.00013921

0.00056855
0.000284275

0.000116856

0.000167419

0.000132115
0.00026423

0.000132115

0.000276496
0.000138248

0.000138248

0.000445502
0.000222751

0.000222751

0.000271583
0.000543166

0.000144095

0.000127488

0.001216573
0.002433146

0.00015662

0.000173772

0.000133134

0.000156173

0.000182429

0.00013992

0.000274525

0.000521326
0.000260663

0.00013582

0.000124843

0.000506801
0.001013602

0.000506801

0.000322538
0.000645076

0.000120882

0.000201656

0.000111594
0.000223188

0.000111594

0.000127974
0.000255948

0.000127974

0.03726015
0.012489007

0.000206871

0.01149293
0.02298586

0.000242297

0.000239463

0.00601677

0.000207934

0.000490921

0.000489996

0.000201082

0.000293284

0.000200329

0.000371039

0.000375857

0.00049786

0.00021781

0.000313687

0.000383864

0.000203215

0.00047183

0.000275692

0.001578412
0.000789206

0.00024577

0.000543436

0.000146155

0.1593503026
0.0531814962

0.000194186

0.000137336
0.000274672

0.000137336

0.002869002
0.001434501

0.000380323

0.000146545

0.000202179

0.000202896

0.000182831

0.000189838

0.000129889

0.000673628
0.000336814

0.000336814

0.000663704
0.000331852

0.000182472

0.00014938

0.00445173
0.00890346

0.000276538

0.000143192

0.000251804

0.000144144

0.000174527

0.000169898

0.000146122

0.000121928

0.000282843

0.000138617

0.000124273

0.000154671

0.000149182

0.000412341

0.000750573

0.000161098

0.000193961

0.000159146

0.000276597

0.000220275

0.088807194
0.044403597

0.000214336

0.000133026

0.000123953

0.000138189

0.000155292

0.000149068

0.000133171

0.000105439

0.00011976

0.000143767

0.000133524

0.000114585

0.000112944

0.000137455

0.000140497

0.00024816

0.000129818

0.000119356

0.000322083

9.98939e-05

0.000142756

9.0857e-05

9.62623e-05

0.000187584

0.00011341

0.00014074

0.000134244

0.000186977

0.000156171

0.00019661

0.000152732

0.000139207

0.000136361

0.000156715

0.000182329

0.000162382

0.000142887

0.00017306

0.000149981

0.000130146

0.000107305

0.00011214

0.000187569

0.000142806

0.000119418

0.000142037

0.000138838

0.000108675

0.00015327

0.000317678

0.000227354

0.000141312

0.000152525

0.000122246

0.000110713

0.000169483

0.000114129

0.000111799

0.000130157

0.000134863

0.000169495

0.000196618

0.000405688

0.000125891

0.000127886

0.00255868

0.000195607

0.000145734

9.5335e-05

0.00010999

0.000135711

0.000159028

0.000111392

0.000132242

0.000135104

0.000122717

0.000130333

0.000157965

0.000139787

0.000137125

0.000103832

0.000119438

0.000116919

0.000152557

0.000139362

0.000102634

0.000114888

0.000102631

0.000113011

0.000221806

0.000131637

0.000310961

0.000115555

0.000160558

0.000135879

0.000149133

0.000123794

0.000133462

0.000137221

0.000125181

0.0002823

0.000209004

0.000227654

0.000130889

0.000102349

0.000105024

0.000140469

0.000142887

0.000138684

0.000175197

0.000262041

0.000156763

0.000143345

0.000139046

0.000132261

0.000156186

0.000439502

0.000197331

0.000126767

0.000114278

0.000195886

0.000142537

0.000151703

0.000139252

0.000602044

0.000142358

0.000113885

0.000133009

0.000187735

0.000127718

0.000142771

0.000143717

0.00011011

0.000147789

0.0003012

0.000134377

0.000129926

0.000141707

0.000150239

0.000118048

0.000147088

0.000156378

0.000104315

0.000148934

0.000165137

0.000188716

0.000662599

0.000144812

0.000175088

0.00014999

0.00014028

0.000168081

0.000142876

0.000116852

0.000264027

0.000133262

0.000101062

0.000160689

0.000127621

0.00014159

0.000140282

0.000144374

0.000293718

0.00015178

0.000205347

0.000146403

0.000135649

0.000202042

0.000127956

0.000136959

0.000136243

0.000130163

0.000140813

0.00012253

0.000136697

0.000140285

0.000126708

0.000143839

0.000106233

0.00016067

0.000140471

0.000126592

0.000131868

0.000141171

0.00014789

0.000146173

0.000161383

0.000125035

0.000160054

0.000102276

0.000114151

0.000135425

0.000380051

0.000140299

0.000239752

0.000148719

0.00013766

0.000218833

0.000198459

0.000158576

9.95082e-05

0.000165453

0.000118718

0.000130644

0.000156091

8.14652e-05

0.000175541

0.000116425

0.000144638

0.000143364

0.000135009

0.000143107

0.000116169

0.000247433

0.000158476

0.00014762

0.00312732

0.000173551

0.000126919

9.73394e-05

0.000103524

0.000116722

0.000110027

0.000161612

0.00012156

0.00181619

0.0001356

0.000114749

0.000133681

0.000151258

0.000150305

0.000105549

0.000180963

0.000117651

0.000216292

0.000354731

0.000894697

0.0016159524
0.0008079762

0.000243524

0.000315809

9.95052e-05

0.000149138

0.001083504
0.002167008

0.000123195

0.000112332

0.000122635

0.000133001

0.000184129

0.000135114

0.000133001

0.000140097

0.000138261

0.000372652
0.000745304

0.000186585

0.000186067

0.000141959

0.000761138
0.000380569

0.000380569

0.542914171002
0.181207383497

0.001300356
0.000650178

0.000167029

0.000150066

0.00015981

0.000173273

0.00081219
0.00162438

0.000176459

0.000155423

0.000170451

0.000156711

0.000153146

0.133813129109
0.267626258217

0.000198752

0.000103514

0.00013023

0.000146435

0.000157108

0.000278874

0.000150165

0.000131334

0.000133242

0.000254254

0.000166969

0.000120826

0.000114882

0.000125131

0.000172899

0.000104774

0.000469657

0.000152837

9.56921e-05

0.000373502

9.53026e-08

0.000106315

0.000171513

0.000172673

0.000189811

0.000160388

0.000105152

0.000159116

0.000118046

0.000144457

0.000119261

0.000166636

0.000127814

0.000318511

0.000146559

0.000238084

0.000130192

0.000153019

0.000189541

0.000254983

9.85414e-05

9.62511e-05

0.00012729

0.000124112

0.000230123

0.00021644

0.000174679

0.000137913

0.000114302

0.000135076

0.000352294

0.000146743

0.000127144

0.000165567

0.000132841

0.00055699

0.000153581

0.000160846

0.000122097

0.000194037

0.000124793

9.09536e-05

0.000193638

0.000156417

0.000168659

0.000126505

0.000122714

0.000156469

0.000133539

0.00029771

9.36214e-05

9.26231e-05

0.000143032

0.000155374

0.000160054

0.000147153

0.000222947

0.000264323

0.000173244

0.000164129

0.000245688

0.000121162

0.000160343

0.000176203

9.45263e-05

0.0001048

0.00014214

0.000174148

0.000322845

0.00011324

0.000127389

0.000127051

0.000186704

0.000134069

0.000108437

0.000116649

0.000211612

7.82253e-05

0.000105335

0.000180668

0.000100014

0.00011637

0.00021803

0.000276599

0.00025217

0.000177312

0.000125994

0.000182529

0.000165629

0.00010275

0.000173552

0.000114713

0.000110433

0.000111472

0.000126427

0.000137647

9.09964e-05

0.000261966

0.000125582

9.1654e-05

0.000107105

0.000108798

0.000183251

9.56494e-05

0.000101493

9.63815e-05

0.00010992

0.000125602

0.000593453

0.00013005

0.000138807

0.000140659

0.000133231

9.65349e-05

0.00011181

0.000121544

8.36595e-05

0.000140954

0.000135543

8.90921e-05

0.000196271

0.000194705

0.000126267

0.000140032

0.000123783

0.000146024

0.00011042

0.000172007

0.00013237

0.000138386

0.00010992

0.000167941

0.000139092

0.0142002205

0.000193295

0.00532272

0.000544082

0.000472702

0.000132259

0.000182754

3.50595e-05

0.000224028

0.000130047

0.000194226

0.0039638

0.000143991

0.000825696

0.000163283

0.000118209

0.000971128

0.000582941

0.000139036

0.00011093

0.000138054

0.000160598

0.000154067

0.000178372

0.000120346

0.000168083

0.00014097

0.000105061

0.000211715

0.000103426

0.00011713

0.000165518

0.00027557

0.000111247

0.000147582

0.000171495

0.00033467

9.88582e-05

0.000125176

0.000160137

0.000136241

0.000143964

0.000595276

0.000122312

0.000115716

0.000140641

0.000146162

0.000460659

0.000148165

0.000111099

0.000263809

0.000119379

0.000161939

0.000139713

0.000148275

9.35219e-05

0.000158621

0.000200964

0.000165967

0.000193767

0.000248231

0.000142409

0.000103487

0.000212036

0.000260177

0.000414316

9.66688e-05

0.000160309

9.5889e-05

0.000304726

0.00013034

8.89576e-05

0.000119737

0.000126844

8.90468e-05

0.0002969

0.000137266

0.000347525

0.0001397

0.000135681

0.000129006

0.000211967

0.000163222

0.000181468

0.000104461

0.000153281

0.000206099

0.000108294

0.000195893

0.000144803

0.000153776

0.000127001

0.000113759

0.000152035

0.000125296

0.000246024

0.000149427

0.000154834

0.000414927

0.000252999

0.000361731

0.000118268

0.000169955

0.000141407

0.000241161

0.000287955

9.5339e-05

0.000163589

0.000145311

0.0001111

0.000138924

0.000133964

0.000109515

0.03275

0.00010259

0.000141218

0.000122494

0.00016172

0.000217762

0.000155946

0.000122128

9.92898e-05

0.000171926

9.62145e-05

0.000165136

0.000137566

0.000175667

0.000107222

0.000109512

0.000138481

0.000100128

0.000153426

1.38442e-08

0.000145324

0.000126769

0.000143229

0.000102755

0.000251903

0.000254977

0.000206234

0.00013156

0.000143166

0.000135325

0.000187332

0.00100949

0.000147365

0.000132716

0.000122336

0.000120472

0.000162976

0.000120695

0.000119415

8.21366e-05

0.000120079

0.000318333

9.64396e-05

0.000160977

0.000129141

0.000119752

0.00013046

8.796e-05

0.000115732

0.000185528

0.000178009

0.000132737

8.94858e-05

0.00011266

0.000136282

0.000157098

0.000147412

0.000180431

0.0001691

0.000135718

0.000178476

9.3861e-05

0.000145258

0.000126887

0.000592739

0.000105165

0.000188136

0.00014459

0.000798996

0.000235394

8.91419e-05

0.000109848

0.000185659

0.000166483

0.000116865

0.00013837

0.00015164

0.000326031

0.000168583

0.000204055

0.000136673

0.000138422

0.000210327

0.000191222

0.00019069

0.000145605

0.000111511

0.000236218

0.000105682

0.000174406

8.91419e-05

0.00015364

0.000149291

0.000170143

0.000401172

8.8971e-05

0.000120545

0.000248466

0.000156399

8.84239e-05

0.000186614

0.000219336

0.000152638

0.000400128

0.000153657

0.000137004

0.000199256

0.000165411

0.000109672

0.000216739

0.000140152

0.000163173

0.000138167

0.000240449

0.00019391

0.000145621

0.000135442

0.000157106

0.000117588

0.000171649

0.000104266

2.74052e-05

0.00016079

0.000113056

0.000108466

0.000145346

0.000150099

0.000100243

9.22426e-05

0.000113746

0.000131279

9.9992e-05

0.000189988

0.000214406

0.000126947

0.000125131

0.000167677

0.000189516

0.000237763

9.37112e-05

9.95816e-05

0.000230474

0.000226366

9.98546e-05

0.000342184

0.000124946

0.000172206

0.000173443

0.000107103

0.000160253

0.000150988

0.000132524

0.000152503

0.000140594

0.000157309

0.000143026

8.19635e-05

0.00013336

0.000116724

0.00015905

0.000181569

0.000148038

0.000114018

0.000224054

0.00016561

0.000148594

0.00010788

9.21235e-05

0.000128827

0.000189994

0.000115642

0.000100547

0.000124447

8.11172e-05

0.000114266

0.000485804

0.000142717

8.28315e-05

0.000136497

0.000210062

1.47904e-05

0.000115322

0.000124494

0.000168839

0.000128702

0.000125326

0.000116047

0.000138367

0.000181477

0.000287325

0.000198439

0.000106226

0.000171633

0.00014952

0.000145406

0.000110768

0.000103407

3.97274e-05

0.000154687

0.000419426

0.000181967

0.000126388

0.000176101

9.82078e-05

0.000193132

0.000108081

2.52806e-08

0.000128741

0.000138699

0.000195148

0.000145122

0.000110683

0.000173517

0.000135689

0.006709602

0.000493265

0.000146301

0.000346964

0.000424081

0.000342236

0.000360154

0.000158783

0.000256175

0.000712653

0.002146435

0.000899437

0.00032532

0.000921678

0.00181582

0.000157991

9.57616e-05

0.000181004

0.000182171

0.000195467

0.000356539

0.000147205

9.8946e-05

0.000181135

0.000905387

0.000106552

0.000136602

0.00015825

0.000116058

0.000107033

0.000114533

0.000115907

0.000117901

2.52806e-08

0.000110506

0.000119308

9.72614e-05

0.000230048

0.000157904

0.000472539

0.000131957

0.000122674

0.000186154

0.000406892
0.000813784

0.000269655

0.000137237

0.000237804
0.000118902

0.000118902

0.00152617
0.000763085

0.000122536

0.000111513

0.00011057

0.000114596

0.000130325

0.000173545

0.002107936
0.004215872

0.000117415

0.000125928

0.000131196

0.000115696

0.000217204

0.00011522

0.0001213

0.000128028

0.00013559

0.000136896

0.000192209

0.000106097

0.000165884

0.000159966

0.000139307

0.000247266
0.000123633

0.000123633

0.0004149598
0.0002074799

0.000121516

8.59639e-05

0.002560182
0.001280091

0.0002394

0.000139273

0.000150537

0.000106887

0.000252489

0.000135361

0.000125639

0.000130505

0.0117579548
0.0058789774

0.000115078

0.000133839

0.000119063

0.000119238

0.000120688

0.000101487

0.000209233

0.000274647

8.7842e-05

0.000995883

0.000231995

0.000124416

0.000136861

0.000263524

0.000239087

0.000150019

0.000115297

0.00014646

0.000122457

0.000112195

0.000129693

0.00013209

0.000121225

9.47272e-05

0.000127077

0.000120124

0.000163481

0.000101739

0.000127608

0.000104539

0.000258431

0.000105757

0.000128832

0.000113703

0.000129057

0.000329064

8.96927e-05

0.000124833

0.000125765

9.37625e-05

0.000104673

0.000129678

0.001097607
0.002195214

0.000337975

0.000299719

0.000155893

0.000103516

0.000200504

0.000949018
0.000474509

0.000110921

0.000188102

0.000175486

0.000608441
0.001216882

0.000142418

0.000132003

0.000195262

0.000138758

0.0001792714
8.96357e-05

8.96357e-05

0.000104981
0.000209962

0.000104981

0.000855817
0.001711634

0.000210312

0.000155998

0.000135118

0.000181931

0.000172458

0.000143828
0.000287656

0.000143828

0.000816158
0.001632316

0.000816158

0.001020398
0.000510199

0.000510199

0.000150314

0.017824296
0.008912148

0.000228802

0.000230299

0.0001173

0.000172911

0.000130493

0.00014507

0.000122681

0.000168

0.000138288

0.000160747

0.000253488

0.00062484

0.000174819

0.000225467

0.00018488

0.000146002

0.000162323

0.000202717

0.000130401

0.00080859

0.000275102

0.000156139

0.000149823

0.000186678

0.00016624

0.000160315

0.000115911

0.000139365

0.000125265

0.000152709

0.000140224

0.000144031

0.000477562

0.000224698

0.000220306

0.000231818

0.000156139

0.000146413

0.000177152

0.000167091

0.000220285

0.000140241

0.00015711

0.000153413

0.00782726
0.00391363

0.000183124

0.000158668

0.000199247

0.000111457

0.000150977

0.000131676

0.000145032

0.000114684

0.0001621

0.000110309

0.000162922

0.000132387

0.000175678

0.000210245

0.000121738

0.000148212

0.000118892

0.000153812

0.00036235

0.000296992

0.000563128

0.002773033
0.005546066

0.000285359

0.000166336

0.000234857

0.000140103

0.000190296

0.000153606

0.000134213

0.000118438

0.000158614

0.000308734

0.000143122

0.000124464

0.000177771

0.000131491

0.000175775

0.000129854

0.000153057

0.000544958
0.001089916

0.000116673

0.000124671

0.000155356

0.000148258

0.006047956
0.003023978

0.000148908

0.000107859

0.000131358

0.000240207

0.000148091

0.000168466

0.000152999

0.000115151

0.000151459

0.000155498

0.000136973

0.000461401

0.000249133

0.00013134

0.000145929

0.000143063

0.000120632

0.000115511

0.000515174
0.000257587

0.000257587

0.001107446
0.002214892

0.000221859

0.000132917

0.000116897

0.000127071

0.00011694

0.000159865

0.000116522

0.000115375

0.000326758
0.000163379

0.000163379

0.000610018
0.000305009

0.000146895

0.000158114

0.000176175
0.00035235

0.000176175

0.000267126
0.000133563

0.000133563

0.0007374998
0.0003687499

0.00010655

0.000167259

9.49409e-05

0.00125503
0.000627515

0.000148748

0.000163121

0.000154297

0.000161349

0.000286412
0.000143206

0.000143206

0.000319034
0.000159517

0.000159517

0.000187584

0.00024809
0.000124045

0.000124045

0.0002086073
0.0004172146

0.000119223

8.93843e-05

0.000356032
0.000712064

0.0002048

0.000151232

3.16488e-07

0.002474406
0.001237203

0.000136536

0.000119718

0.000214026

0.000232916

0.000534007

0.0007103897
0.0014207794

0.000115615

0.00011977

9.52827e-05

0.000153663

0.00011784

0.000108219

0.000216708

0.000310422
0.000620844

0.000133669

0.000176753

0.000108959
0.000217918

0.000108959

0.000183798
0.000367596

0.000183798

0.000286126
0.000143063

0.000143063

0.000155828
0.000311656

0.000155828

0.002848602
0.001424301

0.000154744

0.000204035

0.000198353

0.000131499

0.000121857

0.000148024

0.000152573

0.00011356

0.000199656

0.000442186
0.000221093

0.000101677

0.000119416

0.000599746
0.000299873

0.000299873

0.001843432
0.000921716

0.000199915

0.00021434

0.000132491

0.000244456

0.000130514

0.000444164
0.000888328

0.00010531

0.000105914

0.00023294

0.000352694
0.000176347

0.000176347

0.001998469
0.005646949

0.000283953
0.000567906

0.000150756

0.000133197

0.001172249
0.002344498

0.000170673

0.000293556

0.000184167

0.000170227

0.000176143

0.000177483

0.000193809
0.000387618

0.000193809

0.000160294

0.000188164

0.36560601
0.73121202

0.177834

0.00339838

0.00115369

0.00397548

0.00385931

0.0220693

0.00435885

0.148957

2.35367876498
11.335165188673

0.000276197
0.001104788

0.000828591
0.000276197

0.000552394
0.000276197

0.000122182

0.000154015

0.000256312

0.000290118

0.000314745

8.92820992509301
2.33907376158

0.000295572

0.000238939

0.000888129
0.001776258

0.000246162

0.000641967

0.000237792
0.000713376

0.000237792
0.000475584

0.000237792

0.00072327
0.00024109

0.00048218
0.00024109

0.00024109

0.000327898

0.079095813
0.233896639

0.003436864
0.001718432

0.000334953

0.000302516

0.000245938

0.000550851

0.000284174

0.000418146
0.000209073

0.000209073

0.000175437
0.000350874

0.000175437

0.000290501

0.000278997

0.000245803

0.000220486
0.000440972

0.000220486

0.001916642
0.003833284

0.00157772

0.000338922

0.0002131

0.001093392
0.000546696

0.000546696

0.000211027

0.012489597
0.024979194

0.000317338

0.000184374

0.000371336

0.000443447

0.000180151

0.000231671

0.000324319

0.00026681

0.000205991

0.000441812

0.00027997

0.000246184

0.00720064

0.000269241

0.000195811

0.000220442

0.000308364

0.000801696

0.000254516

0.000271615

0.000495424
0.000247712

0.000247712

0.000223365

0.000244274

0.001810756
0.003621512

0.000328508

0.000325004

0.000401214

0.000399514

0.000356516

0.000411074
0.000822148

0.000411074

0.000432106
0.000216053

0.000216053

0.000188918
0.000377836

0.000188918

0.00052891
0.000264455

0.000264455

0.000490423
0.000980846

0.000490423

0.000671942
0.000335971

0.000335971

0.000289642

0.009001332
0.018002664

0.00153325

0.000371257

0.000242421

0.000583377

0.000231394

0.000470908

0.000318971

0.000243947

0.000253146

0.000359168

0.000312454

0.000279367

0.000376814

0.000858535

0.000536701

0.000916922

0.0011127

0.000258141
0.000516282

0.000258141

0.000283727

0.00119788
0.00059894

0.000365314

0.000233626

0.000614994
0.000307497

0.000307497

0.000418879
0.000837758

0.000418879

0.000186092
0.000372184

0.000186092

0.00112358
0.00224716

0.00112358

0.00049725
0.000248625

0.000248625

0.000759106
0.000379553

0.000379553

0.000263786
0.000527572

0.000263786

0.007030405
0.01406081

0.00669754

0.000332865

0.00057582
0.00028791

0.00028791

0.005699446
0.011398892

0.000287054

0.00148298

0.00336879

0.000288975

0.000271647

0.004707678
0.002353839

0.000382149

0.000405899

0.000255628

0.000310216

0.000460288

0.000266117

0.000273542

0.023079011
0.046158022

0.0213382

0.000317407

0.000314079

0.000214856

0.000310725

0.000583744

0.001735458
0.000867729

0.000295348

0.000572381

0.000343108

0.000563922
0.000281961

0.000281961

0.000290222
0.000580444

0.000290222

0.00178634
0.00357268

0.00178634

0.000241125

0.000887984
0.001775968

0.000318758

0.000569226

0.002194152
0.000731384

0.001462768
0.000731384

0.000205806

0.000207647

0.000136748

0.000181183

0.000323073
0.000969219

0.000323073
0.000646146

0.000158106

0.000164967

0.000422673

0.000299978
0.000149989

0.000149989

0.000518051

0.001757538
0.005272614

0.003515076
0.001757538

0.000204675

0.00022931

0.000454449

0.000869104

0.000300156
0.000900468

0.000600312
0.000300156

0.000129573

0.000170583

0.001372329
0.000457443

0.000233882
0.000467764

0.000233882

0.000223561
0.000447122

0.000223561

0.000298278

0.000291691

0.00417674

0.000471264
0.000942528

0.000471264

0.003711996
0.007423992

0.00028882

0.000226334

0.000221202

0.000266524

0.000216858

0.000288491

0.000267944

0.000268039

0.00026304

0.000250425

0.000245666

0.000908653

0.000185149

0.000393117

0.000388023

0.000296646

0.000668702

0.000665268
0.000332634

0.000332634

0.000299778

0.005174286
0.001724762

0.000139701
0.000279402

0.000139701

0.001264364
0.002528728

0.000922096

0.000154345

0.000187923

0.000133629
0.000267258

0.000133629

0.000187068
0.000374136

0.000187068

0.008154237
0.002848428

0.004914762
0.002457381

0.000309907

0.00031169

0.000430593

0.000414261

0.000431587

0.00031168

0.000247663

0.000391047

0.0004221
0.0012663

0.0004221
0.0008442

0.0004221

0.000479051

0.000475692
0.000237846

0.000237846

0.000198743

0.001750302
0.000583434

0.000318565
0.00063713

0.000318565

0.000264869
0.000529738

0.000264869

0.000528585

0.000612672151
0.001838016453

0.000340200151
0.000680400302

0.000244218

1.31551e-07

9.58506e-05

0.000272472
0.000544944

0.000140954

0.000131518

0.000538155

0.000430507
0.000861014

0.000220051

0.000210456

0.00034676

0.000252715

0.000277961
0.000555922

0.000277961

0.057206961
0.01938971

0.000903728
0.001807456

0.00026373

0.000232477

0.000241256

0.000166265

0.015354401
0.030708802

0.00045685

0.000254313

0.000305797

0.000594457

0.000802476

0.00164945

0.000248477

0.000347153

0.000404755

0.000369594

0.000704243

0.00114594

0.00115003

0.000679687

0.000921094

0.000571573

0.000400478

0.000521401

0.000462514

0.000395084

0.000712782

0.000339688

0.00038847

0.00108959

0.000438505

0.000201048
0.000402096

0.000201048

0.000439816
0.000219908

0.000219908

0.000690453

0.00024601
0.00049202

0.00024601

0.00060709
0.00121418

0.00029974

0.00030735

0.000895356
0.001790712

0.000895356

0.000271716

0.000405471

0.000258692

0.000817511

3.2962818836
1.1238774952

0.000370006

0.000299653

0.793744271
1.587488542

0.0895344

0.000752138

0.549506

0.000777354

0.000878709

0.000727919

0.000839041

0.0609619

0.00494535

0.0839746

0.00084686

0.000411546

0.000350246

0.000487064

0.000616787

0.00498436
0.00249218

0.00205197

0.00044021

0.006423354
0.003211677

0.000407598

0.0011658

0.000432939

0.00120534

0.000312828

0.000533818

0.000485481

0.00020781

0.000693756
0.000346878

0.000346878

0.0004157

0.015019301
0.030038602

0.000436459

0.000555521

0.00186203

0.000491927

0.00046971

0.000299454

0.00165209

0.00167255

0.000486644

0.000351097

0.000420187

0.000298979

0.00041585

0.00105428

0.000510121

0.000729198

0.000333476

0.0011467

0.000304021

0.00055966

0.000460043

0.000509304

0.000305117

0.000519953

0.000259427

0.000461125

0.04920202
0.02460101

0.00042572

0.00137674

0.00130001

0.000637421

0.00023186

0.000389833

0.00209659

0.000516447

0.000445407

0.000380989

0.000397634

0.00118873

0.00032606

0.000322779

0.000724306

0.000871218

0.0011557

0.000414007

0.000581095

0.000283154

0.000240152

0.000341837

0.000988032

0.000349805

0.000431214

0.000571481

0.000830348

0.000296725

0.00205114

0.000497545

0.000351669

0.000525753

0.0003906

0.000433875

0.000468438

0.00150908

0.000257616

0.000316337

0.024807289
0.049614578

0.000279273

0.00034744

0.000396491

0.00111016

0.000371244

0.00522048

0.000438318

0.000687424

0.00123653

0.000486293

0.00038474

0.000319889

0.000424635

0.000633268

0.000428354

0.000477185

0.000325265

0.000491202

0.000371982

0.000433433

0.000389893

0.000387168

0.000327928

0.000377849

0.000411858

0.000381916

0.00036009

0.00047147

0.000312184

0.00041492

0.000412508

0.00036974

0.000368697

0.000406949

0.000348292

0.000391032

0.000351441

0.000387857

0.000510435

0.000322943

0.0018258

0.000412713

0.0134628
0.0269256

0.0134628

0.000374843

0.000464393

0.000383497

0.000363809

0.000260384

0.00040892

0.000399504

0.000372397

0.000363553

0.000362052

0.000392264

0.00040105

0.000478278

0.001111778
0.000555889

0.000555889

0.00573548
0.00286774

0.00123794

0.0016298

0.000429585

0.00699593

0.000271497

0.000481173

0.000297725

0.000282935

0.000347935

0.000317491

0.000367054

0.00224759

0.000759957
0.001519914

0.000759957

0.000667064
0.000333532

0.000333532

0.000920615
0.00184123

0.000280137

0.000378884

0.000261594

0.000242366

0.00051415
0.0010283

0.00051415

0.000557782

0.000265625
0.00053125

0.000265625

0.00272349

0.003499944
0.001749972

0.000893429

0.000856543

0.000339815

0.000385248

0.005649918
0.002824959

0.000266001

0.00132598

0.000503193

0.000343957

0.000385828

0.000496784
0.000993568

0.000496784

0.001683036
0.000841518

0.000490929

0.000350589

0.00039497

0.00811096

0.000286803

0.000435769

0.000420566

0.000338284

0.000487676

0.005018892
0.002509446

0.00207644

0.000433006

0.000396217
0.000792434

0.000396217

0.000404645

0.005488901
0.010977802

0.00130544

0.000281611

0.00035546

0.000431433

0.000295967

0.000303958

0.000289715

0.000553443

0.000370293

0.000377803

0.000311622

0.000296369

0.000315787

0.003103278
0.001551639

0.000464689

0.00108695

0.000534661
0.001069322

0.000242937

0.000291724

0.001273511
0.002547022

0.000305485

0.00035117

0.000616856

0.000867028
0.000433514

0.000433514

0.000612806
0.000306403

0.000306403

0.000263669

0.000445661

0.000886726
0.000443363

0.000443363

0.000301163

0.001349862
0.002699724

0.000286669

0.000418434

0.000644759

0.000856946
0.000428473

0.000428473

0.1039314
0.2078628

0.0251867

0.0251897

0.0313138

0.0222412

0.000340567

0.0089735424
0.0044867712

0.000867268

0.000806455

0.000468007

1.04442e-05

0.00140149

0.000513266

0.000419841

0.00264892
0.00132446

0.00132446

0.000353166

0.00852515

0.000421279

0.004465576
0.008931152

0.000755193

0.000621154

0.0015508

0.000316998

0.000343516

0.000357924

0.000175224

0.000344767

0.0179106

0.000357807

0.000861426
0.000430713

0.000430713

0.00398203

0.000334719

0.000280708

0.000683913

0.00400856
0.00200428

0.00200428

0.00158871
0.000794355

0.000794355

0.001415501
0.002831002

0.000440657

0.000343491

0.000631353

0.000340158

0.000406325

0.000464859

0.000334828

0.000342615

0.0502834
0.0251417

0.0251417

0.000324601

0.000391653

0.03799701
0.012757557

0.000488068
0.000244034

0.000244034

0.000257417
0.000514834

0.000257417

0.000308358
0.000616716

0.000308358

0.000295897
0.000591794

0.000295897

0.002661366
0.001330683

0.000196844

0.000174756

0.00015695

0.000149336

0.000173859

0.000153346

0.000325592

0.000264614
0.000132307

0.000132307

0.000191998
0.000383996

0.000191998

0.000291942
0.000583884

0.000291942

0.000275661

0.000847658
0.000423829

0.000230179

0.00019365

0.006625268
0.003312634

0.000201937

0.000264731

0.000166017

0.000260819

0.000239057

0.000194049

0.000256888

0.000182898

0.000214471

0.000201326

0.00024507

0.000252975

0.000260255

0.000195333

0.000176808

0.00188591
0.00377182

0.00067702

0.00026634

0.000237175

0.00024716

0.000226935

0.00023128

0.000274376
0.000137188

0.000137188

0.000301078
0.000150539

0.000150539

0.00703832
0.00351916

0.000192648

0.000178311

0.000220607

0.000280129

0.000102161

0.000211765

0.000110669

0.000213571

0.000138631

0.000158726

0.000118257

0.000252247

0.000247946

0.000156499

0.000167426

0.000129207

0.000204113

0.000157683

0.000278564

0.00026946

1.25149103146
0.41745457223

0.000246643

0.00132456
0.00066228

0.000198653

0.00017912

0.000284507

0.000220597
0.000441194

0.000220597

0.000222993
0.000445986

0.000222993

9.47123e-06

0.000234549
0.000469098

0.000234549

0.000738216
0.000369108

0.000369108

0.000476128
0.000952256

0.000476128

0.001223494
0.000611747

0.000112767

0.000118781

0.000256434

0.000123765

0.794481104
0.397240552

0.000371673

0.000727821

0.000164933

0.00022551

0.000508243

0.000521063

0.000318355

0.000869697

0.000197277

0.000320035

0.000170487

0.00024581

0.000320101

0.000401579

0.000254868

0.000510157

0.000382026

0.000232932

0.000162055

0.000475952

0.000193342

0.000260136

0.000239108

0.000181973

0.000200687

0.0040946

0.000705158

0.000309151

0.000421091

0.000210731

0.000232765

0.00016138

0.000198365

0.000264305

0.000237761

0.000298008

0.000211415

0.000571436

0.000443018

0.000220314

0.000368034

0.000244051

0.000387813

0.000619088

0.000260376

0.000590234

0.000226198

0.000310045

0.000450805

0.000257388

0.00017979

0.000228006

0.00022829

0.000594588

0.000683969

0.00038752

0.000260658

0.000270111

0.000778975

0.000200177

0.000317657

0.000266193

0.000359482

0.00020729

0.000604493

0.000518466

0.000194599

0.000255774

0.00017268

0.00882013

0.000374267

0.000227802

0.000303713

0.00035191

0.000389988

0.000447565

0.000413261

0.000268371

0.00041214

0.000209137

0.00225637

0.29789

0.000454797

0.000240173

0.000265749

0.000397057

0.000181465

0.000267164

0.000346555

0.000294642

0.000273272

0.000543437

0.000242621

0.000282312

0.00032348

0.000552686

0.000400169

0.000278847

0.000268832

0.000448304

0.000381422

0.000190583

0.000394066

0.000663761

0.000222012

0.000478925

0.000430144

0.000267997

0.00268004

0.000572197

0.000432724

0.000388729

0.000528693

0.000228468

0.000200959

0.000246637

0.00038827

0.00023268

0.00861393

0.000208992

0.000200473

0.000221864

0.000561063

0.000310396

0.000222638

0.000287364

0.0001941

0.000647999

0.000236654

0.000516111

0.000242421

0.00040564

0.000248109

0.000271337

0.000329107

0.000387217

0.000218551

0.000715733

0.000260459

0.000273518

0.000253747

0.000374279

0.0142199

0.000409359

0.000204421

0.000236391

0.000256366

0.000204133

0.000294299

0.000259611

0.000227295

0.000661338

0.000195906

0.000444609

0.000225725

0.000405695

0.000231718

0.00382677

0.000717582

0.000394356

0.000229862

0.000210336

0.000181818

0.000232147

0.00051032

0.000229101

0.000207847

0.000198669

0.000208746

0.000287636

0.000314303

0.000616571

0.000237587
0.000475174

0.000237587

0.000886022
0.000443011

0.000443011

0.000448796
0.000897592

0.000448796

0.000259829
0.000519658

0.000259829

0.00054522
0.00027261

0.00027261

0.000667792
0.000333896

0.000333896

0.000349307
0.000698614

0.000349307

0.000510848
0.000255424

0.000255424

0.000849958
0.000424979

0.000196861

0.000228118

0.00611082
0.01222164

0.00611082

0.000631714
0.000315857

0.000315857

0.000365767
0.000731534

0.000190377

0.00017539

0.001103774
0.002207548

0.000761222

0.000342552

0.00044899
0.00089798

0.00044899

0.002957676
0.001478838

0.000167891

0.000240185

0.000215934

0.000197216

0.000444058

0.000213554

0.00024509
0.00049018

0.00024509

0.000504202
0.000252101

0.000252101

0.000202752
0.000405504

0.000202752

0.000240645
0.00048129

0.000240645

0.001538338
0.000769169

0.000769169

0.000743606
0.000371803

0.000371803

0.000277734
0.000555468

0.000277734

0.00220476
0.00110238

0.00030952

0.000207002

0.000293534

0.000292324

0.000465548
0.000232774

0.000232774

0.000308106

0.621554835
0.209526579

0.0023943
0.0047886

0.000460299

0.000364215

0.000415234

0.000385322

0.000372945

0.000396285

0.000233845
0.00046769

0.000233845

0.001581828
0.000790914

0.000433313

0.000357601

0.085065284
0.170130568

0.000720984

0.0843443

0.00167985

0.00179084

0.000279868

0.0023542
0.0011771

0.0011771

0.00041289
0.00082578

0.00041289

0.00154995

0.000809511
0.001619022

0.000809511

0.001083606
0.002167212

0.000389526

0.00069408

0.000326288

0.000419862

0.09657058
0.19314116

0.00252225

0.0507501

0.00220681

0.02149

0.00408932

0.0155121

0.000809767
0.001619534

0.000286601

0.000284828

0.000238338

0.001504306
0.000752153

0.000752153

0.0026553
0.00132765

0.00132765

0.000350604

0.000526071
0.001052142

0.0003058

0.000220271

0.00952715
0.0190543

0.00426228

0.00526487

0.00062764

0.002041712
0.001020856

0.000371483

0.000649373

0.000265763

0.000355826

0.000419757

0.000220861

0.004056088
0.002028044

0.000294372

0.000257649

0.000294177

0.000259251

0.000373642

0.000244199

0.000304754

0.000609883

0.000429046

0.000452978

0.441025533
1.024924594

0.259962

0.0252931

0.000650334
0.000325167

0.000325167

0.2782001
0.13910005

0.0056129

0.00589605

0.116403

0.0111881

0.000451235
0.00090247

0.000451235

0.001455236
0.000727618

0.000727618

0.00130545
0.0026109

0.00130545

0.001928016
0.000964008

0.000964008

0.000316965

0.00268938

0.00881717

0.00107339

0.000331656
0.000663312

0.000331656

0.0383896216
0.0095974054

0.0099781446
0.0033260482

0.0031237404
0.0015618702

0.000124391

0.000118809

0.000116912

8.82186e-05

0.000118023

0.000119427

0.000119974

8.53993e-05

0.000203919

0.000120493

0.000133261

0.000114735

9.83083e-05

0.00031863
0.000159315

0.000159315

0.000201808
0.000100904

0.000100904

0.0003295
0.00016475

0.00016475

0.002378444
0.001189222

0.000119889

0.000217076

0.000184386

0.000357026

0.000188658

0.000122187

0.000149987
0.000299974

0.000149987

0.017756778
0.005918926

0.001778602
0.000889301

0.000135293

0.000363972

0.000129149

0.000131195

0.000129692

0.000158468
0.000316936

0.000158468

0.000150282
0.000300564

0.000150282

0.000131994
0.000263988

0.000131994

0.002340223
0.004680446

0.000149092

0.000149076

0.000126933

0.000132519

0.000143431

0.000177498

0.000106254

0.000177034

0.000142173

0.000140273

0.000138146

0.000123551

0.000134796

0.000129423

0.000217311

0.000152713

0.000116709
0.000233418

0.000116709

0.000350834
0.000701668

0.000172577

0.000178257

0.000711978
0.001423956

0.000160868

0.00016926

0.000241402

0.000140448

0.000334878
0.000167439

0.000167439

0.000568032
0.000284016

0.000284016

0.000646174
0.000323087

0.000150871

0.000172216

0.000294595
0.00058919

0.000294595

0.000188815
0.000566445

0.000188815
0.00037763

0.000188815

0.0001636162
0.0004908486

0.0001636162
0.0003272324

7.09368e-05

9.26794e-05

0.000200695

0.000219814

0.000226669

0.000206152

0.012067584
0.003016896

0.001520966
0.004562898

0.000316634
0.000158317

0.000158317

0.000750432
0.000375216

0.000126944

0.000117922

0.00013035

0.000336368
0.000168184

0.000168184

0.001058434
0.000529217

0.000143709

0.000385508

0.000125507
0.000251014

0.000125507

0.00032905
0.000164525

0.000164525

0.00448779
0.00149593

0.000195021
0.000390042

0.000195021

0.000236356
0.000118178

0.000118178

0.002365462
0.001182731

0.000131039

0.000144285

0.000147136

0.00024477

0.000360332

0.000155169

0.000292823

0.000362983

8.95817567323
44.78572596315

3.583125276
0.895781319

2.647783656
0.882594552

0.0244492
0.0122246

0.0122246

0.85468308
1.70936616

0.0064305

0.0606727

0.0137401

0.116245

0.1302

0.0164538

0.00114596

0.00997182

0.0112407

0.0180663

0.00204904

0.0194214

0.00120642

0.198534

0.0136555

0.132462

0.0260758

0.00588911

0.00597092

0.0108239

0.0259062

0.00105021

0.0168151

0.0106566

0.001536751
0.003073502

0.000774243

0.000762508

0.01236726
0.00618363

0.002395

0.00169812

0.00209051

0.00217982
0.00108991

0.00108991

0.003988309
0.007976618

0.00141479

0.00122234

0.000701108

0.000650071

0.002888272
0.005776544

0.000589215

0.000557057

0.001742

0.013186767
0.039560301

0.000648044
0.001296088

0.000648044

0.001342038
0.000671019

0.000671019

0.000638797
0.001277594

0.000638797

0.006680682
0.003340341

0.00130381

0.00082627

0.000639411

0.00057085

0.00453874
0.00907748

0.000999237

0.000877952

0.00060485

0.000963774

0.000568534

0.000524393

0.000655167
0.001310334

0.000655167

0.000542517
0.001085034

0.000542517

0.001247517
0.002495034

0.000774077

0.00047344

0.000447667
0.000895334

0.000447667

0.000913916
0.000456958

0.000456958

8.05514158723
32.21541394592

24.16027235869
8.05514158723

0.00130597
0.00261194

0.00130597

0.001165426
0.000582713

0.000582713

15.51941295646
7.75970647823

1.05788

0.0337908

1.56875

0.36143

0.444585

0.0128661

0.296739

0.0454592

0.0307943

0.225999

0.360417

0.0094675

0.39236

0.0384741

1.57823e-06

0.0425991

0.294538

0.402289

0.262452

0.765997

0.262667

0.267134

0.0063308

0.330921

0.245765

0.26248206
0.52496412

0.00210181

0.00142833

0.00120034

0.00190939

0.0030503

0.00166704

0.00156447

0.00128082

0.00186593

0.00334278

0.00120833

0.00133105

0.00392923

0.00164288

0.00185724

0.00152947

0.00119154

0.00276611

0.227615

0.00642448
0.01284896

0.00642448

0.00312662

0.00139842

0.000627363

0.00171461
0.00342922

0.00171461

0.025892706
0.012946353

0.000877894

0.00114868

0.00763577

0.00244093

0.000843079

0.00965304
0.00482652

0.00482652

0.029011068
0.007252767

0.007252767
0.021758301

0.009954196
0.004977098

0.000429213

0.000520041

0.000563747

0.000709703

0.000771204

0.00100627

0.00097692

0.0014672
0.0007336

0.0007336

0.000757748
0.001515496

0.000757748

0.001568642
0.000784321

0.000784321

0.000156545

0.489885754
0.100458188

0.000434394
0.000868788

0.000254546

0.000179848

0.000922374

0.000318087

0.000545804
0.000272902

0.000272902

0.000923536
0.000461768

0.000228772

0.000232996

0.095889937
0.382913995

0.0381024
0.114103468

0.073607348
0.036803674

0.000247174

0.0365565

0.00161886
0.00080943

0.000247529

0.000325581

0.00023632

0.000571128
0.000285564

0.000285564

0.000203732

0.002175507
0.000725169

0.000481768
0.000240884

0.000240884

0.000484285
0.00096857

0.000244748

0.000239537

0.000442021
0.000884042

0.000206567

0.000235454

0.169861041
0.056620347

0.00259793
0.00519586

0.00259793

0.022389506
0.011194753

0.000319328

0.000454297

0.000298754

0.000387178

0.000417208

0.000803153

0.000475061

0.000709372

0.00157998

0.000169217

0.000326621

0.000437339

0.00103204

0.000448097

0.000442147

0.000328706

0.000333558

0.00148743

0.000568679

0.000176588

0.02424216
0.04848432

0.00274605

0.00117112

0.00215529

0.0181697

0.00098262
0.00196524

0.000271384

0.000358232

0.000353004

0.001320528
0.000660264

0.000316231

0.000344033

0.000396818
0.000793636

0.000396818

0.014374804
0.028749608

0.000181822

0.000356618

0.000550744

0.000477177

0.000591938

0.000601206

0.000202835

0.000139331

0.00059599

0.000265403

0.000273594

0.000498135

0.000557972

0.000328629

0.000382606

0.000468893

0.000743445

0.000415808

0.000400665

0.000506348

0.000484209

0.00111987

0.000352807

0.000285856

0.000588728

0.00174747

0.000712039

0.000544666

0.000608331
0.001216662

0.000178249

0.000430082

0.000413138
0.000826276

0.000413138

0.000390876
0.000195438

0.000195438

0.001908182
0.000954091

0.000325613

0.000628478

0.00138247

0.000607141
0.001214282

0.00033723

0.000269911

0.00033823
0.000169115

0.000169115

6.94855e-07

0.000302048
0.001812288

0.000302048
0.00151024

0.001208192
0.000302048

0.000906144
0.000302048

0.000143871
0.000287742

0.000143871

0.000158177
0.000316354

0.000158177

0.000228663
0.001371978

0.000228663
0.001143315

0.000914652
0.000228663

0.000685989
0.000228663

0.000228663
0.000457326

0.000121027

0.000107636

6.81449e-05
0.0002044347

0.0001362898
6.81449e-05

6.81449e-05

0.0301041526
0.0053452663

0.0022749525
0.0005178765

0.000116918
0.000467672

0.000350754
0.000116918

0.000233836
0.000116918

0.000116918

0.000157215
0.00031443

0.000157215

0.000974974

0.0002437435
0.000974974

0.0007312305

0.0002437435
0.0007312305

0.000177541
8.87705e-05

8.87705e-05

0.000154973
0.000309946

0.000154973

0.001461495
0.000292299

0.001169196
0.000292299

0.000876897
0.000292299

0.000584598
0.000292299

0.000292299

0.001914195
0.000382839

0.000150174
0.000600696

0.000450522
0.000150174

0.000300348
0.000150174

0.000150174

0.000232665
0.00093066

0.000232665
0.000697995

0.000232665
0.00046533

0.000106959

0.000125706

0.006445595
0.001289119

0.004728628
0.001182157

0.000723327

0.000241109
0.000723327

0.000241109
0.000482218

0.000114883

0.000126226

0.002823144

0.000674642
0.002023926

0.000674642
0.001349284

0.00010178

0.000120784

0.000116074

0.000114672

0.000221332

0.000266406
0.000799218

0.0003063
0.00015315

0.00015315

0.000113256
0.000226512

0.000113256

0.000427848
0.000106962

0.000320886
0.000106962

0.000106962
0.000213924

0.000106962

0.001741425
0.000348285

0.000348285
0.00139314

0.000348285
0.001044855

0.00034772
0.00017386

0.00017386

0.000174425
0.00034885

0.000174425

0.000795455
0.000159091

0.000636364
0.000159091

0.000159091
0.000477273

0.000318182
0.000159091

0.000159091

0.0007636
0.0032039678

0.0004565839
0.0018263356

0.0004565839
0.0013697517

0.0009131678
0.0004565839

0.000378374

7.82099e-05

0.0006140322
0.0003070161

9.54912e-05

7.46838e-05

7.55568e-05

6.12843e-05

0.000214005

0.006707796
0.0013781518

0.0053296442
0.0013781518

0.0013781518
0.0039514924

0.000219336
0.000438672

0.00010632

0.000113016

8.95658e-05
0.0001791316

8.95658e-05

0.000182963

0.000171948
0.000343896

0.000171948

0.000258158
0.000129079

0.000129079

0.000279146
0.000139573

0.000139573

0.000105854
0.000211708

0.000105854

0.000102667
0.000205334

0.000102667

0.000134582
0.000269164

0.000134582

0.000102584
0.000205168

0.000102584

27.1185097648698
4.52228314079

5.95132e-05

4.8748e-05

0.239039449
1.192934337

0.009604239
0.037669188

0.028064949
0.009604239

0.000580318
0.000290159

0.000140176

0.000149983

0.000270844
0.000135422

0.000135422

0.000524098
0.000262049

0.000262049

0.00110747
0.000553735

0.000553735

0.002006306
0.001003153

0.000161909

0.000311033

0.00019596

0.00015375

0.000180501

0.000327724
0.000163862

0.000163862

0.001095497
0.002190994

0.000429134

0.000156017

0.000183832

0.000128241

0.000198273

0.000189941
0.000379882

0.000189941

0.000747768

0.000315728
0.000157864

0.000157864

0.000529327
0.001058654

0.000529327

0.000827881
0.001655762

0.000205912

0.000621969

0.007295162
0.003647581

0.000192878

0.00153863

0.000265253

0.00165082

0.916225699999999
0.22943521

0.020712527
0.007173343

0.004686033
0.009372066

0.000191261

0.000568634

0.000217359

0.000205773

0.000204742

0.000173209

0.00018506

0.000187105

0.000215329

0.000188869

0.000229549

0.000195016

0.00108742

0.00021518

0.000192775

0.000200746

0.000228006

0.000183454

0.000260533

0.00031059
0.000155295

0.000155295

0.000651278
0.000325639

0.000325639

0.000337708
0.000168854

0.000168854

0.00103002
0.00206004

0.000240332

0.000204365

0.000301045

0.000284278

0.000363515

0.000353819

0.665724144
0.221908048

0.000260036
0.000520072

0.000260036

0.209401495
0.41880299

0.0717403

0.000356727

0.0633743

0.000289383

0.000259134

0.00072089

0.00042152

0.00289994

0.00145356

0.063205

0.000252871

0.00145883

0.00296904

0.012246517
0.024493034

0.000299634

0.000425243

0.00186729

0.000291965

0.00036262

0.000352272

0.000328411

0.000348781

0.000368289

0.000866245

0.000381416

0.000377387

0.000799737

0.000824914

0.000396875

0.000347494

0.00131287

0.000370634

0.000341699

0.000408618

0.000332906

0.000841217

0.001831785
0.000366357

0.000639028
0.000159757

0.000479271
0.000159757

0.000159757
0.000319514

0.000159757

0.0008264
0.0002066

0.0002066
0.0006198

0.0002066
0.0004132

0.0002066

0.0005824449
0.0029122245

0.0005824449
0.0023297796

0.0012940707
0.0004313569

0.0001846242
9.23121e-05

9.23121e-05

0.0001017902
5.08951e-05

5.08951e-05

0.000123489
0.000246978

0.000123489

0.0001618336
8.09168e-05

8.09168e-05

0.0001674878
8.37439e-05

8.37439e-05

0.000151088
0.000453264

0.000151088
0.000302176

0.000151088

5.10322e-05

0.00188832
0.000377664

0.000377664
0.001510656

0.001132992
0.000377664

0.000377664
0.000755328

0.000131043

0.000111692

0.000134929

0.002502179
0.011916241

0.000478032
0.000119508

0.000358524
0.000119508

0.000239016
0.000119508

0.000119508

0.002382671
0.00893603

0.000297327

0.001827069
0.000609023

0.000609023
0.001218046

0.000291823

0.0003172

0.002114406
0.000704802

0.000704802
0.001409604

0.000215089

0.000261397

0.000228316

0.002314557
0.000771519

0.001543038
0.000771519

0.000266506

0.000505013

21.3845247942798
4.27919612456

0.00014407

0.342245282
0.0859315175

0.0136767325
0.0410301975

0.001017456
0.000508728

0.000156417

0.000166544

0.000185767

0.001834768
0.000917384

0.000290424

0.000173916

0.000192855

0.000260189

0.000301064
0.000150532

0.000150532

0.000314678
0.000157339

0.000157339

0.0019994953
0.0039989906

0.000193355

9.435e-05

0.000146558

0.000116605

0.000131328

0.000219786

9.73347e-05

0.000120212

0.00015575

0.000152155

0.000111202

9.79716e-05

0.000245001

0.000117887

0.00022161
0.000110805

0.000110805

0.000490156
0.000980312

0.000170663

0.000138931

0.000180562

0.000783214
0.000391607

0.000391607

0.00014905
0.0002981

0.00014905

0.0166691484
0.0083345742

0.000181684

0.000154907

0.00018286

0.000208628

0.000174149

0.000161448

0.000145099

9.90442e-05

0.000127261

0.000172401

0.000178299

0.000193415

0.000134888

0.000156179

0.000143306

0.000126191

0.000206292

0.000149079

0.000222387

0.000147084

0.000112293

0.000250643

0.000194333

0.000168887

0.000135474

0.000310759

0.000130504

0.000205795

0.000119975

0.000146566

0.000202705

0.000197657

0.000159187

0.000235147

0.000133761

0.000150183

0.000167134

0.000213738

0.000132543

0.000168618

0.000924368

0.000116872

0.000193936

0.000151724

0.000147171

0.000467062
0.000934124

0.000140105

0.000206053

0.000120904

0.215283567
0.072254785

0.000110288

0.003989724
0.001994862

0.000194459

0.000189422

0.000214815

0.000140524

0.000224784

0.00022685

0.000192987

0.000256495

0.00018194

0.000172586

0.00031509
0.000157545

0.000157545

0.0506524
0.0253262

0.0253262

0.031906961
0.063813922

0.00118159

0.000169978

0.000286379

0.000274871

0.00150806

0.00026537

0.00114342

0.00154724

0.00117003

0.00171665

0.00141005

0.000188414

0.000151922

0.00101045

0.00167852

0.00114128

0.00149999

0.00017001

0.00134706

0.00156859

0.00134705

0.00159427

0.00111736

0.000186896

0.000436795

0.00116791

0.00168771

0.000293434

0.00128639

0.000509568

0.00121562

0.000518864

0.00111522

0.00081584
0.00163168

0.000175589

0.000145842

0.000494409

0.000140057

0.000844396
0.001688792

0.000245416

0.000149395

0.000449585

0.000212166

0.000336342
0.000168171

0.000168171

0.000112005

0.000315835
0.00063167

0.000180465

0.00013537

0.000132352
0.000264704

0.000132352

0.000408448
0.000204224

0.000204224

0.000391374
0.000195687

0.000195687

0.000112973

0.000128747

0.000130626

0.000391193
0.000782386

0.000391193

0.000111025

0.016641462
0.008320731

0.000744651

0.00575546

0.000542366

0.000759639

0.000518615

0.00015026

0.000154652

0.000117989

0.001530392
0.000382598

0.001147794
0.000382598

0.000382598
0.000765196

0.000109516

0.000139475

0.000133607

0.000543474
0.000271737

0.000271737

0.000171654

7.0885040522
1.7725278986

9.03066e-05

0.000304595

5.315581252
1.772132997

0.003229192
0.001614596

0.000291144

0.000311282

0.00101217

0.001099567
0.002199134

0.000202521

0.000205907

0.000209849

0.00048129

0.000817739

3.478645312
1.739322656

0.000233243

0.000516008

0.05374

0.0603079

0.012942

0.00862856

0.00151572

0.0356803

0.0018364

0.00162084

0.00108033

0.00266644

0.0985751

0.00143861

0.0366803

0.000916744

0.000243555

0.0127822

0.000284252

0.000592961

0.000308341

0.0122721

0.000252985

0.0016475

0.000492774

0.000837527

0.271699

0.0115429

0.00108296

0.000308608

0.0139826

0.000597913

0.0320304

0.00338851

0.000529096

0.00139409

0.000282154

0.000604078

0.000934424

0.013736

0.0891499

0.0822232

0.000252528

0.199643

0.000313473

0.0312055

0.00551963

0.0269945

0.00243449

0.000226955

0.00388246

0.000254985

0.00112613

0.0276619

0.0892884

0.00251088

0.0241939

0.000834056

0.135936

0.000336197

0.00192569

0.004654

0.0178125

0.04042

0.000251245

0.0935226

0.00141289

0.000488581

0.000536191

0.000215107

0.0104167

0.00117674

0.0370681

0.00261204

0.0018868

0.00165829

0.00156191

0.0648527

0.000287523

0.00851483

0.000241232

0.020646

0.00106429

0.00190319

0.050003542
0.025001771

0.000341281

0.0240669

0.000197465

0.000396125

0.0010414
0.0020828

0.000450715

0.000590685

0.002978104
0.001489052

0.00026983

0.000248652

0.000348238

0.000622332

0.000251991
0.000503982

0.000251991

0.00298845
0.001494225

0.000368309

0.000240728

0.000885188

6.70217e-05

0.03163065871
0.12652263484

0.03163065871
0.09489197613

0.02129781291
0.04259562582

0.000128741

0.000160529

0.000165507

0.00031557

0.000100973

0.000165537

0.000233067

8.13956e-05

0.000430709

0.000100778

0.000288324

0.000144466

0.00391847

0.000118523

0.0001029

0.000100822

0.000131063

0.000128971

0.000147065

0.000204466

0.000142999

7.93262e-05

0.000371703

0.000130543

0.00222326

8.3578e-05

0.00016828

0.000266225

0.000242532

0.000192783

0.000124995

0.00027047

0.000133649

0.000122525

0.000252829

0.000151453

5.56991e-06

0.000592505

0.000124756

8.42587e-05

0.000138286

0.000164806

0.000143407

0.000169187

0.00017095

8.7779e-05

9.59953e-05

0.000121771

7.80727e-05

0.000223275

0.000106172

7.90086e-05

0.000115309

0.000173881

0.00362205

0.000423464

0.000134003

0.000231062

9.8367e-05

0.000191544

0.000195483

9.80149e-05

0.000166858

0.000315551

0.000215004

0.000156565

0.000114818

0.000535013

0.0001578548
7.89274e-05

7.89274e-05

0.002374396
0.001187198

0.000274097

0.000198801

0.00012648

0.00058782

6.38744e-05
0.0001277488

6.38744e-05

0.000280037
0.000560074

0.000280037

0.001060048
0.000530024

0.000405316

0.000124708

0.000284074
0.000568148

0.000151713

0.000132361

0.015817422
0.007908711

0.000649439

0.000525544

0.000525041

0.000329517

0.00587917

5.99641e-05

0.000230028

0.13400359728836
0.0335008993221

0.10050269796626
0.0335008993221

0.0098917603
0.0197835206

0.000214552

0.000141411

0.000271492

0.000208334

0.000177029

0.000230222

0.000121199

0.000142375

0.000344471

9.57457e-05

0.000164391

0.000107431

0.000205437

0.00028118

0.000182677

0.00020719

0.000179311

0.000139679

0.000285122

0.000357125

0.000206304

0.000119891

9.04206e-05

0.000151089

0.000205984

0.0001477

0.000110199

0.000238741

0.000164344

0.000275812

0.000407132

0.000204786

0.000622617

0.000129147

0.00018033

0.000129174

0.000129515

0.000195039

0.000107176

0.000123964

0.000494671

0.00011687

0.000274081

0.0010104

0.000640928
0.001281856

0.000152326

0.000222208

0.000266394

0.00135373
0.000676865

0.000241962

0.000434903

0.00051556
0.00025778

0.00025778

0.0021703528
0.0010851764

0.000140812

0.000241118

8.35944e-05

0.000100843

0.000167271

0.000111787

0.000239751

0.00329534
0.00164767

0.000317806

0.000192736

0.000193993

0.000171264

0.000155327

0.000233733

0.000168844

0.000213967

0.003715802
0.001857901

0.000244021

0.00161388

0.000405318
0.000202659

0.000202659

0.00392669
0.001963345

0.000162773

0.000131309

0.000191699

0.000187481

0.000213345

0.000388086

0.000247429

0.000441223

0.001815233
0.003630466

0.000128315

0.000109161

0.000230428

0.000131353

7.85785e-05

0.000109671

0.000124945

0.000102018

0.000311476

7.97959e-05

0.000138492

9.53926e-05

0.000175607

0.00110657
0.000553285

0.000364283

0.000189002

0.002156384
0.001078192

0.000131718

0.000142387

0.000198237

0.000184451

0.000212804

0.000208595

0.002911064
0.001455532

0.000250458

0.000148237

0.000154222

0.000234893

0.000227872

0.000154191

0.000148635

0.000137024

0.0001719254
8.59627e-05

8.59627e-05

0.000280827
0.000561654

0.000280827

0.001444636
0.000722318

0.000177847

0.000119703

0.000424768

0.000629831
0.001259662

0.000257283

0.000140079

0.000232469

0.012646692
0.006323346

0.000354599

0.000164675

0.000224463

0.000132854

0.000128396

0.000105463

9.67902e-05

9.75623e-05

0.000177716

0.00132482

0.000128332

0.000186626

0.000158215

0.000173613

0.000113332

0.000112296

0.000111116

0.000246219

9.70325e-05

0.000311872

0.000110112

0.000154568

0.000251048

0.000204524

0.000110551

0.00012517

0.000113565

0.000239906

0.000231969

0.00020684

0.000129101

0.000678168
0.000339084

0.000161482

0.000177602

0.000882954
0.000441477

0.000197428

0.000244049

0.001820808
0.000910404

0.000514644

0.000225625

0.000170135

0.00016778692208
0.00033557384416

6.92208e-09

0.00016778

0.000360329
0.000720658

0.000117141

0.000105194

0.000137994

0.000113207
0.000226414

0.000113207

0.0032944732
0.0008236183

0.0024708549
0.0008236183

0.0008236183
0.0016472366

0.000120592

0.000238461

7.02682e-05

5.60729e-05

5.91567e-05

8.79497e-05

7.44528e-05

0.000116665

0.00027514
6.8785e-05

0.000206355
6.8785e-05

6.8785e-05
0.00013757

6.8785e-05

0.001314865
0.00525946

0.000890451
0.000296817

0.000593634
0.000296817

0.000296817

0.001018048
0.003054144

0.001018048
0.002036096

0.000777048

0.000241

0.006705576
0.026245416

0.006705576
0.01953984

0.000552794
0.001105588

0.00012835

0.000214575

0.000209869

0.000218608

0.003713346
0.001856673

0.000108123

0.000151618

0.000106389

0.000233788

0.000117399

0.000140047

0.000163049

0.000183241

0.000150957

0.000156405

0.000127107

0.00021855

0.003092786
0.006185572

0.00018659

0.000158826

0.00015604

0.000158192

0.00015307

0.000134315

0.000285501

0.000445543

0.000202216

0.000142556

0.000175609

0.000147065

0.00022957

0.00015809

0.000222175

0.000137428

0.000185545

0.00125287
0.000626435

0.000173938

0.000135687

0.00014201

0.0001748

0.000172735

0.000241239

0.0248227555
0.006279343

0.000100977

0.001345314
0.000448438

0.000896876
0.000448438

0.00022265

0.000225788

0.000185325
9.26625e-05

9.26625e-05

0.0054082385
0.0162247155

0.00026822
0.00053644

0.000148708

0.000119512

0.0051400185
0.010280037

0.000103164

0.000123279

6.37033e-05

0.000147157

0.000244461

0.00013882

6.74352e-05

6.35433e-05

0.000218598

0.000165702

0.000125496

9.82489e-05

9.13527e-05

0.000131593

0.000109172

0.00023082

7.41392e-05

6.81905e-05

0.000175653

0.000237244

0.000211441

5.59609e-05

0.000164486

6.41537e-05

0.00018029

0.000119255

6.14388e-05

0.000189392

0.000166417

0.000253658

0.000190549

0.000169632

0.000160826

0.000194119

0.000146989

0.000133639

0.000687081
0.000229027

0.000229027
0.000458054

0.000229027

0.0185586229
0.0742344916

0.0556758687
0.0185586229

0.000177024
0.000354048

0.000177024

0.003312855
0.00662571

0.00023372

0.000238084

0.000175104

0.000248351

0.000175905

0.000199171

0.000227571

0.000209485

0.000180618

0.000210156

0.000137641

0.000298954

0.000131758

0.000206148

0.000216023

0.000224166

0.000339257
0.000678514

0.000339257

0.0001879404
9.39702e-05

9.39702e-05

0.000245986
0.000491972

0.000245986

0.000393394
0.000196697

0.000196697

0.0007926
0.0003963

0.000201338

0.000194962

0.00042038
0.00021019

0.00021019

0.000120965
0.00024193

0.000120965

0.000227276
0.000113638

0.000113638

0.0194349994
0.0097174997

0.000156382

0.000152721

0.000102519

0.000116983

0.000142562

0.00013965

0.000137867

0.00014758

0.000163529

8.8082e-05

0.000143425

0.000259323

9.52504e-05

0.00013619

0.000147979

9.52931e-05

0.000137615

0.000102043

0.000149124

0.000149222

0.000203633

9.56575e-05

0.000165262

0.000137764

0.000162596

0.000199454

0.000136206

0.000160252

0.000309443

0.000104966

0.000144437

0.000109224

9.05518e-05

0.000118718

0.00011794

0.000234939

0.000141063

0.000122084

0.000206832

0.000126426

0.000155125

0.000159686

0.000162112

0.00017349

8.9248e-05

0.000284088

0.000133677

9.16438e-05

0.000108921

0.000186855

0.000135647

0.000121899

9.79011e-05

0.000221129

0.000203385

0.000128165

0.000136656

0.00017968

0.000156715

0.000235528

0.000156401

0.00014141

0.000135002

0.000138995

0.000133353

0.000533351
0.001066702

0.000175226

0.000121758

0.000102416

0.000133951

0.000216206
0.000432412

0.000216206

0.00148543
0.00297086

0.000471833

0.000788611

0.000224986

0.00015453
0.00030906

0.00015453

0.002489448
0.001244724

0.000136065

0.000187967

0.000146856

0.000773836

0.09123942968
0.02286701867

0.0273509736
0.0091169912

0.000219336
0.000438672

0.000219336

0.0022715524
0.0011357762

1.21061e-05

0.000214372

0.000402728

0.000258305

6.28391e-05

0.000185426

0.000186349
0.000372698

0.000186349

0.002877176
0.001438588

0.00027624

0.000203926

0.000192922

0.000192814

0.000163763

0.000408923

0.002288214
0.001144107

0.000202849

0.000235306

0.000223664

0.000482288

0.00042779
0.000213895

0.000213895

0.000259251
0.000518502

0.000259251

0.00014027
0.00028054

0.00014027

0.001352902
0.002705804

0.000283691

0.000233554

0.000165586

0.000189766

0.000480305

0.006053034
0.003026517

0.000203736

0.000183738

0.000316376

0.00023694

0.000268208

0.000243747

0.000204908

0.000283702

0.00022506

0.000232716

0.000393358

0.000234028

0.02238590841
0.00746196947

0.000148204
0.000296408

0.000148204

0.000489618
0.000244809

0.000244809

0.000242448
0.000121224

0.000121224

0.000154254
0.000308508

0.000154254

0.01200184494
0.00600092247

0.000111662

0.000186098

0.000209119

0.000144425

8.08647e-06

0.00019817

0.000357683

0.000166412

0.00015391

0.000159149

0.000129954

0.000148733

0.00010523

0.000312846

0.000292012

0.000166072

0.000267292

0.000185872

0.000100201

0.000181796

0.000306049

0.000170405

0.000189368

0.000123812

0.000207071

0.000544123

0.000223198

0.000332828

0.000132298

0.000187048

0.001585112
0.000792556

0.000173209

0.000147845

0.000471502

0.018178239
0.006059413

0.000295402
0.000147701

0.000147701

0.000381964
0.000190982

0.000190982

0.005221462
0.010442924

0.00024813

0.000308596

0.00031546

0.00020647

0.000186808

0.000203148

0.00026744

0.000266704

0.000258592

0.00020537

0.000244362

0.000173399

0.000251603

0.000517163

0.000183949

0.000184775

0.000188273

0.00101122

0.000556684
0.000278342

0.000278342

0.000441852
0.000220926

0.000220926

0.000228645
0.00045729

0.000228645

0.00019705

0.001578014
0.006312056

0.004734042
0.001578014

0.00059829
0.00119658

0.000300531

0.000297759

0.000203104
0.000406208

0.000203104

0.00025471
0.000127355

0.000127355

0.00129853
0.000649265

0.000197978

0.000259765

0.000191522

0.6727234723845
0.168202009046

0.5045214633385
0.168202009046

0.3221181366925
0.161059068346

0.000120529

0.000109905

0.000111417

0.000213433

0.000213884

0.000105673

2.49195e-07

0.000169869

0.00016956

0.000215191

0.000184829

0.000197299

8.73267e-05

0.000165843

9.69623e-05

0.000142067

0.00014308

0.00015218

0.000104826

0.000206778

0.00014775

0.000172478

0.000155872

7.65213e-05

0.000157063

0.000154821

0.000152127

0.000116257

0.00028102

9.41939e-05

0.000188366

0.000131475

0.000220181

0.000168415

0.000141234

9.23198e-05

0.000174246

0.000114067

0.000130861

0.000131514

0.000107092

0.00024786

0.000146402

0.000184879

0.000134999

0.000203623

0.000239238

0.000132658

0.000984793

0.000101534

0.000179557

8.85609e-05

0.000211949

0.000102649

0.000164985

0.000189547

0.000179183

6.21118e-05

0.000112725

0.000125936

0.000228746

0.0001036

0.000116753

0.000202632

9.36342e-05

0.000233844

0.000108356

0.000148335

0.000239789

0.000109433

0.000128361

9.76379e-05

0.000378238

0.000207543

0.000180214

0.000208985

0.000174704

6.75758e-05

0.000117861

5.91869e-07

8.5938e-05

0.000185342

0.00121569

4.33445e-07

0.000102423

0.000100265

0.000157987

0.000133783

0.000136866

0.000283361

0.000150683

0.000160459

7.91254e-05

0.000266359

0.000107423

0.000163485

0.00027375

0.00024696

2.07662e-07

8.1111e-05

0.000172296

0.000182968

0.000189953

0.000135457

9.84456e-05

0.000105668

0.000223526

0.000145578

0.000105168

0.000153669

0.000165917

8.48171e-05

0.000113218

7.11045e-05

0.000167528

0.000158951

0.000224886

0.000160407

0.000110583

0.00011855

8.32038e-05

0.000461061

0.000152421

0.000157154

0.000168835

0.000158426

0.000197166

0.000121316

0.000157625

0.000192032

6.58734e-05

2.44031e-07

0.000202526

0.000134346

0.000108719

7.79089e-05

0.000208615

0.000135108

0.000247447

0.000102493

0.000154918

1.46925e-08

0.000127418

9.26767e-05

0.000189201

0.000133206

0.000154718

9.7903e-05

0.000115199

0.000175775

0.00032934

0.000222004

0.000122488

0.000161789

0.000106073

0.000228503

0.000148982

0.000146676

0.00020308

0.00015876

8.91606e-05

0.000163785

0.000109361

0.000182385

0.000179499

0.000337482

0.000131599

0.00017606

0.00027688

0.000162737

0.000223473

0.000304716

0.000174053

1.27792e-08

7.86323e-05

0.000144679

0.000138209

0.000116883

0.000178523

0.000138946

0.00020374

0.00011669

0.00072225

0.00072225

0.000116479

0.000138121

0.000111931

0.000120603

0.000194252

0.000220335

0.000114913

0.000101079

0.000168172

0.000187721

0.000174533

0.000200444

0.000124735

0.000154646

0.000139442

0.00014189

0.000182747

0.000111541

0.000155897

0.000177972

7.84596e-05

0.000741313

0.000154626

8.72353e-05

0.000143931

9.4197e-05

0.000157992

0.000117278

8.2595e-05

0.000145376

0.000307737

0.000191407

4.46624e-09

0.000123809

0.000119874

0.000132025

0.000110481

0.000196121

0.000109511

6.70409e-06

9.25173e-05

0.000215759

0.000254694

0.000128467

0.000180134

8.71408e-05

0.000224406

0.000101224

8.91177e-05

0.000143215

6.27036e-05

0.000179062

0.000159104

0.000178085

8.01682e-05

0.000312739

0.000189675

0.000148881

0.000161689

0.000172321

0.000145078

0.000170532

0.000220174

0.000183588

0.000238167

0.000500804

0.000174909

0.000151077

6.83156e-05

0.000181356

0.00017148

0.000135798

6.8737e-05

0.000112634

0.000126768

0.000210073

0.000112227

8.19968e-05

0.000154575

0.000113913

0.000144448

0.000155231

0.000113121

0.00012969

0.000138636

9.44417e-05

0.000105081

0.000121371

0.000679275

0.000453755

0.000129624

0.000180202

0.000122571

0.000168921

0.000270281

0.000204054

0.000144816

0.000120303

9.66238e-05

0.000199326

0.000141489

0.000176398

0.000148805

0.00112762

3.41702e-07

0.000193949

0.000111049

0.000312201

0.000110604

0.00030011

0.000160505

3.93555e-09

0.000210397

0.000109013

0.000115187

0.000797866

0.000163593

9.51701e-05

9.81474e-05

0.000116023

0.000391187

0.000133525

0.000100649

0.000178915

0.000130747

8.85555e-05

0.000138789

7.2739e-05

0.000181392

9.13216e-05

0.000109735

0.000151141

0.000129696

0.000118624

0.000106001

0.000217853

0.000175948

9.46571e-05

9.82599e-05

0.000168559

0.000122396

9.65485e-05

0.000198345

0.000103289

9.90758e-05

0.000213747

0.000144677

0.000141044

0.000206562

0.00013558

0.00015924

0.000173966

0.000209469

0.000114573

0.000318398

0.000148124

0.000141972

0.000105539

0.000190692

0.000110803

0.000398533

0.000184833

0.000122297

0.000190158

0.0012602311

0.000910138

9.20748e-05

8.00993e-05

9.22468e-05

8.56722e-05

5.11978e-07

0.000128347

0.000368091

0.000164945

0.000180166

0.000149865

9.45924e-05

0.000168935

0.000132397

0.000205115

0.000144562

0.000109082

5.66405e-05

0.000195611

9.93573e-05

0.00010923

0.000157484

9.44686e-05

0.000155002

0.000126548

0.000246796

0.000117252

7.77448e-05

0.000521494

0.000101876

0.000141696

0.000162328

7.26921e-05

0.000361309

0.000159561

0.000113995

0.00015177

0.000247094

0.000107092

0.000135821

0.00212129

0.000205059

0.000142643

0.000158831

0.000211841

0.000143252

7.14393e-05

0.000225472

0.000102898

0.000166262

0.000115384

9.74181e-05

0.000203764

0.000129575

0.00026648

0.000179684

0.000171599

0.000167291

0.000167474

0.000284983

0.000152406

0.000155888

0.000150368

0.000140653

0.000165531

0.000180276

0.000203517

0.000214876

0.000178484

7.26935e-05

0.000131191

0.000212876

0.000128577

0.000181691

0.000121141

0.000194377

0.000207643

7.29007e-05

8.08809e-05

6.92715e-05

0.000187035

0.000502404

0.000297088

0.000117977

0.000127152

0.00028004

0.000299429

0.00860289

0.000115018

0.00017125

0.000180582

0.000616028

0.000174888

0.000113881

8.3065e-08

0.000381944

0.000182557

0.000111573

0.000107164

9.78809e-05

0.000325205

0.000182934

0.000108539

0.000149406

0.000114803

8.38402e-05

0.000203608

8.3336e-05

0.00013191

0.000140377

0.000161991

0.000162915

8.85692e-05

0.000103143

0.000100924

0.000982149

0.000115107

0.000392434

0.000203748

0.000154266

3.762e-08

0.000168616

0.000133047

0.000135392

0.000188705

8.43036e-05

0.000312988

0.000692313

0.000407871

8.31997e-05

0.000172795

0.000132587

0.000149884

0.00016901

0.000259129

0.000178944

9.3448e-05

0.000210122

0.000132739

0.000106332

0.000332056

0.000214886

0.000260392

0.000193961

0.000240779

0.000479946

0.000178848

0.000130384

0.000151432

0.000152164

0.000155365

0.001006

8.59241e-05

0.000122292

0.000113201

0.000275303

0.000458015

8.30151e-05

0.000179507

0.000136411

0.000109194

0.00026672

0.000420218

0.000250485

0.000195304

0.000181341

0.000162776

0.000131561

0.000164963

8.52975e-05

0.000152692

8.17656e-05

0.000161602

0.000114224

0.00017914

0.000176699

0.000208362

0.000151122

6.79882e-05

0.000492339

0.000212662

9.52846e-05

0.000131079

0.000149422

7.91465e-05

0.000171745

7.63664e-05

0.000217416

0.000194728

6.34497e-05

0.000182609

0.000126066

0.000111342

9.89562e-05

0.00011103

0.000261612

8.24262e-05

0.00018992

0.00018867

0.000341909

0.000145819

2.63039e-08

9.63969e-05

0.000127966

0.000698109

0.000133785

0.000207499

0.000171599

4.77971e-06

0.000187467

8.71313e-05

0.000283842

0.000189782

0.000158411

0.000107144

0.000114806

9.16942e-05

0.000170736

0.00010908

0.000136341

0.000151

0.000273497

0.000106056

0.000144851

9.03753e-05

0.000205114

0.000105621

8.38553e-05

0.000122727

0.000130137

0.000230809

0.000182721

0.000194378

0.000105727

0.000109858

0.000124812

0.000171979

8.74771e-05

0.000126406

0.000215541

7.72359e-05

0.000103222

0.000227248

0.000140643

0.0056658

0.000250631

0.000138699

9.48417e-05

0.00015994

0.000155978

9.52999e-05

0.000235615

0.000139125

0.000150112

6.16025e-05

0.000175704

9.44505e-05

0.00019784

0.000435217

1.99381e-07

0.000311102

0.000105332

9.93678e-05

0.000153073

0.000123481

0.000239287

0.000153471

8.19282e-05

9.72691e-05

0.000384741

0.000123485

0.00015983

0.000297336

0.000140466

7.79628e-05

0.000139172

8.55091e-05

0.000147882

8.20025e-05

9.68791e-05

0.000138237

0.000125932

8.44509e-05

0.000288998

9.93872e-05

0.000152192

0.000277284

0.000114412

0.000100983

0.000156077

0.000152291

5.45638e-05

8.68419e-05

0.000236227

0.000159454

0.00010872

0.000214919

0.000167546

0.000144348

0.000269042

0.000182288

0.000174027

7.76534e-05

0.00011178

0.000218407

0.000107553

0.00021041

0.000173271

0.000190184

0.000102944

1.25914e-08

0.000163289

0.000244741

0.000226392

0.000331776

0.000155146

0.000146044

7.671e-05

0.00035291

0.000167704

8.93682e-05

7.22285e-05

0.000142045

0.000483691

0.000486205

0.000194928

0.000184758

0.00013882

0.000548476

0.00014438

0.000145115

0.000141453

0.000193749

0.000146481

0.000196274

6.86971e-09

0.000277551

0.000408545

0.000168466

5.3763e-06

0.000261197

0.000105043

0.000136907

0.000115503

0.000164143

0.000220729

0.000247607

0.000153592

0.000123571

0.000191288

0.000107219

8.27414e-05

0.000154747

0.000236956

0.000108909

0.000156547

0.000154034

0.000226829

0.000216688

8.7166e-05

0.000164674

0.00016463

0.000253976

0.000104191

8.50727e-05

0.000117748

0.000158484

0.000417307

0.000228739

0.000207731

0.000208738

0.000194921

0.000157651

0.000140394

0.000123401

0.0001834

0.000197187

0.000115986

6.64046e-05

0.00019586

8.88357e-05

0.000105555

0.00020104

0.000230128

0.000293731

0.000103897

0.000107046

8.46447e-05

0.000125007

0.000139114

0.000175997

0.000302331

0.000208415

0.000271002

0.00131073

0.000149341

0.000101255

0.000172116

0.000126569

9.8722e-05

0.000107272

0.000269258

0.000248692

7.4082e-05

0.000135037

0.000191397

0.000225264

0.000100429

0.000157328

0.000259796

0.003042932

9.77125e-05

0.000125664

8.65124e-05

0.000128826

7.19218e-05

9.98524e-05

0.000191473

6.98708e-05

0.000100736

0.00121776

9.723e-05

0.000125249

8.49574e-05

8.1995e-05

6.90981e-05

0.000303799

9.02746e-05

0.000241915

0.000214696

0.000812071

0.000171509

7.72894e-05

0.000117557

0.000218387

0.000141657

0.000134401

0.000253758

0.000277599

0.000135375

9.95243e-05

9.18041e-05

6.94216e-05

0.000139637

0.000108138

8.46894e-05

0.000120862

0.000126659

0.000165481

0.00552638

0.000139639

0.00016679

0.000112457

0.000126489

0.000177851

8.31438e-05

5.6506e-06

4.89936e-06

0.000142618

8.75911e-05

6.45659e-05

0.000193285

7.84404e-05

0.000181368

7.85898e-05

0.00229743
0.00459486

0.000167039

0.000241791

0.000200044

0.000341454

0.000430332

0.000222221

0.000251272

0.00018604

0.000257237

8.45638e-05

0.0095218938
0.0047609469

0.000239564

0.000160286

0.000133557

0.000276593

0.0002293

0.000197343

0.000185997

0.000389863

0.000262532

0.00019827

0.000277761

0.000285304

0.000146494

0.000372184

6.98179e-05

0.000197207

0.000224077

0.000229024

0.000512181

0.000173592

9.52299e-05

0.325873201119
1.303038915476

0.00014915
0.0002983

0.00014915

0.3719578653
0.1239859551

0.0100008572
0.0050004286

4.14752e-05

0.00369873

0.00010436

4.34858e-05

4.02042e-05

4.06429e-05

3.70099e-05

0.000131957

4.14146e-05

3.73067e-05

3.89522e-05

5.62712e-05

4.02732e-05

4.25413e-05

4.15281e-05

0.00014386

0.000215842

3.53268e-05

3.92125e-05

0.000130035

0.0028948656
0.0014474328

8.07224e-05

5.69397e-05

6.97744e-05

0.000245604

9.55007e-05

5.61433e-05

7.3657e-05

8.12416e-05

0.000337782

6.7586e-05

5.88068e-05

5.71971e-05

5.67368e-05

0.000109741

0.0116055156
0.0232110312

0.000134988

0.000247516

0.000135345

6.59568e-05

0.000252357

9.62728e-05

0.000152629

0.000240034

0.000170444

0.000132732

0.000827936

0.000300905

6.87835e-05

0.000752981

7.67139e-05

8.90063e-05

0.000182172

0.000202162

6.5443e-05

7.04563e-05

0.000146308

0.00014238

0.000160836

0.000136027

0.000702013

0.00012835

0.000277659

0.000158717

0.000385408

0.000185308

0.0001007

0.000197497

0.000102185

0.000130057

8.44271e-05

5.4801e-05

6.65171e-05

0.000180842

7.6614e-05

0.000137788

0.000138637

9.33466e-05

0.00117202

0.000330899

9.6396e-05

0.000150669

0.000159075

8.25475e-05

0.000216957

9.13881e-05

0.000304643

8.55217e-05

8.61376e-05

0.000131522

5.80093e-05

0.000136928

4.98825e-05

0.000107448

0.000134959

8.18297e-05

7.74318e-05

0.000328008
0.000656016

0.000103344

0.000224664

0.2109385448
0.1054692724

6.43499e-05

0.000127461

8.6897e-05

4.91735e-05

5.44187e-05

0.0022265224

9.8025e-05

0.000212069

5.80502e-05

0.000109835

0.000161062

0.000140691

9.7688e-05

7.58379e-05

5.26845e-05

8.58027e-05

0.000134365

0.000228342

7.46401e-05

0.00010497

0.000108363

6.90329e-05

0.00026629

6.37278e-05

8.50463e-05

4.30486e-05

7.56575e-05

0.000141468

5.90119e-05

9.82614e-05

0.000183082

9.36908e-05

0.000103568

5.53823e-05

6.55668e-05

6.54194e-05

7.76846e-05

5.48967e-05

0.0725473293

9.39493e-05

0.0716641

3.13416e-05

7.36568e-05

3.44943e-05

5.48629e-05

4.41573e-05

7.54492e-05

3.44069e-05

3.21519e-05

3.38189e-05

0.000245564

3.14681e-05

9.79081e-05

5.70661e-05

5.68839e-05

0.000577175

7.38188e-05

8.42552e-05

4.64292e-05

8.9161e-05

9.34131e-05

6.63136e-05

0.000126473

8.97223e-05

5.5676e-05

5.04338e-05

8.03747e-05

0.000173401

2.8961e-05

0.000124792

5.90119e-05

9.82908e-05

4.62124e-05

6.85755e-05

7.29499e-05

5.97423e-05

0.000615111

0.000156113

7.65368e-05

7.08117e-05

4.25899e-05

7.48946e-05

5.67533e-05

7.39304e-05

6.0917e-05

0.000207624

4.58597e-05

8.20981e-05

4.74764e-05

5.6066e-05

5.31281e-05

8.22819e-05

9.66862e-05

5.68215e-05

7.27548e-05

0.000312856

8.732e-05

4.69827e-05

7.01085e-05

7.68763e-05

8.56319e-05

7.18143e-05

4.37022e-05

8.82362e-05

0.00288744

0.000101888

0.00014483

4.85998e-05

0.00010523

4.38511e-05

0.000114836

5.33611e-05

4.58035e-05

4.83404e-05

5.47679e-05

6.61423e-05

0.000181458

5.24691e-05

6.3191e-05

3.83054e-05

6.33475e-05

0.000283653

0.000112241

5.31423e-05

5.70852e-05

0.000131727

8.33707e-05

4.38078e-05

0.000150446

0.000106051

9.90229e-05

8.84431e-05

5.7899e-05

4.76032e-05

0.00011808

8.15345e-05

5.76294e-05

4.61345e-05

0.000886495

9.05291e-05

8.4221e-05

5.4438e-05

9.23444e-05

6.74369e-05

6.56187e-05

9.38557e-05

8.23146e-05

8.50951e-05

9.77947e-05

4.69088e-05

8.36139e-05

9.94166e-05

5.68882e-05

5.17875e-05

4.60215e-05

5.74791e-05

8.59366e-05

4.10678e-05

8.84748e-05

7.81145e-05

0.000112338

4.99906e-05

8.22763e-05

9.80916e-05

7.44937e-05

8.61878e-05

9.46764e-05

0.000117012

5.50035e-05

6.50023e-05

5.77177e-05

4.43269e-05

5.89373e-05

9.43957e-05

5.11056e-05

5.77309e-05

7.2662e-05

4.9743e-05

5.58379e-05

6.62781e-05

0.000605878

5.18615e-05

7.26763e-05

0.00016764

8.30031e-05

8.3686e-05

7.63701e-05

0.000497198

6.57045e-05

4.6101e-05

0.00304732

0.000103197

6.93543e-05

5.95815e-05

9.69219e-05

5.22257e-05

5.61223e-05

0.000141843

5.32703e-05

0.000101834

8.93576e-05

0.000111838

4.90613e-05

6.08312e-05

3.06739e-05

0.000126323

6.54103e-05

6.24201e-05

0.00011277

4.6812e-05

5.81605e-05

8.26471e-05

5.03541e-05

6.28239e-05

7.80266e-05

0.000105806

6.68751e-05

8.33475e-05

6.16235e-05

6.00368e-05

0.000112416

4.97753e-05

4.96858e-05

0.000133748

0.000144142

9.64222e-05

6.22508e-05

8.42054e-05

0.004339621

0.000451867

0.000552097

0.00236813

8.12084e-05

6.74963e-05

0.000251273

0.000161565

7.76324e-05

5.12646e-05

7.52947e-05

5.82956e-05

0.000143497

7.31543e-05

5.87601e-05

5.81064e-05

7.27285e-05

0.000105166

9.66597e-05

5.65561e-05

5.18412e-05

5.24434e-05

9.87861e-05

0.000105496

6.1817e-05

8.98689e-05

4.76735e-05

5.82618e-05

0.000289628

6.3343e-05

7.07341e-05

5.5525e-05

5.95382e-05

9.08078e-05

9.13532e-05

8.50885e-05

5.22242e-05

5.89057e-05

5.32732e-05

7.84812e-05

0.000116249

7.21525e-05

0.0002705954
0.0001352977

5.81669e-05

7.71308e-05

0.001310399
0.003931197

0.001310399
0.002620798

0.00032476

0.000154879

0.000113849

0.00037503

0.000341881

0.000287364
0.000143682

0.000143682

0.0084871986
0.0254615958

0.000256586
0.000128293

0.000128293

0.000211376
0.000105688

0.000105688

0.0165064352
0.0082532176

8.30256e-05

0.000202154

0.000624861

0.000102535

7.97041e-05

0.00012458

0.000267391

0.000120377

0.000190291

9.63621e-05

0.000851177

0.000121142

0.00018343

0.000106285

0.000108484

0.000100227

0.000208139

0.000101836

7.57313e-05

0.00026678

0.000175967

0.000109884

8.64393e-05

8.78526e-05

0.000484658

0.000163021

7.68626e-05

8.22531e-05

7.71059e-05

6.4313e-05

8.72942e-05

7.3922e-05

0.00015142

7.9686e-05

0.000110431

0.00010717

0.000116473

8.97348e-05

0.000169397

0.00016926

0.000157694

0.000133836

0.000124725

9.9283e-05

0.000142138

0.000484658

0.000533227

0.0024487443
0.0008162481

0.0016324962
0.0008162481

9.87209e-05

8.97685e-05

9.26283e-05

0.000163866

8.04167e-05

8.93355e-05

7.95662e-05

0.000121946

0.006450642
0.002150214

0.004300428
0.002150214

0.000148257

0.00014977

0.000320662

0.000161264

0.000138338

0.000148442

0.000145421

0.00012668

0.000133035

0.000185885

0.000107526

9.8897e-05

0.000125004

0.000161033

0.037045884311
0.111137652933

0.01413382603
0.02826765206

0.00102672

0.000251152

7.26919e-05

0.000198746

8.22737e-05

0.000199012

0.000106042

0.000141892

0.00039937

8.39444e-05

0.000117819

0.000116597

0.000121444

0.000150627

0.000111157

0.000246483

0.000103303

0.000130045

0.000255518

0.000109912

7.10556e-05

0.00012517

7.74717e-05

6.62513e-05

8.09032e-05

7.54622e-05

0.000277258

0.000112464

0.000128829

0.000149273

0.000861028

8.78524e-05

0.000127028

8.44469e-05

0.000115914

0.000114309

0.000133391

0.000158041

8.60686e-05

0.000293327

0.000214863

0.000116273

0.000202347

0.000191204

7.01928e-05

0.000123147

8.51285e-05

0.000132101

9.38145e-05

8.48289e-05

0.00018249

8.58877e-05

0.000103418

0.000115171

0.000118946

0.000138082

9.76352e-05

0.00015383

0.000123517

9.4089e-05

0.000798281

0.000143936

0.00014568

1.6613e-07

7.66494e-05

7.38061e-05

9.48339e-05

0.000157492

7.3125e-05

6.39116e-05

7.22524e-05

0.000177433

9.20791e-05

0.000180174

0.000143292

7.17649e-05

0.000113963

7.43856e-05

0.00018513

0.000425873

9.30089e-05

9.79982e-05

0.000101163

0.000108733

0.000110705

9.28201e-05

8.23492e-05

7.39871e-05

0.000103911

7.27186e-05

7.76783e-05

0.000177267

0.022736690281
0.045473380562

4.65605e-05

4.69489e-05

9.13978e-05

6.78303e-05

5.71027e-05

7.1469e-05

5.20878e-05

0.000116483

7.45819e-05

0.000142838

0.000143591

5.10199e-05

4.85266e-05

5.54779e-05

9.49937e-05

0.000150009

6.4101e-05

0.0001093

0.00280878

4.51202e-05

0.000140997

0.00149507

7.87087e-05

7.2579e-05

7.08807e-05

5.29983e-05

0.000153857

5.03644e-05

8.96773e-05

7.88145e-05

0.000192738

5.46974e-05

5.4151e-05

0.000128504

5.71121e-05

7.88828e-05

4.73021e-05

6.53274e-05

0.00198728

9.24786e-05

6.89207e-05

5.31847e-05

0.000271439

4.70649e-05

9.74183e-05

5.07969e-05

0.00029099

0.000117556

0.000166209

4.97989e-05

6.26823e-05

5.52546e-05

5.00602e-05

8.51498e-05

4.62769e-05

9.87312e-05

6.17481e-05

8.51097e-05

6.17402e-05

7.29079e-05

8.68179e-05

0.00126843

7.61413e-05

1.4701e-07

0.00111138

9.8427e-05

4.86844e-05

8.17355e-05

6.36041e-05

5.31634e-05

7.0291e-05

8.91368e-05

4.74501e-05

0.000111304

7.00632e-05

6.04572e-05

4.38774e-05

0.000116367

0.000109173

9.08996e-05

0.000215428

0.00010498

5.14081e-05

0.000716217

5.79618e-05

4.61474e-05

9.99751e-05

0.000142967

7.44109e-05

4.69947e-05

0.000127261

0.00030798

6.90114e-05

4.7021e-05

0.000300156

0.000159422

6.16157e-05

6.10634e-05

0.000103925

9.32292e-05

4.27213e-05

7.05349e-05

4.43083e-05

0.00010242

7.89918e-05

9.1433e-05

4.68356e-05

0.000115651

4.79027e-05

4.63531e-05

4.62974e-05

6.76991e-05

9.59255e-05

0.000222714

6.09524e-05

4.98915e-05

4.64753e-05

5.12816e-05

0.000341216

8.01935e-05

5.74824e-05

8.06273e-05

0.00048757

5.77273e-05

8.70998e-05

5.50608e-05

5.94689e-05

6.60762e-05

6.0179e-05

9.09921e-05

0.000122435

5.32664e-05

8.79459e-05

4.70252e-05

0.00011987

4.41324e-05

4.80681e-05

6.81063e-05

6.46915e-05

0.000125968

6.54889e-05

4.14559e-05

0.000144096

4.18922e-05

5.66036e-05

6.28445e-05

5.19097e-05

5.50729e-05

0.000104122

6.8725e-05

8.84779e-05

4.8877e-05

6.66848e-05

6.64522e-05

0.000152408

4.88133e-05

8.03662e-05

4.06171e-07

0.000175368
0.000350736

0.000175368

0.0002375825
0.0007127475

0.0002375825
0.000475165

0.00014227

9.53125e-05

0.454479605524
0.151546887508

0.0008685156
0.0017370312

9.07841e-05

8.53125e-05

0.000105131

0.000337852

8.38913e-05

8.10009e-05

8.45438e-05

0.150517314908
0.301034629816

9.36019e-05

0.000149227

7.29303e-05

6.89816e-05

6.57664e-05

7.68624e-05

0.000206382

7.75232e-05

0.000128507

0.000188605

0.000113088

8.11547e-05

7.19016e-05

6.87203e-05

7.83793e-05

0.000523249

0.000117023

0.000107322

0.000178275

0.000209483

0.000114774

6.7398e-05

7.78798e-05

9.90724e-05

0.000112925

0.000171909

0.000177251

6.04276e-05

0.00035063

0.000117929

0.000265747

6.63019e-05

0.000113108

0.000187563

0.000102628

0.000140442

0.00015097

0.000122339

0.000201491

9.63028e-05

0.000114171

9.2954e-05

7.59499e-05

7.51904e-05

0.000124083

7.8897e-05

0.000111567

0.000147645

0.000184636

0.000258427

9.53944e-05

0.000211909

0.000276787

9.71329e-05

0.000100967

0.000425312

0.000257638

0.00107017

0.000139407

0.000123689

0.000228449

8.24698e-05

8.11217e-05

0.000219659

0.000432071

0.00015073

0.000102196

6.71128e-05

9.14649e-05

0.000283791

0.000110194

8.33811e-05

6.18928e-05

8.41106e-05

0.000144183

9.11447e-05

0.0005693

8.68093e-05

0.00021061

0.000129033

0.000489351

0.000100477

8.62146e-05

8.35039e-05

0.000127699

0.000105845

8.35209e-05

0.000118483

0.000101645

7.15817e-05

0.000810488

0.000105341

9.65202e-05

8.0141e-05

9.63823e-05

0.000181768

0.000110896

8.01837e-05

0.000274901

0.000153263

0.0001131

8.8129e-05

0.000103254

6.99329e-05

6.70809e-05

0.000276551

0.000609796

8.78435e-05

0.000148015

0.000528361

0.000390728

6.59836e-05

0.000209806

0.000188442

0.000192205

9.60026e-05

0.000432171

0.00012789

0.0667702

0.00139295

8.13454e-05

9.91123e-05

9.90191e-05

0.000120058

0.000106852

0.000112288

0.000144657

7.91489e-05

0.000117358

6.94858e-05

7.76063e-05

9.24935e-05

0.000212889

8.81058e-05

0.000116984

0.000157291

0.000192611

6.46415e-05

9.92564e-05

7.9148e-05

0.000133174

0.000105593

9.73883e-05

7.90636e-05

8.18881e-05

8.40288e-05

0.000164915

7.39714e-05

6.6288e-05

0.000204167

0.000127847

0.000243152

8.51441e-05

0.000259257

7.97548e-05

7.85971e-05

6.27254e-05

0.000432404

7.55195e-05

7.52271e-05

0.000153372

8.57238e-05

0.000120347

0.000168011

0.0451554

0.00032966

0.000124832

0.000173325

0.000121292

0.000121482

0.000163646

8.76297e-05

9.8095e-05

0.000129898

0.000359799

0.000207024

8.01466e-05

0.000227736

0.000123516

0.000134358

8.54208e-07

0.000331495

5.99389e-05

8.52922e-05

0.000154356

0.000109301

0.000230185

7.38833e-05

0.000257461

9.0247e-05

8.19194e-05

7.10261e-05

0.000121436

0.000174257

9.82393e-05

0.000188152

9.69269e-05

0.000109495

0.000338593

0.000188523

0.000134686

0.000712969

0.000101114

0.000187738

7.66429e-05

0.000129824

6.259e-05

6.66516e-05

0.000257357

0.000250245

7.61713e-05

7.35846e-05

0.000591107

8.46219e-05

0.000106991

0.000132692

0.00033334

0.000131812

5.59274e-05

0.000364405

9.09386e-05

9.20347e-05

0.00170387

0.000383593

0.00030671

0.000161057

8.90358e-05

0.001967146
0.0004917865

0.0014753595
0.0004917865

0.0001374145
0.000274829

4.06235e-05

5.63964e-05

4.03946e-05

0.000708744
0.000354372

4.96427e-05

6.0051e-05

5.27856e-05

6.69275e-05

6.81145e-05

5.68507e-05

0.001595259
0.006381036

0.001595259
0.004785777

0.000258274
0.000129137

0.000129137

0.001466122
0.002932244

0.00045081

0.00017875

0.000433404

0.000195966

0.000207192

0.000164991

7.18655524525099
1.79708995421

0.029346333
0.088038999

0.000382058
0.000191029

0.000191029

0.000917083
0.001834166

0.000295161

0.0001938

0.000174397

0.000253725

0.000594742
0.000297371

0.000297371

0.055263162
0.027631581

0.00112242

0.000191399

0.000206988

0.000237963

0.0212114

0.00025036

0.000197392

0.000191953

0.000247051

0.000224001

0.000157484

0.000188338

0.000239236

0.000224722

0.000379954

0.00037289

0.000224299

0.000257328

0.000175805

0.000653526

0.000290044

0.000189679

0.000197349

0.000309269
0.000618538

0.000309269

0.015380872119
0.005126957373

0.005126957373
0.010253914746

0.000113626

0.00010541

0.000463082

0.00020939

9.1365e-05

0.000498024

0.000122816

0.000282538

2.89173e-07

0.000536783

0.000124141

0.000121812

0.000123355

0.000130427

0.000168953

0.000137438

0.000155086

0.000128307

0.000116318

0.000110296

9.28257e-05

0.000301128

0.000625693

8.60355e-05

0.000152344

0.000129475

0.00028828
0.00014414

0.00014414

0.0603178637
0.1805950136

0.000225238
0.000112619

0.000112619

0.000795008
0.000397504

7.18974e-05

8.17179e-05

7.98504e-05

6.65489e-05

9.74894e-05

0.000103647
0.000207294

0.000103647

0.000333984
0.000166992

0.000166992

0.001087697
0.002175394

0.000140333

0.000107679

0.000145151

0.000179325

0.00013612

0.000232037

0.000147052

0.0002529
0.00012645

5.58517e-05

7.05983e-05

6.25975e-05
0.000125195

6.25975e-05

0.000155561

0.004323346
0.002161673

0.000243537

0.000134168

0.000163546

0.000244665

0.00101557

0.00014845

0.000211737

0.000237271
0.000474542

0.000120121

0.00011715

0.0001337852
6.68926e-05

6.68926e-05

7.20255e-05

0.004978274
0.009956548

5.97797e-05

8.95147e-05

5.44003e-05

7.68512e-05

5.46315e-05

6.64076e-05

0.000126287

5.76347e-05

5.53572e-05

5.75316e-05

6.64204e-05

0.000114056

6.77192e-05

7.28493e-05

7.29172e-05

5.87592e-05

5.34531e-05

5.61695e-05

6.69866e-05

5.15423e-05

5.78616e-05

5.87698e-05

5.9371e-05

5.5979e-05

5.2867e-05

0.000129719

5.89679e-05

5.57043e-05

5.24209e-05

5.75762e-05

5.47829e-05

5.60863e-05

5.41836e-05

5.79614e-05

0.00016449

6.10083e-05

5.32482e-05

5.69744e-05

5.41057e-05

9.57615e-05

7.60718e-05

5.91841e-05

5.72415e-05

8.09343e-05

0.000213343

5.57307e-05

7.97956e-05

0.000151976

0.000105307

0.000114917

5.70136e-05

0.000131896

0.000262486

6.78031e-05

5.3053e-05

5.99516e-05

5.55429e-05

5.57647e-05

6.30179e-05

5.93841e-05

0.000114496

9.89751e-05

5.77563e-05

5.7848e-05

6.16774e-05

0.0045449173
0.0090898346

8.93564e-05

0.000136167

0.000101408

0.000101738

0.000113111

0.000113894

0.000116286

0.000166437

0.000156687

0.000130192

0.000126238

0.000200507

0.000113054

0.000133405

0.000130521

0.000196793

0.000117924

0.000107774

0.000122915

0.000125849

0.000163128

0.000110116

9.19079e-05

0.000117316

0.000182359

0.000145544

0.000107443

0.000127643

0.000138762

0.000155014

0.000107107

0.000102277

0.000129916

0.000134475

0.000131653

0.0047608518
0.0023804259

6.45619e-05

7.92733e-05

8.27497e-05

6.82299e-05

8.34843e-05

0.000116129

7.24672e-05

9.93745e-05

0.000103765

6.65547e-05

7.67e-05

0.000100345

7.53817e-05

9.24599e-05

6.89875e-05

7.87347e-05

7.47288e-05

9.13387e-05

0.000108441

0.000117675

7.63755e-05

9.9908e-05

6.67899e-05

0.00010898

0.000106058

0.000102696

9.82367e-05

0.000223762
0.000111881

0.000111881

0.0010304114
0.0020608228

9.74007e-05

0.000134461

9.74007e-05

0.000159548

0.000113419

0.000139323

0.000186229

0.00010263

0.0255523515
0.051104703

0.00022504

9.76243e-05

0.000104073

0.000250468

8.81187e-05

6.87426e-05

0.000108313

0.000102759

7.07678e-05

0.000110489

7.80032e-05

0.000103141

0.000210674

0.000107955

0.000127767

8.81058e-05

9.61155e-05

9.81289e-05

0.000178647

9.96866e-05

0.000242119

0.000112347

0.000100067

9.48905e-05

0.000115604

0.000161544

9.67631e-05

0.00014728

9.26919e-05

0.000101251

0.00896973

0.000121567

9.06754e-05

8.42221e-05

0.00010907

0.00010136

0.000271676

0.000100524

0.000105748

0.000112286

0.000222863

0.000247676

9.68391e-05

0.000125279

0.000127147

0.00011486

9.63963e-05

9.82102e-05

0.00015138

7.46023e-05

0.000117548

0.000104573

0.000133189

0.000221255

9.19068e-05

0.000143831

0.000126712

0.000133393

8.9748e-05

0.000156095

7.20586e-05

0.000100525

0.000109382

0.00010752

0.000103823

0.00021653

7.19052e-05

0.00011905

0.00011047

0.00010326

7.07145e-05

9.00172e-05

8.87712e-05

0.000118847

0.000215945

0.0001095

0.000107806

0.000106358

0.000157494

0.000627809

0.000152592

0.000114074

0.000134168

0.00010904

0.000109133

7.65184e-05

8.91267e-05

8.20167e-05

8.50375e-05

0.000131284

9.54706e-05

0.000329017

0.000141476

0.000102312

0.000101817

8.56039e-05

0.000120295

0.000113675

9.24241e-05

0.000303193

0.000158842

9.09946e-05

0.000108371

8.08672e-05

9.74257e-05

0.000109235

0.000352734

8.49343e-05

0.000120942

0.000103108

8.62284e-05

9.14531e-05

9.32324e-05

8.82684e-05

0.000104757

0.000240261

0.000142334

9.59793e-05

0.000115288

0.000164807

0.000215832

9.41715e-05

8.56616e-05

0.000110189

8.87528e-05

0.000157524

9.18345e-05

0.000382725

0.0004107972
0.0008215944

0.000183509

0.000144258

8.30302e-05

0.000357555
0.00071511

0.000145355

0.0002122

0.0018445885
0.003689177

0.000188454

9.86958e-05

0.000149179

0.000114336

0.000130273

0.000104278

0.000152441

0.000147619

9.95417e-05

0.000154463

0.000159618

0.000121039

0.000224651

0.000366186
0.000732372

0.000180407

0.000185779

0.0011991562
0.0005995781

6.31448e-05

0.00019715

8.59341e-05

8.42572e-05

0.000169092

0.0015624495
0.003124899

0.000148909

0.000105103

9.75682e-05

0.00011515

0.000153808

0.000106506

0.000150797

0.000127016

0.000117303

9.8153e-05

9.64803e-05

0.000245656

0.0057443276
0.0028721638

8.5808e-05

8.51318e-05

0.000370122

0.000127138

0.000143884

0.00012265

0.000187526

0.000115182

8.53142e-05

0.00010045

0.000100966

8.65545e-05

8.74018e-05

0.000277762

0.000121899

0.000131139

9.29811e-05

9.34187e-05

0.000149052

8.74387e-05

0.000134869

8.5476e-05

0.0002529322
0.0001264661

5.97353e-05

6.67308e-05

0.000253954
0.000126977

0.000126977

0.0019638
0.0009819

0.0009819

0.0024857783
0.0049715566

0.00018001

0.000108555

8.29877e-05

0.000227485

0.000119882

7.45689e-05

0.000113254

8.25297e-05

0.000181957

0.000113107

0.000318006

0.000166

0.000185788

0.00013069

0.00018617

0.000214788

0.000484885
0.00096977

0.000323322

0.000161563

0.000118233
0.000236466

0.000118233

0.000130991

0.000258638
0.000129319

0.000129319

0.000131171
0.000262342

0.000131171

0.000250002
0.000125001

0.000125001

0.000127469
0.000254938

0.000127469

0.0001717645
0.000343529

7.73445e-05

9.442e-05

0.0031392498
0.0062784996

0.000120515

0.000112054

0.000166407

8.72455e-05

0.000481116

8.54242e-05

0.000127577

9.63919e-05

8.72455e-05

8.55267e-05

0.000205578

8.29115e-05

0.000124965

0.000101391

9.81614e-05

8.24045e-05

8.24154e-05

0.000108076

0.000370314

0.000124918

8.54152e-05

0.000135915

8.7282e-05

0.0006761502
0.0013523004

7.76707e-05

8.00671e-05

0.000106444

7.77671e-05

8.89192e-05

7.44803e-05

8.78673e-05

8.29345e-05

0.000252344
0.000504688

0.000252344

0.001180616
0.003541848

0.000276582
0.000138291

0.000138291

0.000159244
0.000318488

0.000159244

0.000222228
0.000444456

0.000222228

0.000581768
0.000290884

0.000290884

0.000369969
0.000739938

0.00020436

0.000165609

0.133521988
0.400351223

0.002757633
0.005515266

0.00120497

0.000191984

0.000213825

0.000354003

0.000246413

0.000165432

0.000210422

0.000170584

0.0645883
0.1291766

0.0645883

0.000442916
0.000221458

0.000221458

0.000214741

0.00150257
0.000751285

0.000246775

0.000208058

0.000296452

0.129977142
0.064988571

0.000164109

0.000212074

0.0641661

0.000167658

0.00027863

0.00152039
0.000574576

0.000203338

0.000203703
0.000407406

0.000203703

0.000167535
0.00033507

0.000167535

0.002063982
0.000687994

0.000498836
0.000249418

0.000249418

0.000202288
0.000404576

0.000202288

0.000236288
0.000472576

0.000236288

0.03173609
0.010632831

0.00237032
0.00118516

0.000194225

0.000285454

0.000145466

0.000121002

0.000139958

0.00014085

0.000158205

0.001551614
0.000775807

0.000168502

0.000166116

0.000249341

0.000191848

0.000146534
0.000293068

0.000146534

0.000968338
0.000484169

0.000166916

0.000317253

0.000130872
0.000261744

0.000130872

0.002682292
0.005364584

0.000164356

0.000369303

0.000201202

0.000200103

0.000141769

0.000245229

0.000751734

0.000228766

0.000146298

0.000233532

0.000923115
0.00184623

0.000923115

0.001327743
0.002655486

0.000126302

0.000235792

0.000117249

0.000142772

0.000168116

0.000161155

0.000229159

0.000147198

0.001045308
0.000522654

0.000169151

0.000155607

0.000197896

0.00155416
0.00077708

0.000133306

0.000164849

0.000121223

0.000172971

0.000184731

0.000388631
0.000777262

0.000181025

0.000207606

0.000162403

0.000937255
0.00187451

0.000135216

0.000144001

0.000126875

0.000178179

0.000171461

0.000181523

0.000189116
0.000378232

0.000189116

0.0391630311
0.1174890933

0.075307732
0.037653866

0.00014632

0.037088

0.000127942

0.000139975

0.000151629

0.000325909
0.000651818

0.000180392

0.000145517

0.0001747422
8.73711e-05

8.73711e-05

0.00219177
0.001095885

0.000194554

0.000505885

0.000207306

0.00018814

0.148254132
0.049418044

0.098836088
0.049418044

0.000174496

0.0477582

0.00026384

0.000197349

0.00020482

0.000253006

0.000214116

0.000352217

1.45596411583
4.367727833522

0.000511205
0.00102241

0.000406323

0.000104882

0.017563666306
0.035127332612

0.000129209

0.00011624

0.000104856

0.000109305

0.000124996

0.000118504

0.000132452

8.98538e-05

0.000131948

0.000121716

9.70037e-05

0.000129923

0.0001282

0.000169895

8.00161e-05

0.000187382

0.000130969

0.000369413

9.298e-05

0.000126636

0.000112635

0.000185771

0.000117164

0.000112061

0.000115688

2.7227e-07

0.000105295

0.000236044

2.9883e-07

9.88666e-05

9.35522e-05

0.00014924

8.75572e-05

0.000101161

0.000120187

0.000123406

0.000104439

8.84229e-05

0.000187893

0.000116462

8.97878e-05

4.26124e-07

0.000218282

0.000122013

1.32458e-06

9.59524e-05

0.000228497

0.000151107

0.000105349

0.000171307

0.000116043

0.000201408

8.40125e-05

0.000142875

0.000136624

0.000120469

9.63579e-05

4.23555e-07

0.000117961

0.000115044

0.00014492

9.88161e-05

0.000146517

0.000311145

0.000103926

9.75186e-05

0.000121569

2.53435e-07

8.9404e-05

9.87906e-05

0.000120458

0.000106821

9.60554e-05

9.99638e-05

0.000103835

0.000103153

0.000153213

0.000130725

9.57111e-05

0.000119858

9.35199e-05

0.000139723

0.000111629

0.000103829

0.000111416

0.000106113

0.000196519

0.000123009

0.000103738

0.00010655

0.000118656

0.00010884

0.00010061

0.000152873

0.000105812

0.000124055

0.000121457

0.00027515

0.000137458

9.5695e-05

0.000117934

0.000148823

9.04227e-05

0.000374721

0.000181901

0.000101432

8.99411e-05

0.000208574

0.000105261

0.000110824

8.6408e-05

0.000152103

0.000108159

9.30124e-05

8.93782e-05

0.000109676

0.000493572

0.000167292

0.000114548

8.80998e-05

0.000104209

0.000334664

7.45812e-07

0.000116254

0.000143409

0.000112746

0.000106802

9.36081e-05

0.00011113

0.000122673

9.58939e-05

0.000169838

9.27996e-05

0.000172663

9.47302e-05

0.000417652

9.92841e-05

0.000251134
0.000125567

0.000125567

0.000201844
0.000100922

0.000100922

0.0001613234
8.06617e-05

8.06617e-05

0.000606194
0.000303097

0.000175846

0.000127251

0.000959696
0.001919392

0.000102255

0.000283092

0.000109425

0.00011928

0.000111089

0.000118888

0.000115667

0.00049588
0.00024794

0.00024794

0.0016807646
0.0008403823

0.000107112

0.000115901

0.000129682

8.84233e-05

0.000140526

0.000122376

0.000136362

0.006780458
0.013560916

0.000187768

0.000279271

0.00108184

0.000180341

0.000281126

0.00017686

0.000224165

0.000302095

0.000407209

0.000216091

0.000202867

0.000406989

0.00029917

0.00056982

0.000749505

0.000168113

0.000596018

0.000183322

0.000267888

0.000657952
0.000328976

0.000181859

0.000147117

0.0001403184
7.01592e-05

7.01592e-05

0.000366335
0.00073267

0.000185566

0.000180769

0.000221862
0.000110931

0.000110931

0.000530249
0.001060498

0.000155499

0.000176276

0.000198474

2.2323378529
1.11616892645

0.00890988

0.00977462

0.0313624

0.0305399

0.0268441

0.00966479

0.000393584

0.054906

0.0253263

0.0302746

0.030263

4.77045e-06

0.0322709

0.0281391

0.0325996

0.0279374

0.00912256

0.0377211

0.0326511

0.00962313

0.0396425

0.0365809

0.177716

0.000303142

0.0284024

0.0314568

0.00986442

0.0326264

0.0254249

0.0113736

0.0264809

0.00960084

0.0101735

0.0319329

0.0222866

0.0223252

0.00915679

0.00936219

0.0346142

0.00933203

0.00873733

0.0196421

0.0308589

0.00994555

0.000111243
0.000222486

0.000111243

0.002417116
0.001208558

0.000121973

0.000219478

0.000629653

0.000125336

0.000112118

0.30873439589
0.61746879178

0.00015559

0.000133616

5.03914e-06

0.000166735

0.305086

7.27809e-06

0.000162191

0.000171143

0.0021785

1.67866e-06

0.000159308

0.000177116

0.000171707

0.000158494

0.000164514

0.000656233
0.001312466

0.000101045

0.000555188

7.06421e-05
0.0001412842

7.06421e-05

0.000116936

0.017273313
0.005757771

0.000238256
0.000119128

0.000119128

0.004662328
0.009324656

0.000205648

0.000206803

0.000210696

0.000191826

0.000187232

0.000238285

0.000178395

0.000225804

0.000163566

0.000263213

0.000240327

0.000145023

0.000180661

0.000222045

0.000212862

0.00140207

0.000187872

0.000300088
0.000150044

0.000150044

0.001652542
0.000826271

0.000134191

0.000167242

0.000267796

0.000135536

0.000121506

0.0040847351
0.0122542053

0.002104445
0.00420889

0.000155101

0.000169771

0.000158236

0.000264849

0.000282022

0.000364748

0.000235024

0.000158227

0.000157803

0.000158664

0.0019802901
0.0039605802

0.000107104

0.000133178

0.000114355

0.000141481

0.000140107

0.00015723

0.000185185

0.000176882

0.000123795

0.000119332

0.000148907

8.30581e-05

0.000109812

0.000134572

0.000105292

0.000359633
0.001078899

0.000359633
0.000719266

0.000113123

0.00024651

0.000369403
0.001108209

0.000369403
0.000738806

0.000369403

0.0020424792
0.0081699168

0.0061274376
0.0020424792

0.0001746644
8.73322e-05

8.73322e-05

0.000144611
0.000289222

0.000144611

0.001810536
0.003621072

0.000425735

0.000249074

0.000234384

0.000166865

0.000173213

0.000178804

0.000200727

0.000181734

5.96289e-05

0.00109479

0.0005023447
0.0022110934

6.9284e-05

0.000952085
0.000190417

0.000761668
0.000190417

0.000190417
0.000571251

0.000380834
0.000190417

0.000190417

7.11076e-05

0.00055592
0.000111184

0.000444736
0.000111184

0.000111184
0.000333552

0.000111184
0.000222368

0.000111184

6.03521e-05

0.0005016499
0.0027816362

5.70658e-05

0.0001682651
0.0008413255

0.0006730604
0.0001682651

0.0001682651
0.0005047953

0.0003365302
0.0001682651

0.000109501

5.87641e-05

0.001381595
0.000276319

0.001105276
0.000276319

0.000828957
0.000276319

0.000276319
0.000552638

0.000139866

0.000136453

3.3192e-08

9.60434e-05
0.0005762604

9.60434e-05
0.000480217

0.0003841736
9.60434e-05

9.60434e-05
0.0002881302

0.0001920868
9.60434e-05

9.60434e-05

0.000580362
9.6727e-05

0.000483635
9.6727e-05

0.000386908
9.6727e-05

9.6727e-05
0.000290181

9.6727e-05
0.000193454

9.6727e-05

0.000262837

1.9576e-08

0.001993461
0.0118781376

0.001993461
0.0098846766

0.0015126077
0.0059678024

0.0007477198
0.0022431594

0.0002076019
0.0004152038

5.2117e-05

5.37393e-05

5.0467e-05

5.12786e-05

0.0002104
0.0001052

0.0001052

0.000444148
0.000222074

4.33185e-05

4.33185e-05

4.33185e-05

4.88e-05

4.33185e-05

8.60921e-05
0.0001721842

8.60921e-05

0.0001267518
0.0002535036

5.3855e-05

7.28968e-05

0.0001652568
8.26284e-05

8.26284e-05

0.0006822595
0.0020467785

0.0002995014
0.0005990028

8.52694e-05

6.26309e-05

7.81936e-05

7.34075e-05

0.0007655162
0.0003827581

0.000101328

9.08397e-05

9.06642e-05

9.99262e-05

0.0004808533
0.0019234132

0.0004808533
0.0014425599

0.000385556
0.000771112

0.000103197

8.99905e-05

9.81764e-05

9.41921e-05

0.0001905946
9.52973e-05

9.52973e-05

0.041204196
0.006867366

0.006745457
0.033727285

0.006745457
0.026981828

0.006745457
0.020236371

0.013490914
0.006745457

0.000217745

0.000160409

0.00018334

0.000161127

0.000157205

0.000229978

0.00036043

0.000217237

0.000160092

0.000165736

0.000221703

0.000161337

0.000160074

0.000215348

0.000179413

0.00018483

0.000173307

0.000158228

0.0001593

0.000176575

0.000181973

0.000163057

0.00016847

0.000200072

0.000175864

0.000290465

0.00020181

0.000176213

0.000174941

0.000175942

0.00016883

0.000199376

0.000180496

0.000315616

0.000168918

0.000609545
0.000121909

0.000121909
0.000487636

0.000365727
0.000121909

0.000243818
0.000121909

0.000121909

4.88617e-09

2.10371e-07

3.0707e-09

1.9576e-08

0.000143855

0.1688917896466
0.0247484544996

4.37976e-06
2.18988e-05

1.751904e-05
4.37976e-06

4.37976e-06
1.313928e-05

4.37976e-06
8.75952e-06

4.37976e-06

9.918e-06
4.959e-06

4.959e-06

1.21404e-05
3.64212e-05

1.21404e-05
2.42808e-05

1.21404e-05

0.1365374627779
0.0229282206296

9.82218e-05
0.0001964436

7.4757e-05

2.34648e-05

9.1698e-05
1.83396e-05

7.33584e-05
1.83396e-05

5.50188e-05
1.83396e-05

1.83396e-05
3.66792e-05

1.83396e-05

9.3514e-05
1.87028e-05

1.87028e-05
7.48112e-05

1.87028e-05
5.61084e-05

1.87028e-05
3.74056e-05

1.87028e-05

0.0006944266475
0.0034721332375

0.00277770659
0.0006944266475

0.0020832799425
0.0006944266475

0.0001165962488
0.0002331924976

5.20488e-08

1.28683e-05

1.54301e-05

1.30785e-05

1.94754e-05

1.959e-05

1.72491e-05

1.88528e-05

6.09044e-05
3.04522e-05

1.66797e-05

1.37725e-05

0.0010947563974
0.0005473781987

3.93423e-05

1.02821e-05

2.5155e-05

3.69222e-05

1.36568e-05

1.02964e-05

1.8828e-05

1.34747e-05

1.60806e-05

1.52707e-05

1.35835e-05

1.30945e-08

1.65941e-05

1.45696e-05

1.11969e-05

1.7402e-05

1.41578e-05

1.21725e-05

1.28361e-05

2.37821e-08

1.22973e-05

2.10599e-05

1.72753e-05

1.25498e-05

1.22238e-05

1.53292e-05

4.49767e-05

1.04371e-05

1.84703e-05

1.6613e-07

1.08678e-05

1.71087e-05

9.06681e-06

1.92904e-05

1.4377e-05

2.37821e-08

0.017087794
0.0034175588

5.63841e-05
0.0002255364

0.0001691523
5.63841e-05

0.0001127682
5.63841e-05

2.71055e-05

2.92786e-05

0.0076294144
0.0019073536

0.0003979254
0.0001326418

0.0001326418
0.0002652836

3.20189e-05

5.47481e-05

4.58748e-05

6.91478e-05
0.0002074434

6.91478e-05
0.0001382956

4.62243e-05

2.29235e-05

0.0050169951
0.0016723317

8.3933e-05
4.19665e-05

4.19665e-05

0.0003275432
0.0001637716

3.81648e-05

4.87917e-05

5.18069e-05

2.50082e-05

0.0001767685
0.000353537

2.91434e-05

4.29299e-05

2.98424e-05

3.69606e-05

3.78922e-05

1.68364e-05
3.36728e-05

1.68364e-05

0.0002476416
0.0001238208

2.78854e-05

4.89818e-05

4.69536e-05

0.0022408282
0.0011204141

3.54809e-05

2.61202e-05

2.97608e-05

5.18814e-05

5.71072e-05

4.97505e-05

2.52339e-05

2.69661e-05

3.353e-05

3.595e-05

2.43602e-05

2.63417e-05

4.9947e-05

4.22266e-05

0.000273291

3.14162e-05

3.09323e-05

3.08241e-05

0.000117594

2.27748e-05

2.94607e-05

3.95596e-05

2.99049e-05

2.87538e-05
5.75076e-05

2.87538e-05

9.96969e-05
3.32323e-05

3.32323e-05
6.64646e-05

3.32323e-05

0.0058152844
0.0014538211

0.0026550429
0.0008850143

0.0001040274
5.20137e-05

2.20782e-05

2.99355e-05

0.0006213873
0.0012427746

3.67966e-05

2.64656e-05

3.49903e-05

3.03896e-05

2.72894e-05

2.90658e-05

0.000199582

0.000236808

0.0002081902
0.0001040951

4.04758e-05

6.36193e-05

9.34166e-05
4.67083e-05

4.67083e-05

6.73004e-05
3.36502e-05

3.36502e-05

2.71597e-05
5.43194e-05

2.71597e-05

0.0004298936
0.0012896808

0.0008597872
0.0004298936

0.000360344

3.21673e-05

3.73823e-05

0.0001003131
0.0003009393

3.18626e-05
1.59313e-05

1.59313e-05

5.00529e-05
0.0001001058

2.56607e-05

2.43922e-05

3.43289e-05
6.86578e-05

3.43289e-05

3.86001e-05
0.0001158003

3.86001e-05
7.72002e-05

3.86001e-05

0.0001168448
0.000584224

0.0001168448
0.0004673792

0.0001168448
0.0003505344

6.02364e-05
3.01182e-05

1.23012e-05

1.7817e-05

2.03607e-05
4.07214e-05

2.03607e-05

0.0001327318
6.63659e-05

1.55258e-05

1.20471e-05

2.25994e-05

1.61936e-05

0.0003351535
0.0016757675

0.0003351535
0.001340614

0.0001970871
0.0005912613

6.32006e-05
3.16003e-05

1.41538e-05

1.74465e-05

0.0003309736
0.0001654868

9.57413e-05

2.94321e-05

2.71484e-05

1.3165e-05

0.0004141992
0.0001380664

0.0002056974
0.0001028487

1.0651e-05

1.49932e-05

1.94236e-05

1.47601e-05

1.50385e-05

1.19695e-05

1.60128e-05

7.04354e-05
3.52177e-05

1.74875e-05

1.77302e-05

0.0015439587
0.0077197935

0.0061758348
0.0015439587

0.0045757212
0.0015252404

0.0001280308
0.0002560616

2.69427e-05

4.78819e-05

2.30556e-05

3.01506e-05

0.0003389519
0.0006779038

1.4491e-05

0.000296146

1.46755e-05

1.36394e-05

0.0014565694
0.0007282847

4.65105e-05

3.49375e-05

2.57628e-05

3.0944e-05

2.76035e-05

2.50047e-05

6.05234e-05

3.86835e-05

3.36892e-05

6.76816e-05

4.96266e-05

5.86318e-05

2.51216e-05

5.16775e-05

2.40836e-05

3.28125e-05

9.49904e-05

0.000329973
0.000659946

1.68663e-05

1.9761e-05

2.44917e-05

2.15355e-05

4.29586e-05

2.28437e-05

2.41163e-05

2.16746e-05

5.86544e-05

2.45093e-05

2.04159e-05

3.21457e-05

1.87183e-05
5.61549e-05

3.74366e-05
1.87183e-05

1.87183e-05

0.0810127169608001
0.0163089006322

0.00519281983215
0.0205433884286

8.52019e-05

0.0033658956
0.0011219652

0.0005004109
0.0010008218

0.000210361

0.000149321

3.80322e-05

6.34266e-05

3.92701e-05

0.000118244
0.000236488

0.000118244

0.0002378458
0.0001189229

5.87687e-05

6.01542e-05

4.28646e-05
8.57292e-05

4.28646e-05

0.0003997158
0.0001998579

2.76235e-05

3.41175e-05

6.08775e-05

2.24664e-05

5.4773e-05

1.9865e-05
3.973e-05

1.9865e-05

6.58764e-05
3.29382e-05

3.29382e-05

6.60584e-05
0.0001321168

6.60584e-05

4.56066e-05
2.28033e-05

2.28033e-05

0.0034736058
0.0011578686

0.0023157372
0.0011578686

5.25314e-05

5.88035e-05

3.15727e-05

0.000820401

7.34742e-05

3.26862e-05

8.83996e-05

0.00842586529645
0.00282778413215

0.000181049
0.000362098

0.000181049

8.87713e-05
0.0001775426

3.60802e-05

5.26911e-05

0.002042623
0.0010213115

7.11457e-05

0.000912494

3.76718e-05

0.0004110388
0.0002055194

0.000100268

5.14006e-05

5.38508e-05

0.0021454810643
0.00107274053215

3.51278e-05

0.000331762

9.01834e-05

2.52218e-05

2.50811e-05

2.95537e-05

2.6136e-05

1.97166e-05

1.43215e-09

4.8194e-05

2.42281e-05

2.55666e-05

1.96601e-05

0.000330361

4.19469e-05

0.0002098
0.0001049

0.0001049

0.0001920106
9.60053e-05

3.22405e-05

3.0922e-05

3.28428e-05

5.74871e-05

0.0308618744
0.0077766669

0.0026439482
0.0077446386

0.0002232767
0.0004465534

3.92254e-05

6.79866e-05

5.37554e-05

6.23093e-05

0.000187206

1.92486e-05
3.84972e-05

1.92486e-05

0.0002328452
0.0004656904

6.36841e-05

8.14558e-05

3.0103e-05

5.76023e-05

0.000105448
5.2724e-05

5.2724e-05

0.0001776083
0.0003552166

5.95233e-05

0.000118085

0.0002955246
0.0001477623

9.37482e-05

5.40141e-05

0.0005441906
0.0002720953

3.49223e-05

0.000110942

0.000126231

0.0013311818
0.0026623636

3.79727e-05

3.90527e-05

4.09848e-05

0.000276068

5.95409e-05

6.4367e-05

4.11018e-05

4.88656e-05

4.51949e-05

5.1826e-05

9.02123e-05

5.18292e-05

6.32764e-05

5.58666e-05

0.000114246

2.98153e-05

7.43528e-05

6.60281e-05

3.14913e-05

4.90894e-05

0.0051039251
0.0153117753

0.000107677
0.000215354

0.000107677

7.63471e-05
0.0001526942

7.63471e-05

6.88083e-05
0.0001376166

6.88083e-05

9.18167e-05
0.0001836334

9.18167e-05

0.0001487656
0.0002975312

0.000112964

3.58016e-05

6.62542e-05
0.0001325084

6.62542e-05

0.0004155016
0.0008310032

7.73549e-05

6.68387e-05

8.0905e-05

0.000190403

0.0041287546
0.0082575092

4.19192e-05

3.84558e-05

0.000145099

5.78789e-05

5.84027e-05

8.86402e-05

4.26734e-05

7.32595e-05

6.30572e-05

8.65259e-05

4.19256e-05

7.08611e-05

5.95833e-05

5.41188e-05

6.85701e-05

3.41536e-05

5.39205e-05

8.42766e-05

0.00162538

7.22787e-05

0.000105205

3.52838e-05

5.68548e-05

5.1555e-05

5.90002e-05

7.2037e-05

6.00921e-05

0.000152957

6.66792e-05

7.01033e-05

0.000131156

7.00591e-05

9.55711e-05

5.75322e-05

8.56988e-05

9.79899e-05

2.87936e-05

0.0132985535
0.0033394139

0.0099591396
0.0033394139

0.0004342128
0.0002171064

8.43291e-05

6.08697e-05

7.19076e-05

0.000203372
0.000101686

3.32583e-05

2.68123e-05

4.16154e-05

0.0002602109
0.0005204218

6.14058e-05

8.75576e-05

5.61946e-05

5.50529e-05

0.0002282258
0.0004564516

7.08649e-05

8.49894e-05

7.23715e-05

5.91021e-05

5.93752e-05
0.0001187504

5.93752e-05

0.0002298316
0.0001149158

6.49952e-05

4.99206e-05

0.000162491
0.000324982

1.7706e-05

0.000144785

5.10672e-05
0.0001021344

5.10672e-05

0.0006261484
0.0003130742

3.21775e-05

3.77532e-05

4.32633e-05

4.25279e-05

5.06723e-05

2.64463e-05

8.02337e-05

0.0013837966
0.0006918983

5.96845e-05

0.00011394

5.74206e-05

7.5713e-05

3.60697e-05

9.67612e-05

4.60695e-05

0.000177041

2.91988e-05

0.0005658012
0.0002829006

7.49958e-05

7.94366e-05

4.3581e-05

3.12581e-05

2.6243e-05

2.73861e-05

0.0015947208
0.0007973604

3.46405e-05

3.21508e-05

3.06136e-05

0.000404578

5.5528e-05

3.85031e-05

6.78397e-05

4.04879e-05

3.15081e-05

6.15107e-05

0.00167515735
0.00037611335

0.00047388
0.00013581615

3.46923e-05

0.00017302752
5.767584e-05

2.695094e-05
5.390188e-05

1.0132e-05

9.87465e-06

6.94429e-06

3.07249e-05
6.14498e-05

1.49169e-05

1.5808e-05

1.807841e-05
5.423523e-05

1.807841e-05
3.615682e-05

7.69651e-06

1.03819e-05

1.574168e-05
4.722504e-05

3.148336e-05
1.574168e-05

7.19916e-06

8.54252e-06

2.888376e-05
9.62792e-06

1.925584e-05
9.62792e-06

9.62792e-06

4.53416e-05

0.0007798224
0.0001949556

0.0001949556
0.0005848668

0.000172408
8.6204e-05

4.14259e-05

4.47781e-05

0.0001634978
8.17489e-05

4.20216e-05

3.97273e-05

5.40054e-05
2.70027e-05

2.70027e-05

0.00121007803
0.00400557429

6.89618e-05
3.44809e-05

1.16645e-05

1.12e-05

1.16164e-05

0.001946509
0.0009732545

0.000961154

1.21005e-05

0.00056059444
0.00014014861

0.00042044583
0.00014014861

0.00021231694
0.00010615847

1.76933e-05

1.68465e-05

7.6105e-06

9.23101e-06

4.65419e-05

8.23526e-06

2.255684e-05
4.511368e-05

7.71084e-06

1.4846e-05

1.14333e-05
2.28666e-05

1.14333e-05

1.83428e-05
3.66856e-05

1.83428e-05

0.00017361775
3.472355e-05

9.9724e-05
2.4931e-05

2.4931e-05
7.4793e-05

2.4931e-05
4.9862e-05

2.4931e-05

3.91702e-05
9.79255e-06

2.937765e-05
9.79255e-06

9.79255e-06
1.95851e-05

9.79255e-06

9.12767e-06

0.0035320600791
0.00058867667985

0.00294338339925
0.00058867667985

0.00077610543
0.0001940263575

0.00011709798
3.903266e-05

3.903266e-05
7.806532e-05

6.38449e-06

5.6611e-06

6.95277e-06

2.00343e-05

0.0001549936975
0.0004649810925

8.0623455e-05
4.03117275e-05

1.11015e-06

1.64122e-06

1.38048e-05

6.61263e-07

2.22782e-05

7.92671e-07

2.34235e-08

2.813318e-05
5.626636e-05

5.63978e-06

1.23602e-05

1.01332e-05

5.219918e-05
0.00010439836

4.37595e-05

8.43968e-06

3.434961e-05
6.869922e-05

6.86066e-06

6.22694e-06

1.23936e-05

8.86841e-06

0.0007918570324
0.0001979642581

0.000164529
5.4843e-05

6.51394e-05
3.25697e-05

1.88456e-05

1.37241e-05

1.04772e-05
2.09544e-05

1.04772e-05

1.17961e-05
2.35922e-05

1.17961e-05

0.0001431212581
0.0004293637743

6.02726e-05
3.01363e-05

1.11354e-05

1.90009e-05

8.80138e-06
4.40069e-06

4.40069e-06

2.168703e-05
4.337406e-05

8.91273e-06

1.27743e-05

0.0001061922
5.30961e-05

1.45374e-05

3.85587e-05

1.89840381e-05
3.79680762e-05

6.49381e-08

1.89191e-05

1.48171e-05
2.96342e-05

5.76771e-06

9.04939e-06

5.835484e-05
1.458871e-05

6.13045e-06
1.839135e-05

1.22609e-05
6.13045e-06

6.13045e-06

2.537478e-05
8.45826e-06

1.691652e-05
8.45826e-06

8.45826e-06

0.00072835904
0.00018208976

0.00018208976
0.00054626928

3.300428e-05
6.600856e-05

6.22856e-06

1.75956e-05

9.18012e-06

3.861258e-05
1.930629e-05

6.65869e-06

1.26476e-05

0.00012977919
0.00025955838

5.16173e-05

1.29198e-05

9.97982e-06

6.37601e-06

2.64915e-05

7.38206e-06

1.50127e-05

7.59425e-09
3.0377e-08

7.59425e-09
2.278275e-08

1.51885e-08
7.59425e-09

7.59425e-09
